# Multi-site co-mutations and 5’UTR CpG immunity escape drive the evolution of SARS-CoV-2

**DOI:** 10.1101/2020.07.21.213405

**Authors:** Jingsong Zhang, Junyan Kang, Mofang Liu, Benhao Han, Li Li, Yongqun He, Zhigang Yi, Luonan Chen

## Abstract

The SARS-CoV-2 infected cases and the caused mortalities have been surging since the COVID-19 pandemic. Viral mutations emerge during the virus circulating in the population, which is shaping the viral infectivity and pathogenicity. Here we extensively analyzed 6698 SARS-CoV-2 whole genome sequences with specific sample collection dates in NCBI database. We found that four mutations, i.e., 5’UTR_c-241-t, NSP3_c-3037-t, NSP12_c-14408-t, and S_a-23403-g, became the dominant variants and each of them represented nearly 100% of all virus sequences since the middle May, 2020. Notably, we found that co-occurrence rates of three significant multi-site co-mutational patterns, i.e., (i) S_a-23403-g, NSP12_c-14408-t, 5’UTR_c-241-t, NSP3_c-3037-t, and ORF3a_c-25563-t; (ii) ORF8_t-28144-c, NSP4_c-8782-t, NSP14_c-18060-t, NSP13_a-17858-g, and NSP13_c-17747-t; and (iii) N_g-28881-a, N_g-28882-a, and N_g-28883-c, reached 66%, 90%, and nearly 100% of recent sequences, respectively. Moreover, we found significant decrease of CpG dinucleotide at positions 241(c)-242(g) in the 5’UTR during the evolution, which was verified as a potential target of human zinc finger antiviral protein (ZAP). The four dominant mutations, three significant multi-site co-mutations, and the potential escape mutation of ZAP-target in 5’UTR region contribute to the rapid evolution of SARS-CoV-2 virus in the population, thus shaping the viral infectivity and pathogenicity. This study provides valuable clues and frameworks to dissect the viral replication and virus-host interactions for designing effective therapeutics.

**One Sentence Summary:** Four dominant mutations, three significant multi-site co-mutations, and 5’UTR CpG escape contribute to the rapid evolution of SARS-CoV-2 virus.

## Main Text

Since the outbreak of COVID-19 in December 2019, it has been pandemic in over 200 countries. The infected cases and the mortalities have been surging, which is an ongoing threat to the public health (*1, 2*). COVID-19 is caused by infection with a novel coronavirus SARS-CoV-2 (*3–5*). Even though as a coronavirus, SARS-CoV-2 has genetic proofreading mechanisms (*6–8*), the persistent natural selective pressure in the population drives the virus to gradually accumulate favorable mutations (*6, 9*). Considerable attention is given to the mutation and evolution of SARS-CoV-2, for that viral mutations have important impact on the infection and pathogenicity of viruses (*10*). The beneficial mutants can better evolve and adapt to host (*9*), either strengthening or weakening the infectivity and pathogenicity. In addition, the variants may generate drug resistance and shrink the efficacy of vaccine and therapeutics (*11, 12*). Dissecting the evolutional trajectory of the virus in the population provides important clues to understand the viral replication and virus-host interactions and helps designing effective therapeutics.

In this study, we used a NCBI dataset consisting of 6698 high-quality SARS-CoV-2 whole genome sequences with sample collection dates ranging from Dec. 20, 2019 to Jun. 8, 2020. By extensive sequence analysis, we identified the significant and convergent features of the accumulated viral mutations and CpG variations over time. Specifically, in the 29903nt viral genome, four significant mutations, i.e., 5’UTR_c-241-t, NSP3_c-3037-t, NSP12_ c-14408-t, and S_ a-23403-g, were found to become the dominant variants since early March, 2020, and each of them reached almost 100% of all virus sequences. By global statistical analyzing, we identified 14 mutation sites with significant high rates. In addition, we evaluated the mutation trajectories by each day and every 10 days, and notably identified three co-mutation patterns consisting of these 13 sites (among these 14 sites) with surprisingly high co-occurrence rates. Moreover, we found the significant decrease of CpG dinucleotides in the viral genome over time, suggesting an evolutional escape of host innate immunity of CpG (*13–15*). The dissected evolution trajectory that the four dominant mutations, three significant multi-site co-mutations, and CpG (decrease) mutation contribute to the rapid evolution of SARS-CoV-2 virus in the population, which shapes the viral infectivity and pathogenicity. This study provides valuable clues and frameworks to dissect the viral replication and virus-host interactions for designing effective therapeutics.

## RESULTS

### Dominant mutations appeared in SARS-CoV-2 in COVID-19 population over time

To explore the mutational landscape of SARS-CoV-2 during virus circulating in the COVID-19 population since the outbreak of COVID-19, we aligned 6698 high quality fulllength genome sequences across all major regions with viral sample collection dates ranging from Dec. 20, 2019 to Jun. 8, 2020 (table S1). As the mutation landscape was massive up to date, we identified 82 mutation sites with mutation rate >1% to draw a heatmap (Fig. 1A). As shown in Fig. 1, the Y-axis represents the collection dates of COVID-19 samples, each of which contains 1~225 sequences. According to the first posted viral sequence (NC_045512), there were accumulated mutations during the virus circulating and apparently new mutation sites gradually emerged since the end of Feb. 2020. Noted that several highly mutated sites appeared before Feb. 22, 2020, which was likely due to the limited collected sequences available at that time and accidental random mutations, i.e., a high mutation rate resulted from even one mutation site. Up to now, there were at least four dominant mutations (5’UTR_c-241-t, NSP3_c-3037-t, NSP12_c-14408-t, and S_a-23403-g) (Fig. 1A), where S_ a-23403-g mutation resulted in the amino acid change (D614G) that enhances viral infectivity (*6, 16*), albeit debate exists (*10*). In particular, each of them covered almost 100% of all virus sequences since the middle May 2020. Focusing on eight mutation sites (5’UTR_c-241-t, NSP3_c-3037-t, NSP12_ c-14408-t, S_a-23403-g, ORF3a_g-25563-t, N_ g-28881-a, N_ g-28882-a, and N_g-28883-c), all the sites sites began to have very high mutation rates since May 2020 (Fig. 1B to 1F). Notably, mutations in three adjacent sites in N (N_g-28881-a, N_g-28882-a, and N_g-28883-c) co-occurred (Fig. 1G-I), suggesting a strong selection pressure.

**Fig. 1.**
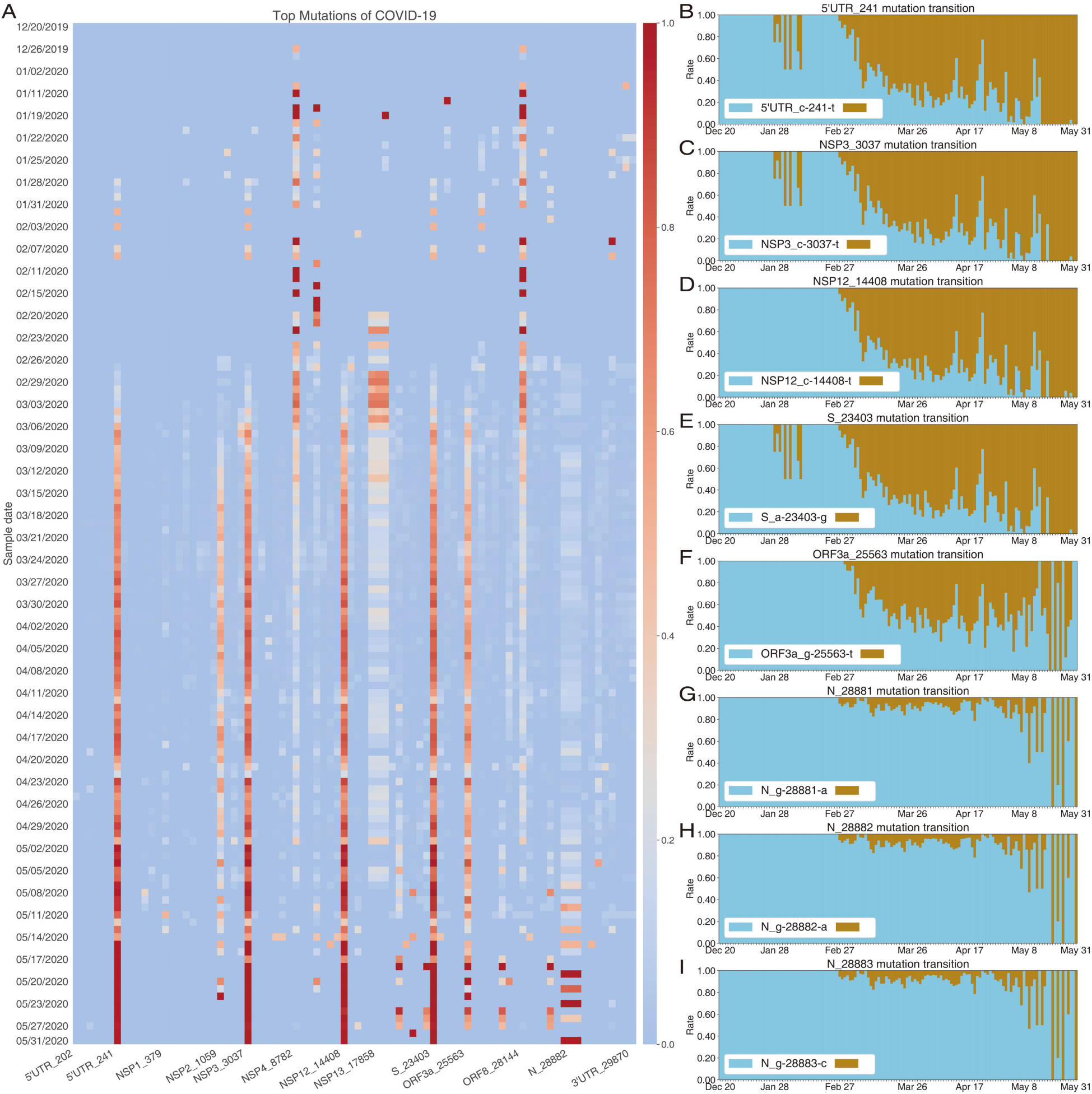
Ongoing and dominant mutations of SARS-CoV-2 over time. **(A)** The global mutational landscape of top 1% mutation rates from Dec. 20, 2019 to May 31, 2020. The mutation of SARS-CoV-2 is clearly ongoing and yet with a rapid rate. **(B)-(I)** show the mutation transitions of 8 sites from Dec. 20, 2019 to May 31, 2020. The sky-blue represents the rates of the original nucleic acids in reference sequence, and the dark-golden the rates of the mutant nucleic acids. Note that the 5 mutations (subfigures B to F), i.e., 5’UTR_c-241-t, NSP3_c-3037-t, NSP12_c-14408-t, and S_a-23403-g, and ORF3a_c-25563-t, especially the top 4 ones have clearly become the dominant mutants. The three adjacent mutations (subfigures G to I, N_ g-28881-a, N_ g-28882-a, and N_g-28883-c) increase daily on the whole. The transitions of the six other significant mutation sites are shown in figs. S1F and figs. S2A-E.

### Strong co-occurrent mutations appeared on multiple sites over time

We assessed the mutations of all residues of the SARS-CoV-2 based on the collected genome sequences. The top 34 mutation sites (with mutation rate>2%) were listed in Fig. 2A. Clearly, there were four dominant mutants (S_a-23403-g, NSP12_c-14408-t, 5’UTR_c-241-t, and NSP3_c-3037-t). The three adjacent sites in Nucleocapsid (N) also had considerably high mutation rates (>0.08). By analyzing the top 34 mutations, there were biased mutation patterns, e.g. the ratio of c-to-t (c-t) was more than 44% (Fig. 2B). We then studied the global cooccurrence relationships of the 34 mutations. We found that there were strong co-occurrence site pairs/associations (Fig. 2C) such as the following three multi-site patterns (i) S_a-23403-g, NSP12_c-14408-t, 5’UTR_c-241-t, NSP3_c-3037-t, ORF3a_c-25563-t, and ORF3a_g-25563-t; (ii) ORF8_t-28144-c, NSP4_c-8782-t, NSP14_c-18060-t, NSP13_a-17858-g, and NSP13_c-17747-t; and (iii) N_g-28881-a, N_g-28882-a, and N_g-28883-c. We further quantified the cooccurrence significance (the ratio of co-occurrence mutants to all sequence examples, also called Support (*17–19*) in data mining) among the top 11 mutation sites (mutation rate>10%) (Fig. 2E to 2I). The Y-axes represented the co-occurrence site pairs. For simplicity, we used gene names instead of their mutation sites in Fig. 2E to 2I. The corresponding mutational positions of genes were given in Fig. 2E in details. From Fig. 2F, every mutational association was significant (>0.10) in all collected genome sequences and almost all associations (except NSP4-NSP13a pair) met strong co-occurrence relationships (>0.60). Figs. 2H–2J show 3-to-6 mutation-site (multi-site) co-occurrences. Interestingly, all mutational associations in Fig. 2H to 2J follow significant and strong co-occurrence relationships. The significant co-occurrences of multi-site mutations may suggest that these sites closely interact with each other during the evolution.

**Fig. 2.**
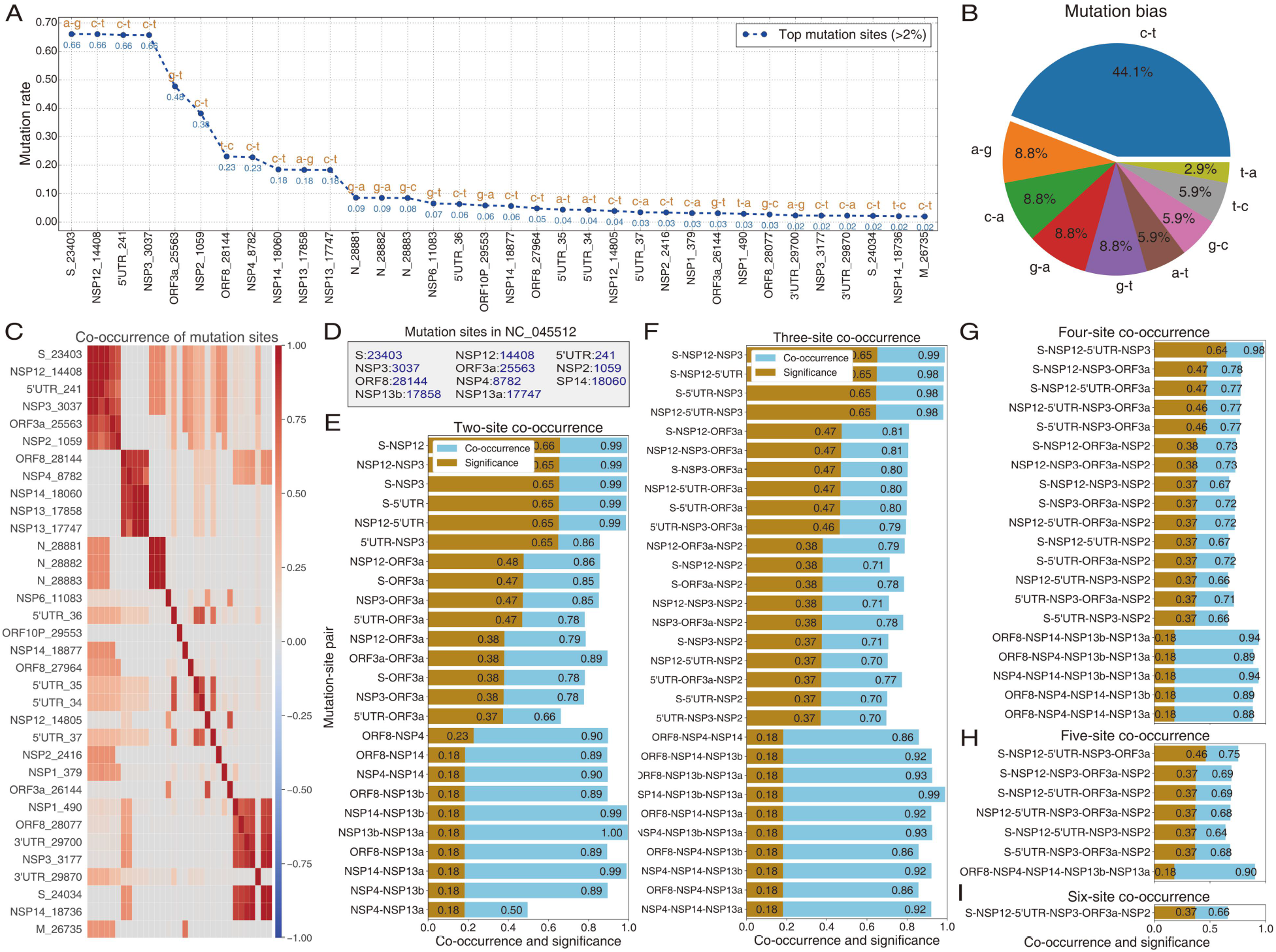
Co-ocurrence mutations of SARS-CoV-2 over time. **(A)** Top 2% mutation sites involve 34 sties. The top 11 sites hold significant high mutation rates. **(B)** Mutation bias of the top 34 mutants. **(C)** Co-occurrence heatmap of top 2% mutation sites in order of mutational significance. The X-axis and Y-axis share the same tick labels (mutation sites along with their positions in reference sequence NC_045512) as shown in Y-axis. The top 14 sites were clustered into three high co-occurrence groups. **(D)** Query table of top 11 mutation sites. We used gene names instead of their mutation sites in subfigures (E) to (I), for simplicity. **(E)** Co-occurrences of 2 mutation sites. Each tick label of Y-axis, like S-NSP12, represents the mutation-site association/pattern of gene S (position 23403) and NSP12 (position 14408). The detailed positions of these mutations are shown in subfigures (E). **(F)** Co-occurrences of 3 mutation sites. **(G)** Co-occurrences of 4 mutation sites. **(H)** Co-occurrences of 5 mutation sites. **(I)** Cooccurrences of 6 mutation sites.

### Co-evolution of multi-sites in COVID-19 population

The strong co-occurrence relationships of multi-sites in COVID-19 virus suggested co-mutational evolution. We then investigated the co-occurrences of three groups involving 14 sites on a time scale of about per ten days. As shown in Fig. 3A, the mutation rates of the first group containing 6 sites (S_23403, NSP12_14408, 5’UTR_241, NSP3_3037, ORF3a_25563, ORF3a_25563, and NSP2_1059) exhibited very similar trends. Strikingly, 4 mutant sites (S_23403, NSP12_14408, 5’UTR_241, and NSP3_3037) almost shared a same mutation rate curve. A second group involved 5 sites (ORF8_28144, NSP4_8782, NSP14_18060, NSP13_17858, and NSP13_17747) and two mutant sites (ORF8_28144 and NSP4_8782) shared a same mutation rate curve whereas the other three mutant sites (NSP14_18060, NSP13_17858, and NSP13_17747) almost had a same mutation rate curve (Fig. 3B). We analyzed the correlations of the 11 mutant sites of the first and the second groups on a 15 intervals by Pearson Correlation Coefficient (PCC) (*20–22*), and expressed them by heatmap (Fig. 3C). There were two red (6*6 and 5*5) regions that corresponded to the first and the second mutant groups, respectively. As expected, the top 5 mutation sites, especially the top 4 sites in the first group exhibited a very strong correlation. In the second red (5*5) region corresponded to the second group, there were two subgroups containing a two-site (ORF8_28144 and NSP4_8782) and a triple-site (NSP14_18060, NSP13_17858 and NSP13_17747) exhibited very strong correlations, respectively (Fig. 3C).

**Fig. 3.**
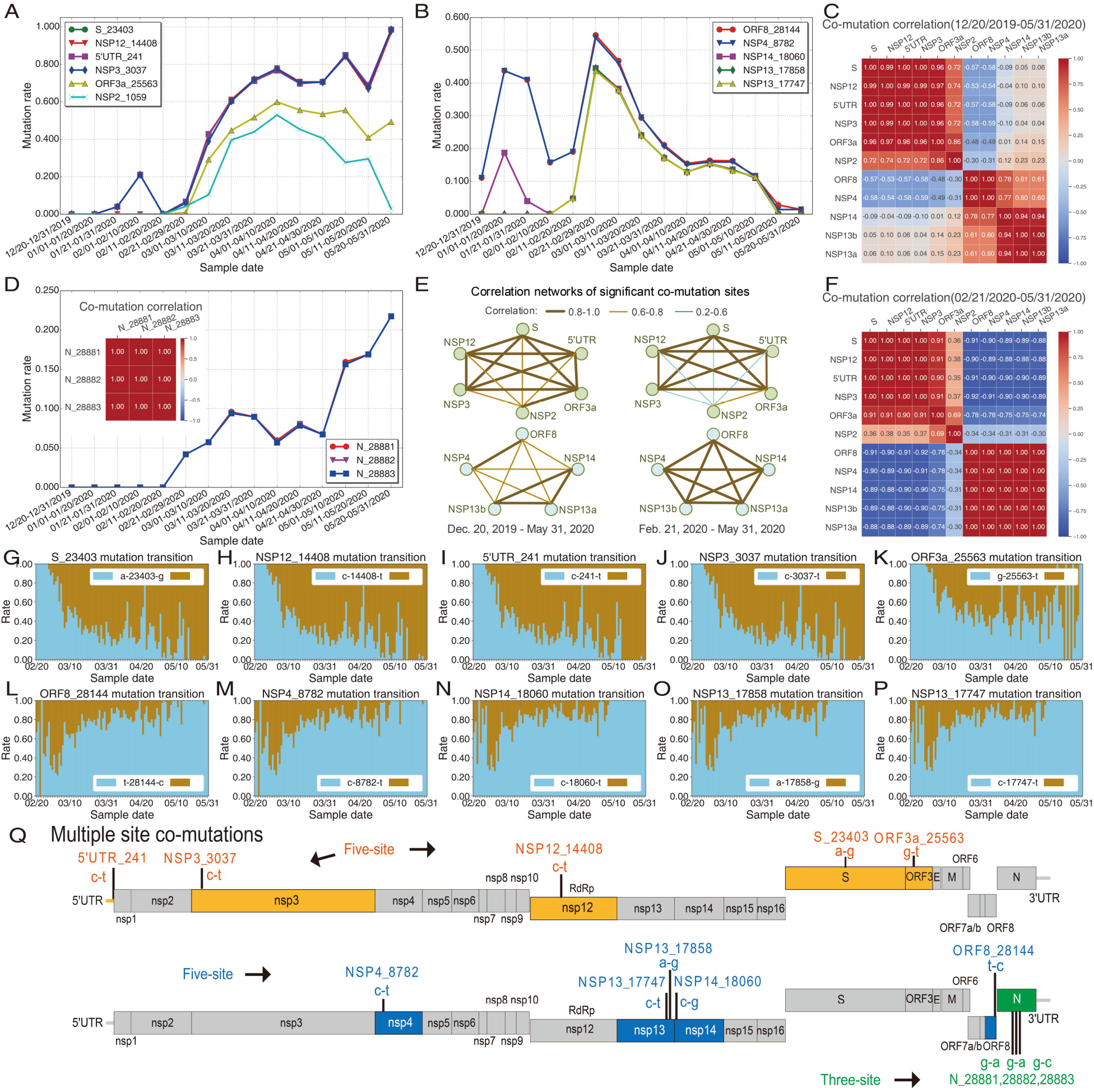
Co-evolution of multi-sites in COVID-19 population over time. **(A)** Mutation trends of top 6 sites in Fig. 2A from Dec. 20, 2019 to May 31, 2020. **(B)** Mutation trends from the 7th to 11th sites in Fig. 2A. **(C)** Global co-mutational heatmap of top 11 in Fig. 2A sites since Dec. 20, 2019. **(D)** Mutation trends and co-mutational heatmap of three adjacent Nucleocapsid sites (from 12th to 14th sites in Fig.2A). **(E)** Correlation networks of significant co-mutation sites. The left two sites show the correlation relationships from Dec. 20, 2019 to May 31, 2020, and the right two from Feb. 21, 2020 to May 31, 2020. **(F)** Co-mutational heatmap of top 11 sites with sample collection dates from Feb. 21, 2020 to May 31, 2020. **(G)-(K)** Mutation transitions of five-site co-mutations in the first group. **(L)-(P)** Mutation transitions of five-site co-mutations of the second group. The mutation transitions of the third group are shown in Fig. 1G-I. **(Q)** The mutational positions of multi-site co-mutations consisting of 5-, 5- and 3-site co-mutations.

Unlike the first and the second groups, the third group consisted of three *adjacent* mutation sites in nucleocapsid region (N_28881, N_28882, and N_28883). The mutation rate curve of these sites almost overlapped with each other and the mutation rates of these sites increased over time (Fig. 3D). The correlations of these sites were nearly 1.0 (Fig. 3D), suggesting a strong coevolution.

Based on the relationships of the top two multi-site mutation groups, we illustrated the correlation networks of these mutations. As shown in Fig. 3E, the correlation networks of the first group mutation sites showed a six-pointed star and that of the second group showed a five-pointed star network. The mutation sites exhibited strong correlations with each other (Fig. 3E) within each star/group. Furthermore, we evaluated the correlations from Feb. 21 to May 31, 2020 and found that five sites (S_23403, NSP12_14408, 5’UTR_241, NSP3_3037, ORF3a_25563, and ORF3a_25563) of the first group and all sites (ORF8_28144, NSP4_8782, NSP14_18060, NSP13_17858, and NSP13_17747) of the second group very strongly correlated with each other within their groups (Fig. 3F, Figs. 3G-P). In short, such three significant multi-site co-mutations involving 13 sites indicated the evolution of SARS-CoV-2 with a rapid rate.

The mutations either changed the amino acid sequence or not. The mutation NSP2_c-1059-t resulted in an amino acid change (T85I) in the NSP2; the mutation NSP12_c-14408-t resulted in an amino acid change in the viral RNA-dependent RNA polymerase NSP12 (P323L) (Fig. 4B); the mutation S_a-23403-g resulted in an amino acid change in Spike protein (S)(D614G), which enhances viral infectivity (*6, 16*); the mutation ORF3a_g-25563-t resulted in an amino acid change (Q57H) in Spike protein in ORF3a; the mutation NSP13_c-17747-t and NSP13_a-17858-g resulted in an amino acid changes P504L and Y541C in the NTPase/helicase domain NSP13, respectively; the mutation ORF8_t-28144-c resulted in an amino acid change (L84S) in the ORF8; the mutations N_g-28881-a, N_g-28882-a, N_g-28883-c resulted amino acid changes R203K, R203S and G203R in the nucleocapsid N, respectively (Fig. 3P). In contrast, the mutations NSP14_c-18060-t and NSP3_c-3037-t were synonymous mutations without amino acid changes (Fig. 3P). The mutation 5’UTR_c-241-t located in the 5’-non-translated region and did not change the predicted RNA structure of 5’UTR (Fig. 4A).

**Fig. 4.**
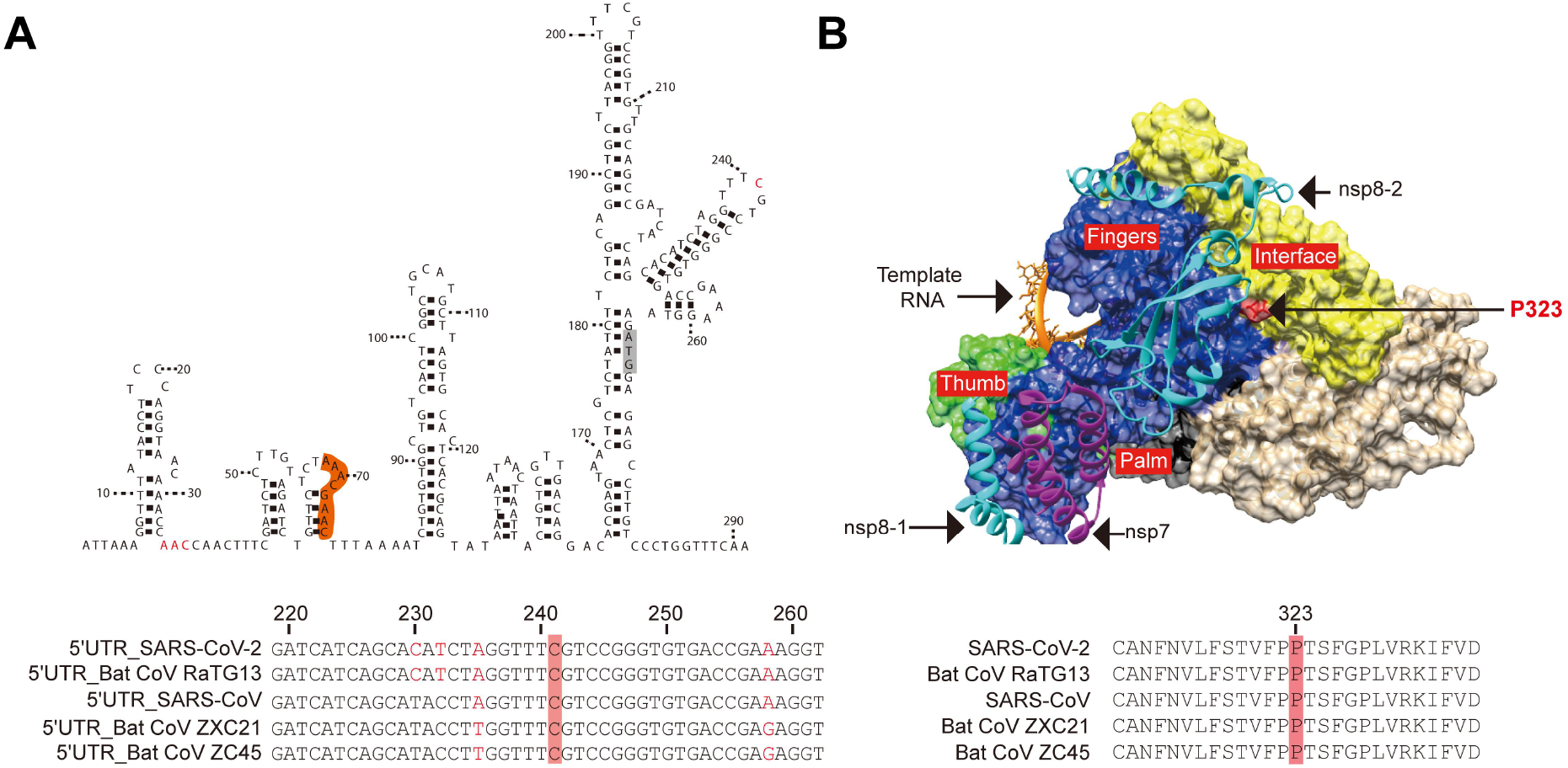
Structure of 5’UTR and SARS-CoV-2 RdRp/RNA complex. **(A)** Predicted RNA structure of the SARS-CoV-2 5’UTR. RNA structure of the 400-nt 5’UTR was predicted by “RNAstructure” (http://rna.urmc.rochester.edu/RNAstructureWeb). The start codon for nsp1 was in grey. The TRS-L was in orange. The mutated nucleotides were in red. The bottom panel, alignment of the 5’UTR of SARS-CoV-2 with 5’UTRs of related viruses. The c241 was highlighted. **(B)** Structure of SARS-CoV-2 RdRp/RNA complex. The structure of SARS-CoV-2 RdRp/RNA complex (PDB, 6X2G) was visualized by Chimera (UCSF). The P323 mutation (in red) was indicated. The alignment of the amino acid sequences of SARS-CoV-2 and related viruses near the P323 position. The P323 was highlighted.

### Significant decrease of CpG dinucleotide content in 5’UTR in COVID-19 population over time

The CpG content of viral genome is restricted by host intrinsic zinc finger antiviral protein that interacts with CpG rich-region and mediates depletion of foreign viral RNAs (*14, 23*). Comparing with other coronaviruses, SARS-CoV-2 genome exhibits extreme CpG deficiency (*13*). However, the evolutional trajectory of SARS-CoV-2 CpG-content within the same species is still unclear. We investigated the CpG-content changes in SARS-CoV-2 since the outbreak. As shown in Fig. 5A and 5B, the CpG dinucleotide content exhibited a decreased trend over time. The CpG-content in each SARS-CoV-2 genome regions varied, with high CpG-contents the 5’UTR, NSP1, E and ORF10 regions and low CpG-contents in NSP8, ORF6 regions. Notably, the NSP7 region was free of CpG dinucleotide (Fig. 5C). Comparing with the first posted SARS-CoV-2 genome (NC_045512), in the very recent SARS-CoV-2 genomes, only the CpG-contents of 5’UTR decreased significantly but not the other CpG high content regions NSP1, E and ORF10 (Fig. 5D, E), suggesting a biased evolution pressure on this region.

**Fig. 5.**
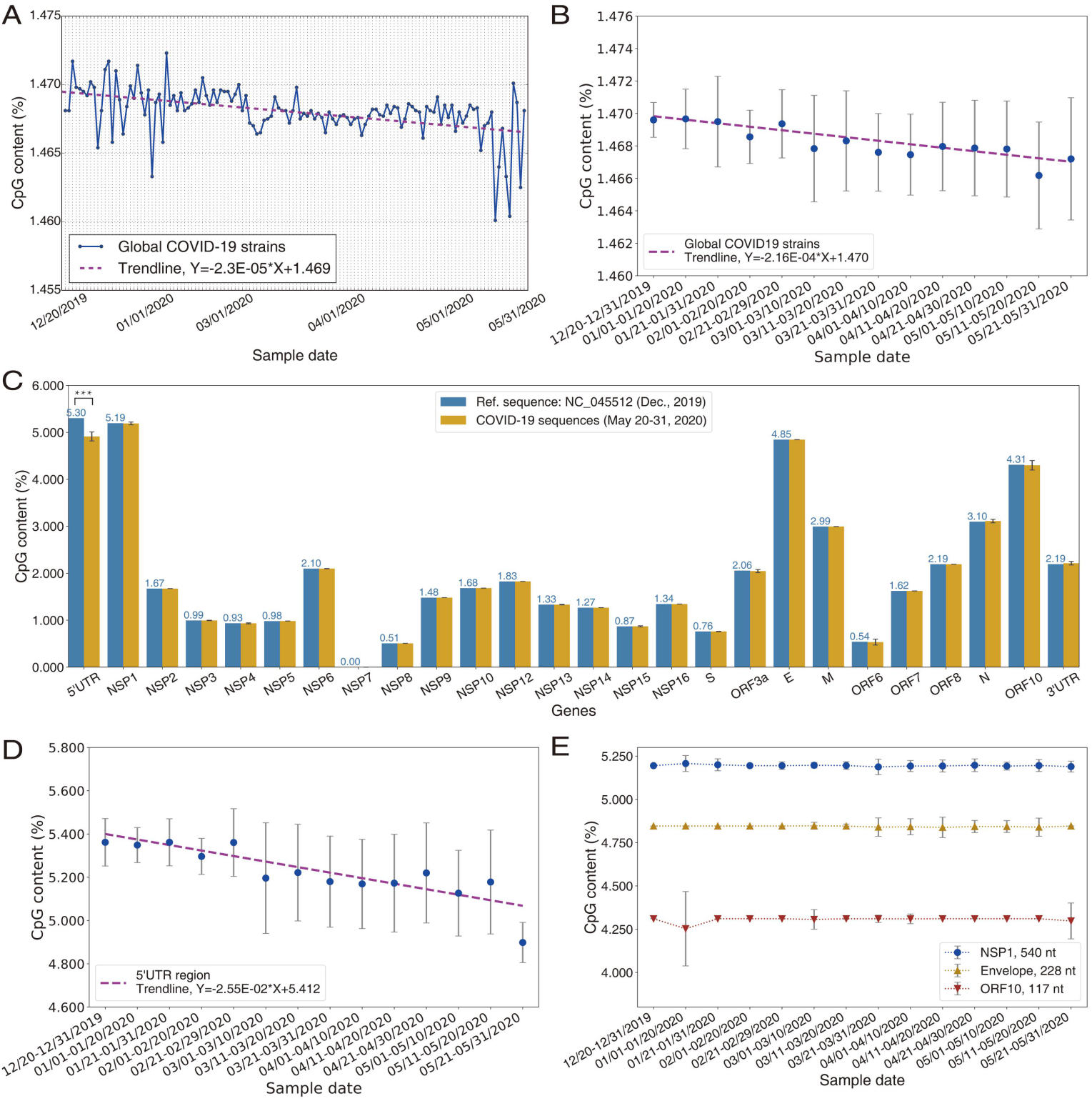
CpG-content decrease in COVID-19 population over time. **(A)** Trend of CpG decrease of COVID-19 genome per day. **(B)** Trend of CpG decrease of COVID-19 genome by intervals of about 10 days. **(C)** CpG changes of all genes. The significant decrease of only 5’UTR region indicates that 5’UTR is the potential target gene of ZAP. **(D)** Trend of CpG decrease of 5’UTR region by about 10 days. **(E)** CpG change trends of NSP1, Envelope, and ORF10.

## CONCLUSION

Our comprehensive and massive mutation and correlation analyses identified four dominant mutation sites (5’UTR_c-241-t, NSP3_c-3037-t, NSP12_c-14408-t, and S_a-23403-g) and revealed three significant multi-site co-mutational patterns (S_a-23403-g, NSP12_c-14408-t, 5’UTR_c-241-t, NSP3_c-3037-t, ORF3a_c-25563-t; ORF8_t-28144-c, NSP4_c-8782-t, NSP14_c-18060-t, NSP13_a-17858-g, NSP13_c-17747-t; and N_g-28881-a, N_g-28882-a, N_g-28883-c). Some of the mutations changed the amino acid sequence in the viral RNA-dependent RNA polymerase nsp12 (P323L), Spike protein (S) (D614G), ORF3a (Q57H), NTPase/helicase domain nsp13 (P504L, Y541C), ORF8 (L84S) and nucleocapsid N (R203K, R203S, G203R), or did not change the amino acid sequence (NSP14_c-18060-t and NSP3_c-3037-t), which may affect viral replication or virus-host interaction. Other mutations were synonymous mutations without amino acid changes (Fig. 3G). And mutations located in the 5’-non-translated region (5’UTR_c-241-t) that did not change the predicted RNA structure. Moreover, we found gradual but significant decrease of CpG-content in 5’UTR region over time, which suggested the 5’UTR region as a potential ZAP target. Taken together, our study provides valuable clues and frameworks to dissect the viral replication and virus-host interactions for designing effective therapeutics.

## Acknowledgments

We thank associate prof. T.Z. and Dr. S.T.H. for useful comments on the manuscript.

## Author contributions

L.N.C. and J.S.Z. designed the study. J.S.Z. and Z.G.Y. designed the experiments. J.S.Z. analyzed data. J.S.Z., Z.G.Y., and J.Y.K. designed the figures. H.B.H. checked the experiments. J.S.Z. wrote the manuscript. Z.G.Y., J.Y.K., M.F.L., L.L., and Y.Q.H polished the manuscript.

## Data and materials availability

All data are available in the main text or supplementary materials or from the corresponding author upon request.

## Supplementary Materials

Figs. S1 and S3

Table. S1

