## Supplementary material for "Multi-site co-mutations and 5’UTR CpG immunity escape drive the evolution of SARS-CoV-2": SI02

**This PDF file includes:**

Figs. S1-S3

Table S1

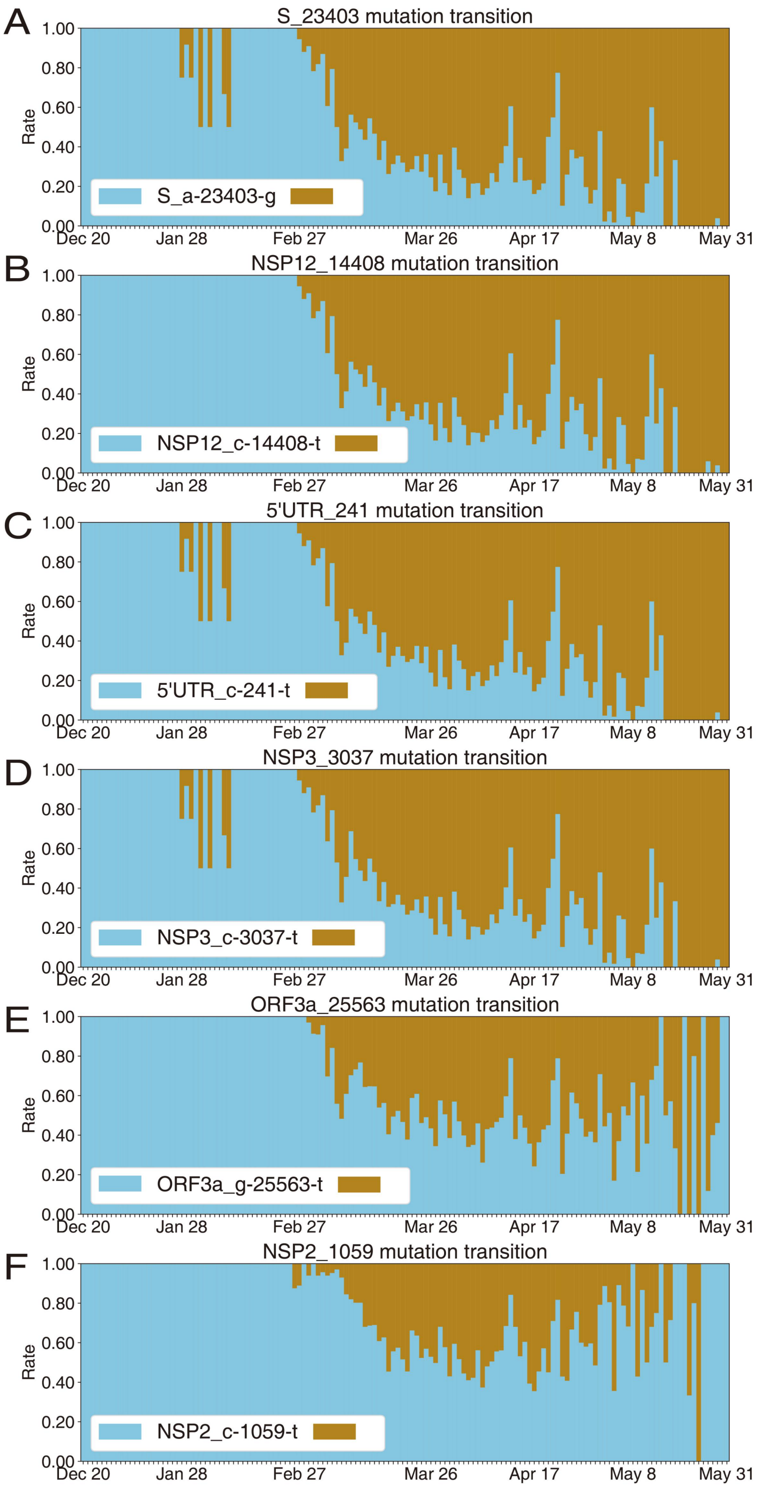


**Figure S1. Mutation transitions of the top 6 mutant sites.** **(A)-(D)** Four sites, i.e., 5'UTR_c-241-t, NSP3_c-3037-t, NSP12_c-14408-t, and S_a-23403-g, became the dominant variants and each of them represented nearly 100% of all virus sequences since the middle May, 2020. **(E)-(F)** Two sites (ORF3a_g-25563-t and NSP2_C-1059-T) overall increased since the end of Jan. 2020.

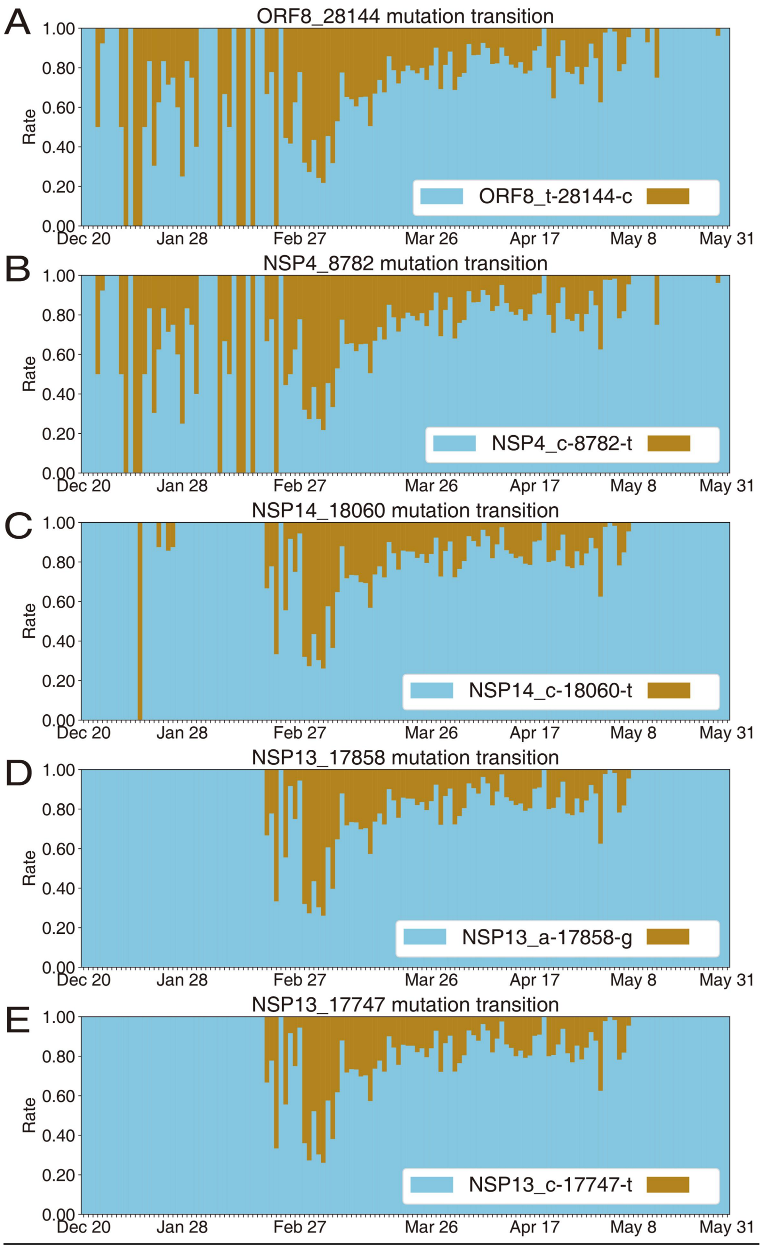


**Figure S2. Mutation transitions of the 7th to 11th mutant sites. (A)-(E)** These mutation sites over decreased since the end of Feb. 2020.

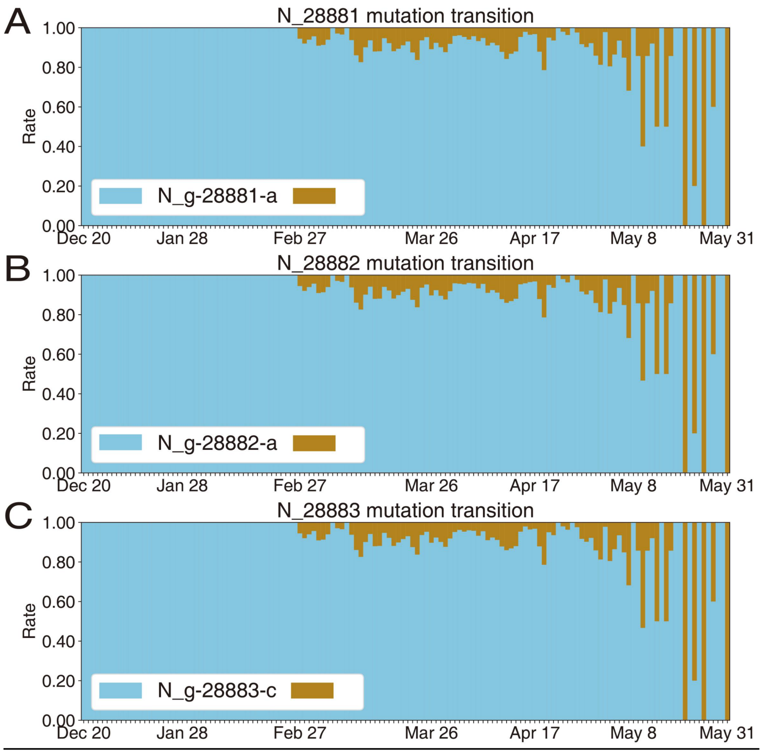


**Figure S3. Mutation transitions of the 12th to 14th mutant sites. (A)-(C)** These three adjacent mutation sites overall share a same increase trend.

**Table S1. Dataset.** The dataset consists of 6698 SARS-CoV-2 whole genome sequences with specific sample collection dates in NCBI database.

| Accession | Collection Date | Locality |
|---|---|---|
| NC_045512.2 | 12/2019 | China |
| MN908947.3 | 12/20/2019 | China |
| MT019529.1 | 12/23/2019 | China:Hubei-Wuhan |
| LR757995.1 | 12/26/2019 | China:Wuhan |
| LR757998.1 | 12/26/2019 | China:Wuhan |
| MN996527.1 | 12/30/2019 | China:Wuhan |
| MN996528.1 | 12/30/2019 | China:Wuhan |
| MN996529.1 | 12/30/2019 | China:Wuhan |
| MN996530.1 | 12/30/2019 | China:Wuhan |
| MN996531.1 | 12/30/2019 | China:Wuhan |
| MT019530.1 | 12/30/2019 | China:Hubei-Wuhan |
| MT019531.1 | 12/30/2019 | China:Hubei-Wuhan |
| MT019532.1 | 12/30/2019 | China:Hubei-Wuhan |
| MT291826.1 | 12/30/2019 | China:Wuhan |
| MT291827.1 | 12/30/2019 | China:Wuhan |
| MT291828.1 | 12/30/2019 | China:Wuhan |
| MT291829.1 | 12/30/2019 | China:Wuhan |
| MT291830.1 | 12/30/2019 | China:Wuhan |
| LR757996.1 | 01/01/2020 | China:Wuhan |
| MT019533.1 | 01/01/2020 | China:Hubei-Wuhan |
| MN988668.1 | 01/02/2020 | China |
| MN988669.1 | 01/02/2020 | China |
| MT093631.2 | 01/08/2020 | China |
| MT447155.1 | 01/08/2020 | Thailand |
| MN938384.1 | 01/10/2020 | China:Shenzhen |
| MT259226.1 | 01/10/2020 | China:Hubei-Wuhan |
| MN975262.1 | 01/11/2020 | China |
| MT072688.1 | 01/13/2020 | Nepal |
| MT049951.1 | 01/17/2020 | China:Yunnan |
| MN985325.1 | 01/19/2020 | USA |
| MT233526.1 | 01/19/2020 | USA:WA |
| MT246667.1 | 01/19/2020 | USA:WA |
| MT039873.1 | 01/20/2020 | China:Hangzhou |
| MT226610.1 | 01/20/2020 | China |
| MN988713.1 | 01/21/2020 | USA:Illinois |
| MT253701.1 | 01/21/2020 | China:Zhejiang-Hangzhou |
| MT253702.1 | 01/21/2020 | China:Zhejiang-Hangzhou |
| MT253708.1 | 01/21/2020 | China:Zhejiang-Hangzhou |
| MT253709.1 | 01/21/2020 | China:Zhejiang-Hangzhou |
| MT253710.1 | 01/21/2020 | China:Zhejiang-Hangzhou |
| MN994468.1 | 01/22/2020 | USA:CA |
| MN997409.1 | 01/22/2020 | USA:AZ |

| | | |
|---|---|---|
| MT039874.1 | 01/22/2020 | China |
| MT079843.1 | 01/22/2020 | China |
| MT079844.1 | 01/22/2020 | China |
| MT079845.1 | 01/22/2020 | China |
| MT079846.1 | 01/22/2020 | China |
| MT079847.1 | 01/22/2020 | China |
| MT079848.1 | 01/22/2020 | China |
| MT079849.1 | 01/22/2020 | China |
| MT079850.1 | 01/22/2020 | China |
| MT079851.1 | 01/22/2020 | China |
| MT079852.1 | 01/22/2020 | China |
| MT079853.1 | 01/22/2020 | China |
| MT079854.1 | 01/22/2020 | China |
| MT192772.1 | 01/22/2020 | Viet-Nam:Ho-Chi-Minh-city |
| MT192773.1 | 01/22/2020 | Viet-Nam:Ho-Chi-Minh-city |
| MT230904.1 | 01/22/2020 | Hong-Kong |
| MT253705.1 | 01/22/2020 | China:Zhejiang-Hangzhou |
| MT253706.1 | 01/22/2020 | China:Zhejiang-Hangzhou |
| MT407649.1 | 01/22/2020 | China:Zhejiang |
| MT407650.1 | 01/22/2020 | China:Zhejiang |
| MT407651.1 | 01/22/2020 | China:Zhejiang |
| MN994467.1 | 01/23/2020 | USA:CA |
| MT114412.1 | 01/23/2020 | Hong-Kong |
| MT253696.1 | 01/23/2020 | China:Zhejiang-Hangzhou |
| MT253697.1 | 01/23/2020 | China:Zhejiang-Hangzhou |
| MT253698.1 | 01/23/2020 | China:Zhejiang-Hangzhou |
| MT447156.1 | 01/23/2020 | Thailand |
| MT447166.1 | 01/23/2020 | Thailand |
| MT622319.1 | 01/23/2020 | China |
| MT114413.1 | 01/24/2020 | Hong-Kong |
| MT114414.1 | 01/24/2020 | Hong-Kong |
| MT253699.1 | 01/24/2020 | China:Zhejiang-Hangzhou |
| MT291831.1 | 01/24/2020 | China:Beijing |
| MT365028.1 | 01/24/2020 | Hong-Kong |
| MT450919.1 | 01/24/2020 | Australia:Victoria |
| MT007544.1 | 01/25/2020 | Australia:Victoria |
| MT020880.1 | 01/25/2020 | USA:WA |
| MT020881.1 | 01/25/2020 | USA:WA |
| MT192759.1 | 01/25/2020 | Taiwan |
| MT253700.1 | 01/25/2020 | China:Zhejiang-Hangzhou |
| MT253703.1 | 01/25/2020 | China:Zhejiang-Hangzhou |
| MT253704.1 | 01/25/2020 | China:Zhejiang-Hangzhou |
| MT253707.1 | 01/25/2020 | China:Zhejiang-Hangzhou |
| MT259230.1 | 01/25/2020 | China:Hubei-Wuhan |
| MT259231.1 | 01/25/2020 | China:Hubei-Wuhan |
| MT291832.1 | 01/25/2020 | China:Beijing |

| | | |
|---|---|---|
| MT447157.1 | 01/25/2020 | Thailand |
| MT447158.1 | 01/25/2020 | Thailand |
| MT450920.1 | 01/25/2020 | Australia:Victoria |
| MT135041.1 | 01/26/2020 | China:Beijing |
| MT259227.1 | 01/26/2020 | China:Hubei-Wuhan |
| MT259228.1 | 01/26/2020 | China:Hubei-Wuhan |
| MT259229.1 | 01/26/2020 | China:Hubei-Wuhan |
| MT407652.1 | 01/26/2020 | China:Zhejiang |
| MT407653.1 | 01/26/2020 | China:Zhejiang |
| MT447159.1 | 01/26/2020 | Thailand |
| MT534630.1 | 01/26/2020 | China:Jiangsu-Changzhou |
| MT044258.1 | 01/27/2020 | USA:CA |
| MT114415.1 | 01/27/2020 | Hong-Kong |
| MT114416.1 | 01/27/2020 | Hong-Kong |
| MT114418.1 | 01/27/2020 | Hong-Kong |
| MT123292.2 | 01/27/2020 | China:Guangzhou |
| MT291835.2 | 01/27/2020 | China:Beijing |
| MT044257.1 | 01/28/2020 | USA:IL |
| MT135042.1 | 01/28/2020 | China:Beijing |
| MT135043.1 | 01/28/2020 | China:Beijing |
| MT135044.1 | 01/28/2020 | China:Beijing |
| MT270101.1 | 01/28/2020 | Germany:Bavaria |
| MT270103.1 | 01/28/2020 | Germany:Bavaria |
| MT291833.1 | 01/28/2020 | China:Beijing |
| MT291834.1 | 01/28/2020 | China:Beijing |
| MT020781.2 | 01/29/2020 | Finland |
| MT027062.1 | 01/29/2020 | USA:CA |
| MT027063.1 | 01/29/2020 | USA:CA |
| MT027064.1 | 01/29/2020 | USA:CA |
| MT039888.1 | 01/29/2020 | USA:MA |
| MT114417.1 | 01/29/2020 | Hong-Kong |
| MT114419.1 | 01/29/2020 | Hong-Kong |
| MT123291.2 | 01/29/2020 | China:Guangzhou |
| MT123293.2 | 01/29/2020 | China:Guangzhou |
| MT270102.1 | 01/29/2020 | Germany:Bavaria |
| MT291836.1 | 01/29/2020 | China:Beijing |
| MT447160.1 | 01/29/2020 | Thailand |
| MT066156.1 | 01/30/2020 | Italy |
| MT270112.1 | 01/30/2020 | Germany:Bavaria |
| MT365029.1 | 01/30/2020 | Hong-Kong |
| MT365032.1 | 01/30/2020 | Hong-Kong |
| MT050493.1 | 01/31/2020 | India:Kerala-State |
| MT066175.1 | 01/31/2020 | Taiwan |
| MT077125.1 | 01/31/2020 | Italy |
| MT365030.1 | 01/31/2020 | Hong-Kong |
| MT450930.1 | 01/31/2020 | Australia:Victoria |

| | | |
|---|---|---|
| MT270106.1 | 02/01/2020 | Germany:Bavaria |
| MT447154.1 | 02/01/2020 | Thailand |
| MT121215.1 | 02/02/2020 | China:Shanghai |
| MT365031.1 | 02/02/2020 | Hong-Kong |
| MT450931.1 | 02/02/2020 | Australia:Victoria |
| MT270107.1 | 02/03/2020 | Germany:Bavaria |
| MT447161.1 | 02/03/2020 | Thailand |
| MT066176.1 | 02/05/2020 | Taiwan |
| MT123290.1 | 02/05/2020 | China:Guangdong-Guangzhou |
| MT446312.1 | 02/05/2020 | China:Guangdong-Guangzhou |
| MT106052.1 | 02/06/2020 | USA:CA |
| MT093571.1 | 02/07/2020 | Sweden |
| MT270110.1 | 02/07/2020 | Germany:Bavaria |
| MT447162.1 | 02/07/2020 | Thailand |
| MT270111.1 | 02/08/2020 | Germany:Bavaria |
| MT450924.1 | 02/08/2020 | Australia:Victoria |
| LC528232.1 | 02/10/2020 | null |
| LC528233.1 | 02/10/2020 | null |
| MT106053.1 | 02/10/2020 | USA:CA |
| MT106054.1 | 02/11/2020 | USA:TX |
| MT510728.1 | 02/13/2020 | China |
| LC534418.1 | 02/14/2020 | Japan |
| MT510727.1 | 02/15/2020 | China |
| MT159705.2 | 02/17/2020 | USA |
| MT159706.2 | 02/17/2020 | USA |
| MT159707.2 | 02/17/2020 | USA |
| MT159708.2 | 02/17/2020 | USA |
| MT159710.2 | 02/17/2020 | USA |
| MT159717.2 | 02/17/2020 | USA |
| MT184911.1 | 02/17/2020 | USA |
| MT184912.1 | 02/17/2020 | USA |
| MT159713.2 | 02/18/2020 | USA |
| MT159714.2 | 02/18/2020 | USA |
| MT159718.2 | 02/18/2020 | USA |
| MT159719.2 | 02/18/2020 | USA |
| MT184907.2 | 02/18/2020 | USA |
| MT184910.1 | 02/18/2020 | USA |
| MT159709.2 | 02/20/2020 | USA |
| MT159711.2 | 02/20/2020 | USA |
| MT598633.1 | 02/20/2020 | USA:King-County-WA |
| MT159720.2 | 02/21/2020 | USA |
| MT159721.2 | 02/21/2020 | USA |
| MT159722.2 | 02/21/2020 | USA |
| MT184908.2 | 02/21/2020 | USA |
| MT184909.2 | 02/21/2020 | USA |
| MT476384.1 | 02/21/2020 | USA:FL |

| | | |
|---|---|---|
| MT627270.1 | 02/21/2020 | USA:Washington-King-County |
| MT641645.1 | 02/21/2020 | Australia:Northern-Territory |
| MT641646.1 | 02/21/2020 | Australia:Northern-Territory |
| MT447163.1 | 02/22/2020 | Thailand |
| MT627288.1 | 02/22/2020 | USA:Washington-King-County |
| MT627630.1 | 02/22/2020 | USA:Washington-King-County |
| MT118835.1 | 02/23/2020 | USA:CA |
| MT152824.1 | 02/24/2020 | USA:Snohomish-County-WA |
| MT159715.2 | 02/24/2020 | USA |
| MT159716.2 | 02/24/2020 | USA |
| MT184913.1 | 02/24/2020 | USA |
| MT215194.1 | 02/24/2020 | Hong-Kong |
| MT412134.1 | 02/24/2020 | China |
| MT598636.1 | 02/24/2020 | USA:WA |
| MT598638.1 | 02/24/2020 | USA:King-County-WA |
| MT632586.1 | 02/24/2020 | USA:Washington-Snohomish-County |
| MT641648.1 | 02/24/2020 | Australia:Northern-Territory |
| MT159712.2 | 02/25/2020 | USA |
| MT215195.1 | 02/25/2020 | Hong-Kong |
| MT447164.1 | 02/25/2020 | Thailand |
| MT447165.1 | 02/25/2020 | Thailand |
| MT568634.1 | 02/25/2020 | China:Guangzhou |
| MT568635.1 | 02/25/2020 | China:Guangzhou |
| MT568637.1 | 02/25/2020 | China:Guangzhou |
| MT568639.1 | 02/25/2020 | China:Guangzhou |
| MT568640.1 | 02/25/2020 | China:Guangzhou |
| MT568641.1 | 02/25/2020 | China:Guangzhou |
| MT582498.1 | 02/25/2020 | Germany:Heinsberg |
| MT632542.1 | 02/25/2020 | USA:Washington-King-County |
| MT233520.1 | 02/26/2020 | Spain:Valencia |
| MT276324.1 | 02/26/2020 | USA:CA |
| MT370516.1 | 02/26/2020 | Taiwan |
| MT582495.1 | 02/26/2020 | Germany:Heinsberg |
| MT582496.1 | 02/26/2020 | Germany:Heinsberg |
| MT582497.1 | 02/26/2020 | Germany:Heinsberg |
| MT627263.1 | 02/26/2020 | USA:Washington-King-County |
| MT627627.1 | 02/26/2020 | USA:Washington-King-County |
| MT163716.1 | 02/27/2020 | USA:WA |
| MT233519.1 | 02/27/2020 | Spain:Valencia |
| MT233521.1 | 02/27/2020 | Spain:Valencia |
| MT276325.2 | 02/27/2020 | USA:WA |
| MT276328.2 | 02/27/2020 | USA:OR |
| MT292580.1 | 02/27/2020 | Spain |
| MT292581.1 | 02/27/2020 | Spain |
| MT304474.1 | 02/27/2020 | South-Korea |
| MT370517.1 | 02/27/2020 | Taiwan |

| | | |
|---|---|---|
| MT370518.1 | 02/27/2020 | Taiwan |
| MT419830.1 | 02/27/2020 | USA:CA |
| MT419831.1 | 02/27/2020 | USA:CA |
| MT419832.1 | 02/27/2020 | USA:CA |
| MT568636.1 | 02/27/2020 | China:Guangzhou |
| MT582492.1 | 02/27/2020 | Germany:Heinsberg |
| MT582493.1 | 02/27/2020 | Germany:Heinsberg |
| MT582494.1 | 02/27/2020 | Germany:Heinsberg |
| MT627276.1 | 02/27/2020 | USA:Washington |
| MT126808.1 | 02/28/2020 | Brazil |
| MT163717.1 | 02/28/2020 | USA:WA |
| MT276323.1 | 02/28/2020 | USA:RI |
| MT276329.1 | 02/28/2020 | USA:FL |
| MT276330.2 | 02/28/2020 | USA:FL |
| MT419829.1 | 02/28/2020 | USA:CA |
| MT527178.1 | 02/28/2020 | Italy:Lazio |
| MT582490.1 | 02/28/2020 | Germany:Heinsberg |
| MT582491.1 | 02/28/2020 | Germany:Heinsberg |
| MT582499.1 | 02/28/2020 | Germany:Heinsberg |
| MT598634.1 | 02/28/2020 | USA:WA |
| MT627219.1 | 02/28/2020 | USA:Washington-King-County |
| MT627226.1 | 02/28/2020 | USA:Washington-King-County |
| MT627235.1 | 02/28/2020 | USA:Washington-King-County |
| MT627252.1 | 02/28/2020 | USA:Washington-King-County |
| MT627253.1 | 02/28/2020 | USA:Washington-King-County |
| MT627258.1 | 02/28/2020 | USA:Washington-Snohomish-County |
| MT627274.1 | 02/28/2020 | USA:Washington-King-County |
| MT627275.1 | 02/28/2020 | USA:Washington |
| MT627286.1 | 02/28/2020 | USA:Washington-Snohomish-County |
| MT627290.1 | 02/28/2020 | USA:Washington-King-County |
| MT627295.1 | 02/28/2020 | USA:Washington-King-County |
| MT627318.1 | 02/28/2020 | USA:Washington-King-County |
| MT627325.1 | 02/28/2020 | China |
| MT632567.1 | 02/28/2020 | USA:Washington-King-County |
| MT163718.1 | 02/29/2020 | USA:WA |
| MT276326.2 | 02/29/2020 | USA:GA |
| MT276327.1 | 02/29/2020 | USA:GA |
| MT304475.1 | 02/29/2020 | South-Korea |
| MT304476.1 | 02/29/2020 | South-Korea |
| MT304484.1 | 02/29/2020 | USA:NH |
| MT304489.1 | 02/29/2020 | USA:TX |
| MT370904.1 | 02/29/2020 | USA:NY |
| MT419841.1 | 02/29/2020 | USA:CA |
| MT479223.1 | 02/29/2020 | Taiwan |
| MT598639.1 | 02/29/2020 | USA:King-County-WA |
| MT598640.1 | 02/29/2020 | USA:King-County-WA |

| | | |
|---|---|---|
| MT598641.1 | 02/29/2020 | USA:King-County-WA |
| MT627216.1 | 02/29/2020 | USA:Washington-King-County |
| MT627233.1 | 02/29/2020 | USA:Washington-Umatilla-County |
| MT627242.1 | 02/29/2020 | USA:Washington-King-County |
| MT627243.1 | 02/29/2020 | USA:Washington-Umatilla-County |
| MT627245.1 | 02/29/2020 | USA:Washington-King-County |
| MT627248.1 | 02/29/2020 | USA:Washington-King-County |
| MT627261.1 | 02/29/2020 | USA:Washington-King-County |
| MT627267.1 | 02/29/2020 | USA:Washington-Snohomish-County |
| MT627269.1 | 02/29/2020 | USA:Washington-King-County |
| MT627279.1 | 02/29/2020 | USA:Washington-Snohomish-County |
| MT627293.1 | 02/29/2020 | USA:Washington-King-County |
| MT627299.1 | 02/29/2020 | USA:Washington |
| MT627304.1 | 02/29/2020 | USA:Washington-King-County |
| MT627307.1 | 02/29/2020 | USA:Washington |
| MT627312.1 | 02/29/2020 | USA:Washington-King-County |
| MT627315.1 | 02/29/2020 | USA:Washington-King-County |
| MT627625.1 | 02/29/2020 | USA:Washington-King-County |
| MT632545.1 | 02/29/2020 | USA:Washington-King-County |
| MT632605.1 | 02/29/2020 | USA:Washington |
| MT632612.1 | 02/29/2020 | USA:Washington-King-County |
| MT163719.1 | 03/01/2020 | USA:WA |
| MT163720.1 | 03/01/2020 | USA:WA |
| MT304482.1 | 03/01/2020 | USA:IL |
| MT304483.1 | 03/01/2020 | USA:IL |
| MT304487.1 | 03/01/2020 | USA:OR |
| MT304488.1 | 03/01/2020 | USA:RI |
| MT304490.1 | 03/01/2020 | USA:TX |
| MT304491.1 | 03/01/2020 | USA:TX |
| MT415320.1 | 03/01/2020 | India |
| MT447174.1 | 03/01/2020 | Thailand |
| MT525950.1 | 03/01/2020 | Italy:Lazio |
| MT531537.2 | 03/01/2020 | Italy |
| MT598635.1 | 03/01/2020 | USA:King-County-WA |
| MT627225.1 | 03/01/2020 | USA:Washington-King-County |
| MT627244.1 | 03/01/2020 | USA:Washington-Clark-County |
| MT627246.1 | 03/01/2020 | USA:Washington-King-County |
| MT627254.1 | 03/01/2020 | USA:Washington-King-County |
| MT627287.1 | 03/01/2020 | USA:Washington-King-County |
| MT627298.1 | 03/01/2020 | USA:Washington-Snohomish-County |
| MT632564.1 | 03/01/2020 | USA:Washington-King-County |
| MT632585.1 | 03/01/2020 | USA:Washington-King-County |
| MT632597.1 | 03/01/2020 | USA:Washington-King-County |
| MT632607.1 | 03/01/2020 | USA:Washington-King-County |
| MT233522.1 | 03/02/2020 | Spain:Valencia |
| MT256917.1 | 03/02/2020 | Spain:Valencia |

| | | |
|---|---|---|
| MT292574.1 | 03/02/2020 | Spain |
| MT304477.1 | 03/02/2020 | USA:AZ |
| MT304478.1 | 03/02/2020 | USA:FL |
| MT304485.1 | 03/02/2020 | USA:NH |
| MT304486.1 | 03/02/2020 | USA:NY |
| MT325565.1 | 03/02/2020 | USA:FL |
| MT419854.1 | 03/02/2020 | USA:CA |
| MT434813.1 | 03/02/2020 | USA:NY |
| MT627217.1 | 03/02/2020 | USA:Washington |
| MT627228.1 | 03/02/2020 | USA:Washington-Snohomish-County |
| MT627231.1 | 03/02/2020 | USA:Washington-Snohomish-County |
| MT627241.1 | 03/02/2020 | USA:Washington-Snohomish-County |
| MT627249.1 | 03/02/2020 | USA:Washington-Grant-County |
| MT627251.1 | 03/02/2020 | USA:Washington |
| MT627262.1 | 03/02/2020 | USA:Washington-Snohomish-County |
| MT627266.1 | 03/02/2020 | USA:Washington-Snohomish-County |
| MT627271.1 | 03/02/2020 | USA:Washington |
| MT627272.1 | 03/02/2020 | USA:Washington-Snohomish-County |
| MT627273.1 | 03/02/2020 | USA:Washington-King-County |
| MT627280.1 | 03/02/2020 | USA:Washington-Grant-County |
| MT627281.1 | 03/02/2020 | USA:Washington-King-County |
| MT627283.1 | 03/02/2020 | USA:Washington-Snohomish-County |
| MT627296.1 | 03/02/2020 | USA:Washington-King-County |
| MT627305.1 | 03/02/2020 | USA:Washington-King-County |
| MT627309.1 | 03/02/2020 | USA:Washington |
| MT627316.1 | 03/02/2020 | USA:Washington |
| MT627317.1 | 03/02/2020 | USA:Washington |
| MT632518.1 | 03/02/2020 | USA:Washington-King-County |
| MT632548.1 | 03/02/2020 | USA:Washington-King-County |
| MT632549.1 | 03/02/2020 | USA:Washington |
| MT632581.1 | 03/02/2020 | USA:Washington-King-County |
| MT304479.1 | 03/03/2020 | USA:GA |
| MT304480.1 | 03/03/2020 | USA:GA |
| MT419847.1 | 03/03/2020 | USA:CA |
| MT450921.1 | 03/03/2020 | Australia:Victoria |
| MT512424.1 | 03/03/2020 | USA |
| MT512444.1 | 03/03/2020 | USA |
| MT582489.1 | 03/03/2020 | Germany:Dusseldorf |
| MT598642.1 | 03/03/2020 | USA:King-County-WA |
| MT627221.1 | 03/03/2020 | USA:Washington-Snohomish-County |
| MT627223.1 | 03/03/2020 | USA:Washington-King-County |
| MT627237.1 | 03/03/2020 | USA:Washington-King-County |
| MT627255.1 | 03/03/2020 | USA:Washington-Snohomish-County |
| MT627265.1 | 03/03/2020 | USA:Washington |
| MT627284.1 | 03/03/2020 | USA:Washington |
| MT627285.1 | 03/03/2020 | USA:Washington-King-County |

| | | |
|---|---|---|
| MT627292.1 | 03/03/2020 | USA:Washington-King-County |
| MT627297.1 | 03/03/2020 | USA:Washington-King-County |
| MT627303.1 | 03/03/2020 | USA:Washington |
| MT627308.1 | 03/03/2020 | USA:Washington-Snohomish-County |
| MT627311.1 | 03/03/2020 | USA:Washington |
| MT627609.1 | 03/03/2020 | USA:Washington-King-County |
| MT632558.1 | 03/03/2020 | USA:Washington-Snohomish-County |
| MT632606.1 | 03/03/2020 | USA:Washington-Snohomish-County |
| MT198651.1 | 03/04/2020 | Spain:Valencia |
| MT233523.1 | 03/04/2020 | Spain:Valencia |
| MT240479.1 | 03/04/2020 | Pakistan:Gilgit |
| MT304481.1 | 03/04/2020 | USA:GA |
| MT325569.1 | 03/04/2020 | USA:GA |
| MT325587.1 | 03/04/2020 | USA:MD |
| MT325596.1 | 03/04/2020 | USA:NJ |
| MT325622.1 | 03/04/2020 | USA:NJ |
| MT325627.1 | 03/04/2020 | USA:NY |
| MT325632.1 | 03/04/2020 | USA:MA |
| MT325633.1 | 03/04/2020 | USA:MA |
| MT344953.1 | 03/04/2020 | USA:MD |
| MT370842.1 | 03/04/2020 | USA:NY |
| MT419842.1 | 03/04/2020 | USA:CA |
| MT419846.1 | 03/04/2020 | USA:CA |
| MT419848.1 | 03/04/2020 | USA:CA |
| MT434786.1 | 03/04/2020 | USA:NY |
| MT472622.1 | 03/04/2020 | USA:MD |
| MT520191.1 | 03/04/2020 | USA:Massachusetts |
| MT520325.1 | 03/04/2020 | USA:Massachusetts |
| MT528239.1 | 03/04/2020 | Italy:Lazio |
| MT539161.1 | 03/04/2020 | USA:Georgia |
| MT627240.1 | 03/04/2020 | USA:Washington-King-County |
| MT627260.1 | 03/04/2020 | USA:Washington-King-County |
| MT627277.1 | 03/04/2020 | USA:Washington-King-County |
| MT627291.1 | 03/04/2020 | USA:Washington-Snohomish-County |
| MT627300.1 | 03/04/2020 | USA:Washington-Snohomish-County |
| MT627306.1 | 03/04/2020 | USA:Washington-King-County |
| MT627618.1 | 03/04/2020 | USA:Washington-King-County |
| MT627638.1 | 03/04/2020 | USA:Washington-King-County |
| MT632571.1 | 03/04/2020 | USA:Washington |
| MT632604.1 | 03/04/2020 | USA:Washington |
| MT632616.1 | 03/04/2020 | USA:Washington-King-County |
| MT198652.2 | 03/05/2020 | Spain:Valencia |
| MT252801.1 | 03/05/2020 | USA:WA |
| MT325566.1 | 03/05/2020 | USA:FL |
| MT325567.1 | 03/05/2020 | USA:FL |
| MT325570.1 | 03/05/2020 | USA:GA |

| | | |
|---|---|---|
| MT325582.1 | 03/05/2020 | USA:MA |
| MT325592.1 | 03/05/2020 | USA:NE |
| MT325599.1 | 03/05/2020 | USA:PA |
| MT325600.1 | 03/05/2020 | USA:PA |
| MT325605.1 | 03/05/2020 | USA:RI |
| MT325609.1 | 03/05/2020 | USA:UT |
| MT325623.1 | 03/05/2020 | USA:SC |
| MT325626.1 | 03/05/2020 | USA:SC |
| MT325629.1 | 03/05/2020 | USA:MA |
| MT325630.1 | 03/05/2020 | USA:MA |
| MT325631.1 | 03/05/2020 | USA:MA |
| MT325634.1 | 03/05/2020 | USA:MA |
| MT334522.1 | 03/05/2020 | USA:UT |
| MT334523.1 | 03/05/2020 | USA:UT |
| MT334524.1 | 03/05/2020 | USA:UT |
| MT344948.1 | 03/05/2020 | USA:HI |
| MT344954.1 | 03/05/2020 | USA:MN |
| MT374102.1 | 03/05/2020 | Taiwan |
| MT419833.1 | 03/05/2020 | USA:CA |
| MT419834.1 | 03/05/2020 | USA:CA |
| MT419835.1 | 03/05/2020 | USA:CA |
| MT419836.1 | 03/05/2020 | USA:CA |
| MT419837.1 | 03/05/2020 | USA:CA |
| MT419838.1 | 03/05/2020 | USA:CA |
| MT419839.1 | 03/05/2020 | USA:CA |
| MT419855.1 | 03/05/2020 | USA:CA |
| MT419856.1 | 03/05/2020 | USA:CA |
| MT419858.1 | 03/05/2020 | USA:CA |
| MT434782.1 | 03/05/2020 | USA:NY |
| MT450922.1 | 03/05/2020 | Australia:Victoria |
| MT459897.1 | 03/05/2020 | Greece:Athens |
| MT472623.1 | 03/05/2020 | USA:VT |
| MT512417.1 | 03/05/2020 | USA |
| MT520282.1 | 03/05/2020 | USA:Massachusetts |
| MT520503.1 | 03/05/2020 | USA:Massachusetts |
| MT605816.1 | 03/05/2020 | USA:Washington-King-County |
| MT627227.1 | 03/05/2020 | USA:Washington-King-County |
| MT627229.1 | 03/05/2020 | USA:Washington |
| MT627232.1 | 03/05/2020 | USA:Washington-King-County |
| MT627234.1 | 03/05/2020 | USA:Washington-Snohomish-County |
| MT627236.1 | 03/05/2020 | USA:Washington-King-County |
| MT627238.1 | 03/05/2020 | USA:Washington-King-County |
| MT627247.1 | 03/05/2020 | USA:Washington-King-County |
| MT627250.1 | 03/05/2020 | USA:Washington-King-County |
| MT627256.1 | 03/05/2020 | USA:Washington-Snohomish-County |
| MT627257.1 | 03/05/2020 | USA:Washington-King-County |

| | | |
|---|---|---|
| MT627259.1 | 03/05/2020 | USA:Washington-King-County |
| MT627268.1 | 03/05/2020 | USA:Washington-Snohomish-County |
| MT627278.1 | 03/05/2020 | USA:Washington-King-County |
| MT627282.1 | 03/05/2020 | USA:Washington-King-County |
| MT627294.1 | 03/05/2020 | USA:Washington-King-County |
| MT627302.1 | 03/05/2020 | USA:Washington-King-County |
| MT627314.1 | 03/05/2020 | USA:Washington-King-County |
| MT627612.1 | 03/05/2020 | USA:Washington-King-County |
| MT627623.1 | 03/05/2020 | USA:Washington-King-County |
| MT627626.1 | 03/05/2020 | USA:Washington-King-County |
| MT627632.1 | 03/05/2020 | USA:Washington-King-County |
| MT627637.1 | 03/05/2020 | USA:Washington-Snohomish-County |
| MT252822.1 | 03/06/2020 | USA:WA |
| MT252824.1 | 03/06/2020 | USA:WA |
| MT256918.1 | 03/06/2020 | Spain:Valencia |
| MT292582.1 | 03/06/2020 | Spain |
| MT325563.1 | 03/06/2020 | USA:DC |
| MT325564.1 | 03/06/2020 | USA:FL |
| MT325568.1 | 03/06/2020 | USA:FL |
| MT325580.1 | 03/06/2020 | USA:KS |
| MT325583.1 | 03/06/2020 | USA:MA |
| MT325584.1 | 03/06/2020 | USA:MA |
| MT325585.1 | 03/06/2020 | USA:MA |
| MT325588.1 | 03/06/2020 | USA:MO |
| MT325591.1 | 03/06/2020 | USA:NC |
| MT325594.1 | 03/06/2020 | USA:NH |
| MT325595.1 | 03/06/2020 | USA:NH |
| MT325604.1 | 03/06/2020 | USA:PA |
| MT325606.1 | 03/06/2020 | USA:SC |
| MT325616.1 | 03/06/2020 | USA:MA |
| MT325628.1 | 03/06/2020 | USA:VA |
| MT325635.1 | 03/06/2020 | USA:MA |
| MT325636.1 | 03/06/2020 | USA:MA |
| MT325637.1 | 03/06/2020 | USA:MA |
| MT325638.1 | 03/06/2020 | USA:MA |
| MT325640.1 | 03/06/2020 | USA:MA |
| MT450923.1 | 03/06/2020 | Australia:Victoria |
| MT472624.1 | 03/06/2020 | USA:FL |
| MT520408.1 | 03/06/2020 | USA:Massachusetts |
| MT627289.1 | 03/06/2020 | USA:Washington-King-County |
| MT627310.1 | 03/06/2020 | USA:Washington-Snohomish-County |
| MT627614.1 | 03/06/2020 | USA:Washington-King-County |
| MT627624.1 | 03/06/2020 | USA:Washington-King-County |
| MT627629.1 | 03/06/2020 | USA:Washington-King-County |
| MT627633.1 | 03/06/2020 | USA:Washington-King-County |
| MT632619.1 | 03/06/2020 | USA:Washington-King-County |

| | | |
|---|---|---|
| MT188340.1 | 03/07/2020 | USA:MN |
| MT252823.1 | 03/07/2020 | USA:WA |
| MT324062.1 | 03/07/2020 | South-Africa:KwaZulu-Natal |
| MT325577.1 | 03/07/2020 | USA:IL |
| MT325578.1 | 03/07/2020 | USA:IL |
| MT325586.1 | 03/07/2020 | USA:MA |
| MT325590.1 | 03/07/2020 | USA:NC |
| MT325593.1 | 03/07/2020 | USA:NE |
| MT325601.1 | 03/07/2020 | USA:PA |
| MT325607.1 | 03/07/2020 | USA:SC |
| MT325610.1 | 03/07/2020 | USA:UT |
| MT325615.1 | 03/07/2020 | USA:IA |
| MT325621.1 | 03/07/2020 | USA:IA |
| MT325624.1 | 03/07/2020 | USA:SC |
| MT325625.1 | 03/07/2020 | USA:SC |
| MT325639.1 | 03/07/2020 | USA:MA |
| MT334525.1 | 03/07/2020 | USA:UT |
| MT334526.1 | 03/07/2020 | USA:UT |
| MT344945.1 | 03/07/2020 | USA:GA |
| MT344949.1 | 03/07/2020 | USA:HI |
| MT344950.1 | 03/07/2020 | USA:IN |
| MT344956.1 | 03/07/2020 | USA:OH |
| MT344957.1 | 03/07/2020 | USA:PA |
| MT344958.1 | 03/07/2020 | USA:PA |
| MT419857.1 | 03/07/2020 | USA:CA |
| MT450932.1 | 03/07/2020 | Australia:Victoria |
| MT450933.1 | 03/07/2020 | Australia:Victoria |
| MT459898.1 | 03/07/2020 | Greece:Athens |
| MT459899.1 | 03/07/2020 | Greece:Athens |
| MT472626.1 | 03/07/2020 | USA:IA |
| MT509463.1 | 03/07/2020 | USA |
| MT512425.1 | 03/07/2020 | USA |
| MT512427.1 | 03/07/2020 | USA |
| MT512430.1 | 03/07/2020 | USA |
| MT520178.1 | 03/07/2020 | USA:Massachusetts |
| MT520192.1 | 03/07/2020 | USA:Massachusetts |
| MT520233.1 | 03/07/2020 | USA:Massachusetts |
| MT520287.1 | 03/07/2020 | USA:Massachusetts |
| MT520303.1 | 03/07/2020 | USA:Massachusetts |
| MT520310.1 | 03/07/2020 | USA:Massachusetts |
| MT520314.1 | 03/07/2020 | USA:Massachusetts |
| MT520316.1 | 03/07/2020 | USA:Massachusetts |
| MT520320.1 | 03/07/2020 | USA:Massachusetts |
| MT520341.1 | 03/07/2020 | USA:Massachusetts |
| MT520375.1 | 03/07/2020 | USA:Massachusetts |
| MT520388.1 | 03/07/2020 | USA:Massachusetts |

| | | |
|---|---|---|
| MT520396.1 | 03/07/2020 | USA:Massachusetts |
| MT520428.1 | 03/07/2020 | USA:Massachusetts |
| MT520441.1 | 03/07/2020 | USA:Massachusetts |
| MT520446.1 | 03/07/2020 | USA:Massachusetts |
| MT520459.1 | 03/07/2020 | USA:Massachusetts |
| MT520470.1 | 03/07/2020 | USA:Massachusetts |
| MT520472.1 | 03/07/2020 | USA:Massachusetts |
| MT520509.1 | 03/07/2020 | USA:Massachusetts |
| MT520517.1 | 03/07/2020 | USA:Massachusetts |
| MT528237.1 | 03/07/2020 | Italy:Lazio |
| MT605817.1 | 03/07/2020 | USA:Washington-King-County |
| MT632576.1 | 03/07/2020 | USA:Washington-King-County |
| MT632593.1 | 03/07/2020 | USA:Washington-Snohomish-County |
| MT646069.1 | 03/07/2020 | USA |
| MT646117.1 | 03/07/2020 | USA |
| LC547518.1 | 03/08/2020 | Japan:Kochi |
| MT198653.1 | 03/08/2020 | Spain:Valencia |
| MT252802.1 | 03/08/2020 | USA:WA |
| MT252804.1 | 03/08/2020 | USA:WA |
| MT252805.1 | 03/08/2020 | USA:WA |
| MT252810.1 | 03/08/2020 | USA:WA |
| MT252811.1 | 03/08/2020 | USA:WA |
| MT276600.1 | 03/08/2020 | Hong-Kong |
| MT292577.1 | 03/08/2020 | Spain |
| MT325572.1 | 03/08/2020 | USA:GA |
| MT325576.1 | 03/08/2020 | USA:IA |
| MT325589.1 | 03/08/2020 | USA:NC |
| MT325598.1 | 03/08/2020 | USA:OH |
| MT325602.1 | 03/08/2020 | USA:PA |
| MT325608.1 | 03/08/2020 | USA:SC |
| MT325611.1 | 03/08/2020 | USA:VA |
| MT325612.1 | 03/08/2020 | USA:VA |
| MT325613.1 | 03/08/2020 | USA:VA |
| MT325614.1 | 03/08/2020 | USA:VA |
| MT325617.1 | 03/08/2020 | USA:IA |
| MT325618.1 | 03/08/2020 | USA:IA |
| MT325619.1 | 03/08/2020 | USA:IA |
| MT325620.1 | 03/08/2020 | USA:IA |
| MT344944.1 | 03/08/2020 | USA:CT |
| MT344951.1 | 03/08/2020 | USA:IN |
| MT344955.1 | 03/08/2020 | USA:OH |
| MT344959.1 | 03/08/2020 | USA:PA |
| MT344963.1 | 03/08/2020 | USA:NC |
| MT419859.1 | 03/08/2020 | USA:CA |
| MT434794.1 | 03/08/2020 | USA:NY |
| MT434816.1 | 03/08/2020 | USA:NY |

| | | |
|---|---|---|
| MT450934.1 | 03/08/2020 | Australia:Victoria |
| MT450935.1 | 03/08/2020 | Australia:Victoria |
| MT459900.1 | 03/08/2020 | Greece:Athens |
| MT472621.1 | 03/08/2020 | USA:PA |
| MT472625.1 | 03/08/2020 | USA:PR |
| MT472627.1 | 03/08/2020 | USA:DC |
| MT509474.1 | 03/08/2020 | USA |
| MT509478.1 | 03/08/2020 | USA |
| MT512426.1 | 03/08/2020 | USA |
| MT512428.1 | 03/08/2020 | USA |
| MT520327.1 | 03/08/2020 | USA:Massachusetts |
| MT520429.1 | 03/08/2020 | USA:Massachusetts |
| MT605815.1 | 03/08/2020 | USA:Washington-King-County |
| MT627616.1 | 03/08/2020 | USA:Washington-King-County |
| MT632502.1 | 03/08/2020 | USA:Washington-Clark-County |
| MT632575.1 | 03/08/2020 | USA:Washington-Clark-County |
| MT632951.1 | 03/08/2020 | USA:Washington |
| LC534419.1 | 03/09/2020 | Japan |
| LC547521.1 | 03/09/2020 | Japan:Ishikawa |
| MT188339.1 | 03/09/2020 | USA:MN |
| MT252787.1 | 03/09/2020 | USA:WA |
| MT252788.1 | 03/09/2020 | USA:WA |
| MT252793.1 | 03/09/2020 | USA:WA |
| MT252799.1 | 03/09/2020 | USA:WA |
| MT252803.1 | 03/09/2020 | USA:WA |
| MT252806.1 | 03/09/2020 | USA:WA |
| MT252807.1 | 03/09/2020 | USA:WA |
| MT252808.1 | 03/09/2020 | USA:WA |
| MT252809.1 | 03/09/2020 | USA:WA |
| MT281530.2 | 03/09/2020 | Iran |
| MT292569.1 | 03/09/2020 | Spain |
| MT292571.1 | 03/09/2020 | Spain |
| MT292573.1 | 03/09/2020 | Spain |
| MT292578.1 | 03/09/2020 | Spain |
| MT292579.1 | 03/09/2020 | Spain |
| MT308692.1 | 03/09/2020 | USA:MI |
| MT308696.1 | 03/09/2020 | USA:MI |
| MT320891.2 | 03/09/2020 | Iran |
| MT325561.1 | 03/09/2020 | USA:AZ |
| MT325562.1 | 03/09/2020 | USA:AZ |
| MT325579.1 | 03/09/2020 | USA:IN |
| MT325581.1 | 03/09/2020 | USA:LA |
| MT325597.1 | 03/09/2020 | USA:NV |
| MT325603.1 | 03/09/2020 | USA:PA |
| MT344952.1 | 03/09/2020 | USA:IN |
| MT344960.1 | 03/09/2020 | USA:RI |

| | | |
|---|---|---|
| MT374103.1 | 03/09/2020 | Taiwan |
| MT419849.1 | 03/09/2020 | USA:CA |
| MT434783.1 | 03/09/2020 | USA:NY |
| MT434784.1 | 03/09/2020 | USA:NY |
| MT434787.1 | 03/09/2020 | USA:NY |
| MT434788.1 | 03/09/2020 | USA:NY |
| MT434789.1 | 03/09/2020 | USA:NY |
| MT434793.1 | 03/09/2020 | USA:NY |
| MT434795.1 | 03/09/2020 | USA:NY |
| MT434796.1 | 03/09/2020 | USA:NY |
| MT434798.1 | 03/09/2020 | USA:NY |
| MT434799.1 | 03/09/2020 | USA:NY |
| MT434800.1 | 03/09/2020 | USA:NY |
| MT434801.1 | 03/09/2020 | USA:NY |
| MT434807.1 | 03/09/2020 | USA:NY |
| MT434814.1 | 03/09/2020 | USA:NY |
| MT450936.1 | 03/09/2020 | Australia:Victoria |
| MT450937.1 | 03/09/2020 | Australia:Victoria |
| MT450938.1 | 03/09/2020 | Australia:Victoria |
| MT450939.1 | 03/09/2020 | Australia:Victoria |
| MT459849.1 | 03/09/2020 | Greece:Athens |
| MT459863.1 | 03/09/2020 | Greece:Athens |
| MT459901.1 | 03/09/2020 | Greece:Athens |
| MT459914.1 | 03/09/2020 | Greece:Athens |
| MT507271.1 | 03/09/2020 | Jamaica |
| MT510999.1 | 03/09/2020 | Netherlands:Leiden |
| MT512415.1 | 03/09/2020 | USA |
| MT512418.1 | 03/09/2020 | USA |
| MT512419.1 | 03/09/2020 | USA |
| MT512429.1 | 03/09/2020 | USA |
| MT512431.1 | 03/09/2020 | USA |
| MT512437.1 | 03/09/2020 | USA |
| MT512438.1 | 03/09/2020 | USA |
| MT512439.1 | 03/09/2020 | USA |
| MT520386.1 | 03/09/2020 | USA:Massachusetts |
| MT632990.1 | 03/09/2020 | USA:Washington |
| LC547519.1 | 03/10/2020 | Japan:Chiba |
| LC547520.1 | 03/10/2020 | Japan:Chiba |
| MT252677.1 | 03/10/2020 | USA:WA |
| MT252678.1 | 03/10/2020 | USA:WA |
| MT252679.1 | 03/10/2020 | USA:WA |
| MT252680.1 | 03/10/2020 | USA:WA |
| MT252681.1 | 03/10/2020 | USA:WA |
| MT252682.1 | 03/10/2020 | USA:WA |
| MT252683.1 | 03/10/2020 | USA:WA |
| MT252684.1 | 03/10/2020 | USA:WA |

| | | |
|---|---|---|
| MT252685.1 | 03/10/2020 | USA:WA |
| MT252686.1 | 03/10/2020 | USA:WA |
| MT252687.1 | 03/10/2020 | USA:WA |
| MT252688.1 | 03/10/2020 | USA:WA |
| MT252689.1 | 03/10/2020 | USA:WA |
| MT252690.1 | 03/10/2020 | USA:WA |
| MT252694.1 | 03/10/2020 | USA:WA |
| MT252695.1 | 03/10/2020 | USA:WA |
| MT252765.1 | 03/10/2020 | USA:WA |
| MT252792.1 | 03/10/2020 | USA:WA |
| MT252794.1 | 03/10/2020 | USA:WA |
| MT252795.1 | 03/10/2020 | USA:WA |
| MT252797.1 | 03/10/2020 | USA:WA |
| MT252798.1 | 03/10/2020 | USA:WA |
| MT252800.1 | 03/10/2020 | USA:WA |
| MT263074.1 | 03/10/2020 | Peru |
| MT281577.1 | 03/10/2020 | China:Anhui-Fuyang |
| MT292570.1 | 03/10/2020 | Spain |
| MT292572.1 | 03/10/2020 | Spain |
| MT292575.1 | 03/10/2020 | Spain |
| MT292576.1 | 03/10/2020 | Spain |
| MT295464.1 | 03/10/2020 | USA |
| MT308693.1 | 03/10/2020 | USA:MI |
| MT308695.1 | 03/10/2020 | USA:MI |
| MT325575.1 | 03/10/2020 | USA:GA |
| MT334529.1 | 03/10/2020 | USA:UT |
| MT371048.1 | 03/10/2020 | Sri-Lanka |
| MT374104.1 | 03/10/2020 | Taiwan |
| MT412135.1 | 03/10/2020 | USA:Michigan |
| MT412239.1 | 03/10/2020 | USA |
| MT434805.1 | 03/10/2020 | USA:NY |
| MT434806.1 | 03/10/2020 | USA:NY |
| MT439230.1 | 03/10/2020 | USA:MI |
| MT439231.1 | 03/10/2020 | USA:MI |
| MT439232.1 | 03/10/2020 | USA:MI |
| MT439233.1 | 03/10/2020 | USA:MI |
| MT450940.1 | 03/10/2020 | Australia:Victoria |
| MT450941.1 | 03/10/2020 | Australia:Victoria |
| MT450942.1 | 03/10/2020 | Australia:Victoria |
| MT450943.1 | 03/10/2020 | Australia:Victoria |
| MT450944.1 | 03/10/2020 | Australia:Victoria |
| MT450945.1 | 03/10/2020 | Australia:Victoria |
| MT450946.1 | 03/10/2020 | Australia:Victoria |
| MT450947.1 | 03/10/2020 | Australia:Victoria |
| MT450948.1 | 03/10/2020 | Australia:Victoria |
| MT450949.1 | 03/10/2020 | Australia:Victoria |

| | | |
|---|---|---|
| MT450950.1 | 03/10/2020 | Australia:Victoria |
| MT450951.1 | 03/10/2020 | Australia:Victoria |
| MT459850.1 | 03/10/2020 | Greece:Athens |
| MT459851.1 | 03/10/2020 | Greece:Athens |
| MT459902.1 | 03/10/2020 | Greece:Athens |
| MT459907.1 | 03/10/2020 | Greece:Athens |
| MT459908.1 | 03/10/2020 | Greece:Athens |
| MT459909.1 | 03/10/2020 | Greece:Athens |
| MT459915.1 | 03/10/2020 | Greece:Athens |
| MT459916.1 | 03/10/2020 | Greece:Athens |
| MT459917.1 | 03/10/2020 | Greece:Athens |
| MT459918.1 | 03/10/2020 | Greece:Athens |
| MT459919.1 | 03/10/2020 | Greece:Athens |
| MT506199.1 | 03/10/2020 | USA:Michigan |
| MT512416.1 | 03/10/2020 | USA |
| MT512433.1 | 03/10/2020 | USA |
| MT512434.1 | 03/10/2020 | USA |
| MT512435.1 | 03/10/2020 | USA |
| MT512436.1 | 03/10/2020 | USA |
| MT512440.1 | 03/10/2020 | USA |
| MT512441.1 | 03/10/2020 | USA |
| MT512442.1 | 03/10/2020 | USA |
| MT512445.1 | 03/10/2020 | USA |
| MT539163.1 | 03/10/2020 | USA:Georgia |
| MT582451.1 | 03/10/2020 | Germany:Dusseldorf |
| MT582488.1 | 03/10/2020 | Germany:Dusseldorf |
| MT627617.1 | 03/10/2020 | USA:Washington-King-County |
| MT627621.1 | 03/10/2020 | USA:Washington-King-County |
| MT632609.1 | 03/10/2020 | USA:Washington-King-County |
| MT632837.1 | 03/10/2020 | USA:Washington |
| MT646118.1 | 03/10/2020 | USA |
| LC547523.1 | 03/11/2020 | Japan:Saitama |
| MT192765.1 | 03/11/2020 | USA:CA-San-Diego-County |
| MT252691.1 | 03/11/2020 | USA:WA |
| MT252692.1 | 03/11/2020 | USA:WA |
| MT252693.1 | 03/11/2020 | USA:WA |
| MT252748.1 | 03/11/2020 | USA:WA |
| MT252749.1 | 03/11/2020 | USA:WA |
| MT252756.1 | 03/11/2020 | USA:WA |
| MT252757.1 | 03/11/2020 | USA:WA |
| MT252758.1 | 03/11/2020 | USA:WA |
| MT252766.1 | 03/11/2020 | USA:WA |
| MT252767.1 | 03/11/2020 | USA:WA |
| MT252768.1 | 03/11/2020 | USA:WA |
| MT252769.1 | 03/11/2020 | USA:WA |
| MT252770.1 | 03/11/2020 | USA:WA |

| | | |
|---|---|---|
| MT252771.1 | 03/11/2020 | USA:WA |
| MT252772.1 | 03/11/2020 | USA:WA |
| MT252773.1 | 03/11/2020 | USA:WA |
| MT252774.1 | 03/11/2020 | USA:WA |
| MT252775.1 | 03/11/2020 | USA:WA |
| MT252778.1 | 03/11/2020 | USA:WA |
| MT252779.1 | 03/11/2020 | USA:WA |
| MT252780.1 | 03/11/2020 | USA:WA |
| MT252781.1 | 03/11/2020 | USA:WA |
| MT252782.1 | 03/11/2020 | USA:WA |
| MT252783.1 | 03/11/2020 | USA:WA |
| MT252784.1 | 03/11/2020 | USA:WA |
| MT252785.1 | 03/11/2020 | USA:WA |
| MT256924.2 | 03/11/2020 | Colombia:Antioquia |
| MT308694.1 | 03/11/2020 | USA:MI |
| MT308697.1 | 03/11/2020 | USA:MI |
| MT334527.1 | 03/11/2020 | USA:UT |
| MT334530.1 | 03/11/2020 | USA:UT |
| MT334541.1 | 03/11/2020 | USA:UT |
| MT334542.1 | 03/11/2020 | USA:UT |
| MT370892.1 | 03/11/2020 | USA:NY |
| MT412242.1 | 03/11/2020 | USA |
| MT412245.1 | 03/11/2020 | USA |
| MT412246.1 | 03/11/2020 | USA:WA |
| MT412252.1 | 03/11/2020 | USA:WA |
| MT412253.1 | 03/11/2020 | USA:WA |
| MT415321.1 | 03/11/2020 | India |
| MT439234.1 | 03/11/2020 | USA:MI |
| MT439235.1 | 03/11/2020 | USA:MI |
| MT444148.1 | 03/11/2020 | USA:CA |
| MT447167.1 | 03/11/2020 | Thailand |
| MT450952.1 | 03/11/2020 | Australia:Victoria |
| MT450953.1 | 03/11/2020 | Australia:Victoria |
| MT450954.1 | 03/11/2020 | Australia:Victoria |
| MT450955.1 | 03/11/2020 | Australia:Victoria |
| MT450956.1 | 03/11/2020 | Australia:Victoria |
| MT450957.1 | 03/11/2020 | Australia:Victoria |
| MT450958.1 | 03/11/2020 | Australia:Victoria |
| MT450959.1 | 03/11/2020 | Australia:Victoria |
| MT450960.1 | 03/11/2020 | Australia:Victoria |
| MT450961.1 | 03/11/2020 | Australia:Victoria |
| MT459852.1 | 03/11/2020 | Greece:Athens |
| MT459853.1 | 03/11/2020 | Greece:Athens |
| MT459854.1 | 03/11/2020 | Greece:Athens |
| MT459864.1 | 03/11/2020 | Greece:Athens |
| MT459910.1 | 03/11/2020 | Greece:Athens |

| | | |
|---|---|---|
| MT459912.1 | 03/11/2020 | Greece:Athens |
| MT459913.1 | 03/11/2020 | Greece:Athens |
| MT459920.1 | 03/11/2020 | Greece:Athens |
| MT459921.1 | 03/11/2020 | Greece:Athens |
| MT459922.1 | 03/11/2020 | Greece:Athens |
| MT507272.1 | 03/11/2020 | Jamaica |
| MT509472.1 | 03/11/2020 | USA |
| MT509475.1 | 03/11/2020 | USA |
| MT509669.1 | 03/11/2020 | USA:Michigan |
| MT512443.1 | 03/11/2020 | USA |
| MT520473.1 | 03/11/2020 | USA:Massachusetts |
| MT520515.1 | 03/11/2020 | USA:Massachusetts |
| MT520542.1 | 03/11/2020 | USA:Massachusetts |
| MT520547.1 | 03/11/2020 | USA:Massachusetts |
| MT582450.1 | 03/11/2020 | Germany:Dusseldorf |
| MT582483.1 | 03/11/2020 | Germany:Dusseldorf |
| MT582484.1 | 03/11/2020 | Germany:Dusseldorf |
| MT582485.1 | 03/11/2020 | Germany:Dusseldorf |
| MT582486.1 | 03/11/2020 | Germany:Dusseldorf |
| MT582487.1 | 03/11/2020 | Germany:Dusseldorf |
| MT612274.1 | 03/11/2020 | Australia:Victoria |
| MT614520.1 | 03/11/2020 | USA:UT |
| MT632570.1 | 03/11/2020 | USA:Washington-Skagit-County |
| MT646071.1 | 03/11/2020 | USA |
| MT646078.1 | 03/11/2020 | USA |
| LC547524.1 | 03/12/2020 | Japan:Saitama |
| LC547525.1 | 03/12/2020 | Japan:Saitama |
| LC547531.1 | 03/12/2020 | Japan:Chiba |
| MT246454.1 | 03/12/2020 | USA:WA |
| MT246455.1 | 03/12/2020 | USA:WA |
| MT246458.1 | 03/12/2020 | USA:WA |
| MT246459.1 | 03/12/2020 | USA:WA |
| MT246460.1 | 03/12/2020 | USA:WA |
| MT246472.1 | 03/12/2020 | USA:WA |
| MT251974.1 | 03/12/2020 | USA:CT |
| MT252701.1 | 03/12/2020 | USA:WA |
| MT252726.1 | 03/12/2020 | USA:WA |
| MT252741.1 | 03/12/2020 | USA:WA |
| MT252742.1 | 03/12/2020 | USA:WA |
| MT252745.1 | 03/12/2020 | USA:WA |
| MT252746.1 | 03/12/2020 | USA:WA |
| MT252747.1 | 03/12/2020 | USA:WA |
| MT252761.1 | 03/12/2020 | USA:WA |
| MT252763.1 | 03/12/2020 | USA:WA |
| MT252764.1 | 03/12/2020 | USA:WA |
| MT263460.1 | 03/12/2020 | USA:IL |

| | | |
|---|---|---|
| MT308698.1 | 03/12/2020 | USA:MI |
| MT308699.1 | 03/12/2020 | USA:MI |
| MT308700.1 | 03/12/2020 | USA:MI |
| MT334528.1 | 03/12/2020 | USA:UT |
| MT334531.1 | 03/12/2020 | USA:UT |
| MT334532.1 | 03/12/2020 | USA:UT |
| MT334549.1 | 03/12/2020 | USA:UT |
| MT412136.1 | 03/12/2020 | USA:Michigan |
| MT412139.1 | 03/12/2020 | USA:Michigan |
| MT412143.1 | 03/12/2020 | USA:Michigan |
| MT412236.1 | 03/12/2020 | USA:WA |
| MT412237.1 | 03/12/2020 | USA:WA |
| MT412238.1 | 03/12/2020 | USA:WA |
| MT412240.1 | 03/12/2020 | USA:WA |
| MT412241.1 | 03/12/2020 | USA:WA |
| MT412244.1 | 03/12/2020 | USA:WA |
| MT412247.1 | 03/12/2020 | USA:WA |
| MT412248.1 | 03/12/2020 | USA:WA |
| MT412249.1 | 03/12/2020 | USA:WA |
| MT412250.1 | 03/12/2020 | USA:WA |
| MT412251.1 | 03/12/2020 | USA:WA |
| MT439236.1 | 03/12/2020 | USA:MI |
| MT439237.1 | 03/12/2020 | USA:MI |
| MT439239.1 | 03/12/2020 | USA:MI |
| MT444560.1 | 03/12/2020 | USA |
| MT444561.1 | 03/12/2020 | USA |
| MT444562.1 | 03/12/2020 | USA |
| MT444563.1 | 03/12/2020 | USA |
| MT447168.1 | 03/12/2020 | Thailand |
| MT447169.1 | 03/12/2020 | Thailand |
| MT450962.1 | 03/12/2020 | Australia:Victoria |
| MT450963.1 | 03/12/2020 | Australia:Victoria |
| MT450964.1 | 03/12/2020 | Australia:Victoria |
| MT450965.1 | 03/12/2020 | Australia:Victoria |
| MT450966.1 | 03/12/2020 | Australia:Victoria |
| MT450967.1 | 03/12/2020 | Australia:Victoria |
| MT450968.1 | 03/12/2020 | Australia:Victoria |
| MT450969.1 | 03/12/2020 | Australia:Victoria |
| MT450970.1 | 03/12/2020 | Australia:Victoria |
| MT450971.1 | 03/12/2020 | Australia:Victoria |
| MT450972.1 | 03/12/2020 | Australia:Victoria |
| MT450973.1 | 03/12/2020 | Australia:Victoria |
| MT452574.1 | 03/12/2020 | USA:NY |
| MT459855.1 | 03/12/2020 | Greece:Athens |
| MT459856.1 | 03/12/2020 | Greece:Athens |
| MT459857.1 | 03/12/2020 | Greece:Athens |

| | | |
|---|---|---|
| MT459865.1 | 03/12/2020 | Greece:Athens |
| MT459870.1 | 03/12/2020 | Greece:Athens |
| MT459871.1 | 03/12/2020 | Greece:Athens |
| MT459872.1 | 03/12/2020 | Greece:Athens |
| MT459889.1 | 03/12/2020 | Greece:Athens |
| MT506527.1 | 03/12/2020 | USA:Michigan |
| MT506528.1 | 03/12/2020 | USA:Michigan |
| MT506531.1 | 03/12/2020 | USA:Michigan |
| MT506637.1 | 03/12/2020 | USA:Michigan |
| MT614510.1 | 03/12/2020 | USA:UT |
| MT614512.1 | 03/12/2020 | USA:UT |
| MT614515.1 | 03/12/2020 | USA:UT |
| MT614517.1 | 03/12/2020 | USA:UT |
| MT614518.1 | 03/12/2020 | USA:UT |
| MT627419.1 | 03/12/2020 | USA:CO |
| MT627420.1 | 03/12/2020 | USA:HI |
| MT627421.1 | 03/12/2020 | USA:HI |
| MT627422.1 | 03/12/2020 | USA:IN |
| MT627423.1 | 03/12/2020 | USA:NY |
| MT627424.1 | 03/12/2020 | USA:NY |
| MT627425.1 | 03/12/2020 | USA:NY |
| MT627426.1 | 03/12/2020 | USA:NY |
| MT627427.1 | 03/12/2020 | USA:NY |
| MT627428.1 | 03/12/2020 | USA:NY |
| MT627429.1 | 03/12/2020 | USA:NY |
| MT627430.1 | 03/12/2020 | USA:NY |
| MT627431.1 | 03/12/2020 | USA:NY |
| MT627432.1 | 03/12/2020 | USA:NY |
| MT627433.1 | 03/12/2020 | USA:NY |
| MT627434.1 | 03/12/2020 | USA:NY |
| MT627613.1 | 03/12/2020 | USA:Washington-King-County |
| MT627636.1 | 03/12/2020 | USA:Washington-King-County |
| MT632562.1 | 03/12/2020 | USA:Washington-Snohomish-County |
| MT632595.1 | 03/12/2020 | USA:Washington-King-County |
| MT632598.1 | 03/12/2020 | USA:Washington-King-County |
| LC547532.1 | 03/13/2020 | Japan:Chiba |
| LC547533.1 | 03/13/2020 | Japan:Chiba |
| MT246449.1 | 03/13/2020 | USA:WA |
| MT246450.1 | 03/13/2020 | USA:WA |
| MT246451.1 | 03/13/2020 | USA:WA |
| MT246452.1 | 03/13/2020 | USA:WA |
| MT246453.1 | 03/13/2020 | USA:WA |
| MT246456.1 | 03/13/2020 | USA:WA |
| MT246461.1 | 03/13/2020 | USA:WA |
| MT246462.1 | 03/13/2020 | USA:WA |
| MT246463.1 | 03/13/2020 | USA:WA |

| | | |
|---|---|---|
| MT246464.1 | 03/13/2020 | USA:WA |
| MT246465.1 | 03/13/2020 | USA:WA |
| MT246466.1 | 03/13/2020 | USA:WA |
| MT246470.1 | 03/13/2020 | USA:WA |
| MT246475.1 | 03/13/2020 | USA:WA |
| MT246476.1 | 03/13/2020 | USA:WA |
| MT246479.1 | 03/13/2020 | USA:WA |
| MT246480.1 | 03/13/2020 | USA:WA |
| MT246481.1 | 03/13/2020 | USA:WA |
| MT252696.1 | 03/13/2020 | USA:WA |
| MT252697.1 | 03/13/2020 | USA:WA |
| MT252698.1 | 03/13/2020 | USA:WA |
| MT252699.1 | 03/13/2020 | USA:WA |
| MT252700.1 | 03/13/2020 | USA:WA |
| MT252702.1 | 03/13/2020 | USA:WA |
| MT252703.1 | 03/13/2020 | USA:WA |
| MT252704.1 | 03/13/2020 | USA:WA |
| MT252705.1 | 03/13/2020 | USA:WA |
| MT252706.1 | 03/13/2020 | USA:WA |
| MT252707.1 | 03/13/2020 | USA:WA |
| MT252708.1 | 03/13/2020 | USA:WA |
| MT252721.1 | 03/13/2020 | USA:WA |
| MT252723.1 | 03/13/2020 | USA:WA |
| MT252725.1 | 03/13/2020 | USA:WA |
| MT252728.1 | 03/13/2020 | USA:WA |
| MT252729.1 | 03/13/2020 | USA:WA |
| MT252730.1 | 03/13/2020 | USA:WA |
| MT252733.1 | 03/13/2020 | USA:WA |
| MT259235.1 | 03/13/2020 | USA:WA |
| MT259243.1 | 03/13/2020 | USA:WA |
| MT259246.1 | 03/13/2020 | USA:WA |
| MT259249.1 | 03/13/2020 | USA:WA |
| MT259256.1 | 03/13/2020 | USA:WA |
| MT259258.1 | 03/13/2020 | USA:MN |
| MT259263.1 | 03/13/2020 | USA:WA |
| MT259264.1 | 03/13/2020 | USA:WA |
| MT259266.1 | 03/13/2020 | USA:WA |
| MT259268.1 | 03/13/2020 | USA:WA |
| MT262896.1 | 03/13/2020 | USA:WA |
| MT262897.1 | 03/13/2020 | USA:WA |
| MT262898.1 | 03/13/2020 | USA:WA |
| MT262899.1 | 03/13/2020 | USA:WA |
| MT262900.1 | 03/13/2020 | USA:WA |
| MT262901.1 | 03/13/2020 | USA:WA |
| MT262902.1 | 03/13/2020 | USA:WA |
| MT262903.1 | 03/13/2020 | USA:WA |

| | | |
|---|---|---|
| MT262904.1 | 03/13/2020 | USA:WA |
| MT262905.1 | 03/13/2020 | USA:WA |
| MT262906.1 | 03/13/2020 | USA:WA |
| MT262907.1 | 03/13/2020 | USA:WA |
| MT262908.1 | 03/13/2020 | USA:WA |
| MT262909.1 | 03/13/2020 | USA:WA |
| MT262910.1 | 03/13/2020 | USA:WA |
| MT262911.1 | 03/13/2020 | USA:WA |
| MT262912.1 | 03/13/2020 | USA:WA |
| MT262913.1 | 03/13/2020 | USA:WA |
| MT262914.1 | 03/13/2020 | USA:WA |
| MT262915.1 | 03/13/2020 | USA:WA |
| MT262916.1 | 03/13/2020 | USA:WA |
| MT263396.1 | 03/13/2020 | USA:WA |
| MT263406.1 | 03/13/2020 | USA:MN |
| MT263415.1 | 03/13/2020 | USA:MN |
| MT263428.1 | 03/13/2020 | USA:IL |
| MT263429.1 | 03/13/2020 | USA:IL |
| MT263433.1 | 03/13/2020 | USA:IL |
| MT326182.1 | 03/13/2020 | USA |
| MT334533.1 | 03/13/2020 | USA:UT |
| MT334534.1 | 03/13/2020 | USA:UT |
| MT334535.1 | 03/13/2020 | USA:UT |
| MT334536.1 | 03/13/2020 | USA:UT |
| MT334537.1 | 03/13/2020 | USA:UT |
| MT334538.1 | 03/13/2020 | USA:UT |
| MT370881.1 | 03/13/2020 | USA:NY |
| MT370889.1 | 03/13/2020 | USA:NY |
| MT370910.1 | 03/13/2020 | USA:NY |
| MT370915.1 | 03/13/2020 | USA:NY |
| MT374105.1 | 03/13/2020 | Taiwan |
| MT374106.1 | 03/13/2020 | Taiwan |
| MT412137.1 | 03/13/2020 | USA:Michigan |
| MT412138.1 | 03/13/2020 | USA:Michigan |
| MT412141.1 | 03/13/2020 | USA:Michigan |
| MT412142.1 | 03/13/2020 | USA:Michigan |
| MT412145.1 | 03/13/2020 | USA:Michigan |
| MT412233.1 | 03/13/2020 | USA |
| MT412234.1 | 03/13/2020 | USA:WA |
| MT412235.1 | 03/13/2020 | USA:WA |
| MT419851.1 | 03/13/2020 | USA:CA |
| MT434802.1 | 03/13/2020 | USA:NY |
| MT439238.1 | 03/13/2020 | USA:MI |
| MT439240.1 | 03/13/2020 | USA:MI |
| MT439241.1 | 03/13/2020 | USA:MI |
| MT439243.1 | 03/13/2020 | USA:MI |

| | | |
|---|---|---|
| MT439244.1 | 03/13/2020 | USA:MI |
| MT439245.1 | 03/13/2020 | USA:MI |
| MT450974.1 | 03/13/2020 | Australia:Victoria |
| MT450975.1 | 03/13/2020 | Australia:Victoria |
| MT450976.1 | 03/13/2020 | Australia:Victoria |
| MT450977.1 | 03/13/2020 | Australia:Victoria |
| MT450978.1 | 03/13/2020 | Australia:Victoria |
| MT450979.1 | 03/13/2020 | Australia:Victoria |
| MT450980.1 | 03/13/2020 | Australia:Victoria |
| MT450981.1 | 03/13/2020 | Australia:Victoria |
| MT450982.1 | 03/13/2020 | Australia:Victoria |
| MT450983.1 | 03/13/2020 | Australia:Victoria |
| MT450984.1 | 03/13/2020 | Australia:Victoria |
| MT452575.1 | 03/13/2020 | USA:WA |
| MT452576.1 | 03/13/2020 | USA:CA |
| MT459866.1 | 03/13/2020 | Greece:Athens |
| MT459873.1 | 03/13/2020 | Greece:Athens |
| MT459874.1 | 03/13/2020 | Greece:Athens |
| MT459875.1 | 03/13/2020 | Greece:Athens |
| MT466071.1 | 03/13/2020 | Uruguay |
| MT506200.1 | 03/13/2020 | USA:Michigan |
| MT506202.1 | 03/13/2020 | USA:Michigan |
| MT506208.1 | 03/13/2020 | USA:Michigan |
| MT506210.1 | 03/13/2020 | USA:Ohio |
| MT506526.1 | 03/13/2020 | USA:Michigan |
| MT506529.1 | 03/13/2020 | USA:Michigan |
| MT506636.1 | 03/13/2020 | USA:Michigan |
| MT506646.1 | 03/13/2020 | USA:Michigan |
| MT509454.1 | 03/13/2020 | USA |
| MT512446.1 | 03/13/2020 | USA |
| MT520328.1 | 03/13/2020 | USA:Massachusetts |
| MT539162.1 | 03/13/2020 | USA:Georgia |
| MT582449.1 | 03/13/2020 | Germany:Dusseldorf |
| MT582482.1 | 03/13/2020 | Germany:Dusseldorf |
| MT614464.1 | 03/13/2020 | USA:CA |
| MT614471.1 | 03/13/2020 | USA:CA |
| MT614476.1 | 03/13/2020 | USA:GA |
| MT614488.1 | 03/13/2020 | USA:MI |
| MT614504.1 | 03/13/2020 | USA:NY |
| MT614506.1 | 03/13/2020 | USA:PA |
| MT614511.1 | 03/13/2020 | USA:UT |
| MT614513.1 | 03/13/2020 | USA:UT |
| MT614514.1 | 03/13/2020 | USA:UT |
| MT614516.1 | 03/13/2020 | USA:UT |
| MT614519.1 | 03/13/2020 | USA:UT |
| MT614521.1 | 03/13/2020 | USA:WA |

| | | |
|---|---|---|
| MT614522.1 | 03/13/2020 | USA:WA |
| MT614527.1 | 03/13/2020 | USA:FL |
| MT614528.1 | 03/13/2020 | USA:GA |
| MT614529.1 | 03/13/2020 | USA:GA |
| MT614530.1 | 03/13/2020 | USA:GA |
| MT614532.1 | 03/13/2020 | USA:MA |
| MT614536.1 | 03/13/2020 | USA:MD |
| MT614537.1 | 03/13/2020 | USA:NJ |
| MT614539.1 | 03/13/2020 | USA:NJ |
| MT614541.1 | 03/13/2020 | USA:NJ |
| MT614555.1 | 03/13/2020 | USA:TN |
| MT614556.1 | 03/13/2020 | USA:UT |
| MT614557.1 | 03/13/2020 | USA:UT |
| MT614558.1 | 03/13/2020 | USA:UT |
| MT614559.1 | 03/13/2020 | USA:UT |
| MT614560.1 | 03/13/2020 | USA:UT |
| MT614561.1 | 03/13/2020 | USA:GA |
| MT614563.1 | 03/13/2020 | USA:UT |
| MT627407.1 | 03/13/2020 | USA:CA |
| MT627408.1 | 03/13/2020 | USA:CA |
| MT627409.1 | 03/13/2020 | USA:CA |
| MT627410.1 | 03/13/2020 | USA:CA |
| MT627411.1 | 03/13/2020 | USA:CA |
| MT627412.1 | 03/13/2020 | USA:CA |
| MT627413.1 | 03/13/2020 | USA:CA |
| MT627414.1 | 03/13/2020 | USA:CA |
| MT627415.1 | 03/13/2020 | USA:CA |
| MT627416.1 | 03/13/2020 | USA:CA |
| MT627417.1 | 03/13/2020 | USA:CA |
| MT627418.1 | 03/13/2020 | USA:CO |
| MT627435.1 | 03/13/2020 | USA:NY |
| MT627436.1 | 03/13/2020 | USA:OK |
| MT627437.1 | 03/13/2020 | USA:WA |
| MT627615.1 | 03/13/2020 | USA:Washington-King-County |
| MT627619.1 | 03/13/2020 | USA:Washington-King-County |
| MT628124.1 | 03/13/2020 | USA:CA |
| MT628230.1 | 03/13/2020 | USA:CA |
| MT632541.1 | 03/13/2020 | USA:Washington-King-County |
| MT632552.1 | 03/13/2020 | USA:Washington-King-County |
| MT632555.1 | 03/13/2020 | USA:Washington-King-County |
| MT632569.1 | 03/13/2020 | USA:Washington-King-County |
| MT632580.1 | 03/13/2020 | USA:Washington-King-County |
| MT632582.1 | 03/13/2020 | USA:Washington-King-County |
| MT632587.1 | 03/13/2020 | USA:Washington-King-County |
| MT632592.1 | 03/13/2020 | USA:Washington-King-County |
| MT632600.1 | 03/13/2020 | USA:Washington-King-County |

| | | |
|---|---|---|
| MT632602.1 | 03/13/2020 | USA:Washington-King-County |
| MT632603.1 | 03/13/2020 | USA:Washington-King-County |
| MT632608.1 | 03/13/2020 | USA:Washington-King-County |
| MT632610.1 | 03/13/2020 | USA:Washington-King-County |
| MT632613.1 | 03/13/2020 | USA:Washington-King-County |
| MT632617.1 | 03/13/2020 | USA:Washington-King-County |
| MT632618.1 | 03/13/2020 | USA:Washington-King-County |
| MT632623.1 | 03/13/2020 | USA:Washington-King-County |
| MT635233.1 | 03/13/2020 | USA:Connecticut |
| MT646075.1 | 03/13/2020 | USA |
| MT646119.1 | 03/13/2020 | USA |
| MT246467.1 | 03/14/2020 | USA:WA |
| MT246468.1 | 03/14/2020 | USA:WA |
| MT246469.1 | 03/14/2020 | USA:WA |
| MT246471.1 | 03/14/2020 | USA:WA |
| MT246474.1 | 03/14/2020 | USA:WA |
| MT246477.1 | 03/14/2020 | USA:WA |
| MT246478.1 | 03/14/2020 | USA:WA |
| MT246486.1 | 03/14/2020 | USA:WA |
| MT246488.1 | 03/14/2020 | USA:WA |
| MT246489.1 | 03/14/2020 | USA:WA |
| MT251972.1 | 03/14/2020 | USA:WA |
| MT251973.1 | 03/14/2020 | USA:WA |
| MT251975.1 | 03/14/2020 | USA:WA |
| MT251976.1 | 03/14/2020 | USA:WA |
| MT251977.1 | 03/14/2020 | USA:WA |
| MT251978.1 | 03/14/2020 | USA:WA |
| MT251979.1 | 03/14/2020 | USA:WA |
| MT252709.1 | 03/14/2020 | USA:WA |
| MT252710.1 | 03/14/2020 | USA:WA |
| MT252711.1 | 03/14/2020 | USA:WA |
| MT252712.1 | 03/14/2020 | USA:WA |
| MT252715.1 | 03/14/2020 | USA:WA |
| MT252716.1 | 03/14/2020 | USA:WA |
| MT252717.1 | 03/14/2020 | USA:WA |
| MT252734.1 | 03/14/2020 | USA:WA |
| MT252735.1 | 03/14/2020 | USA:WA |
| MT252736.1 | 03/14/2020 | USA:WA |
| MT252737.1 | 03/14/2020 | USA:WA |
| MT252740.1 | 03/14/2020 | USA:WA |
| MT259248.1 | 03/14/2020 | USA:CT |
| MT259250.1 | 03/14/2020 | USA:WA |
| MT259251.1 | 03/14/2020 | USA:WA |
| MT259252.1 | 03/14/2020 | USA:WA |
| MT259262.1 | 03/14/2020 | USA:WA |
| MT259267.1 | 03/14/2020 | USA:WA |

| | | |
|---|---|---|
| MT259269.1 | 03/14/2020 | USA:WA |
| MT259274.1 | 03/14/2020 | USA:WA |
| MT259275.1 | 03/14/2020 | USA:WA |
| MT263386.1 | 03/14/2020 | USA:CT |
| MT263394.1 | 03/14/2020 | USA:WA |
| MT263395.1 | 03/14/2020 | USA:WA |
| MT263407.1 | 03/14/2020 | USA:CT |
| MT263411.1 | 03/14/2020 | USA:CT |
| MT263437.1 | 03/14/2020 | USA:WA |
| MT263447.1 | 03/14/2020 | USA:WA |
| MT326181.1 | 03/14/2020 | USA |
| MT334539.1 | 03/14/2020 | USA:UT |
| MT334540.1 | 03/14/2020 | USA:UT |
| MT334550.1 | 03/14/2020 | USA:UT |
| MT334551.1 | 03/14/2020 | USA:UT |
| MT370839.1 | 03/14/2020 | USA:NY |
| MT370854.1 | 03/14/2020 | USA:NY |
| MT370857.1 | 03/14/2020 | USA:NY |
| MT370919.1 | 03/14/2020 | USA:NY |
| MT374107.1 | 03/14/2020 | Taiwan |
| MT374108.1 | 03/14/2020 | Taiwan |
| MT394864.1 | 03/14/2020 | Germany:Dusseldorf |
| MT412140.1 | 03/14/2020 | USA:Michigan |
| MT412144.1 | 03/14/2020 | USA:Michigan |
| MT412150.1 | 03/14/2020 | USA:Michigan |
| MT419853.1 | 03/14/2020 | USA:CA |
| MT439242.1 | 03/14/2020 | USA:MI |
| MT439246.1 | 03/14/2020 | USA:MI |
| MT444531.1 | 03/14/2020 | USA |
| MT444532.1 | 03/14/2020 | USA |
| MT444556.1 | 03/14/2020 | USA |
| MT444559.1 | 03/14/2020 | USA |
| MT450985.1 | 03/14/2020 | Australia:Victoria |
| MT450986.1 | 03/14/2020 | Australia:Victoria |
| MT450987.1 | 03/14/2020 | Australia:Victoria |
| MT450988.1 | 03/14/2020 | Australia:Victoria |
| MT450989.1 | 03/14/2020 | Australia:Victoria |
| MT450990.1 | 03/14/2020 | Australia:Victoria |
| MT450991.1 | 03/14/2020 | Australia:Victoria |
| MT450992.1 | 03/14/2020 | Australia:Victoria |
| MT450993.1 | 03/14/2020 | Australia:Victoria |
| MT450994.1 | 03/14/2020 | Australia:Victoria |
| MT459836.1 | 03/14/2020 | Greece:Athens |
| MT459858.1 | 03/14/2020 | Greece:Athens |
| MT459876.1 | 03/14/2020 | Greece:Athens |
| MT459877.1 | 03/14/2020 | Greece:Athens |

| | | |
|---|---|---|
| MT459878.1 | 03/14/2020 | Greece:Athens |
| MT459879.1 | 03/14/2020 | Greece:Athens |
| MT459880.1 | 03/14/2020 | Greece:Athens |
| MT459890.1 | 03/14/2020 | Greece:Athens |
| MT459911.1 | 03/14/2020 | Greece:Athens |
| MT506201.1 | 03/14/2020 | USA:Michigan |
| MT506203.1 | 03/14/2020 | USA:Michigan |
| MT506204.1 | 03/14/2020 | USA:Michigan |
| MT506205.1 | 03/14/2020 | USA:Michigan |
| MT506206.1 | 03/14/2020 | USA:Michigan |
| MT506532.1 | 03/14/2020 | USA:Michigan |
| MT506533.1 | 03/14/2020 | USA:Michigan |
| MT506534.1 | 03/14/2020 | USA:Michigan |
| MT507273.1 | 03/14/2020 | Jamaica |
| MT509453.1 | 03/14/2020 | USA |
| MT509459.1 | 03/14/2020 | USA |
| MT509464.1 | 03/14/2020 | USA |
| MT509469.1 | 03/14/2020 | USA |
| MT509485.1 | 03/14/2020 | USA |
| MT509490.1 | 03/14/2020 | USA |
| MT509671.1 | 03/14/2020 | USA:Michigan |
| MT509673.1 | 03/14/2020 | USA:Michigan |
| MT510719.1 | 03/14/2020 | USA |
| MT510721.1 | 03/14/2020 | USA |
| MT510744.1 | 03/14/2020 | Hong-Kong |
| MT510745.1 | 03/14/2020 | Hong-Kong |
| MT520422.1 | 03/14/2020 | USA:Massachusetts |
| MT520490.1 | 03/14/2020 | USA:Massachusetts |
| MT582448.1 | 03/14/2020 | Germany:Dusseldorf |
| MT582476.1 | 03/14/2020 | Germany:Dusseldorf |
| MT582479.1 | 03/14/2020 | Germany:Dusseldorf |
| MT582480.1 | 03/14/2020 | Germany:Dusseldorf |
| MT582481.1 | 03/14/2020 | Germany:Dusseldorf |
| MT614477.1 | 03/14/2020 | USA:MA |
| MT614478.1 | 03/14/2020 | USA:MA |
| MT614483.1 | 03/14/2020 | USA:MA |
| MT614485.1 | 03/14/2020 | USA:MA |
| MT614487.1 | 03/14/2020 | USA:MA |
| MT614493.1 | 03/14/2020 | USA:MI |
| MT614495.1 | 03/14/2020 | USA:NJ |
| MT614496.1 | 03/14/2020 | USA:NJ |
| MT614497.1 | 03/14/2020 | USA:NJ |
| MT614498.1 | 03/14/2020 | USA:NJ |
| MT614509.1 | 03/14/2020 | USA:TX |
| MT614531.1 | 03/14/2020 | USA:MA |
| MT614533.1 | 03/14/2020 | USA:MA |

| | | |
|---|---|---|
| MT614534.1 | 03/14/2020 | USA:MA |
| MT614535.1 | 03/14/2020 | USA:MA |
| MT614562.1 | 03/14/2020 | USA:MA |
| MT627634.1 | 03/14/2020 | USA:Washington-Pierce-County |
| MT628231.1 | 03/14/2020 | USA:CA |
| MT632500.1 | 03/14/2020 | USA:Washington-King-County |
| MT632547.1 | 03/14/2020 | USA:Washington-King-County |
| MT632579.1 | 03/14/2020 | USA:Washington-King-County |
| MT632624.1 | 03/14/2020 | USA:Washington-King-County |
| MT646053.1 | 03/14/2020 | USA |
| MT246482.1 | 03/15/2020 | USA:WA |
| MT246483.1 | 03/15/2020 | USA:WA |
| MT246484.1 | 03/15/2020 | USA:WA |
| MT246485.1 | 03/15/2020 | USA:WA |
| MT246487.1 | 03/15/2020 | USA:WA |
| MT246490.1 | 03/15/2020 | USA:WA |
| MT252713.1 | 03/15/2020 | USA:WA |
| MT252719.1 | 03/15/2020 | USA:WA |
| MT252738.1 | 03/15/2020 | USA:WA |
| MT252739.1 | 03/15/2020 | USA:WA |
| MT259245.1 | 03/15/2020 | USA:WA |
| MT259271.1 | 03/15/2020 | USA:WA |
| MT259273.1 | 03/15/2020 | USA:WA |
| MT259276.1 | 03/15/2020 | USA:WA |
| MT259277.1 | 03/15/2020 | USA:WA |
| MT259278.1 | 03/15/2020 | USA:WA |
| MT259280.1 | 03/15/2020 | USA:WA |
| MT259281.1 | 03/15/2020 | USA:WA |
| MT259282.1 | 03/15/2020 | USA:WA |
| MT259285.1 | 03/15/2020 | USA:WA |
| MT259286.1 | 03/15/2020 | USA:WA |
| MT259287.1 | 03/15/2020 | USA:WA |
| MT263457.1 | 03/15/2020 | USA:WA |
| MT263458.1 | 03/15/2020 | USA:WA |
| MT263469.1 | 03/15/2020 | USA:WA |
| MT359865.1 | 03/15/2020 | Spain |
| MT359866.1 | 03/15/2020 | Spain |
| MT370867.1 | 03/15/2020 | USA:NY |
| MT370874.1 | 03/15/2020 | USA:NY |
| MT412154.1 | 03/15/2020 | USA:Michigan |
| MT444524.1 | 03/15/2020 | USA |
| MT444526.1 | 03/15/2020 | USA |
| MT444530.1 | 03/15/2020 | USA |
| MT444533.1 | 03/15/2020 | USA |
| MT444557.1 | 03/15/2020 | USA |
| MT450925.1 | 03/15/2020 | Australia:Victoria |

| | | |
|---|---|---|
| MT450926.1 | 03/15/2020 | Australia:Victoria |
| MT450995.1 | 03/15/2020 | Australia:Victoria |
| MT450996.1 | 03/15/2020 | Australia:Victoria |
| MT450997.1 | 03/15/2020 | Australia:Victoria |
| MT450998.1 | 03/15/2020 | Australia:Victoria |
| MT450999.1 | 03/15/2020 | Australia:Victoria |
| MT451000.1 | 03/15/2020 | Australia:Victoria |
| MT451001.1 | 03/15/2020 | Australia:Victoria |
| MT451002.1 | 03/15/2020 | Australia:Victoria |
| MT451003.1 | 03/15/2020 | Australia:Victoria |
| MT451043.1 | 03/15/2020 | Australia:Victoria |
| MT466583.1 | 03/15/2020 | USA:AK |
| MT477835.1 | 03/15/2020 | USA:AK |
| MT506207.1 | 03/15/2020 | USA:Michigan |
| MT506209.1 | 03/15/2020 | USA:Michigan |
| MT506216.1 | 03/15/2020 | USA:Michigan |
| MT506530.1 | 03/15/2020 | USA:Michigan |
| MT506537.1 | 03/15/2020 | USA:Michigan |
| MT506639.1 | 03/15/2020 | USA:Michigan |
| MT506641.1 | 03/15/2020 | USA:Michigan |
| MT506642.1 | 03/15/2020 | USA:Michigan |
| MT507281.1 | 03/15/2020 | USA |
| MT509465.1 | 03/15/2020 | USA |
| MT509670.1 | 03/15/2020 | USA:Michigan |
| MT509672.1 | 03/15/2020 | USA:Michigan |
| MT509674.1 | 03/15/2020 | USA:Michigan |
| MT510718.1 | 03/15/2020 | USA |
| MT533232.1 | 03/15/2020 | USA:CA |
| MT582468.1 | 03/15/2020 | Germany:Dusseldorf |
| MT582469.1 | 03/15/2020 | Germany:Dusseldorf |
| MT582470.1 | 03/15/2020 | Germany:Dusseldorf |
| MT582471.1 | 03/15/2020 | Germany:Dusseldorf |
| MT582472.1 | 03/15/2020 | Germany:Dusseldorf |
| MT582473.1 | 03/15/2020 | Germany:Dusseldorf |
| MT582474.1 | 03/15/2020 | Germany:Dusseldorf |
| MT582475.1 | 03/15/2020 | Germany:Dusseldorf |
| MT582478.1 | 03/15/2020 | Germany:Dusseldorf |
| MT614451.1 | 03/15/2020 | USA:CA |
| MT614457.1 | 03/15/2020 | USA:CA |
| MT614459.1 | 03/15/2020 | USA:CA |
| MT614460.1 | 03/15/2020 | USA:CA |
| MT614461.1 | 03/15/2020 | USA:CA |
| MT614462.1 | 03/15/2020 | USA:CA |
| MT614466.1 | 03/15/2020 | USA:CA |
| MT614468.1 | 03/15/2020 | USA:CA |
| MT614469.1 | 03/15/2020 | USA:CA |

| | | |
|---|---|---|
| MT614470.1 | 03/15/2020 | USA:CA |
| MT614475.1 | 03/15/2020 | USA:CA |
| MT614479.1 | 03/15/2020 | USA:MA |
| MT614480.1 | 03/15/2020 | USA:MA |
| MT614481.1 | 03/15/2020 | USA:MA |
| MT614482.1 | 03/15/2020 | USA:MA |
| MT614484.1 | 03/15/2020 | USA:MA |
| MT614486.1 | 03/15/2020 | USA:MA |
| MT614489.1 | 03/15/2020 | USA:MI |
| MT614490.1 | 03/15/2020 | USA:MI |
| MT614491.1 | 03/15/2020 | USA:MI |
| MT614492.1 | 03/15/2020 | USA:MI |
| MT614494.1 | 03/15/2020 | USA:NJ |
| MT614500.1 | 03/15/2020 | USA:NY |
| MT614501.1 | 03/15/2020 | USA:NY |
| MT614505.1 | 03/15/2020 | USA:NY |
| MT614507.1 | 03/15/2020 | USA:TX |
| MT614508.1 | 03/15/2020 | USA:TX |
| MT614524.1 | 03/15/2020 | USA:CA |
| MT614540.1 | 03/15/2020 | USA:NJ |
| MT614542.1 | 03/15/2020 | USA:NY |
| MT614546.1 | 03/15/2020 | USA:NY |
| MT614550.1 | 03/15/2020 | USA:NY |
| MT614552.1 | 03/15/2020 | USA:NY |
| MT614553.1 | 03/15/2020 | USA:NY |
| MT614554.1 | 03/15/2020 | USA:NY |
| MT627635.1 | 03/15/2020 | USA:Washington-Pierce-County |
| MT628126.1 | 03/15/2020 | USA:CA |
| MT630421.1 | 03/15/2020 | Saudi-Arabia:Jeddah |
| MT630422.1 | 03/15/2020 | Saudi-Arabia:Jeddah |
| MT630423.1 | 03/15/2020 | Saudi-Arabia:Jeddah |
| MT630424.1 | 03/15/2020 | Saudi-Arabia:Jeddah |
| MT630425.1 | 03/15/2020 | Saudi-Arabia:Jeddah |
| MT630426.1 | 03/15/2020 | Saudi-Arabia:Jeddah |
| MT630427.1 | 03/15/2020 | Saudi-Arabia:Jeddah |
| MT630428.1 | 03/15/2020 | Saudi-Arabia:Jeddah |
| MT630429.1 | 03/15/2020 | Saudi-Arabia:Jeddah |
| MT630430.1 | 03/15/2020 | Saudi-Arabia:Jeddah |
| MT630431.1 | 03/15/2020 | Saudi-Arabia:Jeddah |
| MT630432.1 | 03/15/2020 | Saudi-Arabia:Jeddah |
| MT632591.1 | 03/15/2020 | USA:Washington-King-County |
| MT632596.1 | 03/15/2020 | USA:Washington-King-County |
| MT632621.1 | 03/15/2020 | USA:Washington-King-County |
| MT635227.1 | 03/15/2020 | USA:Connecticut |
| MT646065.1 | 03/15/2020 | USA |
| MT646077.1 | 03/15/2020 | USA |

| | | |
|---|---|---|
| MT259236.1 | 03/16/2020 | USA:WA |
| MT259237.1 | 03/16/2020 | USA:WA |
| MT259239.1 | 03/16/2020 | USA:WA |
| MT259240.1 | 03/16/2020 | USA:WA |
| MT259241.1 | 03/16/2020 | USA:WA |
| MT259244.1 | 03/16/2020 | USA:WA |
| MT259247.1 | 03/16/2020 | USA:WA |
| MT259253.1 | 03/16/2020 | USA:WA |
| MT259254.1 | 03/16/2020 | USA:WA |
| MT259257.1 | 03/16/2020 | USA:WA |
| MT259260.1 | 03/16/2020 | USA:WA |
| MT259261.1 | 03/16/2020 | USA:WA |
| MT263387.1 | 03/16/2020 | USA:WA |
| MT263388.1 | 03/16/2020 | USA:WA |
| MT263391.1 | 03/16/2020 | USA:WA |
| MT263403.1 | 03/16/2020 | USA:WA |
| MT263405.1 | 03/16/2020 | USA:WA |
| MT263410.1 | 03/16/2020 | USA:WA |
| MT263412.1 | 03/16/2020 | USA:WA |
| MT263413.1 | 03/16/2020 | USA:WA |
| MT263416.1 | 03/16/2020 | USA:WA |
| MT263418.1 | 03/16/2020 | USA:WA |
| MT263419.1 | 03/16/2020 | USA:WA |
| MT263422.1 | 03/16/2020 | USA:WA |
| MT263426.1 | 03/16/2020 | USA:WA |
| MT263427.1 | 03/16/2020 | USA:WA |
| MT263448.1 | 03/16/2020 | USA:WA |
| MT263451.1 | 03/16/2020 | USA:WA |
| MT263463.1 | 03/16/2020 | USA:CT |
| MT263464.1 | 03/16/2020 | USA:WA |
| MT263465.1 | 03/16/2020 | USA:WA |
| MT263467.1 | 03/16/2020 | USA:WA |
| MT270815.1 | 03/16/2020 | Hong-Kong |
| MT293158.1 | 03/16/2020 | USA:WA |
| MT293159.1 | 03/16/2020 | USA:WA |
| MT293160.1 | 03/16/2020 | USA:WA |
| MT293161.1 | 03/16/2020 | USA:WA |
| MT293162.1 | 03/16/2020 | USA:WA |
| MT293177.1 | 03/16/2020 | USA |
| MT326091.1 | 03/16/2020 | USA |
| MT326188.1 | 03/16/2020 | USA |
| MT334543.1 | 03/16/2020 | USA:UT |
| MT370846.1 | 03/16/2020 | USA:NY |
| MT370862.1 | 03/16/2020 | USA:NY |
| MT370873.1 | 03/16/2020 | USA:NY |
| MT370878.1 | 03/16/2020 | USA:NY |

| | | |
|---|---|---|
| MT370884.1 | 03/16/2020 | USA:NY |
| MT370891.1 | 03/16/2020 | USA:NY |
| MT370895.1 | 03/16/2020 | USA:NY |
| MT370900.1 | 03/16/2020 | USA:NY |
| MT370918.1 | 03/16/2020 | USA:NY |
| MT371047.1 | 03/16/2020 | Sri-Lanka |
| MT374112.1 | 03/16/2020 | Taiwan |
| MT412146.1 | 03/16/2020 | USA:Michigan |
| MT412147.1 | 03/16/2020 | USA:Michigan |
| MT412153.1 | 03/16/2020 | USA:Michigan |
| MT412176.1 | 03/16/2020 | USA:Michigan |
| MT415322.1 | 03/16/2020 | India |
| MT439247.1 | 03/16/2020 | USA:MI |
| MT439248.1 | 03/16/2020 | USA:MI |
| MT439258.1 | 03/16/2020 | USA:MI |
| MT444515.1 | 03/16/2020 | USA |
| MT444516.1 | 03/16/2020 | USA |
| MT444517.1 | 03/16/2020 | USA |
| MT444518.1 | 03/16/2020 | USA |
| MT444519.1 | 03/16/2020 | USA |
| MT444520.1 | 03/16/2020 | USA |
| MT444522.1 | 03/16/2020 | USA |
| MT444525.1 | 03/16/2020 | USA |
| MT447171.1 | 03/16/2020 | Thailand |
| MT450927.1 | 03/16/2020 | Australia:Victoria |
| MT450928.1 | 03/16/2020 | Australia:Victoria |
| MT450929.1 | 03/16/2020 | Australia:Victoria |
| MT451004.1 | 03/16/2020 | Australia:Victoria |
| MT451005.1 | 03/16/2020 | Australia:Victoria |
| MT451007.1 | 03/16/2020 | Australia:Victoria |
| MT451008.1 | 03/16/2020 | Australia:Victoria |
| MT451009.1 | 03/16/2020 | Australia:Victoria |
| MT451010.1 | 03/16/2020 | Australia:Victoria |
| MT451011.1 | 03/16/2020 | Australia:Victoria |
| MT451012.1 | 03/16/2020 | Australia:Victoria |
| MT451044.1 | 03/16/2020 | Australia:Victoria |
| MT451045.1 | 03/16/2020 | Australia:Victoria |
| MT451046.1 | 03/16/2020 | Australia:Victoria |
| MT451047.1 | 03/16/2020 | Australia:Victoria |
| MT451567.1 | 03/16/2020 | Australia:Victoria |
| MT459859.1 | 03/16/2020 | Greece:Athens |
| MT459860.1 | 03/16/2020 | Greece:Athens |
| MT459861.1 | 03/16/2020 | Greece:Athens |
| MT459869.1 | 03/16/2020 | Greece:Athens |
| MT459883.1 | 03/16/2020 | Greece:Athens |
| MT459884.1 | 03/16/2020 | Greece:Athens |

| | | |
|---|---|---|
| MT477836.1 | 03/16/2020 | USA:AK |
| MT506211.1 | 03/16/2020 | USA:Michigan |
| MT506535.1 | 03/16/2020 | USA:Michigan |
| MT506536.1 | 03/16/2020 | USA:Michigan |
| MT506638.1 | 03/16/2020 | USA:Michigan |
| MT506885.1 | 03/16/2020 | USA:Wisconsin |
| MT507274.1 | 03/16/2020 | Jamaica |
| MT507275.1 | 03/16/2020 | Jamaica |
| MT507794.1 | 03/16/2020 | Jamaica |
| MT509488.1 | 03/16/2020 | USA |
| MT520299.1 | 03/16/2020 | USA:Massachusetts |
| MT582462.1 | 03/16/2020 | Germany:Dusseldorf |
| MT582463.1 | 03/16/2020 | Germany:Dusseldorf |
| MT582464.1 | 03/16/2020 | Germany:Dusseldorf |
| MT582465.1 | 03/16/2020 | Germany:Dusseldorf |
| MT582466.1 | 03/16/2020 | Germany:Dusseldorf |
| MT582467.1 | 03/16/2020 | Germany:Dusseldorf |
| MT582477.1 | 03/16/2020 | Germany:Dusseldorf |
| MT614447.1 | 03/16/2020 | USA:CA |
| MT614448.1 | 03/16/2020 | USA:CA |
| MT614449.1 | 03/16/2020 | USA:CA |
| MT614450.1 | 03/16/2020 | USA:CA |
| MT614452.1 | 03/16/2020 | USA:CA |
| MT614453.1 | 03/16/2020 | USA:CA |
| MT614454.1 | 03/16/2020 | USA:CA |
| MT614455.1 | 03/16/2020 | USA:CA |
| MT614456.1 | 03/16/2020 | USA:CA |
| MT614463.1 | 03/16/2020 | USA:CA |
| MT614465.1 | 03/16/2020 | USA:CA |
| MT614467.1 | 03/16/2020 | USA:CA |
| MT614472.1 | 03/16/2020 | USA:CA |
| MT614473.1 | 03/16/2020 | USA:CA |
| MT614474.1 | 03/16/2020 | USA:CA |
| MT614499.1 | 03/16/2020 | USA:NY |
| MT614502.1 | 03/16/2020 | USA:NY |
| MT614503.1 | 03/16/2020 | USA:NY |
| MT614523.1 | 03/16/2020 | USA:CA |
| MT614525.1 | 03/16/2020 | USA:CA |
| MT614526.1 | 03/16/2020 | USA:CA |
| MT614538.1 | 03/16/2020 | USA:NJ |
| MT614543.1 | 03/16/2020 | USA:NY |
| MT614544.1 | 03/16/2020 | USA:NY |
| MT614547.1 | 03/16/2020 | USA:NY |
| MT614548.1 | 03/16/2020 | USA:NY |
| MT614549.1 | 03/16/2020 | USA:NY |
| MT614551.1 | 03/16/2020 | USA:NY |

| | | |
|---|---|---|
| MT628125.1 | 03/16/2020 | USA:CA |
| MT632495.1 | 03/16/2020 | USA:Washington-Snohomish-County |
| MT632501.1 | 03/16/2020 | USA:Washington-Snohomish-County |
| MT632535.1 | 03/16/2020 | USA:Washington-King-County |
| MT632543.1 | 03/16/2020 | USA:Washington-King-County |
| MT632560.1 | 03/16/2020 | USA:Washington-King-County |
| MT632614.1 | 03/16/2020 | USA:Washington-King-County |
| MT632622.1 | 03/16/2020 | USA:Washington-Snohomish-County |
| MT635229.1 | 03/16/2020 | USA:Connecticut |
| MT646074.1 | 03/16/2020 | USA |
| MT646076.1 | 03/16/2020 | USA |
| MT646091.1 | 03/16/2020 | USA |
| MT646094.1 | 03/16/2020 | USA |
| MT646097.1 | 03/16/2020 | USA |
| MT646098.1 | 03/16/2020 | USA |
| LC547526.1 | 03/17/2020 | Japan:Saitama |
| MT263452.1 | 03/17/2020 | USA:WA |
| MT263468.1 | 03/17/2020 | USA:WA |
| MT293164.1 | 03/17/2020 | USA:WA |
| MT293165.1 | 03/17/2020 | USA:WA |
| MT293166.1 | 03/17/2020 | USA:WA |
| MT293167.1 | 03/17/2020 | USA:WA |
| MT293168.1 | 03/17/2020 | USA:WA |
| MT293169.1 | 03/17/2020 | USA:WA |
| MT293171.1 | 03/17/2020 | USA:WA |
| MT293172.1 | 03/17/2020 | USA:WA |
| MT293176.1 | 03/17/2020 | USA:WA |
| MT293179.1 | 03/17/2020 | USA:WA |
| MT293180.1 | 03/17/2020 | USA:WA |
| MT293187.1 | 03/17/2020 | USA:WA |
| MT293190.1 | 03/17/2020 | USA:WA |
| MT293191.1 | 03/17/2020 | USA:WA |
| MT326183.1 | 03/17/2020 | USA |
| MT326184.1 | 03/17/2020 | USA |
| MT326185.1 | 03/17/2020 | USA |
| MT327745.1 | 03/17/2020 | Turkey |
| MT339039.1 | 03/17/2020 | USA:Maricopa-County-Arizona |
| MT339041.1 | 03/17/2020 | USA:Maricopa-County-Arizona |
| MT370832.1 | 03/17/2020 | USA:NY |
| MT370838.1 | 03/17/2020 | USA:NY |
| MT370840.1 | 03/17/2020 | USA:NY |
| MT370841.1 | 03/17/2020 | USA:NY |
| MT370843.1 | 03/17/2020 | USA:NY |
| MT370849.1 | 03/17/2020 | USA:NY |
| MT370850.1 | 03/17/2020 | USA:NY |
| MT370852.1 | 03/17/2020 | USA:NY |

| | | |
|---|---|---|
| MT370853.1 | 03/17/2020 | USA:NY |
| MT370859.1 | 03/17/2020 | USA:NY |
| MT370861.1 | 03/17/2020 | USA:NY |
| MT370865.1 | 03/17/2020 | USA:NY |
| MT370866.1 | 03/17/2020 | USA:NY |
| MT370868.1 | 03/17/2020 | USA:NY |
| MT370871.1 | 03/17/2020 | USA:NY |
| MT370872.1 | 03/17/2020 | USA:NY |
| MT370876.1 | 03/17/2020 | USA:NY |
| MT370877.1 | 03/17/2020 | USA:NY |
| MT370879.1 | 03/17/2020 | USA:NY |
| MT370886.1 | 03/17/2020 | USA:NY |
| MT370888.1 | 03/17/2020 | USA:NY |
| MT370893.1 | 03/17/2020 | USA:NY |
| MT370894.1 | 03/17/2020 | USA:NY |
| MT370899.1 | 03/17/2020 | USA:NY |
| MT370901.1 | 03/17/2020 | USA:NY |
| MT370902.1 | 03/17/2020 | USA:NY |
| MT370903.1 | 03/17/2020 | USA:NY |
| MT370905.1 | 03/17/2020 | USA:NY |
| MT370906.1 | 03/17/2020 | USA:NY |
| MT370907.1 | 03/17/2020 | USA:NY |
| MT370909.1 | 03/17/2020 | USA:NY |
| MT370911.1 | 03/17/2020 | USA:NY |
| MT370913.1 | 03/17/2020 | USA:NY |
| MT370914.1 | 03/17/2020 | USA:NY |
| MT370916.1 | 03/17/2020 | USA:NY |
| MT371573.1 | 03/17/2020 | Czech-Republic |
| MT374109.1 | 03/17/2020 | Taiwan |
| MT374110.1 | 03/17/2020 | Taiwan |
| MT374111.1 | 03/17/2020 | Taiwan |
| MT374113.1 | 03/17/2020 | Taiwan |
| MT412148.1 | 03/17/2020 | USA:Michigan |
| MT412149.1 | 03/17/2020 | USA:Michigan |
| MT412151.1 | 03/17/2020 | USA:Michigan |
| MT412152.1 | 03/17/2020 | USA:Michigan |
| MT412155.1 | 03/17/2020 | USA:Michigan |
| MT412156.1 | 03/17/2020 | USA:Michigan |
| MT412157.1 | 03/17/2020 | USA:Michigan |
| MT434785.1 | 03/17/2020 | USA:NY |
| MT434790.1 | 03/17/2020 | USA:NY |
| MT434791.1 | 03/17/2020 | USA:NY |
| MT434792.1 | 03/17/2020 | USA:NY |
| MT434797.1 | 03/17/2020 | USA:NY |
| MT434803.1 | 03/17/2020 | USA:NY |
| MT434804.1 | 03/17/2020 | USA:NY |

| | | |
|---|---|---|
| MT434811.1 | 03/17/2020 | USA:NY |
| MT439249.1 | 03/17/2020 | USA:MI |
| MT439250.1 | 03/17/2020 | USA:MI |
| MT439253.1 | 03/17/2020 | USA:MI |
| MT444521.1 | 03/17/2020 | USA |
| MT444523.1 | 03/17/2020 | USA |
| MT444527.1 | 03/17/2020 | USA |
| MT444528.1 | 03/17/2020 | USA |
| MT444529.1 | 03/17/2020 | USA |
| MT451013.1 | 03/17/2020 | Australia:Victoria |
| MT451014.1 | 03/17/2020 | Australia:Victoria |
| MT451015.1 | 03/17/2020 | Australia:Victoria |
| MT451027.1 | 03/17/2020 | Australia:Victoria |
| MT451048.1 | 03/17/2020 | Australia:Victoria |
| MT451049.1 | 03/17/2020 | Australia:Victoria |
| MT451050.1 | 03/17/2020 | Australia:Victoria |
| MT451051.1 | 03/17/2020 | Australia:Victoria |
| MT451052.1 | 03/17/2020 | Australia:Victoria |
| MT451053.1 | 03/17/2020 | Australia:Victoria |
| MT451054.1 | 03/17/2020 | Australia:Victoria |
| MT451055.1 | 03/17/2020 | Australia:Victoria |
| MT451056.1 | 03/17/2020 | Australia:Victoria |
| MT451057.1 | 03/17/2020 | Australia:Victoria |
| MT451058.1 | 03/17/2020 | Australia:Victoria |
| MT451059.1 | 03/17/2020 | Australia:Victoria |
| MT451060.1 | 03/17/2020 | Australia:Victoria |
| MT451061.1 | 03/17/2020 | Australia:Victoria |
| MT451219.1 | 03/17/2020 | Australia:Victoria |
| MT451220.1 | 03/17/2020 | Australia:Victoria |
| MT451221.1 | 03/17/2020 | Australia:Victoria |
| MT459885.1 | 03/17/2020 | Greece:Athens |
| MT459886.1 | 03/17/2020 | Greece:Athens |
| MT459891.1 | 03/17/2020 | Greece:Athens |
| MT477837.1 | 03/17/2020 | USA:AK |
| MT477838.1 | 03/17/2020 | USA:AK |
| MT506212.1 | 03/17/2020 | USA:Michigan |
| MT506213.1 | 03/17/2020 | USA:Michigan |
| MT506640.1 | 03/17/2020 | USA:Michigan |
| MT506643.1 | 03/17/2020 | USA:Michigan |
| MT506644.1 | 03/17/2020 | USA:Michigan |
| MT506648.1 | 03/17/2020 | USA:Michigan |
| MT506650.1 | 03/17/2020 | USA:Michigan |
| MT506652.1 | 03/17/2020 | USA:Michigan |
| MT507276.1 | 03/17/2020 | Jamaica |
| MT509466.1 | 03/17/2020 | USA |
| MT509471.1 | 03/17/2020 | USA |

| | | |
|---|---|---|
| MT509476.1 | 03/17/2020 | USA |
| MT509487.1 | 03/17/2020 | USA |
| MT509489.1 | 03/17/2020 | USA |
| MT509492.1 | 03/17/2020 | USA |
| MT512447.1 | 03/17/2020 | USA |
| MT520286.1 | 03/17/2020 | USA:Massachusetts |
| MT549878.1 | 03/17/2020 | Kenya |
| MT582460.1 | 03/17/2020 | Germany:Dusseldorf |
| MT582461.1 | 03/17/2020 | Germany:Dusseldorf |
| MT590598.1 | 03/17/2020 | Taiwan |
| MT627620.1 | 03/17/2020 | USA:Washington |
| MT628128.1 | 03/17/2020 | USA:CA |
| MT628129.1 | 03/17/2020 | USA:CA |
| MT632559.1 | 03/17/2020 | USA:Washington-King-County |
| MT632589.1 | 03/17/2020 | USA:Washington-Yakima-County |
| MT646064.1 | 03/17/2020 | USA |
| MT646096.1 | 03/17/2020 | USA |
| MT646099.1 | 03/17/2020 | USA |
| LC547527.1 | 03/18/2020 | Japan:Saitama |
| MT258377.1 | 03/18/2020 | USA:San-Francisco-CA |
| MT258378.1 | 03/18/2020 | USA:San-Francisco-CA |
| MT258379.1 | 03/18/2020 | USA:San-Francisco-CA |
| MT258380.1 | 03/18/2020 | USA:San-Francisco-CA |
| MT258381.1 | 03/18/2020 | USA:San-Francisco-CA |
| MT258382.1 | 03/18/2020 | USA:San-Francisco-CA |
| MT258383.1 | 03/18/2020 | USA:San-Francisco-CA |
| MT270814.1 | 03/18/2020 | Hong-Kong |
| MT293170.1 | 03/18/2020 | USA:WA |
| MT293181.1 | 03/18/2020 | USA:WA |
| MT293182.1 | 03/18/2020 | USA:WA |
| MT293183.1 | 03/18/2020 | USA:WA |
| MT293184.1 | 03/18/2020 | USA:WA |
| MT293189.1 | 03/18/2020 | USA:WA |
| MT293194.1 | 03/18/2020 | USA:WA |
| MT293204.1 | 03/18/2020 | USA:WA |
| MT293218.1 | 03/18/2020 | USA:WA |
| MT293219.1 | 03/18/2020 | USA:WA |
| MT293220.1 | 03/18/2020 | USA:WA |
| MT293221.1 | 03/18/2020 | USA:WA |
| MT293222.1 | 03/18/2020 | USA:WA |
| MT293223.1 | 03/18/2020 | USA:WA |
| MT293224.1 | 03/18/2020 | USA:WA |
| MT293225.1 | 03/18/2020 | USA:WA |
| MT326187.1 | 03/18/2020 | USA |
| MT326189.1 | 03/18/2020 | USA |
| MT326190.1 | 03/18/2020 | USA |

| | | |
|---|---|---|
| MT326191.1 | 03/18/2020 | USA |
| MT328032.1 | 03/18/2020 | Greece |
| MT328033.1 | 03/18/2020 | Greece |
| MT328034.1 | 03/18/2020 | Greece |
| MT328035.1 | 03/18/2020 | Greece |
| MT350282.1 | 03/18/2020 | Brazil |
| MT365025.1 | 03/18/2020 | USA:KY |
| MT365026.1 | 03/18/2020 | USA:KY |
| MT365027.1 | 03/18/2020 | USA:KY |
| MT370831.1 | 03/18/2020 | USA:NY |
| MT370833.1 | 03/18/2020 | USA:NY |
| MT370834.1 | 03/18/2020 | USA:NY |
| MT370835.1 | 03/18/2020 | USA:NY |
| MT370836.1 | 03/18/2020 | USA:NY |
| MT370837.1 | 03/18/2020 | USA:NY |
| MT370844.1 | 03/18/2020 | USA:NY |
| MT370845.1 | 03/18/2020 | USA:NY |
| MT370847.1 | 03/18/2020 | USA:NY |
| MT370851.1 | 03/18/2020 | USA:NY |
| MT370856.1 | 03/18/2020 | USA:NY |
| MT370858.1 | 03/18/2020 | USA:NY |
| MT370860.1 | 03/18/2020 | USA:NY |
| MT370863.1 | 03/18/2020 | USA:NY |
| MT370864.1 | 03/18/2020 | USA:NY |
| MT370869.1 | 03/18/2020 | USA:NY |
| MT370870.1 | 03/18/2020 | USA:NY |
| MT370875.1 | 03/18/2020 | USA:NY |
| MT370882.1 | 03/18/2020 | USA:NY |
| MT370883.1 | 03/18/2020 | USA:NY |
| MT370885.1 | 03/18/2020 | USA:NY |
| MT370887.1 | 03/18/2020 | USA:NY |
| MT370890.1 | 03/18/2020 | USA:NY |
| MT370896.1 | 03/18/2020 | USA:NY |
| MT370897.1 | 03/18/2020 | USA:NY |
| MT370898.1 | 03/18/2020 | USA:NY |
| MT370908.1 | 03/18/2020 | USA:NY |
| MT370920.1 | 03/18/2020 | USA:NY |
| MT370926.1 | 03/18/2020 | USA:NY |
| MT370957.1 | 03/18/2020 | USA:NY |
| MT370960.1 | 03/18/2020 | USA:NY |
| MT370987.1 | 03/18/2020 | USA:NY |
| MT371571.1 | 03/18/2020 | Czech-Republic |
| MT371574.1 | 03/18/2020 | Czech-Republic |
| MT372480.1 | 03/18/2020 | Malaysia |
| MT374114.1 | 03/18/2020 | Taiwan |
| MT374115.1 | 03/18/2020 | Taiwan |

| | | |
|---|---|---|
| MT374116.1 | 03/18/2020 | Taiwan |
| MT385431.1 | 03/18/2020 | USA:CA |
| MT412158.1 | 03/18/2020 | USA:Michigan |
| MT412160.1 | 03/18/2020 | USA:Michigan |
| MT412161.1 | 03/18/2020 | USA:Michigan |
| MT412169.1 | 03/18/2020 | USA:Michigan |
| MT434810.1 | 03/18/2020 | USA:NY |
| MT439252.1 | 03/18/2020 | USA:MI |
| MT439254.1 | 03/18/2020 | USA:MI |
| MT439265.1 | 03/18/2020 | USA:MI |
| MT447172.1 | 03/18/2020 | Thailand |
| MT447173.1 | 03/18/2020 | Thailand |
| MT451017.1 | 03/18/2020 | Australia:Victoria |
| MT451018.1 | 03/18/2020 | Australia:Victoria |
| MT451019.1 | 03/18/2020 | Australia:Victoria |
| MT451020.1 | 03/18/2020 | Australia:Victoria |
| MT451021.1 | 03/18/2020 | Australia:Victoria |
| MT451022.1 | 03/18/2020 | Australia:Victoria |
| MT451024.1 | 03/18/2020 | Australia:Victoria |
| MT451031.1 | 03/18/2020 | Australia:Victoria |
| MT451062.1 | 03/18/2020 | Australia:Victoria |
| MT451063.1 | 03/18/2020 | Australia:Victoria |
| MT451064.1 | 03/18/2020 | Australia:Victoria |
| MT451065.1 | 03/18/2020 | Australia:Victoria |
| MT451066.1 | 03/18/2020 | Australia:Victoria |
| MT451067.1 | 03/18/2020 | Australia:Victoria |
| MT451068.1 | 03/18/2020 | Australia:Victoria |
| MT451069.1 | 03/18/2020 | Australia:Victoria |
| MT451070.1 | 03/18/2020 | Australia:Victoria |
| MT451071.1 | 03/18/2020 | Australia:Victoria |
| MT451072.1 | 03/18/2020 | Australia:Victoria |
| MT451073.1 | 03/18/2020 | Australia:Victoria |
| MT451074.1 | 03/18/2020 | Australia:Victoria |
| MT451075.1 | 03/18/2020 | Australia:Victoria |
| MT451076.1 | 03/18/2020 | Australia:Victoria |
| MT451077.1 | 03/18/2020 | Australia:Victoria |
| MT451078.1 | 03/18/2020 | Australia:Victoria |
| MT451079.1 | 03/18/2020 | Australia:Victoria |
| MT451080.1 | 03/18/2020 | Australia:Victoria |
| MT451081.1 | 03/18/2020 | Australia:Victoria |
| MT451222.1 | 03/18/2020 | Australia:Victoria |
| MT451223.1 | 03/18/2020 | Australia:Victoria |
| MT451224.1 | 03/18/2020 | Australia:Victoria |
| MT451533.1 | 03/18/2020 | Australia:Victoria |
| MT451537.1 | 03/18/2020 | Australia:Victoria |
| MT451538.1 | 03/18/2020 | Australia:Victoria |

| | | |
|---|---|---|
| MT451541.1 | 03/18/2020 | Australia:Victoria |
| MT459887.1 | 03/18/2020 | Greece:Athens |
| MT459888.1 | 03/18/2020 | Greece:Athens |
| MT459892.1 | 03/18/2020 | Greece:Athens |
| MT459893.1 | 03/18/2020 | Greece:Athens |
| MT459906.1 | 03/18/2020 | Greece:Athens |
| MT477840.1 | 03/18/2020 | USA:AK |
| MT477841.1 | 03/18/2020 | USA:AK |
| MT499215.1 | 03/18/2020 | Tunisia |
| MT506538.1 | 03/18/2020 | USA:Michigan |
| MT506645.1 | 03/18/2020 | USA:Michigan |
| MT506647.1 | 03/18/2020 | USA:Michigan |
| MT506649.1 | 03/18/2020 | USA:Michigan |
| MT506651.1 | 03/18/2020 | USA:Michigan |
| MT506657.1 | 03/18/2020 | USA:Michigan |
| MT509486.1 | 03/18/2020 | USA |
| MT509675.1 | 03/18/2020 | USA:Michigan |
| MT509676.1 | 03/18/2020 | USA:Michigan |
| MT520253.1 | 03/18/2020 | USA:Massachusetts |
| MT520337.1 | 03/18/2020 | USA:Massachusetts |
| MT520419.1 | 03/18/2020 | USA:Massachusetts |
| MT520456.1 | 03/18/2020 | USA:Massachusetts |
| MT520488.1 | 03/18/2020 | USA:Massachusetts |
| MT520533.1 | 03/18/2020 | USA:Massachusetts |
| MT582458.1 | 03/18/2020 | Germany:Dusseldorf |
| MT582459.1 | 03/18/2020 | Germany:Dusseldorf |
| MT598172.1 | 03/18/2020 | USA:San-Diego-California |
| MT606515.1 | 03/18/2020 | USA:Maricopa-County-Arizona |
| MT628219.1 | 03/18/2020 | USA:CA |
| MT628232.1 | 03/18/2020 | USA:CA |
| MT628233.1 | 03/18/2020 | USA:CA |
| MT632504.1 | 03/18/2020 | USA:Washington-Whatcom-County |
| MT632531.1 | 03/18/2020 | USA:Washington-King-County |
| MT632533.1 | 03/18/2020 | USA:Washington-Whatcom-County |
| MT632536.1 | 03/18/2020 | USA:Washington-Whatcom-County |
| MT637144.1 | 03/18/2020 | Russia:Moscow-region |
| MT646082.1 | 03/18/2020 | USA |
| MT646084.1 | 03/18/2020 | USA |
| MT646090.1 | 03/18/2020 | USA |
| MT646114.1 | 03/18/2020 | USA |
| LC547530.1 | 03/19/2020 | Japan:Saitama |
| MT293196.1 | 03/19/2020 | USA:WA |
| MT293197.1 | 03/19/2020 | USA:WA |
| MT293198.1 | 03/19/2020 | USA:WA |
| MT293199.1 | 03/19/2020 | USA:WA |
| MT293200.1 | 03/19/2020 | USA:WA |

| | | |
|---|---|---|
| MT293201.1 | 03/19/2020 | USA:WA |
| MT293202.1 | 03/19/2020 | USA:WA |
| MT293205.1 | 03/19/2020 | USA:WA |
| MT293206.1 | 03/19/2020 | USA:WA |
| MT293207.1 | 03/19/2020 | USA:WA |
| MT293208.1 | 03/19/2020 | USA:WA |
| MT293209.1 | 03/19/2020 | USA:WA |
| MT293210.1 | 03/19/2020 | USA:WA |
| MT293211.1 | 03/19/2020 | USA:WA |
| MT293212.1 | 03/19/2020 | USA:WA |
| MT293213.1 | 03/19/2020 | USA:WA |
| MT293214.1 | 03/19/2020 | USA:WA |
| MT293215.1 | 03/19/2020 | USA:WA |
| MT293216.1 | 03/19/2020 | USA:WA |
| MT318827.1 | 03/19/2020 | null |
| MT326086.1 | 03/19/2020 | USA |
| MT326136.1 | 03/19/2020 | USA |
| MT326137.1 | 03/19/2020 | USA |
| MT326141.1 | 03/19/2020 | USA |
| MT326174.1 | 03/19/2020 | USA |
| MT326175.1 | 03/19/2020 | USA |
| MT326177.1 | 03/19/2020 | USA |
| MT326178.1 | 03/19/2020 | USA |
| MT326179.1 | 03/19/2020 | USA |
| MT326180.1 | 03/19/2020 | USA |
| MT334544.1 | 03/19/2020 | USA:UT |
| MT334545.1 | 03/19/2020 | USA:UT |
| MT334546.1 | 03/19/2020 | USA:UT |
| MT334547.1 | 03/19/2020 | USA:UT |
| MT334554.1 | 03/19/2020 | USA:UT |
| MT334558.1 | 03/19/2020 | USA:UT |
| MT350263.1 | 03/19/2020 | USA:WA |
| MT350264.1 | 03/19/2020 | USA:WA |
| MT350265.1 | 03/19/2020 | USA:WA |
| MT350266.1 | 03/19/2020 | USA:WA |
| MT350267.1 | 03/19/2020 | USA:WA |
| MT350268.1 | 03/19/2020 | USA:WA |
| MT350270.1 | 03/19/2020 | USA:WA |
| MT350272.1 | 03/19/2020 | USA:WA |
| MT350273.1 | 03/19/2020 | USA:WA |
| MT350274.1 | 03/19/2020 | USA:WA |
| MT350275.1 | 03/19/2020 | USA:WA |
| MT350276.1 | 03/19/2020 | USA:WA |
| MT350277.1 | 03/19/2020 | USA:WA |
| MT350278.1 | 03/19/2020 | USA:WA |
| MT350279.1 | 03/19/2020 | USA:WA |

| | | |
|---|---|---|
| MT350280.1 | 03/19/2020 | USA:WA |
| MT370855.1 | 03/19/2020 | USA:NY |
| MT370880.1 | 03/19/2020 | USA:NY |
| MT370912.1 | 03/19/2020 | USA:NY |
| MT370917.1 | 03/19/2020 | USA:NY |
| MT370928.1 | 03/19/2020 | USA:NY |
| MT370939.1 | 03/19/2020 | USA:NY |
| MT370940.1 | 03/19/2020 | USA:NY |
| MT370942.1 | 03/19/2020 | USA:NY |
| MT370944.1 | 03/19/2020 | USA:NY |
| MT370951.1 | 03/19/2020 | USA:NY |
| MT370953.1 | 03/19/2020 | USA:NY |
| MT370955.1 | 03/19/2020 | USA:NY |
| MT370956.1 | 03/19/2020 | USA:NY |
| MT370958.1 | 03/19/2020 | USA:NY |
| MT370963.1 | 03/19/2020 | USA:NY |
| MT370965.1 | 03/19/2020 | USA:NY |
| MT370966.1 | 03/19/2020 | USA:NY |
| MT370975.1 | 03/19/2020 | USA:NY |
| MT370981.1 | 03/19/2020 | USA:NY |
| MT370982.1 | 03/19/2020 | USA:NY |
| MT370989.1 | 03/19/2020 | USA:NY |
| MT370993.1 | 03/19/2020 | USA:NY |
| MT370996.1 | 03/19/2020 | USA:NY |
| MT370997.1 | 03/19/2020 | USA:NY |
| MT370998.1 | 03/19/2020 | USA:NY |
| MT371001.1 | 03/19/2020 | USA:NY |
| MT371005.1 | 03/19/2020 | USA:NY |
| MT371007.1 | 03/19/2020 | USA:NY |
| MT371010.1 | 03/19/2020 | USA:NY |
| MT371012.1 | 03/19/2020 | USA:NY |
| MT371020.1 | 03/19/2020 | USA:NY |
| MT371026.1 | 03/19/2020 | USA:NY |
| MT371030.1 | 03/19/2020 | USA:NY |
| MT371032.1 | 03/19/2020 | USA:NY |
| MT371037.1 | 03/19/2020 | USA:NY |
| MT371038.1 | 03/19/2020 | USA:NY |
| MT371049.1 | 03/19/2020 | Sri-Lanka |
| MT371569.1 | 03/19/2020 | Czech-Republic |
| MT371572.1 | 03/19/2020 | Czech-Republic |
| MT412159.1 | 03/19/2020 | USA:Michigan |
| MT412162.1 | 03/19/2020 | USA:Michigan |
| MT412163.1 | 03/19/2020 | USA:Michigan |
| MT412164.1 | 03/19/2020 | USA:Michigan |
| MT412165.1 | 03/19/2020 | USA:Michigan |
| MT412166.1 | 03/19/2020 | USA:Michigan |

| | | |
|---|---|---|
| MT412167.1 | 03/19/2020 | USA:Michigan |
| MT412168.1 | 03/19/2020 | USA:Michigan |
| MT412170.1 | 03/19/2020 | USA:Michigan |
| MT412175.1 | 03/19/2020 | USA:Michigan |
| MT434808.1 | 03/19/2020 | USA:NY |
| MT434809.1 | 03/19/2020 | USA:NY |
| MT434812.1 | 03/19/2020 | USA:NY |
| MT434817.1 | 03/19/2020 | USA:NY |
| MT439251.1 | 03/19/2020 | USA:MI |
| MT439255.1 | 03/19/2020 | USA:MI |
| MT439256.1 | 03/19/2020 | USA:MI |
| MT447175.1 | 03/19/2020 | Thailand |
| MT451016.1 | 03/19/2020 | Australia:Victoria |
| MT451025.1 | 03/19/2020 | Australia:Victoria |
| MT451026.1 | 03/19/2020 | Australia:Victoria |
| MT451029.1 | 03/19/2020 | Australia:Victoria |
| MT451030.1 | 03/19/2020 | Australia:Victoria |
| MT451083.1 | 03/19/2020 | Australia:Victoria |
| MT451084.1 | 03/19/2020 | Australia:Victoria |
| MT451085.1 | 03/19/2020 | Australia:Victoria |
| MT451086.1 | 03/19/2020 | Australia:Victoria |
| MT451087.1 | 03/19/2020 | Australia:Victoria |
| MT451088.1 | 03/19/2020 | Australia:Victoria |
| MT451089.1 | 03/19/2020 | Australia:Victoria |
| MT451090.1 | 03/19/2020 | Australia:Victoria |
| MT451091.1 | 03/19/2020 | Australia:Victoria |
| MT451092.1 | 03/19/2020 | Australia:Victoria |
| MT451093.1 | 03/19/2020 | Australia:Victoria |
| MT451094.1 | 03/19/2020 | Australia:Victoria |
| MT451095.1 | 03/19/2020 | Australia:Victoria |
| MT451096.1 | 03/19/2020 | Australia:Victoria |
| MT451097.1 | 03/19/2020 | Australia:Victoria |
| MT451098.1 | 03/19/2020 | Australia:Victoria |
| MT451099.1 | 03/19/2020 | Australia:Victoria |
| MT451100.1 | 03/19/2020 | Australia:Victoria |
| MT451101.1 | 03/19/2020 | Australia:Victoria |
| MT451102.1 | 03/19/2020 | Australia:Victoria |
| MT451103.1 | 03/19/2020 | Australia:Victoria |
| MT451104.1 | 03/19/2020 | Australia:Victoria |
| MT451105.1 | 03/19/2020 | Australia:Victoria |
| MT451106.1 | 03/19/2020 | Australia:Victoria |
| MT451107.1 | 03/19/2020 | Australia:Victoria |
| MT451108.1 | 03/19/2020 | Australia:Victoria |
| MT451225.1 | 03/19/2020 | Australia:Victoria |
| MT459862.1 | 03/19/2020 | Greece:Athens |
| MT459868.1 | 03/19/2020 | Greece:Athens |

| | | |
|---|---|---|
| MT459894.1 | 03/19/2020 | Greece:Athens |
| MT477839.1 | 03/19/2020 | USA:AK |
| MT499207.1 | 03/19/2020 | USA:CA |
| MT506541.1 | 03/19/2020 | USA:Michigan |
| MT506653.1 | 03/19/2020 | USA:Michigan |
| MT506654.1 | 03/19/2020 | USA:Michigan |
| MT506655.1 | 03/19/2020 | USA:Michigan |
| MT506656.1 | 03/19/2020 | USA:Michigan |
| MT506658.1 | 03/19/2020 | USA:Michigan |
| MT506659.1 | 03/19/2020 | USA:Michigan |
| MT506660.1 | 03/19/2020 | USA:Michigan |
| MT506680.1 | 03/19/2020 | USA:Michigan |
| MT509457.1 | 03/19/2020 | USA |
| MT509460.1 | 03/19/2020 | USA |
| MT509677.1 | 03/19/2020 | USA:Michigan |
| MT509678.1 | 03/19/2020 | USA:Michigan |
| MT509679.1 | 03/19/2020 | USA:Michigan |
| MT509680.1 | 03/19/2020 | USA:Michigan |
| MT510722.1 | 03/19/2020 | USA |
| MT510723.1 | 03/19/2020 | USA |
| MT520257.1 | 03/19/2020 | USA:Massachusetts |
| MT520291.1 | 03/19/2020 | USA:Massachusetts |
| MT520350.1 | 03/19/2020 | USA:Massachusetts |
| MT520511.1 | 03/19/2020 | USA:Massachusetts |
| MT520527.1 | 03/19/2020 | USA:Massachusetts |
| MT520535.1 | 03/19/2020 | USA:Massachusetts |
| MT533204.1 | 03/19/2020 | USA:CA |
| MT533205.1 | 03/19/2020 | USA:CA |
| MT582456.1 | 03/19/2020 | Germany:Dusseldorf |
| MT582457.1 | 03/19/2020 | Germany:Dusseldorf |
| MT594109.1 | 03/19/2020 | USA |
| MT606516.1 | 03/19/2020 | USA:Maricopa-Country-Arizona |
| MT628127.1 | 03/19/2020 | USA:CA |
| MT628172.1 | 03/19/2020 | USA:CA |
| MT632496.1 | 03/19/2020 | USA:Washington-King-County |
| MT632515.1 | 03/19/2020 | USA:Washington-King-County |
| MT632538.1 | 03/19/2020 | USA:Washington-Grant-County |
| MT632539.1 | 03/19/2020 | USA:Washington-King-County |
| MT632550.1 | 03/19/2020 | USA:Washington |
| MT632551.1 | 03/19/2020 | USA:Washington-King-County |
| MT632554.1 | 03/19/2020 | USA:Washington |
| MT632556.1 | 03/19/2020 | USA:Washington-King-County |
| MT632557.1 | 03/19/2020 | USA:Washington-King-County |
| MT632561.1 | 03/19/2020 | USA:Washington-King-County |
| MT632572.1 | 03/19/2020 | USA:Washington-King-County |
| MT632573.1 | 03/19/2020 | USA:Washington-King-County |

| | | |
|---|---|---|
| MT632574.1 | 03/19/2020 | USA:Washington-King-County |
| MT632578.1 | 03/19/2020 | USA:Washington-King-County |
| MT632584.1 | 03/19/2020 | USA:Washington-King-County |
| MT632594.1 | 03/19/2020 | USA:Washington-King-County |
| MT632620.1 | 03/19/2020 | USA:Washington |
| MT635211.1 | 03/19/2020 | USA:Connecticut |
| MT635212.1 | 03/19/2020 | USA:Connecticut |
| MT646068.1 | 03/19/2020 | USA |
| MT646072.1 | 03/19/2020 | USA |
| MT646083.1 | 03/19/2020 | USA |
| LC547522.1 | 03/20/2020 | Japan:Ishikawa |
| LC547528.1 | 03/20/2020 | Japan:Saitama |
| LC547529.1 | 03/20/2020 | Japan:Saitama |
| MT263420.1 | 03/20/2020 | USA:WA |
| MT263444.1 | 03/20/2020 | USA:WA |
| MT326041.1 | 03/20/2020 | USA |
| MT326074.1 | 03/20/2020 | USA |
| MT326075.1 | 03/20/2020 | USA |
| MT326079.1 | 03/20/2020 | USA |
| MT326085.1 | 03/20/2020 | USA |
| MT326112.1 | 03/20/2020 | USA |
| MT326113.1 | 03/20/2020 | USA |
| MT326116.1 | 03/20/2020 | USA |
| MT326119.1 | 03/20/2020 | USA |
| MT326121.1 | 03/20/2020 | USA |
| MT326122.1 | 03/20/2020 | USA |
| MT326123.1 | 03/20/2020 | USA |
| MT326124.1 | 03/20/2020 | USA |
| MT326125.1 | 03/20/2020 | USA |
| MT326126.1 | 03/20/2020 | USA |
| MT326127.1 | 03/20/2020 | USA |
| MT326128.1 | 03/20/2020 | USA |
| MT326129.1 | 03/20/2020 | USA |
| MT326130.1 | 03/20/2020 | USA |
| MT326132.1 | 03/20/2020 | USA |
| MT326133.1 | 03/20/2020 | USA |
| MT326134.1 | 03/20/2020 | USA |
| MT326135.1 | 03/20/2020 | USA |
| MT326138.1 | 03/20/2020 | USA |
| MT326139.1 | 03/20/2020 | USA |
| MT326140.1 | 03/20/2020 | USA |
| MT326143.1 | 03/20/2020 | USA |
| MT326144.1 | 03/20/2020 | USA |
| MT326147.1 | 03/20/2020 | USA |
| MT334548.1 | 03/20/2020 | USA:UT |
| MT334555.1 | 03/20/2020 | USA:UT |

| | | |
|---|---|---|
| MT334556.1 | 03/20/2020 | USA:UT |
| MT334557.1 | 03/20/2020 | USA:UT |
| MT350269.1 | 03/20/2020 | USA:WA |
| MT350271.1 | 03/20/2020 | USA:WA |
| MT370921.1 | 03/20/2020 | USA:NY |
| MT370925.1 | 03/20/2020 | USA:NY |
| MT370929.1 | 03/20/2020 | USA:NY |
| MT370932.1 | 03/20/2020 | USA:NY |
| MT370933.1 | 03/20/2020 | USA:NJ |
| MT370934.1 | 03/20/2020 | USA:NY |
| MT370935.1 | 03/20/2020 | USA:NY |
| MT370936.1 | 03/20/2020 | USA:NY |
| MT370937.1 | 03/20/2020 | USA:NY |
| MT370941.1 | 03/20/2020 | USA:NY |
| MT370946.1 | 03/20/2020 | USA:NY |
| MT370948.1 | 03/20/2020 | USA:NY |
| MT370949.1 | 03/20/2020 | USA:NY |
| MT370950.1 | 03/20/2020 | USA:NY |
| MT370952.1 | 03/20/2020 | USA:NY |
| MT370961.1 | 03/20/2020 | USA:NY |
| MT370968.1 | 03/20/2020 | USA:NY |
| MT370970.1 | 03/20/2020 | USA:NY |
| MT370971.1 | 03/20/2020 | USA:NY |
| MT370972.1 | 03/20/2020 | USA:NY |
| MT370973.1 | 03/20/2020 | USA:NY |
| MT370976.1 | 03/20/2020 | USA:NY |
| MT370980.1 | 03/20/2020 | USA:NY |
| MT370985.1 | 03/20/2020 | USA:NY |
| MT370986.1 | 03/20/2020 | USA:NY |
| MT370988.1 | 03/20/2020 | USA:NY |
| MT370994.1 | 03/20/2020 | USA:NY |
| MT370995.1 | 03/20/2020 | USA:NY |
| MT370999.1 | 03/20/2020 | USA:NY |
| MT371003.1 | 03/20/2020 | USA:NY |
| MT371004.1 | 03/20/2020 | USA:NY |
| MT371006.1 | 03/20/2020 | USA:NY |
| MT371015.1 | 03/20/2020 | USA:NY |
| MT371018.1 | 03/20/2020 | USA:NY |
| MT371021.1 | 03/20/2020 | USA:NY |
| MT371027.1 | 03/20/2020 | USA:NY |
| MT371029.1 | 03/20/2020 | USA:NY |
| MT371031.1 | 03/20/2020 | USA:NY |
| MT371036.1 | 03/20/2020 | USA:NY |
| MT372481.1 | 03/20/2020 | Malaysia |
| MT372482.1 | 03/20/2020 | Malaysia |
| MT380731.1 | 03/20/2020 | USA |

| | | |
|---|---|---|
| MT412171.1 | 03/20/2020 | USA:Michigan |
| MT412172.1 | 03/20/2020 | USA:Michigan |
| MT412173.1 | 03/20/2020 | USA:Michigan |
| MT412174.1 | 03/20/2020 | USA:Michigan |
| MT412177.1 | 03/20/2020 | USA:Michigan |
| MT412178.1 | 03/20/2020 | USA:Michigan |
| MT412188.1 | 03/20/2020 | USA:Michigan |
| MT412189.1 | 03/20/2020 | USA:Michigan |
| MT415323.1 | 03/20/2020 | India |
| MT419860.1 | 03/20/2020 | USA:CA |
| MT428552.1 | 03/20/2020 | Kazakhstan |
| MT428553.1 | 03/20/2020 | Kazakhstan |
| MT428554.1 | 03/20/2020 | Kazakhstan |
| MT439257.1 | 03/20/2020 | USA:MI |
| MT439263.1 | 03/20/2020 | USA:MI |
| MT447176.1 | 03/20/2020 | Thailand |
| MT451023.1 | 03/20/2020 | Australia:Victoria |
| MT451028.1 | 03/20/2020 | Australia:Victoria |
| MT451032.1 | 03/20/2020 | Australia:Victoria |
| MT451033.1 | 03/20/2020 | Australia:Victoria |
| MT451034.1 | 03/20/2020 | Australia:Victoria |
| MT451035.1 | 03/20/2020 | Australia:Victoria |
| MT451109.1 | 03/20/2020 | Australia:Victoria |
| MT451110.1 | 03/20/2020 | Australia:Victoria |
| MT451111.1 | 03/20/2020 | Australia:Victoria |
| MT451112.1 | 03/20/2020 | Australia:Victoria |
| MT451113.1 | 03/20/2020 | Australia:Victoria |
| MT451114.1 | 03/20/2020 | Australia:Victoria |
| MT451115.1 | 03/20/2020 | Australia:Victoria |
| MT451116.1 | 03/20/2020 | Australia:Victoria |
| MT451117.1 | 03/20/2020 | Australia:Victoria |
| MT451118.1 | 03/20/2020 | Australia:Victoria |
| MT451119.1 | 03/20/2020 | Australia:Victoria |
| MT451120.1 | 03/20/2020 | Australia:Victoria |
| MT451121.1 | 03/20/2020 | Australia:Victoria |
| MT451122.1 | 03/20/2020 | Australia:Victoria |
| MT451123.1 | 03/20/2020 | Australia:Victoria |
| MT451124.1 | 03/20/2020 | Australia:Victoria |
| MT451125.1 | 03/20/2020 | Australia:Victoria |
| MT451126.1 | 03/20/2020 | Australia:Victoria |
| MT451127.1 | 03/20/2020 | Australia:Victoria |
| MT451128.1 | 03/20/2020 | Australia:Victoria |
| MT451226.1 | 03/20/2020 | Australia:Victoria |
| MT451227.1 | 03/20/2020 | Australia:Victoria |
| MT451228.1 | 03/20/2020 | Australia:Victoria |
| MT451229.1 | 03/20/2020 | Australia:Victoria |

| | | |
|---|---|---|
| MT451230.1 | 03/20/2020 | Australia:Victoria |
| MT451231.1 | 03/20/2020 | Australia:Victoria |
| MT451232.1 | 03/20/2020 | Australia:Victoria |
| MT451233.1 | 03/20/2020 | Australia:Victoria |
| MT451234.1 | 03/20/2020 | Australia:Victoria |
| MT451235.1 | 03/20/2020 | Australia:Victoria |
| MT451236.1 | 03/20/2020 | Australia:Victoria |
| MT451237.1 | 03/20/2020 | Australia:Victoria |
| MT451238.1 | 03/20/2020 | Australia:Victoria |
| MT451239.1 | 03/20/2020 | Australia:Victoria |
| MT451241.1 | 03/20/2020 | Australia:Victoria |
| MT451242.1 | 03/20/2020 | Australia:Victoria |
| MT451243.1 | 03/20/2020 | Australia:Victoria |
| MT451534.1 | 03/20/2020 | Australia:Victoria |
| MT451535.1 | 03/20/2020 | Australia:Victoria |
| MT451536.1 | 03/20/2020 | Australia:Victoria |
| MT451539.1 | 03/20/2020 | Australia:Victoria |
| MT451540.1 | 03/20/2020 | Australia:Victoria |
| MT459833.1 | 03/20/2020 | Greece:Athens |
| MT459839.1 | 03/20/2020 | Greece:Athens |
| MT459867.1 | 03/20/2020 | Greece:Athens |
| MT459987.1 | 03/20/2020 | Guam |
| MT479226.1 | 03/20/2020 | Taiwan |
| MT479227.1 | 03/20/2020 | Taiwan |
| MT499194.1 | 03/20/2020 | USA:CA |
| MT499195.1 | 03/20/2020 | USA:CA |
| MT499200.1 | 03/20/2020 | USA:CA |
| MT499201.1 | 03/20/2020 | USA:CA |
| MT499203.1 | 03/20/2020 | USA:CA |
| MT499205.1 | 03/20/2020 | USA:CA |
| MT506539.1 | 03/20/2020 | USA:Michigan |
| MT506540.1 | 03/20/2020 | USA:Michigan |
| MT506542.1 | 03/20/2020 | USA:Michigan |
| MT506673.1 | 03/20/2020 | USA:Michigan |
| MT509455.1 | 03/20/2020 | USA |
| MT509456.1 | 03/20/2020 | USA |
| MT509473.1 | 03/20/2020 | USA |
| MT509477.1 | 03/20/2020 | USA |
| MT509681.1 | 03/20/2020 | USA:Michigan |
| MT510643.1 | 03/20/2020 | Russia:Moscow-region |
| MT510725.1 | 03/20/2020 | USA |
| MT510726.1 | 03/20/2020 | USA |
| MT520213.1 | 03/20/2020 | USA:Massachusetts |
| MT520242.1 | 03/20/2020 | USA:Massachusetts |
| MT520413.1 | 03/20/2020 | USA:Massachusetts |
| MT520424.1 | 03/20/2020 | USA:Massachusetts |

| | | |
|---|---|---|
| MT520438.1 | 03/20/2020 | USA:Massachusetts |
| MT520474.1 | 03/20/2020 | USA:Massachusetts |
| MT520481.1 | 03/20/2020 | USA:Massachusetts |
| MT520531.1 | 03/20/2020 | USA:Massachusetts |
| MT534285.1 | 03/20/2020 | Czech-Republic |
| MT582446.1 | 03/20/2020 | Germany:Dusseldorf |
| MT582447.1 | 03/20/2020 | Germany:Dusseldorf |
| MT582453.1 | 03/20/2020 | Germany:Dusseldorf |
| MT582455.1 | 03/20/2020 | Germany:Dusseldorf |
| MT627931.1 | 03/20/2020 | USA:Washington-King-County |
| MT628130.1 | 03/20/2020 | USA:CA |
| MT628131.1 | 03/20/2020 | USA:CA |
| MT628132.1 | 03/20/2020 | USA:CA |
| MT628133.1 | 03/20/2020 | USA:CA |
| MT632493.1 | 03/20/2020 | USA:Washington-Whatcom-County |
| MT632498.1 | 03/20/2020 | USA:Washington-Whatcom-County |
| MT632499.1 | 03/20/2020 | USA:Washington-Whatcom-County |
| MT632503.1 | 03/20/2020 | USA:Washington-Whatcom-County |
| MT632505.1 | 03/20/2020 | USA:Washington-Whatcom-County |
| MT632507.1 | 03/20/2020 | USA:Washington-Whatcom-County |
| MT632508.1 | 03/20/2020 | USA:Washington-Whatcom-County |
| MT632509.1 | 03/20/2020 | USA:Washington |
| MT632511.1 | 03/20/2020 | USA:Washington-Whatcom-County |
| MT632512.1 | 03/20/2020 | USA:Washington-Whatcom-County |
| MT632513.1 | 03/20/2020 | USA:Washington-Whatcom-County |
| MT632514.1 | 03/20/2020 | USA:Washington-King-County |
| MT632516.1 | 03/20/2020 | USA:Washington-Whatcom-County |
| MT632517.1 | 03/20/2020 | USA:Washington-Whatcom-County |
| MT632519.1 | 03/20/2020 | USA:Washington-Whatcom-County |
| MT632520.1 | 03/20/2020 | USA:Washington-Whatcom-County |
| MT632521.1 | 03/20/2020 | USA:Washington-Whatcom-County |
| MT632523.1 | 03/20/2020 | USA:Washington-Whatcom-County |
| MT632525.1 | 03/20/2020 | USA:Washington-Whatcom-County |
| MT632526.1 | 03/20/2020 | USA:Washington |
| MT632527.1 | 03/20/2020 | USA:Washington-Whatcom-County |
| MT632529.1 | 03/20/2020 | USA:Washington-Whatcom-County |
| MT632534.1 | 03/20/2020 | USA:Washington |
| MT632540.1 | 03/20/2020 | USA:Washington-Whatcom-County |
| MT632546.1 | 03/20/2020 | USA:Washington-King-County |
| MT632553.1 | 03/20/2020 | USA:Washington-Whatcom-County |
| MT632563.1 | 03/20/2020 | USA:Washington-King-County |
| MT632577.1 | 03/20/2020 | USA:Washington-King-County |
| MT632583.1 | 03/20/2020 | USA:Washington-Yakima-County |
| MT632588.1 | 03/20/2020 | USA:Washington-King-County |
| MT632590.1 | 03/20/2020 | USA:Washington-Whatcom-County |
| MT632599.1 | 03/20/2020 | USA:Washington |

| | | |
|---|---|---|
| MT632601.1 | 03/20/2020 | USA:Washington-Pierce-County |
| MT635444.1 | 03/20/2020 | Russia:Moscow-region |
| MT641651.1 | 03/20/2020 | Australia:Victoria |
| MT641654.1 | 03/20/2020 | Australia:Northern-Territory |
| MT646048.1 | 03/20/2020 | USA |
| MT646070.1 | 03/20/2020 | USA |
| MT646086.1 | 03/20/2020 | USA |
| MT646100.1 | 03/20/2020 | USA |
| MT326040.1 | 03/21/2020 | USA |
| MT326055.1 | 03/21/2020 | USA |
| MT326089.1 | 03/21/2020 | USA |
| MT326090.1 | 03/21/2020 | USA |
| MT326097.1 | 03/21/2020 | USA |
| MT326118.1 | 03/21/2020 | USA |
| MT326120.1 | 03/21/2020 | USA |
| MT326148.1 | 03/21/2020 | USA |
| MT326149.1 | 03/21/2020 | USA |
| MT326150.1 | 03/21/2020 | USA |
| MT326151.1 | 03/21/2020 | USA |
| MT326152.1 | 03/21/2020 | USA |
| MT326153.1 | 03/21/2020 | USA |
| MT326154.1 | 03/21/2020 | USA |
| MT326155.1 | 03/21/2020 | USA |
| MT326156.1 | 03/21/2020 | USA |
| MT326158.1 | 03/21/2020 | USA |
| MT326159.1 | 03/21/2020 | USA |
| MT326160.1 | 03/21/2020 | USA |
| MT326161.1 | 03/21/2020 | USA |
| MT326162.1 | 03/21/2020 | USA |
| MT326163.1 | 03/21/2020 | USA |
| MT326164.1 | 03/21/2020 | USA |
| MT326165.1 | 03/21/2020 | USA |
| MT326166.1 | 03/21/2020 | USA |
| MT326167.1 | 03/21/2020 | USA |
| MT326168.1 | 03/21/2020 | USA |
| MT326170.1 | 03/21/2020 | USA |
| MT326171.1 | 03/21/2020 | USA |
| MT326172.1 | 03/21/2020 | USA |
| MT345802.1 | 03/21/2020 | USA |
| MT345803.1 | 03/21/2020 | USA:ID |
| MT370923.1 | 03/21/2020 | USA:NY |
| MT370927.1 | 03/21/2020 | USA:NY |
| MT370931.1 | 03/21/2020 | USA:NY |
| MT370938.1 | 03/21/2020 | USA:NY |
| MT370943.1 | 03/21/2020 | USA:NY |
| MT370947.1 | 03/21/2020 | USA:NY |

| | | |
|---|---|---|
| MT370954.1 | 03/21/2020 | USA:NY |
| MT370959.1 | 03/21/2020 | USA:NY |
| MT370962.1 | 03/21/2020 | USA:NY |
| MT370964.1 | 03/21/2020 | USA:NY |
| MT370967.1 | 03/21/2020 | USA:NY |
| MT370969.1 | 03/21/2020 | USA:NY |
| MT370977.1 | 03/21/2020 | USA:NY |
| MT370979.1 | 03/21/2020 | USA:NY |
| MT370983.1 | 03/21/2020 | USA:NY |
| MT370984.1 | 03/21/2020 | USA:NY |
| MT370990.1 | 03/21/2020 | USA:NY |
| MT370991.1 | 03/21/2020 | USA:NY |
| MT371000.1 | 03/21/2020 | USA:NY |
| MT371009.1 | 03/21/2020 | USA:NY |
| MT371013.1 | 03/21/2020 | USA:NY |
| MT371014.1 | 03/21/2020 | USA:NY |
| MT371019.1 | 03/21/2020 | USA:NY |
| MT371023.1 | 03/21/2020 | USA:NY |
| MT371024.1 | 03/21/2020 | USA:NY |
| MT371028.1 | 03/21/2020 | USA:NY |
| MT371033.1 | 03/21/2020 | USA:NY |
| MT371035.1 | 03/21/2020 | USA:NY |
| MT371570.1 | 03/21/2020 | Czech-Republic |
| MT412179.1 | 03/21/2020 | USA:Michigan |
| MT412180.1 | 03/21/2020 | USA:Michigan |
| MT412181.1 | 03/21/2020 | USA:Michigan |
| MT412182.1 | 03/21/2020 | USA:Michigan |
| MT412183.1 | 03/21/2020 | USA:Michigan |
| MT412184.1 | 03/21/2020 | USA:Michigan |
| MT412185.1 | 03/21/2020 | USA:Michigan |
| MT412196.1 | 03/21/2020 | USA:Michigan |
| MT439260.1 | 03/21/2020 | USA:MI |
| MT451036.1 | 03/21/2020 | Australia:Victoria |
| MT451037.1 | 03/21/2020 | Australia:Victoria |
| MT451129.1 | 03/21/2020 | Australia:Victoria |
| MT451130.1 | 03/21/2020 | Australia:Victoria |
| MT451131.1 | 03/21/2020 | Australia:Victoria |
| MT451132.1 | 03/21/2020 | Australia:Victoria |
| MT451133.1 | 03/21/2020 | Australia:Victoria |
| MT451134.1 | 03/21/2020 | Australia:Victoria |
| MT451135.1 | 03/21/2020 | Australia:Victoria |
| MT451136.1 | 03/21/2020 | Australia:Victoria |
| MT451137.1 | 03/21/2020 | Australia:Victoria |
| MT451138.1 | 03/21/2020 | Australia:Victoria |
| MT451139.1 | 03/21/2020 | Australia:Victoria |
| MT451140.1 | 03/21/2020 | Australia:Victoria |

| | | |
|---|---|---|
| MT451141.1 | 03/21/2020 | Australia:Victoria |
| MT451142.1 | 03/21/2020 | Australia:Victoria |
| MT451143.1 | 03/21/2020 | Australia:Victoria |
| MT451144.1 | 03/21/2020 | Australia:Victoria |
| MT451145.1 | 03/21/2020 | Australia:Victoria |
| MT451146.1 | 03/21/2020 | Australia:Victoria |
| MT451147.1 | 03/21/2020 | Australia:Victoria |
| MT451148.1 | 03/21/2020 | Australia:Victoria |
| MT451149.1 | 03/21/2020 | Australia:Victoria |
| MT451150.1 | 03/21/2020 | Australia:Victoria |
| MT451151.1 | 03/21/2020 | Australia:Victoria |
| MT451152.1 | 03/21/2020 | Australia:Victoria |
| MT451153.1 | 03/21/2020 | Australia:Victoria |
| MT451154.1 | 03/21/2020 | Australia:Victoria |
| MT451155.1 | 03/21/2020 | Australia:Victoria |
| MT451156.1 | 03/21/2020 | Australia:Victoria |
| MT451157.1 | 03/21/2020 | Australia:Victoria |
| MT451158.1 | 03/21/2020 | Australia:Victoria |
| MT451159.1 | 03/21/2020 | Australia:Victoria |
| MT451244.1 | 03/21/2020 | Australia:Victoria |
| MT451245.1 | 03/21/2020 | Australia:Victoria |
| MT451246.1 | 03/21/2020 | Australia:Victoria |
| MT451247.1 | 03/21/2020 | Australia:Victoria |
| MT451248.1 | 03/21/2020 | Australia:Victoria |
| MT451249.1 | 03/21/2020 | Australia:Victoria |
| MT451490.1 | 03/21/2020 | Australia:Victoria |
| MT477843.1 | 03/21/2020 | USA:AK |
| MT479225.1 | 03/21/2020 | Taiwan |
| MT499197.1 | 03/21/2020 | USA:CA |
| MT499199.1 | 03/21/2020 | USA:CA |
| MT506543.1 | 03/21/2020 | USA:Michigan |
| MT506664.1 | 03/21/2020 | USA:Michigan |
| MT506669.1 | 03/21/2020 | USA:Michigan |
| MT506670.1 | 03/21/2020 | USA:Michigan |
| MT506671.1 | 03/21/2020 | USA:Michigan |
| MT506672.1 | 03/21/2020 | USA:Michigan |
| MT506675.1 | 03/21/2020 | USA:Michigan |
| MT506676.1 | 03/21/2020 | USA:Michigan |
| MT506677.1 | 03/21/2020 | USA:Michigan |
| MT506678.1 | 03/21/2020 | USA:Michigan |
| MT506679.1 | 03/21/2020 | USA:Michigan |
| MT509452.1 | 03/21/2020 | USA |
| MT509468.1 | 03/21/2020 | USA |
| MT509470.1 | 03/21/2020 | USA |
| MT512420.1 | 03/21/2020 | USA |
| MT520221.1 | 03/21/2020 | USA:Massachusetts |

| | | |
|---|---|---|
| MT520463.1 | 03/21/2020 | USA:Massachusetts |
| MT520499.1 | 03/21/2020 | USA:Massachusetts |
| MT559037.1 | 03/21/2020 | Tunisia:Bizerte |
| MT582454.1 | 03/21/2020 | Germany:Dusseldorf |
| MT598171.1 | 03/21/2020 | USA:San-Diego-California |
| MT627607.1 | 03/21/2020 | USA:Washington-King-County |
| MT628134.1 | 03/21/2020 | USA:CA |
| MT628234.1 | 03/21/2020 | USA:CA |
| MT628235.1 | 03/21/2020 | USA:CA |
| MT628236.1 | 03/21/2020 | USA:CA |
| MT632497.1 | 03/21/2020 | USA:Washington-Whatcom-County |
| MT635443.1 | 03/21/2020 | Russia:Moscow-region |
| MT646105.1 | 03/21/2020 | USA |
| MT646120.1 | 03/21/2020 | USA |
| MT263461.1 | 03/22/2020 | USA:WA |
| MT326043.1 | 03/22/2020 | USA |
| MT326072.1 | 03/22/2020 | USA |
| MT326083.1 | 03/22/2020 | USA |
| MT326084.1 | 03/22/2020 | USA |
| MT326092.1 | 03/22/2020 | USA |
| MT326093.1 | 03/22/2020 | USA |
| MT326094.1 | 03/22/2020 | USA |
| MT326095.1 | 03/22/2020 | USA |
| MT326096.1 | 03/22/2020 | USA |
| MT326098.1 | 03/22/2020 | USA |
| MT326099.1 | 03/22/2020 | USA |
| MT326100.1 | 03/22/2020 | USA |
| MT326101.1 | 03/22/2020 | USA |
| MT326102.1 | 03/22/2020 | USA |
| MT326103.1 | 03/22/2020 | USA |
| MT326104.1 | 03/22/2020 | USA |
| MT326105.1 | 03/22/2020 | USA |
| MT326106.1 | 03/22/2020 | USA |
| MT326107.1 | 03/22/2020 | USA |
| MT326109.1 | 03/22/2020 | USA |
| MT326110.1 | 03/22/2020 | USA |
| MT326111.1 | 03/22/2020 | USA |
| MT326169.1 | 03/22/2020 | USA |
| MT370922.1 | 03/22/2020 | USA:NY |
| MT370924.1 | 03/22/2020 | USA:NY |
| MT370930.1 | 03/22/2020 | USA:NY |
| MT370945.1 | 03/22/2020 | USA:NY |
| MT370974.1 | 03/22/2020 | USA:NY |
| MT370978.1 | 03/22/2020 | USA:NY |
| MT371002.1 | 03/22/2020 | USA:NY |
| MT371008.1 | 03/22/2020 | USA:NY |

| | | |
|---|---|---|
| MT371011.1 | 03/22/2020 | USA:NY |
| MT371016.1 | 03/22/2020 | USA:NY |
| MT371017.1 | 03/22/2020 | USA:NY |
| MT371022.1 | 03/22/2020 | USA:NY |
| MT371025.1 | 03/22/2020 | USA:NY |
| MT371034.1 | 03/22/2020 | USA:NY |
| MT371568.1 | 03/22/2020 | Czech-Republic |
| MT380732.1 | 03/22/2020 | USA |
| MT412186.1 | 03/22/2020 | USA:Michigan |
| MT412187.1 | 03/22/2020 | USA:Michigan |
| MT412192.1 | 03/22/2020 | USA:Michigan |
| MT412193.1 | 03/22/2020 | USA:Michigan |
| MT439259.1 | 03/22/2020 | USA:MI |
| MT439264.1 | 03/22/2020 | USA:MI |
| MT439268.1 | 03/22/2020 | USA:MI |
| MT439271.1 | 03/22/2020 | USA:MI |
| MT439282.1 | 03/22/2020 | USA:MI |
| MT451038.1 | 03/22/2020 | Australia:Victoria |
| MT451039.1 | 03/22/2020 | Australia:Victoria |
| MT451040.1 | 03/22/2020 | Australia:Victoria |
| MT451041.1 | 03/22/2020 | Australia:Victoria |
| MT451160.1 | 03/22/2020 | Australia:Victoria |
| MT451161.1 | 03/22/2020 | Australia:Victoria |
| MT451162.1 | 03/22/2020 | Australia:Victoria |
| MT451163.1 | 03/22/2020 | Australia:Victoria |
| MT451164.1 | 03/22/2020 | Australia:Victoria |
| MT451165.1 | 03/22/2020 | Australia:Victoria |
| MT451166.1 | 03/22/2020 | Australia:Victoria |
| MT451167.1 | 03/22/2020 | Australia:Victoria |
| MT451168.1 | 03/22/2020 | Australia:Victoria |
| MT451169.1 | 03/22/2020 | Australia:Victoria |
| MT451170.1 | 03/22/2020 | Australia:Victoria |
| MT451171.1 | 03/22/2020 | Australia:Victoria |
| MT451172.1 | 03/22/2020 | Australia:Victoria |
| MT451173.1 | 03/22/2020 | Australia:Victoria |
| MT451174.1 | 03/22/2020 | Australia:Victoria |
| MT451175.1 | 03/22/2020 | Australia:Victoria |
| MT451176.1 | 03/22/2020 | Australia:Victoria |
| MT451177.1 | 03/22/2020 | Australia:Victoria |
| MT451178.1 | 03/22/2020 | Australia:Victoria |
| MT451179.1 | 03/22/2020 | Australia:Victoria |
| MT451180.1 | 03/22/2020 | Australia:Victoria |
| MT451181.1 | 03/22/2020 | Australia:Victoria |
| MT451182.1 | 03/22/2020 | Australia:Victoria |
| MT451183.1 | 03/22/2020 | Australia:Victoria |
| MT451250.1 | 03/22/2020 | Australia:Victoria |

| | | |
|---|---|---|
| MT451251.1 | 03/22/2020 | Australia:Victoria |
| MT451252.1 | 03/22/2020 | Australia:Victoria |
| MT451253.1 | 03/22/2020 | Australia:Victoria |
| MT451254.1 | 03/22/2020 | Australia:Victoria |
| MT451255.1 | 03/22/2020 | Australia:Victoria |
| MT451256.1 | 03/22/2020 | Australia:Victoria |
| MT451257.1 | 03/22/2020 | Australia:Victoria |
| MT451258.1 | 03/22/2020 | Australia:Victoria |
| MT451259.1 | 03/22/2020 | Australia:Victoria |
| MT451260.1 | 03/22/2020 | Australia:Victoria |
| MT451261.1 | 03/22/2020 | Australia:Victoria |
| MT451262.1 | 03/22/2020 | Australia:Victoria |
| MT451263.1 | 03/22/2020 | Australia:Victoria |
| MT451264.1 | 03/22/2020 | Australia:Victoria |
| MT451265.1 | 03/22/2020 | Australia:Victoria |
| MT451266.1 | 03/22/2020 | Australia:Victoria |
| MT451351.1 | 03/22/2020 | Australia:Victoria |
| MT451404.1 | 03/22/2020 | Australia:Victoria |
| MT451405.1 | 03/22/2020 | Australia:Victoria |
| MT451406.1 | 03/22/2020 | Australia:Victoria |
| MT459845.1 | 03/22/2020 | Greece:Athens |
| MT459895.1 | 03/22/2020 | Greece:Athens |
| MT499196.1 | 03/22/2020 | USA:CA |
| MT506661.1 | 03/22/2020 | USA:Michigan |
| MT506662.1 | 03/22/2020 | USA:Michigan |
| MT506663.1 | 03/22/2020 | USA:Michigan |
| MT506665.1 | 03/22/2020 | USA:Michigan |
| MT506666.1 | 03/22/2020 | USA:Michigan |
| MT506667.1 | 03/22/2020 | USA:Michigan |
| MT506668.1 | 03/22/2020 | USA:Michigan |
| MT506674.1 | 03/22/2020 | USA:Michigan |
| MT506682.1 | 03/22/2020 | USA:Michigan |
| MT509461.1 | 03/22/2020 | USA |
| MT509467.1 | 03/22/2020 | USA |
| MT510747.1 | 03/22/2020 | Russia:Moscow-region |
| MT520172.1 | 03/22/2020 | USA:Massachusetts |
| MT520193.1 | 03/22/2020 | USA:Massachusetts |
| MT520200.1 | 03/22/2020 | USA:Massachusetts |
| MT520215.1 | 03/22/2020 | USA:Massachusetts |
| MT520241.1 | 03/22/2020 | USA:Massachusetts |
| MT520255.1 | 03/22/2020 | USA:Massachusetts |
| MT520261.1 | 03/22/2020 | USA:Massachusetts |
| MT520306.1 | 03/22/2020 | USA:Massachusetts |
| MT520307.1 | 03/22/2020 | USA:Massachusetts |
| MT520342.1 | 03/22/2020 | USA:Massachusetts |
| MT520360.1 | 03/22/2020 | USA:Massachusetts |

| | | |
|---|---|---|
| MT520391.1 | 03/22/2020 | USA:Massachusetts |
| MT520455.1 | 03/22/2020 | USA:Massachusetts |
| MT520462.1 | 03/22/2020 | USA:Massachusetts |
| MT520498.1 | 03/22/2020 | USA:Massachusetts |
| MT520540.1 | 03/22/2020 | USA:Massachusetts |
| MT628238.1 | 03/22/2020 | USA:CA |
| MT632528.1 | 03/22/2020 | USA:Washington-Whatcom-County |
| MT635445.1 | 03/22/2020 | Russia:Moscow-region |
| MT637143.1 | 03/22/2020 | Russia:Moscow-region |
| MT641664.1 | 03/22/2020 | Australia:Victoria |
| MT641665.1 | 03/22/2020 | Australia:Victoria |
| MT641669.1 | 03/22/2020 | Australia:Victoria |
| MT641678.1 | 03/22/2020 | Australia:Victoria |
| MT641682.1 | 03/22/2020 | Australia:Victoria |
| MT646113.1 | 03/22/2020 | USA |
| LC546038.1 | 03/23/2020 | Japan |
| MT263381.1 | 03/23/2020 | USA:WA |
| MT263382.1 | 03/23/2020 | USA:WA |
| MT263384.1 | 03/23/2020 | USA:WA |
| MT263385.1 | 03/23/2020 | USA:WA |
| MT263400.1 | 03/23/2020 | USA:WA |
| MT263402.1 | 03/23/2020 | USA:WA |
| MT263423.1 | 03/23/2020 | USA:WA |
| MT263462.1 | 03/23/2020 | USA:WA |
| MT326032.1 | 03/23/2020 | USA |
| MT326033.1 | 03/23/2020 | USA |
| MT326034.1 | 03/23/2020 | USA |
| MT326035.1 | 03/23/2020 | USA |
| MT326036.1 | 03/23/2020 | USA |
| MT326039.1 | 03/23/2020 | USA |
| MT326042.1 | 03/23/2020 | USA |
| MT326044.1 | 03/23/2020 | USA |
| MT326046.1 | 03/23/2020 | USA |
| MT326047.1 | 03/23/2020 | USA |
| MT326048.1 | 03/23/2020 | USA |
| MT326049.1 | 03/23/2020 | USA |
| MT326051.1 | 03/23/2020 | USA |
| MT326052.1 | 03/23/2020 | USA |
| MT326053.1 | 03/23/2020 | USA |
| MT326054.1 | 03/23/2020 | USA |
| MT326061.1 | 03/23/2020 | USA |
| MT326064.1 | 03/23/2020 | USA |
| MT326082.1 | 03/23/2020 | USA |
| MT326087.1 | 03/23/2020 | USA |
| MT326088.1 | 03/23/2020 | USA |
| MT334552.1 | 03/23/2020 | USA:UT |

| | | |
|---|---|---|
| MT345799.1 | 03/23/2020 | USA:WA |
| MT345805.1 | 03/23/2020 | USA:WA |
| MT345816.1 | 03/23/2020 | USA |
| MT345828.1 | 03/23/2020 | USA:ID |
| MT345831.1 | 03/23/2020 | USA:ID |
| MT345836.1 | 03/23/2020 | USA:WA |
| MT345838.1 | 03/23/2020 | USA:WA |
| MT345858.1 | 03/23/2020 | USA:ID |
| MT345866.1 | 03/23/2020 | USA:ID |
| MT345867.1 | 03/23/2020 | USA:WA |
| MT380734.1 | 03/23/2020 | USA |
| MT385418.1 | 03/23/2020 | USA:CA |
| MT385440.1 | 03/23/2020 | USA:CA |
| MT385446.1 | 03/23/2020 | USA:CA |
| MT385448.1 | 03/23/2020 | USA:CA |
| MT385458.1 | 03/23/2020 | USA:CA |
| MT385462.1 | 03/23/2020 | USA:CA |
| MT385464.1 | 03/23/2020 | USA:CA |
| MT385466.1 | 03/23/2020 | USA:CA |
| MT385476.1 | 03/23/2020 | USA:CA |
| MT385494.1 | 03/23/2020 | USA:CA |
| MT412190.1 | 03/23/2020 | USA:Michigan |
| MT412191.1 | 03/23/2020 | USA:Michigan |
| MT412194.1 | 03/23/2020 | USA:Michigan |
| MT419810.1 | 03/23/2020 | Puerto-Rico |
| MT419811.1 | 03/23/2020 | Puerto-Rico |
| MT419812.1 | 03/23/2020 | Puerto-Rico |
| MT419813.1 | 03/23/2020 | Puerto-Rico |
| MT419814.1 | 03/23/2020 | Puerto-Rico |
| MT419817.1 | 03/23/2020 | Puerto-Rico |
| MT419818.1 | 03/23/2020 | Puerto-Rico |
| MT439261.1 | 03/23/2020 | USA:MI |
| MT439262.1 | 03/23/2020 | USA:MI |
| MT439266.1 | 03/23/2020 | USA:MI |
| MT439267.1 | 03/23/2020 | USA:MI |
| MT439269.1 | 03/23/2020 | USA:MI |
| MT439270.1 | 03/23/2020 | USA:MI |
| MT439272.1 | 03/23/2020 | USA:MI |
| MT439280.1 | 03/23/2020 | USA:MI |
| MT439281.1 | 03/23/2020 | USA:MI |
| MT451042.1 | 03/23/2020 | Australia:Victoria |
| MT451184.1 | 03/23/2020 | Australia:Victoria |
| MT451185.1 | 03/23/2020 | Australia:Victoria |
| MT451186.1 | 03/23/2020 | Australia:Victoria |
| MT451187.1 | 03/23/2020 | Australia:Victoria |
| MT451188.1 | 03/23/2020 | Australia:Victoria |

| | | |
|---|---|---|
| MT451189.1 | 03/23/2020 | Australia:Victoria |
| MT451190.1 | 03/23/2020 | Australia:Victoria |
| MT451191.1 | 03/23/2020 | Australia:Victoria |
| MT451192.1 | 03/23/2020 | Australia:Victoria |
| MT451193.1 | 03/23/2020 | Australia:Victoria |
| MT451194.1 | 03/23/2020 | Australia:Victoria |
| MT451195.1 | 03/23/2020 | Australia:Victoria |
| MT451196.1 | 03/23/2020 | Australia:Victoria |
| MT451197.1 | 03/23/2020 | Australia:Victoria |
| MT451198.1 | 03/23/2020 | Australia:Victoria |
| MT451199.1 | 03/23/2020 | Australia:Victoria |
| MT451200.1 | 03/23/2020 | Australia:Victoria |
| MT451201.1 | 03/23/2020 | Australia:Victoria |
| MT451202.1 | 03/23/2020 | Australia:Victoria |
| MT451203.1 | 03/23/2020 | Australia:Victoria |
| MT451204.1 | 03/23/2020 | Australia:Victoria |
| MT451267.1 | 03/23/2020 | Australia:Victoria |
| MT451268.1 | 03/23/2020 | Australia:Victoria |
| MT451269.1 | 03/23/2020 | Australia:Victoria |
| MT451270.1 | 03/23/2020 | Australia:Victoria |
| MT451271.1 | 03/23/2020 | Australia:Victoria |
| MT451272.1 | 03/23/2020 | Australia:Victoria |
| MT451273.1 | 03/23/2020 | Australia:Victoria |
| MT451274.1 | 03/23/2020 | Australia:Victoria |
| MT451275.1 | 03/23/2020 | Australia:Victoria |
| MT451276.1 | 03/23/2020 | Australia:Victoria |
| MT451277.1 | 03/23/2020 | Australia:Victoria |
| MT451292.1 | 03/23/2020 | Australia:Victoria |
| MT451350.1 | 03/23/2020 | Australia:Victoria |
| MT451352.1 | 03/23/2020 | Australia:Victoria |
| MT451353.1 | 03/23/2020 | Australia:Victoria |
| MT451354.1 | 03/23/2020 | Australia:Victoria |
| MT451356.1 | 03/23/2020 | Australia:Victoria |
| MT451357.1 | 03/23/2020 | Australia:Victoria |
| MT451358.1 | 03/23/2020 | Australia:Victoria |
| MT451364.1 | 03/23/2020 | Australia:Victoria |
| MT451366.1 | 03/23/2020 | Australia:Victoria |
| MT451370.1 | 03/23/2020 | Australia:Victoria |
| MT451372.1 | 03/23/2020 | Australia:Victoria |
| MT451377.1 | 03/23/2020 | Australia:Victoria |
| MT451378.1 | 03/23/2020 | Australia:Victoria |
| MT451407.1 | 03/23/2020 | Australia:Victoria |
| MT451408.1 | 03/23/2020 | Australia:Victoria |
| MT451568.1 | 03/23/2020 | Australia:Victoria |
| MT451792.1 | 03/23/2020 | Australia:Victoria |
| MT459834.1 | 03/23/2020 | Greece:Athens |

| | | |
|---|---|---|
| MT459835.1 | 03/23/2020 | Greece:Athens |
| MT459840.1 | 03/23/2020 | Greece:Athens |
| MT459842.1 | 03/23/2020 | Greece:Athens |
| MT459843.1 | 03/23/2020 | Greece:Athens |
| MT459844.1 | 03/23/2020 | Greece:Athens |
| MT459903.1 | 03/23/2020 | Greece:Athens |
| MT459905.1 | 03/23/2020 | Greece:Athens |
| MT477842.1 | 03/23/2020 | USA:AK |
| MT477845.1 | 03/23/2020 | USA:AK |
| MT481992.1 | 03/23/2020 | USA |
| MT499183.1 | 03/23/2020 | USA:CA |
| MT506214.1 | 03/23/2020 | USA:Michigan |
| MT506219.1 | 03/23/2020 | USA:Michigan |
| MT506681.1 | 03/23/2020 | USA:Michigan |
| MT506685.1 | 03/23/2020 | USA:Michigan |
| MT509493.1 | 03/23/2020 | USA |
| MT509684.1 | 03/23/2020 | USA:Michigan |
| MT512421.1 | 03/23/2020 | USA |
| MT512422.1 | 03/23/2020 | USA |
| MT512423.1 | 03/23/2020 | USA |
| MT520229.1 | 03/23/2020 | USA:Massachusetts |
| MT520249.1 | 03/23/2020 | USA:Massachusetts |
| MT520297.1 | 03/23/2020 | USA:Massachusetts |
| MT520339.1 | 03/23/2020 | USA:Massachusetts |
| MT520346.1 | 03/23/2020 | USA:Massachusetts |
| MT520366.1 | 03/23/2020 | USA:Massachusetts |
| MT520387.1 | 03/23/2020 | USA:Massachusetts |
| MT520399.1 | 03/23/2020 | USA:Massachusetts |
| MT520466.1 | 03/23/2020 | USA:Massachusetts |
| MT520513.1 | 03/23/2020 | USA:Massachusetts |
| MT520519.1 | 03/23/2020 | USA:Massachusetts |
| MT520521.1 | 03/23/2020 | USA:Massachusetts |
| MT527184.1 | 03/23/2020 | Italy:Lazio |
| MT528235.1 | 03/23/2020 | Italy:Lazio |
| MT528238.1 | 03/23/2020 | Italy:Lazio |
| MT582452.1 | 03/23/2020 | Germany:Dusseldorf |
| MT627930.1 | 03/23/2020 | USA:Washington-King-County |
| MT628135.1 | 03/23/2020 | USA:CA |
| MT628136.1 | 03/23/2020 | USA:CA |
| MT628138.1 | 03/23/2020 | USA:CA |
| MT632510.1 | 03/23/2020 | USA:Washington-Whatcom-County |
| MT632522.1 | 03/23/2020 | USA:Washington-Whatcom-County |
| MT632532.1 | 03/23/2020 | USA:Washington-Whatcom-County |
| MT632544.1 | 03/23/2020 | USA:Washington-King-County |
| MT632630.1 | 03/23/2020 | USA:Washington-King-County |
| MT632864.1 | 03/23/2020 | USA:Washington |

| | | |
|---|---|---|
| MT632881.1 | 03/23/2020 | USA:Washington-Pierce-County |
| MT632906.1 | 03/23/2020 | USA:Washington |
| MT632913.1 | 03/23/2020 | USA:Washington |
| MT632947.1 | 03/23/2020 | USA:Washington-Adams-County |
| MT633004.1 | 03/23/2020 | USA:Washington-Adams-County |
| MT641662.1 | 03/23/2020 | Australia:Northern-Territory |
| MT641676.1 | 03/23/2020 | Australia:Victoria |
| MT641677.1 | 03/23/2020 | Australia:Victoria |
| MT641679.1 | 03/23/2020 | Australia:Victoria |
| MT641681.1 | 03/23/2020 | Australia:Victoria |
| MT641684.1 | 03/23/2020 | Australia:Victoria |
| MT641685.1 | 03/23/2020 | Australia:Victoria |
| MT641686.1 | 03/23/2020 | Australia:Victoria |
| MT641687.1 | 03/23/2020 | Australia:Victoria |
| MT646063.1 | 03/23/2020 | USA |
| MT263383.1 | 03/24/2020 | USA:WA |
| MT263390.1 | 03/24/2020 | USA:WA |
| MT263392.1 | 03/24/2020 | USA:WA |
| MT263398.1 | 03/24/2020 | USA:WA |
| MT263399.1 | 03/24/2020 | USA:WA |
| MT263404.1 | 03/24/2020 | USA:WA |
| MT263408.1 | 03/24/2020 | USA:WA |
| MT263414.1 | 03/24/2020 | USA:WA |
| MT263417.1 | 03/24/2020 | USA:WA |
| MT263421.1 | 03/24/2020 | USA:WA |
| MT263424.1 | 03/24/2020 | USA:WA |
| MT263425.1 | 03/24/2020 | USA:WA |
| MT263430.1 | 03/24/2020 | USA:WA |
| MT263431.1 | 03/24/2020 | USA:WA |
| MT263432.1 | 03/24/2020 | USA:WA |
| MT263434.1 | 03/24/2020 | USA:WA |
| MT263435.1 | 03/24/2020 | USA:WA |
| MT263436.1 | 03/24/2020 | USA:WA |
| MT263439.1 | 03/24/2020 | USA:WA |
| MT263440.1 | 03/24/2020 | USA:WA |
| MT263441.1 | 03/24/2020 | USA:WA |
| MT263442.1 | 03/24/2020 | USA:WA |
| MT263443.1 | 03/24/2020 | USA:WA |
| MT263445.1 | 03/24/2020 | USA:WA |
| MT263446.1 | 03/24/2020 | USA:WA |
| MT263449.1 | 03/24/2020 | USA:WA |
| MT263450.1 | 03/24/2020 | USA:WA |
| MT263453.1 | 03/24/2020 | USA:WA |
| MT263455.1 | 03/24/2020 | USA:WA |
| MT263456.1 | 03/24/2020 | USA:WA |
| MT263459.1 | 03/24/2020 | USA:WA |

| | | |
|---|---|---|
| MT326023.1 | 03/24/2020 | USA |
| MT326024.1 | 03/24/2020 | USA |
| MT326025.1 | 03/24/2020 | USA |
| MT326027.1 | 03/24/2020 | USA |
| MT326028.1 | 03/24/2020 | USA |
| MT326029.1 | 03/24/2020 | USA |
| MT326030.1 | 03/24/2020 | USA |
| MT326031.1 | 03/24/2020 | USA |
| MT326056.1 | 03/24/2020 | USA |
| MT326058.1 | 03/24/2020 | USA |
| MT326063.1 | 03/24/2020 | USA |
| MT326065.1 | 03/24/2020 | USA |
| MT326066.1 | 03/24/2020 | USA |
| MT326067.1 | 03/24/2020 | USA |
| MT326068.1 | 03/24/2020 | USA |
| MT326069.1 | 03/24/2020 | USA |
| MT326070.1 | 03/24/2020 | USA |
| MT326076.1 | 03/24/2020 | USA |
| MT326077.1 | 03/24/2020 | USA |
| MT326078.1 | 03/24/2020 | USA |
| MT326081.1 | 03/24/2020 | USA |
| MT334553.1 | 03/24/2020 | USA:UT |
| MT334559.1 | 03/24/2020 | USA:UT |
| MT345798.1 | 03/24/2020 | USA:WA |
| MT345811.1 | 03/24/2020 | USA:WA |
| MT345814.1 | 03/24/2020 | USA:WA |
| MT345815.1 | 03/24/2020 | USA:WA |
| MT345817.1 | 03/24/2020 | USA:ID |
| MT345840.1 | 03/24/2020 | USA:WA |
| MT345841.1 | 03/24/2020 | USA:WA |
| MT345842.1 | 03/24/2020 | USA:WA |
| MT345843.1 | 03/24/2020 | USA:WA |
| MT345844.1 | 03/24/2020 | USA:WA |
| MT345845.1 | 03/24/2020 | USA:WA |
| MT345846.1 | 03/24/2020 | USA:WA |
| MT345847.1 | 03/24/2020 | USA:WA |
| MT345848.1 | 03/24/2020 | USA:WA |
| MT345849.1 | 03/24/2020 | USA:WA |
| MT345850.1 | 03/24/2020 | USA:WA |
| MT345857.1 | 03/24/2020 | USA:ID |
| MT394528.1 | 03/24/2020 | USA:CA |
| MT394529.1 | 03/24/2020 | USA:CA |
| MT407654.1 | 03/24/2020 | China:Zhejiang |
| MT407655.1 | 03/24/2020 | China:Zhejiang |
| MT407656.1 | 03/24/2020 | China:Zhejiang |
| MT407657.1 | 03/24/2020 | China:Zhejiang |

| | | |
|---|---|---|
| MT407658.1 | 03/24/2020 | China:Zhejiang |
| MT407659.1 | 03/24/2020 | China:Zhejiang |
| MT412195.1 | 03/24/2020 | USA:Michigan |
| MT412197.1 | 03/24/2020 | USA:Michigan |
| MT412198.1 | 03/24/2020 | USA:Michigan |
| MT419815.1 | 03/24/2020 | Puerto-Rico |
| MT419816.1 | 03/24/2020 | Puerto-Rico |
| MT419819.1 | 03/24/2020 | Puerto-Rico |
| MT419820.1 | 03/24/2020 | Puerto-Rico |
| MT439273.1 | 03/24/2020 | USA:MI |
| MT439274.1 | 03/24/2020 | USA:MI |
| MT439275.1 | 03/24/2020 | USA:MI |
| MT439276.1 | 03/24/2020 | USA:MI |
| MT439277.1 | 03/24/2020 | USA:MI |
| MT439278.1 | 03/24/2020 | USA:MI |
| MT439279.1 | 03/24/2020 | USA:MI |
| MT439283.1 | 03/24/2020 | USA:MI |
| MT439284.1 | 03/24/2020 | USA:MI |
| MT439285.1 | 03/24/2020 | USA:MI |
| MT439286.1 | 03/24/2020 | USA:MI |
| MT439287.1 | 03/24/2020 | USA:MI |
| MT439293.1 | 03/24/2020 | USA:MI |
| MT439296.1 | 03/24/2020 | USA:MI |
| MT439297.1 | 03/24/2020 | USA:MI |
| MT439298.1 | 03/24/2020 | USA:MI |
| MT451205.1 | 03/24/2020 | Australia:Victoria |
| MT451206.1 | 03/24/2020 | Australia:Victoria |
| MT451207.1 | 03/24/2020 | Australia:Victoria |
| MT451208.1 | 03/24/2020 | Australia:Victoria |
| MT451209.1 | 03/24/2020 | Australia:Victoria |
| MT451210.1 | 03/24/2020 | Australia:Victoria |
| MT451211.1 | 03/24/2020 | Australia:Victoria |
| MT451212.1 | 03/24/2020 | Australia:Victoria |
| MT451213.1 | 03/24/2020 | Australia:Victoria |
| MT451214.1 | 03/24/2020 | Australia:Victoria |
| MT451215.1 | 03/24/2020 | Australia:Victoria |
| MT451216.1 | 03/24/2020 | Australia:Victoria |
| MT451217.1 | 03/24/2020 | Australia:Victoria |
| MT451218.1 | 03/24/2020 | Australia:Victoria |
| MT451278.1 | 03/24/2020 | Australia:Victoria |
| MT451279.1 | 03/24/2020 | Australia:Victoria |
| MT451280.1 | 03/24/2020 | Australia:Victoria |
| MT451281.1 | 03/24/2020 | Australia:Victoria |
| MT451282.1 | 03/24/2020 | Australia:Victoria |
| MT451283.1 | 03/24/2020 | Australia:Victoria |
| MT451284.1 | 03/24/2020 | Australia:Victoria |

| | | |
|---|---|---|
| MT451316.1 | 03/24/2020 | Australia:Victoria |
| MT451317.1 | 03/24/2020 | Australia:Victoria |
| MT451355.1 | 03/24/2020 | Australia:Victoria |
| MT451359.1 | 03/24/2020 | Australia:Victoria |
| MT451360.1 | 03/24/2020 | Australia:Victoria |
| MT451361.1 | 03/24/2020 | Australia:Victoria |
| MT451362.1 | 03/24/2020 | Australia:Victoria |
| MT451363.1 | 03/24/2020 | Australia:Victoria |
| MT451365.1 | 03/24/2020 | Australia:Victoria |
| MT451367.1 | 03/24/2020 | Australia:Victoria |
| MT451369.1 | 03/24/2020 | Australia:Victoria |
| MT451371.1 | 03/24/2020 | Australia:Victoria |
| MT451373.1 | 03/24/2020 | Australia:Victoria |
| MT451374.1 | 03/24/2020 | Australia:Victoria |
| MT451384.1 | 03/24/2020 | Australia:Victoria |
| MT451387.1 | 03/24/2020 | Australia:Victoria |
| MT451388.1 | 03/24/2020 | Australia:Victoria |
| MT451402.1 | 03/24/2020 | Australia:Victoria |
| MT451448.1 | 03/24/2020 | Australia:Victoria |
| MT451569.1 | 03/24/2020 | Australia:Victoria |
| MT451630.1 | 03/24/2020 | Australia:Victoria |
| MT457402.1 | 03/24/2020 | India:Hyderabad |
| MT459832.1 | 03/24/2020 | Greece:Athens |
| MT459986.1 | 03/24/2020 | Guam |
| MT477844.1 | 03/24/2020 | USA:AK |
| MT477853.1 | 03/24/2020 | USA:AK |
| MT477854.1 | 03/24/2020 | USA:AK |
| MT499191.1 | 03/24/2020 | USA:CA |
| MT499192.1 | 03/24/2020 | USA:CA |
| MT499193.1 | 03/24/2020 | USA:CA |
| MT506215.1 | 03/24/2020 | USA:Michigan |
| MT506218.1 | 03/24/2020 | USA:Michigan |
| MT506220.1 | 03/24/2020 | USA:Michigan |
| MT506222.1 | 03/24/2020 | USA:Michigan |
| MT506224.1 | 03/24/2020 | USA:Michigan |
| MT506225.1 | 03/24/2020 | USA:Michigan |
| MT506683.1 | 03/24/2020 | USA:Michigan |
| MT506684.1 | 03/24/2020 | USA:Michigan |
| MT506686.1 | 03/24/2020 | USA:Michigan |
| MT506687.1 | 03/24/2020 | USA:Michigan |
| MT506688.1 | 03/24/2020 | USA:Michigan |
| MT509458.1 | 03/24/2020 | USA |
| MT509462.1 | 03/24/2020 | USA |
| MT509479.1 | 03/24/2020 | USA |
| MT509682.1 | 03/24/2020 | USA:Michigan |
| MT509683.1 | 03/24/2020 | USA:Michigan |

| | | |
|---|---|---|
| MT520179.1 | 03/24/2020 | USA:Massachusetts |
| MT520220.1 | 03/24/2020 | USA:Massachusetts |
| MT520224.1 | 03/24/2020 | USA:Massachusetts |
| MT520289.1 | 03/24/2020 | USA:Massachusetts |
| MT520315.1 | 03/24/2020 | USA:Massachusetts |
| MT520326.1 | 03/24/2020 | USA:Massachusetts |
| MT520356.1 | 03/24/2020 | USA:Massachusetts |
| MT520378.1 | 03/24/2020 | USA:Massachusetts |
| MT520401.1 | 03/24/2020 | USA:Massachusetts |
| MT520409.1 | 03/24/2020 | USA:Massachusetts |
| MT520450.1 | 03/24/2020 | USA:Massachusetts |
| MT533206.1 | 03/24/2020 | USA:CA |
| MT559038.1 | 03/24/2020 | Tunisia:Ben-Arous |
| MT598174.1 | 03/24/2020 | USA:San-Diego-California |
| MT628137.1 | 03/24/2020 | USA:CA |
| MT628139.1 | 03/24/2020 | USA:CA |
| MT628173.1 | 03/24/2020 | USA:CA |
| MT628240.1 | 03/24/2020 | USA:CA |
| MT628241.1 | 03/24/2020 | USA:CA |
| MT628242.1 | 03/24/2020 | USA:CA |
| MT632780.1 | 03/24/2020 | USA:Washington-Yakima-County |
| MT632791.1 | 03/24/2020 | USA:Washington-Yakima-County |
| MT632799.1 | 03/24/2020 | USA:Washington-Yakima-County |
| MT632808.1 | 03/24/2020 | USA:Washington-Yakima-County |
| MT632814.1 | 03/24/2020 | USA:Washington-Yakima-County |
| MT632816.1 | 03/24/2020 | USA:Washington-Yakima-County |
| MT632820.1 | 03/24/2020 | USA:Washington-King-County |
| MT632826.1 | 03/24/2020 | USA:Washington-Yakima-County |
| MT632831.1 | 03/24/2020 | USA:Washington-Whatcom-County |
| MT632835.1 | 03/24/2020 | USA:Washington-Whatcom-County |
| MT632844.1 | 03/24/2020 | USA:Washington-Whatcom-County |
| MT632854.1 | 03/24/2020 | USA:Washington-Yakima-County |
| MT632869.1 | 03/24/2020 | USA:Washington-Yakima-County |
| MT632878.1 | 03/24/2020 | USA:Washington-Yakima-County |
| MT632912.1 | 03/24/2020 | USA:Washington-Whatcom-County |
| MT632929.1 | 03/24/2020 | USA:Washington |
| MT632935.1 | 03/24/2020 | USA:Washington-Pierce-County |
| MT632943.1 | 03/24/2020 | USA:Washington-King-County |
| MT633001.1 | 03/24/2020 | USA:Washington-Kittitas-County |
| MT641660.1 | 03/24/2020 | Australia:Northern-Territory |
| MT641661.1 | 03/24/2020 | Australia:Northern-Territory |
| MT641667.1 | 03/24/2020 | Australia:Victoria |
| MT641688.1 | 03/24/2020 | Australia:Victoria |
| MT641689.1 | 03/24/2020 | Australia:Victoria |
| MT641713.1 | 03/24/2020 | Australia:Victoria |
| MT646054.1 | 03/24/2020 | USA |

| | | |
|---|---|---|
| MT646062.1 | 03/24/2020 | USA |
| MT646073.1 | 03/24/2020 | USA |
| MT646087.1 | 03/24/2020 | USA |
| MT646088.1 | 03/24/2020 | USA |
| MT646102.1 | 03/24/2020 | USA |
| MT324684.1 | 03/25/2020 | USA |
| MT334560.1 | 03/25/2020 | USA:UT |
| MT334564.1 | 03/25/2020 | USA:UT |
| MT334566.1 | 03/25/2020 | USA:UT |
| MT345800.1 | 03/25/2020 | USA:WA |
| MT345801.1 | 03/25/2020 | USA:WA |
| MT345806.1 | 03/25/2020 | USA:WA |
| MT345808.1 | 03/25/2020 | USA:WA |
| MT345809.1 | 03/25/2020 | USA:WA |
| MT345812.1 | 03/25/2020 | USA:WA |
| MT345818.1 | 03/25/2020 | USA:WA |
| MT345819.1 | 03/25/2020 | USA:WA |
| MT345821.1 | 03/25/2020 | USA:WA |
| MT345822.1 | 03/25/2020 | USA:WA |
| MT345823.1 | 03/25/2020 | USA:WA |
| MT345824.1 | 03/25/2020 | USA:WA |
| MT345825.1 | 03/25/2020 | USA:WA |
| MT345826.1 | 03/25/2020 | USA:WA |
| MT345827.1 | 03/25/2020 | USA:WA |
| MT345829.1 | 03/25/2020 | USA:WA |
| MT345830.1 | 03/25/2020 | USA:WA |
| MT345832.1 | 03/25/2020 | USA:WA |
| MT345833.1 | 03/25/2020 | USA:WA |
| MT345835.1 | 03/25/2020 | USA:WA |
| MT345851.1 | 03/25/2020 | USA:WA |
| MT345852.1 | 03/25/2020 | USA:WA |
| MT345853.1 | 03/25/2020 | USA:WA |
| MT345854.1 | 03/25/2020 | USA:WA |
| MT345855.1 | 03/25/2020 | USA:WA |
| MT345856.1 | 03/25/2020 | USA:WA |
| MT345859.1 | 03/25/2020 | USA:WA |
| MT345860.1 | 03/25/2020 | USA:WA |
| MT345861.1 | 03/25/2020 | USA:WA |
| MT345862.1 | 03/25/2020 | USA:WA |
| MT345865.1 | 03/25/2020 | USA:WA |
| MT358683.1 | 03/25/2020 | USA:WA |
| MT358684.1 | 03/25/2020 | USA:WA |
| MT358692.1 | 03/25/2020 | USA:WA |
| MT358693.1 | 03/25/2020 | USA:CT |
| MT358698.1 | 03/25/2020 | USA:WA |
| MT358701.1 | 03/25/2020 | USA:WA |

| | | |
|---|---|---|
| MT358703.1 | 03/25/2020 | USA:ID |
| MT375447.1 | 03/25/2020 | USA:WA |
| MT385421.1 | 03/25/2020 | USA:CA |
| MT385425.1 | 03/25/2020 | USA:CA |
| MT385436.1 | 03/25/2020 | USA:CA |
| MT385441.1 | 03/25/2020 | USA:CA |
| MT385445.1 | 03/25/2020 | USA:CA |
| MT385459.1 | 03/25/2020 | USA:CA |
| MT385461.1 | 03/25/2020 | USA:CA |
| MT385463.1 | 03/25/2020 | USA:CA |
| MT385469.1 | 03/25/2020 | USA:CA |
| MT385472.1 | 03/25/2020 | USA:CA |
| MT385487.1 | 03/25/2020 | USA:CA |
| MT385488.1 | 03/25/2020 | USA:CA |
| MT385492.1 | 03/25/2020 | USA:CA |
| MT385493.1 | 03/25/2020 | USA:CA |
| MT385497.1 | 03/25/2020 | USA:CA |
| MT412199.1 | 03/25/2020 | USA:Michigan |
| MT412201.1 | 03/25/2020 | USA:Michigan |
| MT412204.1 | 03/25/2020 | USA:Michigan |
| MT439288.1 | 03/25/2020 | USA:MI |
| MT439289.1 | 03/25/2020 | USA:MI |
| MT439290.1 | 03/25/2020 | USA:MI |
| MT439291.1 | 03/25/2020 | USA:MI |
| MT439292.1 | 03/25/2020 | USA:MI |
| MT439299.1 | 03/25/2020 | USA:MI |
| MT439301.1 | 03/25/2020 | USA:MI |
| MT439302.1 | 03/25/2020 | USA:MI |
| MT451285.1 | 03/25/2020 | Australia:Victoria |
| MT451286.1 | 03/25/2020 | Australia:Victoria |
| MT451291.1 | 03/25/2020 | Australia:Victoria |
| MT451293.1 | 03/25/2020 | Australia:Victoria |
| MT451294.1 | 03/25/2020 | Australia:Victoria |
| MT451296.1 | 03/25/2020 | Australia:Victoria |
| MT451297.1 | 03/25/2020 | Australia:Victoria |
| MT451298.1 | 03/25/2020 | Australia:Victoria |
| MT451299.1 | 03/25/2020 | Australia:Victoria |
| MT451300.1 | 03/25/2020 | Australia:Victoria |
| MT451301.1 | 03/25/2020 | Australia:Victoria |
| MT451302.1 | 03/25/2020 | Australia:Victoria |
| MT451305.1 | 03/25/2020 | Australia:Victoria |
| MT451306.1 | 03/25/2020 | Australia:Victoria |
| MT451307.1 | 03/25/2020 | Australia:Victoria |
| MT451318.1 | 03/25/2020 | Australia:Victoria |
| MT451319.1 | 03/25/2020 | Australia:Victoria |
| MT451320.1 | 03/25/2020 | Australia:Victoria |

| | | |
|---|---|---|
| MT451321.1 | 03/25/2020 | Australia:Victoria |
| MT451322.1 | 03/25/2020 | Australia:Victoria |
| MT451323.1 | 03/25/2020 | Australia:Victoria |
| MT451368.1 | 03/25/2020 | Australia:Victoria |
| MT451375.1 | 03/25/2020 | Australia:Victoria |
| MT451376.1 | 03/25/2020 | Australia:Victoria |
| MT451379.1 | 03/25/2020 | Australia:Victoria |
| MT451380.1 | 03/25/2020 | Australia:Victoria |
| MT451381.1 | 03/25/2020 | Australia:Victoria |
| MT451382.1 | 03/25/2020 | Australia:Victoria |
| MT451383.1 | 03/25/2020 | Australia:Victoria |
| MT451385.1 | 03/25/2020 | Australia:Victoria |
| MT451386.1 | 03/25/2020 | Australia:Victoria |
| MT451389.1 | 03/25/2020 | Australia:Victoria |
| MT451390.1 | 03/25/2020 | Australia:Victoria |
| MT451391.1 | 03/25/2020 | Australia:Victoria |
| MT451392.1 | 03/25/2020 | Australia:Victoria |
| MT451396.1 | 03/25/2020 | Australia:Victoria |
| MT451397.1 | 03/25/2020 | Australia:Victoria |
| MT451398.1 | 03/25/2020 | Australia:Victoria |
| MT451399.1 | 03/25/2020 | Australia:Victoria |
| MT451400.1 | 03/25/2020 | Australia:Victoria |
| MT451401.1 | 03/25/2020 | Australia:Victoria |
| MT451403.1 | 03/25/2020 | Australia:Victoria |
| MT451409.1 | 03/25/2020 | Australia:Victoria |
| MT451410.1 | 03/25/2020 | Australia:Victoria |
| MT451411.1 | 03/25/2020 | Australia:Victoria |
| MT451412.1 | 03/25/2020 | Australia:Victoria |
| MT451435.1 | 03/25/2020 | Australia:Victoria |
| MT451638.1 | 03/25/2020 | Australia:Victoria |
| MT451639.1 | 03/25/2020 | Australia:Victoria |
| MT451640.1 | 03/25/2020 | Australia:Victoria |
| MT457403.1 | 03/25/2020 | India:Hyderabad |
| MT474129.1 | 03/25/2020 | USA:CA |
| MT477852.1 | 03/25/2020 | USA:AK |
| MT477885.1 | 03/25/2020 | India |
| MT481928.1 | 03/25/2020 | USA:CA |
| MT499180.1 | 03/25/2020 | USA:CA |
| MT499181.1 | 03/25/2020 | USA:CA |
| MT499186.1 | 03/25/2020 | USA:CA |
| MT499187.1 | 03/25/2020 | USA:CA |
| MT499188.1 | 03/25/2020 | USA:CA |
| MT499189.1 | 03/25/2020 | USA:CA |
| MT506217.1 | 03/25/2020 | USA:Michigan |
| MT506221.1 | 03/25/2020 | USA:Michigan |
| MT506223.1 | 03/25/2020 | USA:Michigan |

| | | |
|---|---|---|
| MT506226.1 | 03/25/2020 | USA:Michigan |
| MT506544.1 | 03/25/2020 | USA:Michigan |
| MT506545.1 | 03/25/2020 | USA:Michigan |
| MT506548.1 | 03/25/2020 | USA:Michigan |
| MT506549.1 | 03/25/2020 | USA:Michigan |
| MT509685.1 | 03/25/2020 | USA:Michigan |
| MT509686.1 | 03/25/2020 | USA:Michigan |
| MT520176.1 | 03/25/2020 | USA:Massachusetts |
| MT520177.1 | 03/25/2020 | USA:Massachusetts |
| MT520180.1 | 03/25/2020 | USA:Massachusetts |
| MT520184.1 | 03/25/2020 | USA:Massachusetts |
| MT520186.1 | 03/25/2020 | USA:Massachusetts |
| MT520194.1 | 03/25/2020 | USA:Massachusetts |
| MT520197.1 | 03/25/2020 | USA:Massachusetts |
| MT520201.1 | 03/25/2020 | USA:Massachusetts |
| MT520203.1 | 03/25/2020 | USA:Massachusetts |
| MT520218.1 | 03/25/2020 | USA:Massachusetts |
| MT520250.1 | 03/25/2020 | USA:Massachusetts |
| MT520259.1 | 03/25/2020 | USA:Massachusetts |
| MT520275.1 | 03/25/2020 | USA:Massachusetts |
| MT520277.1 | 03/25/2020 | USA:Massachusetts |
| MT520281.1 | 03/25/2020 | USA:Massachusetts |
| MT520293.1 | 03/25/2020 | USA:Massachusetts |
| MT520294.1 | 03/25/2020 | USA:Massachusetts |
| MT520367.1 | 03/25/2020 | USA:Massachusetts |
| MT520372.1 | 03/25/2020 | USA:Massachusetts |
| MT520377.1 | 03/25/2020 | USA:Massachusetts |
| MT520385.1 | 03/25/2020 | USA:Massachusetts |
| MT520394.1 | 03/25/2020 | USA:Massachusetts |
| MT520395.1 | 03/25/2020 | USA:Massachusetts |
| MT520410.1 | 03/25/2020 | USA:Massachusetts |
| MT520439.1 | 03/25/2020 | USA:Massachusetts |
| MT520444.1 | 03/25/2020 | USA:Massachusetts |
| MT520496.1 | 03/25/2020 | USA:Massachusetts |
| MT520523.1 | 03/25/2020 | USA:Massachusetts |
| MT520544.1 | 03/25/2020 | USA:Massachusetts |
| MT520545.1 | 03/25/2020 | USA:Massachusetts |
| MT535474.1 | 03/25/2020 | USA |
| MT628140.1 | 03/25/2020 | USA:CA |
| MT628141.1 | 03/25/2020 | USA:CA |
| MT628142.1 | 03/25/2020 | USA:CA |
| MT628243.1 | 03/25/2020 | USA:CA |
| MT632629.1 | 03/25/2020 | USA:Washington-King-County |
| MT632793.1 | 03/25/2020 | USA:Washington-Yakima-County |
| MT632829.1 | 03/25/2020 | USA:Washington-Yakima-County |
| MT632834.1 | 03/25/2020 | USA:Washington |

| | | |
|---|---|---|
| MT632855.1 | 03/25/2020 | USA:Washington-Whatcom-County |
| MT632857.1 | 03/25/2020 | USA:Washington |
| MT632870.1 | 03/25/2020 | USA:Washington |
| MT632885.1 | 03/25/2020 | USA:Washington-Yakima-County |
| MT632902.1 | 03/25/2020 | USA:Washington-Yakima-County |
| MT632908.1 | 03/25/2020 | USA:Washington-Grant-County |
| MT632927.1 | 03/25/2020 | USA:Washington-Yakima-County |
| MT632938.1 | 03/25/2020 | USA:Washington |
| MT632953.1 | 03/25/2020 | USA:Washington-King-County |
| MT632959.1 | 03/25/2020 | USA:Washington-King-County |
| MT632968.1 | 03/25/2020 | USA:Washington-Yakima-County |
| MT632973.1 | 03/25/2020 | USA:Washington-Yakima-County |
| MT632975.1 | 03/25/2020 | USA:Washington-Yakima-County |
| MT632980.1 | 03/25/2020 | USA:Washington |
| MT632986.1 | 03/25/2020 | USA:Washington-Yakima-County |
| MT632988.1 | 03/25/2020 | USA:Washington-Yakima-County |
| MT632989.1 | 03/25/2020 | USA:Washington-Yakima-County |
| MT632994.1 | 03/25/2020 | USA:Washington-Yakima-County |
| MT632996.1 | 03/25/2020 | USA:Washington-Yakima-County |
| MT633014.1 | 03/25/2020 | USA:Washington-Yakima-County |
| MT633015.1 | 03/25/2020 | USA:Washington-Yakima-County |
| MT633026.1 | 03/25/2020 | USA:Washington-Yakima-County |
| MT633028.1 | 03/25/2020 | USA:Washington-Yakima-County |
| MT633032.1 | 03/25/2020 | USA:Washington-Yakima-County |
| MT633034.1 | 03/25/2020 | USA:Washington-Yakima-County |
| MT641657.1 | 03/25/2020 | Australia:Northern-Territory |
| MT641658.1 | 03/25/2020 | Australia:Northern-Territory |
| MT641663.1 | 03/25/2020 | Australia:Northern-Territory |
| MT641671.1 | 03/25/2020 | Australia:Victoria |
| MT641692.1 | 03/25/2020 | Australia:Victoria |
| MT641693.1 | 03/25/2020 | Australia:Victoria |
| MT641695.1 | 03/25/2020 | Australia:Victoria |
| MT641697.1 | 03/25/2020 | Australia:Victoria |
| MT641698.1 | 03/25/2020 | Australia:Victoria |
| MT641728.1 | 03/25/2020 | Australia:Victoria |
| MT646093.1 | 03/25/2020 | USA |
| MT646106.1 | 03/25/2020 | USA |
| MT293178.1 | 03/26/2020 | USA:WA |
| MT334561.1 | 03/26/2020 | USA:UT |
| MT334563.1 | 03/26/2020 | USA:UT |
| MT334565.1 | 03/26/2020 | USA:UT |
| MT334567.1 | 03/26/2020 | USA:UT |
| MT334570.1 | 03/26/2020 | USA:UT |
| MT334571.1 | 03/26/2020 | USA:UT |
| MT345834.1 | 03/26/2020 | USA:WA |
| MT358674.1 | 03/26/2020 | USA:WA |

| | | |
|---|---|---|
| MT358675.1 | 03/26/2020 | USA:WA |
| MT358676.1 | 03/26/2020 | USA:WA |
| MT358677.1 | 03/26/2020 | USA:WA |
| MT358679.1 | 03/26/2020 | USA:WA |
| MT358680.1 | 03/26/2020 | USA:WA |
| MT358681.1 | 03/26/2020 | USA:WA |
| MT358682.1 | 03/26/2020 | USA:WA |
| MT358685.1 | 03/26/2020 | USA:WA |
| MT358686.1 | 03/26/2020 | USA:WA |
| MT358687.1 | 03/26/2020 | USA:WA |
| MT358688.1 | 03/26/2020 | USA:WA |
| MT358689.1 | 03/26/2020 | USA:WA |
| MT358690.1 | 03/26/2020 | USA:WA |
| MT358694.1 | 03/26/2020 | USA:WA |
| MT358695.1 | 03/26/2020 | USA:WA |
| MT358696.1 | 03/26/2020 | USA:WA |
| MT358697.1 | 03/26/2020 | USA:WA |
| MT358699.1 | 03/26/2020 | USA:WA |
| MT358700.1 | 03/26/2020 | USA:WA |
| MT358709.1 | 03/26/2020 | USA:ID |
| MT375441.1 | 03/26/2020 | USA:WA |
| MT375442.1 | 03/26/2020 | USA:WA |
| MT375443.1 | 03/26/2020 | USA:WA |
| MT412200.1 | 03/26/2020 | USA:Michigan |
| MT412202.1 | 03/26/2020 | USA:Michigan |
| MT412203.1 | 03/26/2020 | USA:Michigan |
| MT412205.1 | 03/26/2020 | USA:Michigan |
| MT412206.1 | 03/26/2020 | USA:Michigan |
| MT412207.1 | 03/26/2020 | USA:Michigan |
| MT412208.1 | 03/26/2020 | USA:Michigan |
| MT412212.1 | 03/26/2020 | USA:Michigan |
| MT412216.1 | 03/26/2020 | USA:Michigan |
| MT429183.1 | 03/26/2020 | USA:Wisconsin |
| MT439294.1 | 03/26/2020 | USA:MI |
| MT439295.1 | 03/26/2020 | USA:MI |
| MT439300.1 | 03/26/2020 | USA:MI |
| MT439303.1 | 03/26/2020 | USA:MI |
| MT439305.1 | 03/26/2020 | USA:MI |
| MT439317.1 | 03/26/2020 | USA:MI |
| MT447177.1 | 03/26/2020 | Iran |
| MT451287.1 | 03/26/2020 | Australia:Victoria |
| MT451288.1 | 03/26/2020 | Australia:Victoria |
| MT451289.1 | 03/26/2020 | Australia:Victoria |
| MT451290.1 | 03/26/2020 | Australia:Victoria |
| MT451303.1 | 03/26/2020 | Australia:Victoria |
| MT451304.1 | 03/26/2020 | Australia:Victoria |

| | | |
|---|---|---|
| MT451308.1 | 03/26/2020 | Australia:Victoria |
| MT451309.1 | 03/26/2020 | Australia:Victoria |
| MT451310.1 | 03/26/2020 | Australia:Victoria |
| MT451311.1 | 03/26/2020 | Australia:Victoria |
| MT451324.1 | 03/26/2020 | Australia:Victoria |
| MT451325.1 | 03/26/2020 | Australia:Victoria |
| MT451328.1 | 03/26/2020 | Australia:Victoria |
| MT451329.1 | 03/26/2020 | Australia:Victoria |
| MT451333.1 | 03/26/2020 | Australia:Victoria |
| MT451393.1 | 03/26/2020 | Australia:Victoria |
| MT451394.1 | 03/26/2020 | Australia:Victoria |
| MT451395.1 | 03/26/2020 | Australia:Victoria |
| MT451413.1 | 03/26/2020 | Australia:Victoria |
| MT451418.1 | 03/26/2020 | Australia:Victoria |
| MT451420.1 | 03/26/2020 | Australia:Victoria |
| MT451421.1 | 03/26/2020 | Australia:Victoria |
| MT451422.1 | 03/26/2020 | Australia:Victoria |
| MT451424.1 | 03/26/2020 | Australia:Victoria |
| MT451425.1 | 03/26/2020 | Australia:Victoria |
| MT451436.1 | 03/26/2020 | Australia:Victoria |
| MT451471.1 | 03/26/2020 | Australia:Victoria |
| MT451641.1 | 03/26/2020 | Australia:Victoria |
| MT451642.1 | 03/26/2020 | Australia:Victoria |
| MT451643.1 | 03/26/2020 | Australia:Victoria |
| MT451644.1 | 03/26/2020 | Australia:Victoria |
| MT451645.1 | 03/26/2020 | Australia:Victoria |
| MT451646.1 | 03/26/2020 | Australia:Victoria |
| MT451647.1 | 03/26/2020 | Australia:Victoria |
| MT451648.1 | 03/26/2020 | Australia:Victoria |
| MT459846.1 | 03/26/2020 | Greece:Athens |
| MT459928.1 | 03/26/2020 | Iran |
| MT459985.1 | 03/26/2020 | Guam |
| MT459990.1 | 03/26/2020 | USA:SC |
| MT499175.1 | 03/26/2020 | USA:CA |
| MT499176.1 | 03/26/2020 | USA:CA |
| MT499177.1 | 03/26/2020 | USA:CA |
| MT499178.1 | 03/26/2020 | USA:CA |
| MT499185.1 | 03/26/2020 | USA:CA |
| MT499202.1 | 03/26/2020 | USA:CA |
| MT506227.1 | 03/26/2020 | USA:Michigan |
| MT506547.1 | 03/26/2020 | USA:Michigan |
| MT506552.1 | 03/26/2020 | USA:Michigan |
| MT506689.1 | 03/26/2020 | USA:Michigan |
| MT506690.1 | 03/26/2020 | USA:Michigan |
| MT506693.1 | 03/26/2020 | USA:Michigan |
| MT509480.1 | 03/26/2020 | USA |

| | | |
|---|---|---|
| MT509484.1 | 03/26/2020 | USA |
| MT509491.1 | 03/26/2020 | USA |
| MT509688.1 | 03/26/2020 | USA:Michigan |
| MT517422.1 | 03/26/2020 | Czech-Republic |
| MT517427.1 | 03/26/2020 | Czech-Republic |
| MT517428.1 | 03/26/2020 | Czech-Republic |
| MT520182.1 | 03/26/2020 | USA:Massachusetts |
| MT520189.1 | 03/26/2020 | USA:Massachusetts |
| MT520199.1 | 03/26/2020 | USA:Massachusetts |
| MT520225.1 | 03/26/2020 | USA:Massachusetts |
| MT520245.1 | 03/26/2020 | USA:Massachusetts |
| MT520254.1 | 03/26/2020 | USA:Massachusetts |
| MT520258.1 | 03/26/2020 | USA:Massachusetts |
| MT520274.1 | 03/26/2020 | USA:Massachusetts |
| MT520278.1 | 03/26/2020 | USA:Massachusetts |
| MT520284.1 | 03/26/2020 | USA:Massachusetts |
| MT520296.1 | 03/26/2020 | USA:Massachusetts |
| MT520311.1 | 03/26/2020 | USA:Massachusetts |
| MT520313.1 | 03/26/2020 | USA:Massachusetts |
| MT520336.1 | 03/26/2020 | USA:Massachusetts |
| MT520338.1 | 03/26/2020 | USA:Massachusetts |
| MT520355.1 | 03/26/2020 | USA:Massachusetts |
| MT520365.1 | 03/26/2020 | USA:Massachusetts |
| MT520376.1 | 03/26/2020 | USA:Massachusetts |
| MT520379.1 | 03/26/2020 | USA:Massachusetts |
| MT520406.1 | 03/26/2020 | USA:Massachusetts |
| MT520414.1 | 03/26/2020 | USA:Massachusetts |
| MT520423.1 | 03/26/2020 | USA:Massachusetts |
| MT520430.1 | 03/26/2020 | USA:Massachusetts |
| MT520434.1 | 03/26/2020 | USA:Massachusetts |
| MT520442.1 | 03/26/2020 | USA:Massachusetts |
| MT520443.1 | 03/26/2020 | USA:Massachusetts |
| MT520447.1 | 03/26/2020 | USA:Massachusetts |
| MT520469.1 | 03/26/2020 | USA:Massachusetts |
| MT520484.1 | 03/26/2020 | USA:Massachusetts |
| MT520492.1 | 03/26/2020 | USA:Massachusetts |
| MT520522.1 | 03/26/2020 | USA:Massachusetts |
| MT520526.1 | 03/26/2020 | USA:Massachusetts |
| MT520528.1 | 03/26/2020 | USA:Massachusetts |
| MT520532.1 | 03/26/2020 | USA:Massachusetts |
| MT520537.1 | 03/26/2020 | USA:Massachusetts |
| MT520541.1 | 03/26/2020 | USA:Massachusetts |
| MT535472.1 | 03/26/2020 | USA |
| MT535473.1 | 03/26/2020 | USA |
| MT535475.1 | 03/26/2020 | USA |
| MT535477.1 | 03/26/2020 | USA |

| | | |
|---|---|---|
| MT535478.1 | 03/26/2020 | USA |
| MT612108.1 | 03/26/2020 | Australia:Victoria |
| MT627932.1 | 03/26/2020 | USA:Washington-Snohomish-County |
| MT628143.1 | 03/26/2020 | USA:CA |
| MT628244.1 | 03/26/2020 | USA:CA |
| MT628245.1 | 03/26/2020 | USA:CA |
| MT632781.1 | 03/26/2020 | USA:Washington-Whatcom-County |
| MT632800.1 | 03/26/2020 | USA:Washington-Whatcom-County |
| MT632802.1 | 03/26/2020 | USA:Washington-Whatcom-County |
| MT632806.1 | 03/26/2020 | USA:Washington-King-County |
| MT632813.1 | 03/26/2020 | USA:Washington-Whatcom-County |
| MT632867.1 | 03/26/2020 | USA:Washington-Yakima-County |
| MT632901.1 | 03/26/2020 | USA:Washington-Yakima-County |
| MT632903.1 | 03/26/2020 | USA:Washington-Yakima-County |
| MT632918.1 | 03/26/2020 | USA:Washington-Yakima-County |
| MT632926.1 | 03/26/2020 | USA:Washington-Yakima-County |
| MT632944.1 | 03/26/2020 | USA:Washington-Yakima-County |
| MT632956.1 | 03/26/2020 | USA:Washington-King-County |
| MT632965.1 | 03/26/2020 | USA:Washington-Yakima-County |
| MT632991.1 | 03/26/2020 | USA:Washington-Yakima-County |
| MT633002.1 | 03/26/2020 | USA:Washington-Yakima-County |
| MT633005.1 | 03/26/2020 | USA:Washington-King-County |
| MT633007.1 | 03/26/2020 | USA:Washington-Yakima-County |
| MT633044.1 | 03/26/2020 | USA:Washington-Franklin-County |
| MT633045.1 | 03/26/2020 | USA:Washington-Yakima-County |
| MT633046.1 | 03/26/2020 | USA:Washington-Yakima-County |
| MT641655.1 | 03/26/2020 | Australia:Northern-Territory |
| MT641674.1 | 03/26/2020 | Australia:Victoria |
| MT641699.1 | 03/26/2020 | Australia:Victoria |
| MT641715.1 | 03/26/2020 | Australia:Victoria |
| MT646079.1 | 03/26/2020 | USA |
| MT293174.1 | 03/27/2020 | USA:WA |
| MT295465.1 | 03/27/2020 | USA |
| MT334568.1 | 03/27/2020 | USA:UT |
| MT334569.1 | 03/27/2020 | USA:UT |
| MT334572.1 | 03/27/2020 | USA:UT |
| MT345871.1 | 03/27/2020 | USA:ID |
| MT345872.1 | 03/27/2020 | USA:ID |
| MT345877.1 | 03/27/2020 | USA:ID |
| MT345886.1 | 03/27/2020 | USA:ID |
| MT358678.1 | 03/27/2020 | USA:WA |
| MT358691.1 | 03/27/2020 | USA:WA |
| MT358702.1 | 03/27/2020 | USA:WA |
| MT358708.1 | 03/27/2020 | USA:WA |
| MT358710.1 | 03/27/2020 | USA:WA |
| MT358711.1 | 03/27/2020 | USA:WA |

| | | |
|---|---|---|
| MT358712.1 | 03/27/2020 | USA:WA |
| MT358713.1 | 03/27/2020 | USA:WA |
| MT358714.1 | 03/27/2020 | USA:WA |
| MT358715.1 | 03/27/2020 | USA:WA |
| MT358716.1 | 03/27/2020 | USA:WA |
| MT358718.1 | 03/27/2020 | USA:WA |
| MT358719.1 | 03/27/2020 | USA:WA |
| MT358720.1 | 03/27/2020 | USA:WA |
| MT358721.1 | 03/27/2020 | USA:WA |
| MT358722.1 | 03/27/2020 | USA:WA |
| MT358723.1 | 03/27/2020 | USA:WA |
| MT358724.1 | 03/27/2020 | USA:WA |
| MT358725.1 | 03/27/2020 | USA:WA |
| MT358726.1 | 03/27/2020 | USA:WA |
| MT375448.1 | 03/27/2020 | USA:WA |
| MT375451.1 | 03/27/2020 | USA:WA |
| MT375453.1 | 03/27/2020 | USA:WA |
| MT375454.1 | 03/27/2020 | USA:WA |
| MT375456.1 | 03/27/2020 | USA:WA |
| MT412209.1 | 03/27/2020 | USA:Michigan |
| MT412210.1 | 03/27/2020 | USA:Michigan |
| MT412211.1 | 03/27/2020 | USA:Michigan |
| MT412214.1 | 03/27/2020 | USA:Michigan |
| MT412304.1 | 03/27/2020 | USA:ID |
| MT422806.1 | 03/27/2020 | USA:FL |
| MT422807.1 | 03/27/2020 | USA:FL |
| MT439304.1 | 03/27/2020 | USA:MI |
| MT439306.1 | 03/27/2020 | USA:MI |
| MT439307.1 | 03/27/2020 | USA:MI |
| MT439308.1 | 03/27/2020 | USA:MI |
| MT439310.1 | 03/27/2020 | USA:MI |
| MT439311.1 | 03/27/2020 | USA:MI |
| MT439312.1 | 03/27/2020 | USA:MI |
| MT439313.1 | 03/27/2020 | USA:MI |
| MT439314.1 | 03/27/2020 | USA:MI |
| MT439315.1 | 03/27/2020 | USA:MI |
| MT439316.1 | 03/27/2020 | USA:MI |
| MT439318.1 | 03/27/2020 | USA:MI |
| MT444564.1 | 03/27/2020 | USA |
| MT451312.1 | 03/27/2020 | Australia:Victoria |
| MT451313.1 | 03/27/2020 | Australia:Victoria |
| MT451314.1 | 03/27/2020 | Australia:Victoria |
| MT451315.1 | 03/27/2020 | Australia:Victoria |
| MT451326.1 | 03/27/2020 | Australia:Victoria |
| MT451327.1 | 03/27/2020 | Australia:Victoria |
| MT451330.1 | 03/27/2020 | Australia:Victoria |

| | | |
|---|---|---|
| MT451331.1 | 03/27/2020 | Australia:Victoria |
| MT451332.1 | 03/27/2020 | Australia:Victoria |
| MT451414.1 | 03/27/2020 | Australia:Victoria |
| MT451415.1 | 03/27/2020 | Australia:Victoria |
| MT451416.1 | 03/27/2020 | Australia:Victoria |
| MT451417.1 | 03/27/2020 | Australia:Victoria |
| MT451419.1 | 03/27/2020 | Australia:Victoria |
| MT451437.1 | 03/27/2020 | Australia:Victoria |
| MT451459.1 | 03/27/2020 | Australia:Victoria |
| MT451461.1 | 03/27/2020 | Australia:Victoria |
| MT451462.1 | 03/27/2020 | Australia:Victoria |
| MT451463.1 | 03/27/2020 | Australia:Victoria |
| MT451464.1 | 03/27/2020 | Australia:Victoria |
| MT451465.1 | 03/27/2020 | Australia:Victoria |
| MT451466.1 | 03/27/2020 | Australia:Victoria |
| MT451467.1 | 03/27/2020 | Australia:Victoria |
| MT451468.1 | 03/27/2020 | Australia:Victoria |
| MT451469.1 | 03/27/2020 | Australia:Victoria |
| MT451470.1 | 03/27/2020 | Australia:Victoria |
| MT451472.1 | 03/27/2020 | Australia:Victoria |
| MT451473.1 | 03/27/2020 | Australia:Victoria |
| MT451474.1 | 03/27/2020 | Australia:Victoria |
| MT451475.1 | 03/27/2020 | Australia:Victoria |
| MT451484.1 | 03/27/2020 | Australia:Victoria |
| MT451489.1 | 03/27/2020 | Australia:Victoria |
| MT451498.1 | 03/27/2020 | Australia:Victoria |
| MT451499.1 | 03/27/2020 | Australia:Victoria |
| MT451501.1 | 03/27/2020 | Australia:Victoria |
| MT451631.1 | 03/27/2020 | Australia:Victoria |
| MT451649.1 | 03/27/2020 | Australia:Victoria |
| MT451650.1 | 03/27/2020 | Australia:Victoria |
| MT451651.1 | 03/27/2020 | Australia:Victoria |
| MT451652.1 | 03/27/2020 | Australia:Victoria |
| MT451653.1 | 03/27/2020 | Australia:Victoria |
| MT451654.1 | 03/27/2020 | Australia:Victoria |
| MT451655.1 | 03/27/2020 | Australia:Victoria |
| MT451656.1 | 03/27/2020 | Australia:Victoria |
| MT451657.1 | 03/27/2020 | Australia:Victoria |
| MT451658.1 | 03/27/2020 | Australia:Victoria |
| MT451659.1 | 03/27/2020 | Australia:Victoria |
| MT451660.1 | 03/27/2020 | Australia:Victoria |
| MT451661.1 | 03/27/2020 | Australia:Victoria |
| MT451662.1 | 03/27/2020 | Australia:Victoria |
| MT459923.1 | 03/27/2020 | Greece:Athens |
| MT459988.1 | 03/27/2020 | USA:SC |
| MT459989.1 | 03/27/2020 | USA:SC |

| | | |
|---|---|---|
| MT474128.1 | 03/27/2020 | USA:CA |
| MT477848.1 | 03/27/2020 | USA:AK |
| MT481927.1 | 03/27/2020 | USA:CA |
| MT499182.1 | 03/27/2020 | USA:CA |
| MT499184.1 | 03/27/2020 | USA:CA |
| MT499219.1 | 03/27/2020 | Tunisia |
| MT506228.1 | 03/27/2020 | USA:Michigan |
| MT506546.1 | 03/27/2020 | USA:Michigan |
| MT506550.1 | 03/27/2020 | USA:Michigan |
| MT506551.1 | 03/27/2020 | USA:Michigan |
| MT506553.1 | 03/27/2020 | USA:Michigan |
| MT506554.1 | 03/27/2020 | USA:Michigan |
| MT506555.1 | 03/27/2020 | USA:Michigan |
| MT506556.1 | 03/27/2020 | USA:Michigan |
| MT506558.1 | 03/27/2020 | USA:Michigan |
| MT506559.1 | 03/27/2020 | USA:Michigan |
| MT509483.1 | 03/27/2020 | USA |
| MT509687.1 | 03/27/2020 | USA:Michigan |
| MT517420.1 | 03/27/2020 | Czech-Republic |
| MT517421.1 | 03/27/2020 | Czech-Republic |
| MT517429.1 | 03/27/2020 | Czech-Republic |
| MT517430.1 | 03/27/2020 | Czech-Republic |
| MT517431.1 | 03/27/2020 | Czech-Republic |
| MT517432.1 | 03/27/2020 | Czech-Republic |
| MT520183.1 | 03/27/2020 | USA:Massachusetts |
| MT520185.1 | 03/27/2020 | USA:Massachusetts |
| MT520217.1 | 03/27/2020 | USA:Massachusetts |
| MT520231.1 | 03/27/2020 | USA:Massachusetts |
| MT520260.1 | 03/27/2020 | USA:Massachusetts |
| MT520279.1 | 03/27/2020 | USA:Massachusetts |
| MT520304.1 | 03/27/2020 | USA:Massachusetts |
| MT520317.1 | 03/27/2020 | USA:Massachusetts |
| MT520322.1 | 03/27/2020 | USA:Massachusetts |
| MT520333.1 | 03/27/2020 | USA:Massachusetts |
| MT520358.1 | 03/27/2020 | USA:Massachusetts |
| MT520361.1 | 03/27/2020 | USA:Massachusetts |
| MT520363.1 | 03/27/2020 | USA:Massachusetts |
| MT520381.1 | 03/27/2020 | USA:Massachusetts |
| MT520397.1 | 03/27/2020 | USA:Massachusetts |
| MT520404.1 | 03/27/2020 | USA:Massachusetts |
| MT520412.1 | 03/27/2020 | USA:Massachusetts |
| MT520418.1 | 03/27/2020 | USA:Massachusetts |
| MT520437.1 | 03/27/2020 | USA:Massachusetts |
| MT520475.1 | 03/27/2020 | USA:Massachusetts |
| MT520485.1 | 03/27/2020 | USA:Massachusetts |
| MT520500.1 | 03/27/2020 | USA:Massachusetts |

| | | |
|---|---|---|
| MT520502.1 | 03/27/2020 | USA:Massachusetts |
| MT520512.1 | 03/27/2020 | USA:Massachusetts |
| MT520520.1 | 03/27/2020 | USA:Massachusetts |
| MT520534.1 | 03/27/2020 | USA:Massachusetts |
| MT520543.1 | 03/27/2020 | USA:Massachusetts |
| MT520546.1 | 03/27/2020 | USA:Massachusetts |
| MT533233.1 | 03/27/2020 | USA:CA |
| MT535476.1 | 03/27/2020 | USA |
| MT535479.1 | 03/27/2020 | USA |
| MT535480.1 | 03/27/2020 | USA |
| MT628174.1 | 03/27/2020 | USA:CA |
| MT628175.1 | 03/27/2020 | USA:CA |
| MT628248.1 | 03/27/2020 | USA:CA |
| MT628249.1 | 03/27/2020 | USA:CA |
| MT632782.1 | 03/27/2020 | USA:Washington-Skagit-County |
| MT632787.1 | 03/27/2020 | USA:Washington-Skagit-County |
| MT632798.1 | 03/27/2020 | USA:Washington-Adams-County |
| MT632801.1 | 03/27/2020 | USA:Washington-Skagit-County |
| MT632803.1 | 03/27/2020 | USA:Washington-Adams-County |
| MT632807.1 | 03/27/2020 | USA:Washington-Skagit-County |
| MT632818.1 | 03/27/2020 | USA:Washington-Skagit-County |
| MT632819.1 | 03/27/2020 | USA:Washington-Skagit-County |
| MT632838.1 | 03/27/2020 | USA:Washington-Skagit-County |
| MT632848.1 | 03/27/2020 | USA:Washington-Skagit-County |
| MT632888.1 | 03/27/2020 | USA:Washington-Yakima-County |
| MT632889.1 | 03/27/2020 | USA:Washington-Snohomish-County |
| MT632942.1 | 03/27/2020 | USA:Washington-Franklin-County |
| MT641656.1 | 03/27/2020 | Australia:Northern-Territory |
| MT641672.1 | 03/27/2020 | Australia:Victoria |
| MT641673.1 | 03/27/2020 | Australia:Victoria |
| MT641716.1 | 03/27/2020 | Australia:Victoria |
| MT646051.1 | 03/27/2020 | USA |
| MT293185.1 | 03/28/2020 | USA:WA |
| MT293188.1 | 03/28/2020 | USA:WA |
| MT293192.1 | 03/28/2020 | USA:WA |
| MT293195.1 | 03/28/2020 | USA:WA |
| MT345875.1 | 03/28/2020 | USA:ID |
| MT345876.1 | 03/28/2020 | USA:ID |
| MT345879.1 | 03/28/2020 | USA:ID |
| MT345880.1 | 03/28/2020 | USA:ID |
| MT358735.1 | 03/28/2020 | USA:WA |
| MT358737.1 | 03/28/2020 | USA:WA |
| MT358738.1 | 03/28/2020 | USA:WA |
| MT358740.1 | 03/28/2020 | USA:WA |
| MT375446.1 | 03/28/2020 | USA:WA |
| MT375449.1 | 03/28/2020 | USA:WA |

| | | |
|---|---|---|
| MT375450.1 | 03/28/2020 | USA:WA |
| MT375452.1 | 03/28/2020 | USA:WA |
| MT375455.1 | 03/28/2020 | USA:WA |
| MT380733.1 | 03/28/2020 | USA |
| MT385414.1 | 03/28/2020 | USA:CA |
| MT385415.1 | 03/28/2020 | USA:CA |
| MT385420.1 | 03/28/2020 | USA:CA |
| MT385422.1 | 03/28/2020 | USA:CA |
| MT385423.1 | 03/28/2020 | USA:CA |
| MT385424.1 | 03/28/2020 | USA:CA |
| MT385426.1 | 03/28/2020 | USA:CA |
| MT385428.1 | 03/28/2020 | USA:CA |
| MT385429.1 | 03/28/2020 | USA:CA |
| MT385430.1 | 03/28/2020 | USA:CA |
| MT385432.1 | 03/28/2020 | USA:CA |
| MT385434.1 | 03/28/2020 | USA:CA |
| MT385437.1 | 03/28/2020 | USA:CA |
| MT385439.1 | 03/28/2020 | USA:CA |
| MT385443.1 | 03/28/2020 | USA:CA |
| MT385453.1 | 03/28/2020 | USA:CA |
| MT385456.1 | 03/28/2020 | USA:CA |
| MT385457.1 | 03/28/2020 | USA:CA |
| MT385460.1 | 03/28/2020 | USA:CA |
| MT385465.1 | 03/28/2020 | USA:CA |
| MT385473.1 | 03/28/2020 | USA:CA |
| MT385474.1 | 03/28/2020 | USA:CA |
| MT385480.1 | 03/28/2020 | USA:CA |
| MT385481.1 | 03/28/2020 | USA:CA |
| MT385482.1 | 03/28/2020 | USA:CA |
| MT385483.1 | 03/28/2020 | USA:CA |
| MT385489.1 | 03/28/2020 | USA:CA |
| MT385490.1 | 03/28/2020 | USA:CA |
| MT385495.1 | 03/28/2020 | USA:CA |
| MT412213.1 | 03/28/2020 | USA:Michigan |
| MT412215.1 | 03/28/2020 | USA:Illinois |
| MT412309.1 | 03/28/2020 | USA:ID |
| MT439309.1 | 03/28/2020 | USA:MI |
| MT439319.1 | 03/28/2020 | USA:MI |
| MT444565.1 | 03/28/2020 | USA |
| MT444566.1 | 03/28/2020 | USA |
| MT444568.1 | 03/28/2020 | USA |
| MT451334.1 | 03/28/2020 | Australia:Victoria |
| MT451335.1 | 03/28/2020 | Australia:Victoria |
| MT451336.1 | 03/28/2020 | Australia:Victoria |
| MT451337.1 | 03/28/2020 | Australia:Victoria |
| MT451338.1 | 03/28/2020 | Australia:Victoria |

| | | |
|---|---|---|
| MT451339.1 | 03/28/2020 | Australia:Victoria |
| MT451426.1 | 03/28/2020 | Australia:Victoria |
| MT451427.1 | 03/28/2020 | Australia:Victoria |
| MT451428.1 | 03/28/2020 | Australia:Victoria |
| MT451430.1 | 03/28/2020 | Australia:Victoria |
| MT451431.1 | 03/28/2020 | Australia:Victoria |
| MT451432.1 | 03/28/2020 | Australia:Victoria |
| MT451438.1 | 03/28/2020 | Australia:Victoria |
| MT451439.1 | 03/28/2020 | Australia:Victoria |
| MT451440.1 | 03/28/2020 | Australia:Victoria |
| MT451446.1 | 03/28/2020 | Australia:Victoria |
| MT451447.1 | 03/28/2020 | Australia:Victoria |
| MT451449.1 | 03/28/2020 | Australia:Victoria |
| MT451450.1 | 03/28/2020 | Australia:Victoria |
| MT451451.1 | 03/28/2020 | Australia:Victoria |
| MT451452.1 | 03/28/2020 | Australia:Victoria |
| MT451453.1 | 03/28/2020 | Australia:Victoria |
| MT451454.1 | 03/28/2020 | Australia:Victoria |
| MT451455.1 | 03/28/2020 | Australia:Victoria |
| MT451456.1 | 03/28/2020 | Australia:Victoria |
| MT451457.1 | 03/28/2020 | Australia:Victoria |
| MT451458.1 | 03/28/2020 | Australia:Victoria |
| MT451460.1 | 03/28/2020 | Australia:Victoria |
| MT451481.1 | 03/28/2020 | Australia:Victoria |
| MT451485.1 | 03/28/2020 | Australia:Victoria |
| MT451486.1 | 03/28/2020 | Australia:Victoria |
| MT451487.1 | 03/28/2020 | Australia:Victoria |
| MT451488.1 | 03/28/2020 | Australia:Victoria |
| MT451500.1 | 03/28/2020 | Australia:Victoria |
| MT451570.1 | 03/28/2020 | Australia:Victoria |
| MT451571.1 | 03/28/2020 | Australia:Victoria |
| MT451663.1 | 03/28/2020 | Australia:Victoria |
| MT451664.1 | 03/28/2020 | Australia:Victoria |
| MT451665.1 | 03/28/2020 | Australia:Victoria |
| MT459838.1 | 03/28/2020 | Greece:Athens |
| MT459904.1 | 03/28/2020 | Greece:Athens |
| MT461605.1 | 03/28/2020 | USA:WA |
| MT461606.1 | 03/28/2020 | USA:WA |
| MT461612.1 | 03/28/2020 | USA:WA |
| MT474126.1 | 03/28/2020 | USA:CA |
| MT477847.1 | 03/28/2020 | USA:AK |
| MT499216.1 | 03/28/2020 | Tunisia |
| MT506557.1 | 03/28/2020 | USA:Michigan |
| MT506560.1 | 03/28/2020 | USA:Michigan |
| MT506561.1 | 03/28/2020 | USA:Michigan |
| MT506562.1 | 03/28/2020 | USA:Michigan |

| | | |
|---|---|---|
| MT506563.1 | 03/28/2020 | USA:Michigan |
| MT506564.1 | 03/28/2020 | USA:Michigan |
| MT506691.1 | 03/28/2020 | USA:Michigan |
| MT509689.1 | 03/28/2020 | USA:Michigan |
| MT517423.1 | 03/28/2020 | Czech-Republic |
| MT517433.1 | 03/28/2020 | Czech-Republic |
| MT520190.1 | 03/28/2020 | USA:Massachusetts |
| MT520204.1 | 03/28/2020 | USA:Massachusetts |
| MT520209.1 | 03/28/2020 | USA:Massachusetts |
| MT520212.1 | 03/28/2020 | USA:Massachusetts |
| MT520236.1 | 03/28/2020 | USA:Massachusetts |
| MT520238.1 | 03/28/2020 | USA:Massachusetts |
| MT520267.1 | 03/28/2020 | USA:Massachusetts |
| MT520269.1 | 03/28/2020 | USA:Massachusetts |
| MT520280.1 | 03/28/2020 | USA:Massachusetts |
| MT520345.1 | 03/28/2020 | USA:Massachusetts |
| MT520348.1 | 03/28/2020 | USA:Massachusetts |
| MT520351.1 | 03/28/2020 | USA:Massachusetts |
| MT520354.1 | 03/28/2020 | USA:Massachusetts |
| MT520364.1 | 03/28/2020 | USA:Massachusetts |
| MT520370.1 | 03/28/2020 | USA:Massachusetts |
| MT520389.1 | 03/28/2020 | USA:Massachusetts |
| MT520392.1 | 03/28/2020 | USA:Massachusetts |
| MT520415.1 | 03/28/2020 | USA:Massachusetts |
| MT520449.1 | 03/28/2020 | USA:Massachusetts |
| MT520451.1 | 03/28/2020 | USA:Massachusetts |
| MT520457.1 | 03/28/2020 | USA:Massachusetts |
| MT520468.1 | 03/28/2020 | USA:Massachusetts |
| MT520471.1 | 03/28/2020 | USA:Massachusetts |
| MT520501.1 | 03/28/2020 | USA:Massachusetts |
| MT533234.1 | 03/28/2020 | USA:CA |
| MT628145.1 | 03/28/2020 | USA:CA |
| MT628146.1 | 03/28/2020 | USA:CA |
| MT628246.1 | 03/28/2020 | USA:CA |
| MT628247.1 | 03/28/2020 | USA:CA |
| MT632839.1 | 03/28/2020 | USA:Washington-Skagit-County |
| MT632846.1 | 03/28/2020 | USA:Washington-Pierce-County |
| MT632875.1 | 03/28/2020 | USA:Washington-Yakima-County |
| MT632883.1 | 03/28/2020 | USA:Washington-Yakima-County |
| MT632898.1 | 03/28/2020 | USA:Washington |
| MT632917.1 | 03/28/2020 | USA:Washington-Yakima-County |
| MT632922.1 | 03/28/2020 | USA:Washington-Yakima-County |
| MT632928.1 | 03/28/2020 | USA:Washington-Yakima-County |
| MT632934.1 | 03/28/2020 | USA:Washington-Yakima-County |
| MT632939.1 | 03/28/2020 | USA:Washington-Yakima-County |
| MT632945.1 | 03/28/2020 | USA:Washington-Snohomish-County |

| | | |
|---|---|---|
| MT632948.1 | 03/28/2020 | USA:Washington-Yakima-County |
| MT632954.1 | 03/28/2020 | USA:Washington-Yakima-County |
| MT632963.1 | 03/28/2020 | USA:Washington-Yakima-County |
| MT632967.1 | 03/28/2020 | USA:Washington-Snohomish-County |
| MT632970.1 | 03/28/2020 | USA:Washington-Yakima-County |
| MT632972.1 | 03/28/2020 | USA:Washington-Yakima-County |
| MT632978.1 | 03/28/2020 | USA:Washington-Yakima-County |
| MT632995.1 | 03/28/2020 | USA:Washington-Yakima-County |
| MT632997.1 | 03/28/2020 | USA:Washington-Yakima-County |
| MT632999.1 | 03/28/2020 | USA:Washington-Yakima-County |
| MT633003.1 | 03/28/2020 | USA:Washington-Yakima-County |
| MT633009.1 | 03/28/2020 | USA:Washington-Yakima-County |
| MT633019.1 | 03/28/2020 | USA:Washington-Yakima-County |
| MT633024.1 | 03/28/2020 | USA:Washington-Yakima-County |
| MT633025.1 | 03/28/2020 | USA:Washington-Yakima-County |
| MT633031.1 | 03/28/2020 | USA:Washington-Yakima-County |
| MT633041.1 | 03/28/2020 | USA:Washington-Yakima-County |
| MT641703.1 | 03/28/2020 | Australia:Victoria |
| MT646080.1 | 03/28/2020 | USA |
| MT646101.1 | 03/28/2020 | USA |
| MT293157.1 | 03/29/2020 | USA:WA |
| MT293163.1 | 03/29/2020 | USA:WA |
| MT345878.1 | 03/29/2020 | USA:OR |
| MT345884.1 | 03/29/2020 | USA:ID |
| MT345887.1 | 03/29/2020 | USA:ID |
| MT345888.1 | 03/29/2020 | USA:ID |
| MT358669.1 | 03/29/2020 | USA:WA |
| MT358704.1 | 03/29/2020 | USA:ID |
| MT358705.1 | 03/29/2020 | USA:ID |
| MT358730.1 | 03/29/2020 | USA:WA |
| MT358731.1 | 03/29/2020 | USA:WA |
| MT358732.1 | 03/29/2020 | USA:WA |
| MT358733.1 | 03/29/2020 | USA:WA |
| MT358734.1 | 03/29/2020 | USA:WA |
| MT358736.1 | 03/29/2020 | USA:WA |
| MT358739.1 | 03/29/2020 | USA:WA |
| MT358741.1 | 03/29/2020 | USA:WA |
| MT358742.1 | 03/29/2020 | USA:WA |
| MT358745.1 | 03/29/2020 | USA:ID |
| MT375428.1 | 03/29/2020 | USA:WA |
| MT375429.1 | 03/29/2020 | USA:ID |
| MT375431.1 | 03/29/2020 | USA:ID |
| MT375444.1 | 03/29/2020 | USA:WA |
| MT375445.1 | 03/29/2020 | USA:WA |
| MT380728.1 | 03/29/2020 | USA |
| MT412305.1 | 03/29/2020 | USA:ID |

| | | |
|---|---|---|
| MT412306.1 | 03/29/2020 | USA:ID |
| MT412307.1 | 03/29/2020 | USA:OR |
| MT444567.1 | 03/29/2020 | USA |
| MT451340.1 | 03/29/2020 | Australia:Victoria |
| MT451341.1 | 03/29/2020 | Australia:Victoria |
| MT451342.1 | 03/29/2020 | Australia:Victoria |
| MT451343.1 | 03/29/2020 | Australia:Victoria |
| MT451344.1 | 03/29/2020 | Australia:Victoria |
| MT451345.1 | 03/29/2020 | Australia:Victoria |
| MT451346.1 | 03/29/2020 | Australia:Victoria |
| MT451347.1 | 03/29/2020 | Australia:Victoria |
| MT451429.1 | 03/29/2020 | Australia:Victoria |
| MT451434.1 | 03/29/2020 | Australia:Victoria |
| MT451441.1 | 03/29/2020 | Australia:Victoria |
| MT451442.1 | 03/29/2020 | Australia:Victoria |
| MT451443.1 | 03/29/2020 | Australia:Victoria |
| MT451478.1 | 03/29/2020 | Australia:Victoria |
| MT451480.1 | 03/29/2020 | Australia:Victoria |
| MT451482.1 | 03/29/2020 | Australia:Victoria |
| MT451483.1 | 03/29/2020 | Australia:Victoria |
| MT451493.1 | 03/29/2020 | Australia:Victoria |
| MT451494.1 | 03/29/2020 | Australia:Victoria |
| MT451495.1 | 03/29/2020 | Australia:Victoria |
| MT451496.1 | 03/29/2020 | Australia:Victoria |
| MT451497.1 | 03/29/2020 | Australia:Victoria |
| MT451502.1 | 03/29/2020 | Australia:Victoria |
| MT451503.1 | 03/29/2020 | Australia:Victoria |
| MT451504.1 | 03/29/2020 | Australia:Victoria |
| MT451516.1 | 03/29/2020 | Australia:Victoria |
| MT451517.1 | 03/29/2020 | Australia:Victoria |
| MT451518.1 | 03/29/2020 | Australia:Victoria |
| MT451519.1 | 03/29/2020 | Australia:Victoria |
| MT451528.1 | 03/29/2020 | Australia:Victoria |
| MT451529.1 | 03/29/2020 | Australia:Victoria |
| MT451531.1 | 03/29/2020 | Australia:Victoria |
| MT451532.1 | 03/29/2020 | Australia:Victoria |
| MT451666.1 | 03/29/2020 | Australia:Victoria |
| MT459837.1 | 03/29/2020 | Greece:Athens |
| MT459841.1 | 03/29/2020 | Greece:Athens |
| MT459896.1 | 03/29/2020 | Greece:Athens |
| MT461603.1 | 03/29/2020 | USA |
| MT461604.1 | 03/29/2020 | USA:WA |
| MT461607.1 | 03/29/2020 | USA:WA |
| MT461608.1 | 03/29/2020 | USA:WA |
| MT461609.1 | 03/29/2020 | USA:WA |
| MT461610.1 | 03/29/2020 | USA:WA |

| | | |
|---|---|---|
| MT461611.1 | 03/29/2020 | USA:WA |
| MT477849.1 | 03/29/2020 | USA:AK |
| MT499173.1 | 03/29/2020 | USA:CA |
| MT499174.1 | 03/29/2020 | USA:CA |
| MT506692.1 | 03/29/2020 | USA:Michigan |
| MT509691.1 | 03/29/2020 | USA:Michigan |
| MT509694.1 | 03/29/2020 | USA:Michigan |
| MT509695.1 | 03/29/2020 | USA:Michigan |
| MT511077.1 | 03/29/2020 | Poland |
| MT511079.1 | 03/29/2020 | Poland |
| MT511080.1 | 03/29/2020 | Poland |
| MT517434.1 | 03/29/2020 | Czech-Republic |
| MT520344.1 | 03/29/2020 | USA:Massachusetts |
| MT628220.1 | 03/29/2020 | USA:CA |
| MT628250.1 | 03/29/2020 | USA:CA |
| MT632811.1 | 03/29/2020 | USA:Washington |
| MT632821.1 | 03/29/2020 | USA:Washington-Pierce-County |
| MT632822.1 | 03/29/2020 | USA:Washington |
| MT632833.1 | 03/29/2020 | USA:Washington-Pierce-County |
| MT632841.1 | 03/29/2020 | USA:Washington-Yakima-County |
| MT632849.1 | 03/29/2020 | USA:Washington-Skagit-County |
| MT632904.1 | 03/29/2020 | USA:Washington-Yakima-County |
| MT632924.1 | 03/29/2020 | USA:Washington |
| MT633000.1 | 03/29/2020 | USA:Washington-Snohomish-County |
| MT633033.1 | 03/29/2020 | USA:Washington-Yakima-County |
| MT646081.1 | 03/29/2020 | USA |
| MT646112.1 | 03/29/2020 | USA |
| MT293156.1 | 03/30/2020 | USA:WA |
| MT293173.1 | 03/30/2020 | USA:WA |
| MT293175.1 | 03/30/2020 | USA:WA |
| MT334573.1 | 03/30/2020 | USA:UT |
| MT345868.1 | 03/30/2020 | USA:CT |
| MT345869.1 | 03/30/2020 | USA:CT |
| MT345870.1 | 03/30/2020 | USA:CT |
| MT345873.1 | 03/30/2020 | USA:CT |
| MT345874.1 | 03/30/2020 | USA:CT |
| MT345881.1 | 03/30/2020 | USA:WA |
| MT345885.1 | 03/30/2020 | USA:ID |
| MT358651.1 | 03/30/2020 | USA:CT |
| MT358652.1 | 03/30/2020 | USA:ID |
| MT358655.1 | 03/30/2020 | USA:ID |
| MT358667.1 | 03/30/2020 | USA:WA |
| MT358671.1 | 03/30/2020 | USA:WA |
| MT358672.1 | 03/30/2020 | USA:WA |
| MT358673.1 | 03/30/2020 | USA:CT |
| MT358727.1 | 03/30/2020 | USA:WA |

| | | |
|---|---|---|
| MT375432.1 | 03/30/2020 | USA:WA |
| MT375433.1 | 03/30/2020 | USA:WA |
| MT375439.1 | 03/30/2020 | USA:ID |
| MT412217.1 | 03/30/2020 | USA:Michigan |
| MT412218.1 | 03/30/2020 | USA:Michigan |
| MT412219.1 | 03/30/2020 | USA:Michigan |
| MT412220.1 | 03/30/2020 | USA:Michigan |
| MT412221.1 | 03/30/2020 | USA:Michigan |
| MT412222.1 | 03/30/2020 | USA:Michigan |
| MT412223.1 | 03/30/2020 | USA:Michigan |
| MT412224.1 | 03/30/2020 | USA:Michigan |
| MT412297.1 | 03/30/2020 | USA:ID |
| MT412300.1 | 03/30/2020 | USA:WA |
| MT412301.1 | 03/30/2020 | USA:CT |
| MT412302.1 | 03/30/2020 | USA:CT |
| MT412303.1 | 03/30/2020 | USA:CT |
| MT439320.1 | 03/30/2020 | USA:MI |
| MT444534.1 | 03/30/2020 | USA |
| MT444536.1 | 03/30/2020 | USA |
| MT444537.1 | 03/30/2020 | USA |
| MT444538.1 | 03/30/2020 | USA |
| MT444539.1 | 03/30/2020 | USA |
| MT451348.1 | 03/30/2020 | Australia:Victoria |
| MT451349.1 | 03/30/2020 | Australia:Victoria |
| MT451491.1 | 03/30/2020 | Australia:Victoria |
| MT451492.1 | 03/30/2020 | Australia:Victoria |
| MT451505.1 | 03/30/2020 | Australia:Victoria |
| MT451506.1 | 03/30/2020 | Australia:Victoria |
| MT451507.1 | 03/30/2020 | Australia:Victoria |
| MT451508.1 | 03/30/2020 | Australia:Victoria |
| MT451509.1 | 03/30/2020 | Australia:Victoria |
| MT451510.1 | 03/30/2020 | Australia:Victoria |
| MT451511.1 | 03/30/2020 | Australia:Victoria |
| MT451512.1 | 03/30/2020 | Australia:Victoria |
| MT451513.1 | 03/30/2020 | Australia:Victoria |
| MT451514.1 | 03/30/2020 | Australia:Victoria |
| MT451515.1 | 03/30/2020 | Australia:Victoria |
| MT451521.1 | 03/30/2020 | Australia:Victoria |
| MT451523.1 | 03/30/2020 | Australia:Victoria |
| MT451524.1 | 03/30/2020 | Australia:Victoria |
| MT451525.1 | 03/30/2020 | Australia:Victoria |
| MT451526.1 | 03/30/2020 | Australia:Victoria |
| MT451530.1 | 03/30/2020 | Australia:Victoria |
| MT451542.1 | 03/30/2020 | Australia:Victoria |
| MT451544.1 | 03/30/2020 | Australia:Victoria |
| MT451545.1 | 03/30/2020 | Australia:Victoria |

| | | |
|---|---|---|
| MT451546.1 | 03/30/2020 | Australia:Victoria |
| MT451547.1 | 03/30/2020 | Australia:Victoria |
| MT451548.1 | 03/30/2020 | Australia:Victoria |
| MT451549.1 | 03/30/2020 | Australia:Victoria |
| MT451550.1 | 03/30/2020 | Australia:Victoria |
| MT451551.1 | 03/30/2020 | Australia:Victoria |
| MT451632.1 | 03/30/2020 | Australia:Victoria |
| MT451633.1 | 03/30/2020 | Australia:Victoria |
| MT451634.1 | 03/30/2020 | Australia:Victoria |
| MT451635.1 | 03/30/2020 | Australia:Victoria |
| MT451636.1 | 03/30/2020 | Australia:Victoria |
| MT451667.1 | 03/30/2020 | Australia:Victoria |
| MT451668.1 | 03/30/2020 | Australia:Victoria |
| MT451669.1 | 03/30/2020 | Australia:Victoria |
| MT451670.1 | 03/30/2020 | Australia:Victoria |
| MT451671.1 | 03/30/2020 | Australia:Victoria |
| MT451793.1 | 03/30/2020 | Australia:Victoria |
| MT461624.1 | 03/30/2020 | USA:CT |
| MT461625.1 | 03/30/2020 | USA:ID |
| MT461628.1 | 03/30/2020 | USA:ID |
| MT474136.1 | 03/30/2020 | USA:CA |
| MT499171.1 | 03/30/2020 | USA:CA |
| MT499172.1 | 03/30/2020 | USA:CA |
| MT499217.1 | 03/30/2020 | Tunisia |
| MT506694.1 | 03/30/2020 | USA:Michigan |
| MT509481.1 | 03/30/2020 | USA |
| MT509690.1 | 03/30/2020 | USA:Michigan |
| MT509692.1 | 03/30/2020 | USA:Michigan |
| MT509693.1 | 03/30/2020 | USA:Michigan |
| MT509696.1 | 03/30/2020 | USA:Michigan |
| MT509697.1 | 03/30/2020 | USA:Michigan |
| MT509698.1 | 03/30/2020 | USA:Michigan |
| MT509699.1 | 03/30/2020 | USA:Michigan |
| MT509700.1 | 03/30/2020 | USA:Michigan |
| MT509701.1 | 03/30/2020 | USA:Michigan |
| MT509702.1 | 03/30/2020 | USA:Michigan |
| MT509703.1 | 03/30/2020 | USA:Michigan |
| MT509704.1 | 03/30/2020 | USA:Michigan |
| MT532557.1 | 03/30/2020 | USA |
| MT535481.1 | 03/30/2020 | USA |
| MT612284.1 | 03/30/2020 | Timor-Leste |
| MT627628.1 | 03/30/2020 | USA:Washington |
| MT628147.1 | 03/30/2020 | USA:CA |
| MT628148.1 | 03/30/2020 | USA:CA |
| MT628177.1 | 03/30/2020 | USA:CA |
| MT628180.1 | 03/30/2020 | USA:CA |

| | | |
|---|---|---|
| MT628221.1 | 03/30/2020 | USA:CA |
| MT631807.1 | 03/30/2020 | USA:CA |
| MT631808.1 | 03/30/2020 | USA:CA |
| MT631809.1 | 03/30/2020 | USA:CA |
| MT632775.1 | 03/30/2020 | USA:Washington-King-County |
| MT632812.1 | 03/30/2020 | USA:Washington |
| MT632815.1 | 03/30/2020 | USA:Washington-Snohomish-County |
| MT632851.1 | 03/30/2020 | USA:Washington-Yakima-County |
| MT632861.1 | 03/30/2020 | USA:Washington-Mason-County |
| MT632871.1 | 03/30/2020 | USA:Washington-Yakima-County |
| MT632900.1 | 03/30/2020 | USA:Washington-Adams-County |
| MT632907.1 | 03/30/2020 | USA:Washington-Yakima-County |
| MT632921.1 | 03/30/2020 | USA:Washington-Yakima-County |
| MT632925.1 | 03/30/2020 | USA:Washington-King-County |
| MT632933.1 | 03/30/2020 | USA:Washington-King-County |
| MT632969.1 | 03/30/2020 | USA:Washington-King-County |
| MT632979.1 | 03/30/2020 | USA:Washington-Yakima-County |
| MT632982.1 | 03/30/2020 | USA:Washington-Yakima-County |
| MT633011.1 | 03/30/2020 | USA:Washington-Yakima-County |
| MT633022.1 | 03/30/2020 | USA:Washington-King-County |
| MT641778.1 | 03/30/2020 | Timor-Leste |
| MT646089.1 | 03/30/2020 | USA |
| MT646092.1 | 03/30/2020 | USA |
| MT646110.1 | 03/30/2020 | USA |
| MT646116.1 | 03/30/2020 | USA |
| MT345882.1 | 03/31/2020 | USA:WA |
| MT345883.1 | 03/31/2020 | USA:WA |
| MT358645.1 | 03/31/2020 | USA:WA |
| MT358646.1 | 03/31/2020 | USA:WA |
| MT358647.1 | 03/31/2020 | USA:WA |
| MT358648.1 | 03/31/2020 | USA:WA |
| MT358649.1 | 03/31/2020 | USA:WA |
| MT358650.1 | 03/31/2020 | USA:WA |
| MT358653.1 | 03/31/2020 | USA:CT |
| MT358654.1 | 03/31/2020 | USA:CT |
| MT358656.1 | 03/31/2020 | USA:WA |
| MT358657.1 | 03/31/2020 | USA:WA |
| MT358658.1 | 03/31/2020 | USA:WA |
| MT358659.1 | 03/31/2020 | USA:WA |
| MT358660.1 | 03/31/2020 | USA:WA |
| MT358661.1 | 03/31/2020 | USA:OR |
| MT358662.1 | 03/31/2020 | USA:WA |
| MT358663.1 | 03/31/2020 | USA:WA |
| MT358664.1 | 03/31/2020 | USA:WA |
| MT358665.1 | 03/31/2020 | USA:WA |
| MT358666.1 | 03/31/2020 | USA:WA |

| | | |
|---|---|---|
| MT358668.1 | 03/31/2020 | USA:WA |
| MT358670.1 | 03/31/2020 | USA:WA |
| MT358706.1 | 03/31/2020 | USA:WA |
| MT358707.1 | 03/31/2020 | USA:WA |
| MT358728.1 | 03/31/2020 | USA:WA |
| MT358729.1 | 03/31/2020 | USA:WA |
| MT358746.1 | 03/31/2020 | USA:WA |
| MT358747.1 | 03/31/2020 | USA:WA |
| MT358748.1 | 03/31/2020 | USA:WA |
| MT371050.1 | 03/31/2020 | Sri-Lanka |
| MT375430.1 | 03/31/2020 | USA:WA |
| MT375434.1 | 03/31/2020 | USA:WA |
| MT375435.1 | 03/31/2020 | USA:WA |
| MT375436.1 | 03/31/2020 | USA:WA |
| MT375437.1 | 03/31/2020 | USA:WA |
| MT375438.1 | 03/31/2020 | USA:WA |
| MT375440.1 | 03/31/2020 | USA:WA |
| MT380729.1 | 03/31/2020 | USA |
| MT385444.1 | 03/31/2020 | USA:CA |
| MT385447.1 | 03/31/2020 | USA:CA |
| MT385449.1 | 03/31/2020 | USA:CA |
| MT385450.1 | 03/31/2020 | USA:CA |
| MT385479.1 | 03/31/2020 | USA:CA |
| MT385484.1 | 03/31/2020 | USA:CA |
| MT385491.1 | 03/31/2020 | USA:CA |
| MT385496.1 | 03/31/2020 | USA:CA |
| MT394530.1 | 03/31/2020 | USA:CA |
| MT394531.1 | 03/31/2020 | USA:CA |
| MT412295.1 | 03/31/2020 | USA:WA |
| MT412298.1 | 03/31/2020 | USA:WA |
| MT412299.1 | 03/31/2020 | USA:ID |
| MT412308.1 | 03/31/2020 | USA:WA |
| MT444540.1 | 03/31/2020 | USA |
| MT444545.1 | 03/31/2020 | USA |
| MT444583.1 | 03/31/2020 | USA |
| MT449637.1 | 03/31/2020 | USA:CA |
| MT451520.1 | 03/31/2020 | Australia:Victoria |
| MT451527.1 | 03/31/2020 | Australia:Victoria |
| MT451543.1 | 03/31/2020 | Australia:Victoria |
| MT451552.1 | 03/31/2020 | Australia:Victoria |
| MT451553.1 | 03/31/2020 | Australia:Victoria |
| MT451554.1 | 03/31/2020 | Australia:Victoria |
| MT451555.1 | 03/31/2020 | Australia:Victoria |
| MT451556.1 | 03/31/2020 | Australia:Victoria |
| MT451557.1 | 03/31/2020 | Australia:Victoria |
| MT451558.1 | 03/31/2020 | Australia:Victoria |

| | | |
|---|---|---|
| MT451559.1 | 03/31/2020 | Australia:Victoria |
| MT451637.1 | 03/31/2020 | Australia:Victoria |
| MT451673.1 | 03/31/2020 | Australia:Victoria |
| MT451674.1 | 03/31/2020 | Australia:Victoria |
| MT451675.1 | 03/31/2020 | Australia:Victoria |
| MT451676.1 | 03/31/2020 | Australia:Victoria |
| MT451678.1 | 03/31/2020 | Australia:Victoria |
| MT451679.1 | 03/31/2020 | Australia:Victoria |
| MT451680.1 | 03/31/2020 | Australia:Victoria |
| MT451681.1 | 03/31/2020 | Australia:Victoria |
| MT451682.1 | 03/31/2020 | Australia:Victoria |
| MT451683.1 | 03/31/2020 | Australia:Victoria |
| MT451684.1 | 03/31/2020 | Australia:Victoria |
| MT451685.1 | 03/31/2020 | Australia:Victoria |
| MT451686.1 | 03/31/2020 | Australia:Victoria |
| MT451687.1 | 03/31/2020 | Australia:Victoria |
| MT451688.1 | 03/31/2020 | Australia:Victoria |
| MT451689.1 | 03/31/2020 | Australia:Victoria |
| MT451690.1 | 03/31/2020 | Australia:Victoria |
| MT451794.1 | 03/31/2020 | Australia:Victoria |
| MT451795.1 | 03/31/2020 | Australia:Victoria |
| MT451796.1 | 03/31/2020 | Australia:Victoria |
| MT451797.1 | 03/31/2020 | Australia:Victoria |
| MT459848.1 | 03/31/2020 | Greece:Athens |
| MT459925.1 | 03/31/2020 | Greece:Athens |
| MT461620.1 | 03/31/2020 | USA:WA |
| MT461621.1 | 03/31/2020 | USA:WA |
| MT461622.1 | 03/31/2020 | USA:WA |
| MT461623.1 | 03/31/2020 | USA:WA |
| MT461626.1 | 03/31/2020 | USA:WA |
| MT461627.1 | 03/31/2020 | USA:WA |
| MT461629.1 | 03/31/2020 | USA:WA |
| MT461630.1 | 03/31/2020 | USA:WA |
| MT474131.1 | 03/31/2020 | USA:CA |
| MT474134.1 | 03/31/2020 | USA:CA |
| MT474135.1 | 03/31/2020 | USA:CA |
| MT499220.1 | 03/31/2020 | Tunisia |
| MT506707.1 | 03/31/2020 | USA |
| MT507277.1 | 03/31/2020 | USA |
| MT507282.1 | 03/31/2020 | USA |
| MT509482.1 | 03/31/2020 | USA |
| MT511068.1 | 03/31/2020 | Poland |
| MT511070.1 | 03/31/2020 | Poland |
| MT511074.1 | 03/31/2020 | Poland |
| MT517424.1 | 03/31/2020 | Czech-Republic |
| MT517425.1 | 03/31/2020 | Czech-Republic |

| | | |
|---|---|---|
| MT533235.1 | 03/31/2020 | USA:CA |
| MT628251.1 | 03/31/2020 | USA:CA |
| MT632626.1 | 03/31/2020 | USA:Washington-King-County |
| MT632631.1 | 03/31/2020 | USA:Washington-King-County |
| MT632632.1 | 03/31/2020 | USA:Washington-King-County |
| MT632772.1 | 03/31/2020 | USA:Washington-Pierce-County |
| MT632786.1 | 03/31/2020 | USA:Washington-Pierce-County |
| MT632795.1 | 03/31/2020 | USA:Washington-Pierce-County |
| MT632810.1 | 03/31/2020 | USA:Washington-Skagit-County |
| MT632823.1 | 03/31/2020 | USA:Washington-Pierce-County |
| MT632824.1 | 03/31/2020 | USA:Washington-Pierce-County |
| MT632830.1 | 03/31/2020 | USA:Washington-Pierce-County |
| MT632840.1 | 03/31/2020 | USA:Washington-Pierce-County |
| MT632842.1 | 03/31/2020 | USA:Washington-Pierce-County |
| MT632850.1 | 03/31/2020 | USA:Washington-Pierce-County |
| MT632860.1 | 03/31/2020 | USA:Washington-Pierce-County |
| MT632866.1 | 03/31/2020 | USA:Washington-Yakima-County |
| MT632873.1 | 03/31/2020 | USA:Washington-Pierce-County |
| MT632874.1 | 03/31/2020 | USA:Washington-King-County |
| MT632877.1 | 03/31/2020 | USA:Washington-Pierce-County |
| MT632880.1 | 03/31/2020 | USA:Washington-Pierce-County |
| MT632890.1 | 03/31/2020 | USA:Washington-Yakima-County |
| MT632909.1 | 03/31/2020 | USA:Washington-Pierce-County |
| MT632941.1 | 03/31/2020 | USA:Washington-Yakima-County |
| MT632949.1 | 03/31/2020 | USA:Washington-Yakima-County |
| MT632983.1 | 03/31/2020 | USA:Washington-King-County |
| MT632992.1 | 03/31/2020 | USA:Washington-Yakima-County |
| MT633006.1 | 03/31/2020 | USA:Washington-Yakima-County |
| MT633020.1 | 03/31/2020 | USA:Washington-Yakima-County |
| MT633039.1 | 03/31/2020 | USA:Washington-Yakima-County |
| MT635253.1 | 03/31/2020 | USA:Connecticut |
| MT646049.1 | 03/31/2020 | USA |
| MT358644.1 | 04/01/2020 | USA:WA |
| MT358743.1 | 04/01/2020 | USA:WA |
| MT358744.1 | 04/01/2020 | USA:WA |
| MT380730.1 | 04/01/2020 | USA |
| MT412225.1 | 04/01/2020 | USA:WA |
| MT412226.1 | 04/01/2020 | USA:WA |
| MT412229.1 | 04/01/2020 | USA:WA |
| MT412230.1 | 04/01/2020 | USA:WA |
| MT412232.1 | 04/01/2020 | USA:WA |
| MT412256.1 | 04/01/2020 | USA:WA |
| MT412291.1 | 04/01/2020 | USA:WA |
| MT412292.1 | 04/01/2020 | USA:WA |
| MT412293.1 | 04/01/2020 | USA:WA |
| MT412294.1 | 04/01/2020 | USA:WA |

| | | |
|---|---|---|
| MT412296.1 | 04/01/2020 | USA:WA |
| MT412310.1 | 04/01/2020 | USA:WA |
| MT412311.1 | 04/01/2020 | USA:WA |
| MT412312.1 | 04/01/2020 | USA:WA |
| MT412313.1 | 04/01/2020 | USA:WA |
| MT412314.1 | 04/01/2020 | USA:WA |
| MT412315.1 | 04/01/2020 | USA:WA |
| MT412317.1 | 04/01/2020 | USA:CT |
| MT412318.1 | 04/01/2020 | USA:CT |
| MT412319.1 | 04/01/2020 | USA:CT |
| MT412320.1 | 04/01/2020 | USA:CT |
| MT412322.1 | 04/01/2020 | USA:CT |
| MT412325.1 | 04/01/2020 | USA:WA |
| MT412327.1 | 04/01/2020 | USA:ID |
| MT412329.1 | 04/01/2020 | USA:CT |
| MT412330.1 | 04/01/2020 | USA:CT |
| MT419821.1 | 04/01/2020 | Puerto-Rico |
| MT419822.1 | 04/01/2020 | Puerto-Rico |
| MT449636.1 | 04/01/2020 | USA:CA |
| MT451560.1 | 04/01/2020 | Australia:Victoria |
| MT451561.1 | 04/01/2020 | Australia:Victoria |
| MT451562.1 | 04/01/2020 | Australia:Victoria |
| MT451563.1 | 04/01/2020 | Australia:Victoria |
| MT451564.1 | 04/01/2020 | Australia:Victoria |
| MT451565.1 | 04/01/2020 | Australia:Victoria |
| MT451566.1 | 04/01/2020 | Australia:Victoria |
| MT451572.1 | 04/01/2020 | Australia:Victoria |
| MT451691.1 | 04/01/2020 | Australia:Victoria |
| MT451692.1 | 04/01/2020 | Australia:Victoria |
| MT451693.1 | 04/01/2020 | Australia:Victoria |
| MT451694.1 | 04/01/2020 | Australia:Victoria |
| MT451695.1 | 04/01/2020 | Australia:Victoria |
| MT451696.1 | 04/01/2020 | Australia:Victoria |
| MT451697.1 | 04/01/2020 | Australia:Victoria |
| MT451698.1 | 04/01/2020 | Australia:Victoria |
| MT451699.1 | 04/01/2020 | Australia:Victoria |
| MT451700.1 | 04/01/2020 | Australia:Victoria |
| MT451701.1 | 04/01/2020 | Australia:Victoria |
| MT451798.1 | 04/01/2020 | Australia:Victoria |
| MT451799.1 | 04/01/2020 | Australia:Victoria |
| MT451800.1 | 04/01/2020 | Australia:Victoria |
| MT451801.1 | 04/01/2020 | Australia:Victoria |
| MT451802.1 | 04/01/2020 | Australia:Victoria |
| MT451803.1 | 04/01/2020 | Australia:Victoria |
| MT451804.1 | 04/01/2020 | Australia:Victoria |
| MT451805.1 | 04/01/2020 | Australia:Victoria |

| | | |
|---|---|---|
| MT451806.1 | 04/01/2020 | Australia:Victoria |
| MT451807.1 | 04/01/2020 | Australia:Victoria |
| MT451808.1 | 04/01/2020 | Australia:Victoria |
| MT451809.1 | 04/01/2020 | Australia:Victoria |
| MT451810.1 | 04/01/2020 | Australia:Victoria |
| MT451811.1 | 04/01/2020 | Australia:Victoria |
| MT451812.1 | 04/01/2020 | Australia:Victoria |
| MT451813.1 | 04/01/2020 | Australia:Victoria |
| MT459924.1 | 04/01/2020 | Greece:Athens |
| MT461613.1 | 04/01/2020 | USA:WA |
| MT461614.1 | 04/01/2020 | USA:WA |
| MT461615.1 | 04/01/2020 | USA:WA |
| MT461616.1 | 04/01/2020 | USA:WA |
| MT461617.1 | 04/01/2020 | USA:WA |
| MT461618.1 | 04/01/2020 | USA |
| MT461619.1 | 04/01/2020 | USA |
| MT481929.1 | 04/01/2020 | USA:CA |
| MT481931.1 | 04/01/2020 | USA:CA |
| MT499218.1 | 04/01/2020 | Tunisia |
| MT506565.1 | 04/01/2020 | USA:Michigan |
| MT506893.1 | 04/01/2020 | USA:Dane-County-Wisconsin |
| MT506906.1 | 04/01/2020 | USA:Dane-County-Wisconsin |
| MT511066.1 | 04/01/2020 | Poland |
| MT511073.1 | 04/01/2020 | Poland |
| MT511082.1 | 04/01/2020 | Poland |
| MT517419.1 | 04/01/2020 | Hong-Kong |
| MT520173.1 | 04/01/2020 | USA:Massachusetts |
| MT520175.1 | 04/01/2020 | USA:Massachusetts |
| MT520195.1 | 04/01/2020 | USA:Massachusetts |
| MT520206.1 | 04/01/2020 | USA:Massachusetts |
| MT520208.1 | 04/01/2020 | USA:Massachusetts |
| MT520216.1 | 04/01/2020 | USA:Massachusetts |
| MT520219.1 | 04/01/2020 | USA:Massachusetts |
| MT520223.1 | 04/01/2020 | USA:Massachusetts |
| MT520226.1 | 04/01/2020 | USA:Massachusetts |
| MT520230.1 | 04/01/2020 | USA:Massachusetts |
| MT520232.1 | 04/01/2020 | USA:Massachusetts |
| MT520235.1 | 04/01/2020 | USA:Massachusetts |
| MT520239.1 | 04/01/2020 | USA:Massachusetts |
| MT520244.1 | 04/01/2020 | USA:Massachusetts |
| MT520246.1 | 04/01/2020 | USA:Massachusetts |
| MT520247.1 | 04/01/2020 | USA:Massachusetts |
| MT520256.1 | 04/01/2020 | USA:Massachusetts |
| MT520268.1 | 04/01/2020 | USA:Massachusetts |
| MT520270.1 | 04/01/2020 | USA:Massachusetts |
| MT520271.1 | 04/01/2020 | USA:Massachusetts |

| | | |
|---|---|---|
| MT520273.1 | 04/01/2020 | USA:Massachusetts |
| MT520285.1 | 04/01/2020 | USA:Massachusetts |
| MT520309.1 | 04/01/2020 | USA:Massachusetts |
| MT520318.1 | 04/01/2020 | USA:Massachusetts |
| MT520324.1 | 04/01/2020 | USA:Massachusetts |
| MT520330.1 | 04/01/2020 | USA:Massachusetts |
| MT520335.1 | 04/01/2020 | USA:Massachusetts |
| MT520340.1 | 04/01/2020 | USA:Massachusetts |
| MT520349.1 | 04/01/2020 | USA:Massachusetts |
| MT520357.1 | 04/01/2020 | USA:Massachusetts |
| MT520362.1 | 04/01/2020 | USA:Massachusetts |
| MT520369.1 | 04/01/2020 | USA:Massachusetts |
| MT520383.1 | 04/01/2020 | USA:Massachusetts |
| MT520417.1 | 04/01/2020 | USA:Massachusetts |
| MT520427.1 | 04/01/2020 | USA:Massachusetts |
| MT520454.1 | 04/01/2020 | USA:Massachusetts |
| MT520483.1 | 04/01/2020 | USA:Massachusetts |
| MT520491.1 | 04/01/2020 | USA:Massachusetts |
| MT520505.1 | 04/01/2020 | USA:Massachusetts |
| MT520507.1 | 04/01/2020 | USA:Massachusetts |
| MT520536.1 | 04/01/2020 | USA:Massachusetts |
| MT520538.1 | 04/01/2020 | USA:Massachusetts |
| MT520539.1 | 04/01/2020 | USA:Massachusetts |
| MT533236.1 | 04/01/2020 | USA:CA |
| MT533237.1 | 04/01/2020 | USA:CA |
| MT535482.1 | 04/01/2020 | USA |
| MT535483.1 | 04/01/2020 | USA |
| MT535484.1 | 04/01/2020 | USA |
| MT569470.1 | 04/01/2020 | Serbia:Novi-Sad |
| MT627622.1 | 04/01/2020 | USA:Washington |
| MT627631.1 | 04/01/2020 | USA:Washington |
| MT627924.1 | 04/01/2020 | USA:Washington |
| MT628181.1 | 04/01/2020 | USA:CA |
| MT628184.1 | 04/01/2020 | USA:CA |
| MT628186.1 | 04/01/2020 | USA:CA |
| MT628222.1 | 04/01/2020 | USA:CA |
| MT628252.1 | 04/01/2020 | USA:CA |
| MT628254.1 | 04/01/2020 | USA:CA |
| MT631813.1 | 04/01/2020 | USA:CA |
| MT632804.1 | 04/01/2020 | USA:Washington-Yakima-County |
| MT632836.1 | 04/01/2020 | USA:Washington-Yakima-County |
| MT632858.1 | 04/01/2020 | USA:Washington-King-County |
| MT632859.1 | 04/01/2020 | USA:Washington-Yakima-County |
| MT632863.1 | 04/01/2020 | USA:Washington-Pierce-County |
| MT632865.1 | 04/01/2020 | USA:Washington-Franklin-County |
| MT632876.1 | 04/01/2020 | USA:Washington-Pierce-County |

| | | |
|---|---|---|
| MT632879.1 | 04/01/2020 | USA:Washington-Pierce-County |
| MT632884.1 | 04/01/2020 | USA:Washington-King-County |
| MT632910.1 | 04/01/2020 | USA:Washington-Benton-County |
| MT632914.1 | 04/01/2020 | USA:Washington-Franklin-County |
| MT632916.1 | 04/01/2020 | USA:Washington-Franklin-County |
| MT632919.1 | 04/01/2020 | USA:Washington-Asotin-County |
| MT632920.1 | 04/01/2020 | USA:Washington-Yakima-County |
| MT632931.1 | 04/01/2020 | USA:Washington-Adams-County |
| MT632936.1 | 04/01/2020 | USA:Washington-King-County |
| MT632950.1 | 04/01/2020 | USA:Washington-Pierce-County |
| MT632952.1 | 04/01/2020 | USA:Washington-Yakima-County |
| MT632955.1 | 04/01/2020 | USA:Washington-King-County |
| MT632957.1 | 04/01/2020 | USA:Washington-Yakima-County |
| MT632960.1 | 04/01/2020 | USA:Washington-Yakima-County |
| MT632974.1 | 04/01/2020 | USA:Washington-Yakima-County |
| MT632976.1 | 04/01/2020 | USA:Washington-Yakima-County |
| MT632981.1 | 04/01/2020 | USA:Washington-Yakima-County |
| MT632993.1 | 04/01/2020 | USA:Washington-Yakima-County |
| MT632998.1 | 04/01/2020 | USA:Washington-Yakima-County |
| MT633010.1 | 04/01/2020 | USA:Washington-Yakima-County |
| MT633013.1 | 04/01/2020 | USA:Washington-Yakima-County |
| MT633016.1 | 04/01/2020 | USA:Washington-Yakima-County |
| MT633018.1 | 04/01/2020 | USA:Washington-Yakima-County |
| MT633021.1 | 04/01/2020 | USA:Washington-Yakima-County |
| MT633023.1 | 04/01/2020 | USA:Washington-Yakima-County |
| MT633027.1 | 04/01/2020 | USA:Washington-Yakima-County |
| MT633030.1 | 04/01/2020 | USA:Washington-Yakima-County |
| MT633035.1 | 04/01/2020 | USA:Washington-Yakima-County |
| MT633036.1 | 04/01/2020 | USA:Washington-Yakima-County |
| MT633037.1 | 04/01/2020 | USA:Washington-Yakima-County |
| MT633038.1 | 04/01/2020 | USA:Washington-Yakima-County |
| MT635251.1 | 04/01/2020 | USA:Connecticut |
| MT646050.1 | 04/01/2020 | USA |
| MT300186.2 | 04/02/2020 | USA:North-Carolina |
| MT365033.1 | 04/02/2020 | USA:New-York |
| MT375476.1 | 04/02/2020 | USA:WA |
| MT375477.1 | 04/02/2020 | USA:WA |
| MT375478.1 | 04/02/2020 | USA:WA |
| MT375479.1 | 04/02/2020 | USA:WA |
| MT375480.1 | 04/02/2020 | USA:WA |
| MT375481.1 | 04/02/2020 | USA:WA |
| MT375482.1 | 04/02/2020 | USA:WA |
| MT375483.1 | 04/02/2020 | USA |
| MT412227.1 | 04/02/2020 | USA:WA |
| MT412228.1 | 04/02/2020 | USA:WA |
| MT412254.1 | 04/02/2020 | USA |

| | | |
|---|---|---|
| MT412255.1 | 04/02/2020 | USA:WA |
| MT412257.1 | 04/02/2020 | USA:WA |
| MT412316.1 | 04/02/2020 | USA:WA |
| MT412321.1 | 04/02/2020 | USA:WA |
| MT412323.1 | 04/02/2020 | USA:WA |
| MT412324.1 | 04/02/2020 | USA:WA |
| MT412326.1 | 04/02/2020 | USA:WA |
| MT412328.1 | 04/02/2020 | USA:CT |
| MT444541.1 | 04/02/2020 | USA |
| MT444542.1 | 04/02/2020 | USA |
| MT444543.1 | 04/02/2020 | USA |
| MT444544.1 | 04/02/2020 | USA |
| MT444546.1 | 04/02/2020 | USA |
| MT444548.1 | 04/02/2020 | USA |
| MT444549.1 | 04/02/2020 | USA |
| MT444550.1 | 04/02/2020 | USA |
| MT444577.1 | 04/02/2020 | USA |
| MT444581.1 | 04/02/2020 | USA |
| MT444582.1 | 04/02/2020 | USA |
| MT444584.1 | 04/02/2020 | USA |
| MT444585.1 | 04/02/2020 | USA |
| MT444586.1 | 04/02/2020 | USA |
| MT444587.1 | 04/02/2020 | USA |
| MT451573.1 | 04/02/2020 | Australia:Victoria |
| MT451574.1 | 04/02/2020 | Australia:Victoria |
| MT451702.1 | 04/02/2020 | Australia:Victoria |
| MT451703.1 | 04/02/2020 | Australia:Victoria |
| MT451704.1 | 04/02/2020 | Australia:Victoria |
| MT451705.1 | 04/02/2020 | Australia:Victoria |
| MT451706.1 | 04/02/2020 | Australia:Victoria |
| MT451814.1 | 04/02/2020 | Australia:Victoria |
| MT451815.1 | 04/02/2020 | Australia:Victoria |
| MT451816.1 | 04/02/2020 | Australia:Victoria |
| MT451817.1 | 04/02/2020 | Australia:Victoria |
| MT451818.1 | 04/02/2020 | Australia:Victoria |
| MT451819.1 | 04/02/2020 | Australia:Victoria |
| MT451820.1 | 04/02/2020 | Australia:Victoria |
| MT451821.1 | 04/02/2020 | Australia:Victoria |
| MT461631.1 | 04/02/2020 | USA:ID |
| MT461632.1 | 04/02/2020 | USA:ID |
| MT461633.1 | 04/02/2020 | USA:ID |
| MT461635.1 | 04/02/2020 | USA:ID |
| MT461636.1 | 04/02/2020 | USA:ID |
| MT461637.1 | 04/02/2020 | USA:ID |
| MT461638.1 | 04/02/2020 | USA:ID |
| MT461662.1 | 04/02/2020 | USA:CT |

| | | |
|---|---|---|
| MT461663.1 | 04/02/2020 | USA:CT |
| MT461664.1 | 04/02/2020 | USA:CT |
| MT461666.1 | 04/02/2020 | USA:CT |
| MT461668.1 | 04/02/2020 | USA:CT |
| MT470219.1 | 04/02/2020 | Colombia |
| MT477850.1 | 04/02/2020 | USA:AK |
| MT481930.1 | 04/02/2020 | USA:CA |
| MT506566.1 | 04/02/2020 | USA:Michigan |
| MT506567.1 | 04/02/2020 | USA:Michigan |
| MT506568.1 | 04/02/2020 | USA:Michigan |
| MT506569.1 | 04/02/2020 | USA:Michigan |
| MT507279.1 | 04/02/2020 | USA |
| MT511067.1 | 04/02/2020 | Poland |
| MT511075.1 | 04/02/2020 | Poland |
| MT511078.1 | 04/02/2020 | Poland |
| MT520214.1 | 04/02/2020 | USA:Massachusetts |
| MT520264.1 | 04/02/2020 | USA:Massachusetts |
| MT520272.1 | 04/02/2020 | USA:Massachusetts |
| MT520329.1 | 04/02/2020 | USA:Massachusetts |
| MT520373.1 | 04/02/2020 | USA:Massachusetts |
| MT520405.1 | 04/02/2020 | USA:Massachusetts |
| MT520461.1 | 04/02/2020 | USA:Massachusetts |
| MT520494.1 | 04/02/2020 | USA:Massachusetts |
| MT520518.1 | 04/02/2020 | USA:Massachusetts |
| MT533238.1 | 04/02/2020 | USA:CA |
| MT590599.1 | 04/02/2020 | Taiwan |
| MT627756.1 | 04/02/2020 | USA:Washington-King-County |
| MT628095.1 | 04/02/2020 | USA:CA |
| MT628223.1 | 04/02/2020 | USA:CA |
| MT628224.1 | 04/02/2020 | USA:CA |
| MT628253.1 | 04/02/2020 | USA:CA |
| MT631816.1 | 04/02/2020 | USA:CA |
| MT631817.1 | 04/02/2020 | USA:CA |
| MT631818.1 | 04/02/2020 | USA:CA |
| MT631819.1 | 04/02/2020 | USA:CA |
| MT631820.1 | 04/02/2020 | USA:CA |
| MT631822.1 | 04/02/2020 | USA:CA |
| MT631823.1 | 04/02/2020 | USA:CA |
| MT631824.1 | 04/02/2020 | USA:CA |
| MT631825.1 | 04/02/2020 | USA:CA |
| MT631827.1 | 04/02/2020 | USA:CA |
| MT631828.1 | 04/02/2020 | USA:CA |
| MT631829.1 | 04/02/2020 | USA:CA |
| MT631830.1 | 04/02/2020 | USA:CA |
| MT631831.1 | 04/02/2020 | USA:CA |
| MT631833.1 | 04/02/2020 | USA:CA |

| | | |
|---|---|---|
| MT632766.1 | 04/02/2020 | USA:Washington-Pierce-County |
| MT632771.1 | 04/02/2020 | USA:Washington-Yakima-County |
| MT632776.1 | 04/02/2020 | USA:Washington-Yakima-County |
| MT632784.1 | 04/02/2020 | USA:Washington-Yakima-County |
| MT632788.1 | 04/02/2020 | USA:Washington-Pierce-County |
| MT632789.1 | 04/02/2020 | USA:Washington-Whatcom-County |
| MT632790.1 | 04/02/2020 | USA:Washington-Pierce-County |
| MT632794.1 | 04/02/2020 | USA:Washington-Yakima-County |
| MT632796.1 | 04/02/2020 | USA:Washington-Yakima-County |
| MT632817.1 | 04/02/2020 | USA:Washington-Mason-County |
| MT632827.1 | 04/02/2020 | USA:Washington-Yakima-County |
| MT632828.1 | 04/02/2020 | USA:Washington-Pierce-County |
| MT632853.1 | 04/02/2020 | USA:Washington-Yakima-County |
| MT632882.1 | 04/02/2020 | USA:Washington-Yakima-County |
| MT632887.1 | 04/02/2020 | USA:Washington-Yakima-County |
| MT632894.1 | 04/02/2020 | USA:Washington-Snohomish-County |
| MT632930.1 | 04/02/2020 | USA:Washington-Yakima-County |
| MT632961.1 | 04/02/2020 | USA:Washington-Yakima-County |
| MT632966.1 | 04/02/2020 | USA:Washington-Yakima-County |
| MT632977.1 | 04/02/2020 | USA:Washington-King-County |
| MT633008.1 | 04/02/2020 | USA:Washington-Yakima-County |
| MT633017.1 | 04/02/2020 | USA:Washington-Yakima-County |
| MT633029.1 | 04/02/2020 | USA:Washington-Yakima-County |
| MT641708.1 | 04/02/2020 | Australia:Victoria |
| MT385442.1 | 04/03/2020 | USA:CA |
| MT385452.1 | 04/03/2020 | USA:CA |
| MT385475.1 | 04/03/2020 | USA:CA |
| MT412331.1 | 04/03/2020 | USA:WA |
| MT412332.1 | 04/03/2020 | USA:WA |
| MT412333.1 | 04/03/2020 | USA:WA |
| MT412334.1 | 04/03/2020 | USA:WA |
| MT412335.1 | 04/03/2020 | USA:WA |
| MT412336.1 | 04/03/2020 | USA:WA |
| MT412337.1 | 04/03/2020 | USA:WA |
| MT444547.1 | 04/03/2020 | USA |
| MT444551.1 | 04/03/2020 | USA |
| MT444552.1 | 04/03/2020 | USA |
| MT444553.1 | 04/03/2020 | USA |
| MT444554.1 | 04/03/2020 | USA |
| MT444555.1 | 04/03/2020 | USA |
| MT444569.1 | 04/03/2020 | USA |
| MT444570.1 | 04/03/2020 | USA |
| MT444571.1 | 04/03/2020 | USA |
| MT444572.1 | 04/03/2020 | USA |
| MT444573.1 | 04/03/2020 | USA |
| MT444574.1 | 04/03/2020 | USA |

| | | |
|---|---|---|
| MT444575.1 | 04/03/2020 | USA |
| MT444576.1 | 04/03/2020 | USA |
| MT444578.1 | 04/03/2020 | USA |
| MT446356.1 | 04/03/2020 | USA:Lacombe-LA |
| MT449640.1 | 04/03/2020 | USA:CA |
| MT451575.1 | 04/03/2020 | Australia:Victoria |
| MT451577.1 | 04/03/2020 | Australia:Victoria |
| MT451578.1 | 04/03/2020 | Australia:Victoria |
| MT451579.1 | 04/03/2020 | Australia:Victoria |
| MT451580.1 | 04/03/2020 | Australia:Victoria |
| MT451581.1 | 04/03/2020 | Australia:Victoria |
| MT451582.1 | 04/03/2020 | Australia:Victoria |
| MT451707.1 | 04/03/2020 | Australia:Victoria |
| MT451708.1 | 04/03/2020 | Australia:Victoria |
| MT451709.1 | 04/03/2020 | Australia:Victoria |
| MT451710.1 | 04/03/2020 | Australia:Victoria |
| MT451711.1 | 04/03/2020 | Australia:Victoria |
| MT451712.1 | 04/03/2020 | Australia:Victoria |
| MT451713.1 | 04/03/2020 | Australia:Victoria |
| MT451714.1 | 04/03/2020 | Australia:Victoria |
| MT451715.1 | 04/03/2020 | Australia:Victoria |
| MT451716.1 | 04/03/2020 | Australia:Victoria |
| MT451717.1 | 04/03/2020 | Australia:Victoria |
| MT451718.1 | 04/03/2020 | Australia:Victoria |
| MT451719.1 | 04/03/2020 | Australia:Victoria |
| MT451720.1 | 04/03/2020 | Australia:Victoria |
| MT451721.1 | 04/03/2020 | Australia:Victoria |
| MT451722.1 | 04/03/2020 | Australia:Victoria |
| MT451723.1 | 04/03/2020 | Australia:Victoria |
| MT451724.1 | 04/03/2020 | Australia:Victoria |
| MT461634.1 | 04/03/2020 | USA:ID |
| MT461640.1 | 04/03/2020 | USA:WA |
| MT461641.1 | 04/03/2020 | USA:WA |
| MT461646.1 | 04/03/2020 | USA:WA |
| MT461647.1 | 04/03/2020 | USA:WA |
| MT461648.1 | 04/03/2020 | USA:WA |
| MT461665.1 | 04/03/2020 | USA:CT |
| MT461667.1 | 04/03/2020 | USA:CT |
| MT481933.1 | 04/03/2020 | USA:CA |
| MT499204.1 | 04/03/2020 | USA:CA |
| MT499206.1 | 04/03/2020 | USA:CA |
| MT506570.1 | 04/03/2020 | USA:Michigan |
| MT506571.1 | 04/03/2020 | USA:Michigan |
| MT506572.1 | 04/03/2020 | USA:Michigan |
| MT506573.1 | 04/03/2020 | USA:Michigan |
| MT506577.1 | 04/03/2020 | USA:Michigan |

| | | |
|---|---|---|
| MT506977.1 | 04/03/2020 | USA:FL |
| MT506981.1 | 04/03/2020 | USA:FL |
| MT507280.1 | 04/03/2020 | USA |
| MT511069.1 | 04/03/2020 | Poland |
| MT511072.1 | 04/03/2020 | Poland |
| MT520210.1 | 04/03/2020 | USA:Massachusetts |
| MT520292.1 | 04/03/2020 | USA:Massachusetts |
| MT520323.1 | 04/03/2020 | USA:Massachusetts |
| MT520331.1 | 04/03/2020 | USA:Massachusetts |
| MT520402.1 | 04/03/2020 | USA:Massachusetts |
| MT520452.1 | 04/03/2020 | USA:Massachusetts |
| MT520477.1 | 04/03/2020 | USA:Massachusetts |
| MT520487.1 | 04/03/2020 | USA:Massachusetts |
| MT536184.1 | 04/03/2020 | USA:SLIDELL-LA |
| MT585089.1 | 04/03/2020 | USA |
| MT585091.1 | 04/03/2020 | USA |
| MT585093.1 | 04/03/2020 | USA |
| MT585102.1 | 04/03/2020 | USA |
| MT628149.1 | 04/03/2020 | USA:CA |
| MT628255.1 | 04/03/2020 | USA:CA |
| MT628256.1 | 04/03/2020 | USA:CA |
| MT631835.1 | 04/03/2020 | USA:CA |
| MT631836.1 | 04/03/2020 | USA:CA |
| MT631837.1 | 04/03/2020 | USA:CA |
| MT631838.1 | 04/03/2020 | USA:CA |
| MT631839.1 | 04/03/2020 | USA:CA |
| MT631840.1 | 04/03/2020 | USA:CA |
| MT632797.1 | 04/03/2020 | USA:Washington-Pierce-County |
| MT632832.1 | 04/03/2020 | USA:Washington-Yakima-County |
| MT632872.1 | 04/03/2020 | USA:Washington-Yakima-County |
| MT632932.1 | 04/03/2020 | USA:Washington-King-County |
| MT632985.1 | 04/03/2020 | USA:Washington-Yakima-County |
| MT412274.1 | 04/04/2020 | USA:CT |
| MT412282.1 | 04/04/2020 | USA:CT |
| MT412283.1 | 04/04/2020 | USA:CT |
| MT412284.1 | 04/04/2020 | USA:CT |
| MT444579.1 | 04/04/2020 | USA |
| MT444580.1 | 04/04/2020 | USA |
| MT446343.1 | 04/04/2020 | USA:New-Orleans-LA |
| MT446344.1 | 04/04/2020 | USA:LOCKPORT-LA |
| MT446345.1 | 04/04/2020 | USA:NEW-ORLEANS-LA |
| MT446349.1 | 04/04/2020 | USA:KENNER-LA |
| MT449638.1 | 04/04/2020 | USA:CA |
| MT449639.1 | 04/04/2020 | USA:CA |
| MT449641.1 | 04/04/2020 | USA:CA |
| MT449643.1 | 04/04/2020 | USA:CA |

| | | |
|---|---|---|
| MT449671.1 | 04/04/2020 | USA:CA |
| MT449677.1 | 04/04/2020 | USA:CA |
| MT451583.1 | 04/04/2020 | Australia:Victoria |
| MT451584.1 | 04/04/2020 | Australia:Victoria |
| MT451585.1 | 04/04/2020 | Australia:Victoria |
| MT451586.1 | 04/04/2020 | Australia:Victoria |
| MT451587.1 | 04/04/2020 | Australia:Victoria |
| MT451588.1 | 04/04/2020 | Australia:Victoria |
| MT451589.1 | 04/04/2020 | Australia:Victoria |
| MT451591.1 | 04/04/2020 | Australia:Victoria |
| MT451592.1 | 04/04/2020 | Australia:Victoria |
| MT451593.1 | 04/04/2020 | Australia:Victoria |
| MT451594.1 | 04/04/2020 | Australia:Victoria |
| MT451595.1 | 04/04/2020 | Australia:Victoria |
| MT451596.1 | 04/04/2020 | Australia:Victoria |
| MT451597.1 | 04/04/2020 | Australia:Victoria |
| MT451598.1 | 04/04/2020 | Australia:Victoria |
| MT451599.1 | 04/04/2020 | Australia:Victoria |
| MT451725.1 | 04/04/2020 | Australia:Victoria |
| MT451726.1 | 04/04/2020 | Australia:Victoria |
| MT451727.1 | 04/04/2020 | Australia:Victoria |
| MT451728.1 | 04/04/2020 | Australia:Victoria |
| MT451729.1 | 04/04/2020 | Australia:Victoria |
| MT459847.1 | 04/04/2020 | Greece:Athens |
| MT459979.1 | 04/04/2020 | Serbia:Novi-Pazar |
| MT461642.1 | 04/04/2020 | USA:WA |
| MT461645.1 | 04/04/2020 | USA:WA |
| MT461650.1 | 04/04/2020 | USA:WA |
| MT461651.1 | 04/04/2020 | USA:WA |
| MT461652.1 | 04/04/2020 | USA:WA |
| MT461656.1 | 04/04/2020 | USA:WA |
| MT461657.1 | 04/04/2020 | USA:WA |
| MT461658.1 | 04/04/2020 | USA:WA |
| MT461659.1 | 04/04/2020 | USA:WA |
| MT477851.1 | 04/04/2020 | USA:AK |
| MT481932.1 | 04/04/2020 | USA:CA |
| MT499198.1 | 04/04/2020 | USA:CA |
| MT506574.1 | 04/04/2020 | USA:Michigan |
| MT506575.1 | 04/04/2020 | USA:Michigan |
| MT506576.1 | 04/04/2020 | USA:Michigan |
| MT506979.1 | 04/04/2020 | USA:FL |
| MT506980.1 | 04/04/2020 | USA:FL |
| MT506982.1 | 04/04/2020 | USA:FL |
| MT506984.1 | 04/04/2020 | USA:FL |
| MT506985.1 | 04/04/2020 | USA:FL |
| MT520174.1 | 04/04/2020 | USA:Massachusetts |

| | | |
|---|---|---|
| MT520227.1 | 04/04/2020 | USA:Massachusetts |
| MT520234.1 | 04/04/2020 | USA:Massachusetts |
| MT520240.1 | 04/04/2020 | USA:Massachusetts |
| MT520243.1 | 04/04/2020 | USA:Massachusetts |
| MT520251.1 | 04/04/2020 | USA:Massachusetts |
| MT520252.1 | 04/04/2020 | USA:Massachusetts |
| MT520265.1 | 04/04/2020 | USA:Massachusetts |
| MT520276.1 | 04/04/2020 | USA:Massachusetts |
| MT520283.1 | 04/04/2020 | USA:Massachusetts |
| MT520295.1 | 04/04/2020 | USA:Massachusetts |
| MT520298.1 | 04/04/2020 | USA:Massachusetts |
| MT520300.1 | 04/04/2020 | USA:Massachusetts |
| MT520301.1 | 04/04/2020 | USA:Massachusetts |
| MT520302.1 | 04/04/2020 | USA:Massachusetts |
| MT520305.1 | 04/04/2020 | USA:Massachusetts |
| MT520319.1 | 04/04/2020 | USA:Massachusetts |
| MT520347.1 | 04/04/2020 | USA:Massachusetts |
| MT520359.1 | 04/04/2020 | USA:Massachusetts |
| MT520371.1 | 04/04/2020 | USA:Massachusetts |
| MT520382.1 | 04/04/2020 | USA:Massachusetts |
| MT520403.1 | 04/04/2020 | USA:Massachusetts |
| MT520426.1 | 04/04/2020 | USA:Massachusetts |
| MT520431.1 | 04/04/2020 | USA:Massachusetts |
| MT520436.1 | 04/04/2020 | USA:Massachusetts |
| MT520458.1 | 04/04/2020 | USA:Massachusetts |
| MT520460.1 | 04/04/2020 | USA:Massachusetts |
| MT520465.1 | 04/04/2020 | USA:Massachusetts |
| MT520467.1 | 04/04/2020 | USA:Massachusetts |
| MT520480.1 | 04/04/2020 | USA:Massachusetts |
| MT520495.1 | 04/04/2020 | USA:Massachusetts |
| MT520516.1 | 04/04/2020 | USA:Massachusetts |
| MT520529.1 | 04/04/2020 | USA:Massachusetts |
| MT536188.1 | 04/04/2020 | USA:KENNER-LA |
| MT585084.1 | 04/04/2020 | USA |
| MT585087.1 | 04/04/2020 | USA |
| MT585088.1 | 04/04/2020 | USA |
| MT585090.1 | 04/04/2020 | USA |
| MT585092.1 | 04/04/2020 | USA |
| MT585094.1 | 04/04/2020 | USA |
| MT585100.1 | 04/04/2020 | USA |
| MT585103.1 | 04/04/2020 | USA |
| MT627748.1 | 04/04/2020 | USA:Washington-King-County |
| MT627751.1 | 04/04/2020 | USA:Washington-King-County |
| MT627759.1 | 04/04/2020 | USA:Washington-King-County |
| MT627761.1 | 04/04/2020 | USA:Washington-King-County |
| MT628257.1 | 04/04/2020 | USA:CA |

| | | |
|---|---|---|
| MT632785.1 | 04/04/2020 | USA:Washington-Yakima-County |
| MT632847.1 | 04/04/2020 | USA:Washington-Yakima-County |
| MT632856.1 | 04/04/2020 | USA:Washington-Yakima-County |
| MT632868.1 | 04/04/2020 | USA:Washington-Yakima-County |
| MT632895.1 | 04/04/2020 | USA:Washington-Yakima-County |
| MT632896.1 | 04/04/2020 | USA:Washington-King-County |
| MT632897.1 | 04/04/2020 | USA:Washington-Yakima-County |
| MT632911.1 | 04/04/2020 | USA:Washington-Yakima-County |
| MT632958.1 | 04/04/2020 | USA:Washington-Yakima-County |
| MT635255.1 | 04/04/2020 | USA:Connecticut |
| MT635258.1 | 04/04/2020 | USA:Connecticut |
| MT641729.1 | 04/04/2020 | Australia:Victoria |
| MT358637.1 | 04/05/2020 | India:Rajkot |
| MT375459.1 | 04/05/2020 | USA:WA |
| MT412258.1 | 04/05/2020 | USA:WA |
| MT412259.1 | 04/05/2020 | USA:WA |
| MT412260.1 | 04/05/2020 | USA |
| MT412261.1 | 04/05/2020 | USA:WA |
| MT412262.1 | 04/05/2020 | USA:WA |
| MT412263.1 | 04/05/2020 | USA:WA |
| MT412276.1 | 04/05/2020 | USA:CT |
| MT412279.1 | 04/05/2020 | USA:CT |
| MT412280.1 | 04/05/2020 | USA:CT |
| MT444588.1 | 04/05/2020 | USA |
| MT444589.1 | 04/05/2020 | USA |
| MT444590.1 | 04/05/2020 | USA |
| MT444594.1 | 04/05/2020 | USA |
| MT444596.1 | 04/05/2020 | USA |
| MT444598.1 | 04/05/2020 | USA |
| MT446359.1 | 04/05/2020 | USA:SLIDELL-LA |
| MT449642.1 | 04/05/2020 | USA:CA |
| MT449646.1 | 04/05/2020 | USA:CA |
| MT449678.1 | 04/05/2020 | USA:CA |
| MT449679.1 | 04/05/2020 | USA:CA |
| MT451600.1 | 04/05/2020 | Australia:Victoria |
| MT451601.1 | 04/05/2020 | Australia:Victoria |
| MT451602.1 | 04/05/2020 | Australia:Victoria |
| MT451603.1 | 04/05/2020 | Australia:Victoria |
| MT451604.1 | 04/05/2020 | Australia:Victoria |
| MT451605.1 | 04/05/2020 | Australia:Victoria |
| MT451606.1 | 04/05/2020 | Australia:Victoria |
| MT451607.1 | 04/05/2020 | Australia:Victoria |
| MT451608.1 | 04/05/2020 | Australia:Victoria |
| MT451609.1 | 04/05/2020 | Australia:Victoria |
| MT451610.1 | 04/05/2020 | Australia:Victoria |
| MT451611.1 | 04/05/2020 | Australia:Victoria |

| | | |
|---|---|---|
| MT451612.1 | 04/05/2020 | Australia:Victoria |
| MT451613.1 | 04/05/2020 | Australia:Victoria |
| MT451614.1 | 04/05/2020 | Australia:Victoria |
| MT451615.1 | 04/05/2020 | Australia:Victoria |
| MT451730.1 | 04/05/2020 | Australia:Victoria |
| MT461639.1 | 04/05/2020 | USA:WA |
| MT461643.1 | 04/05/2020 | USA:WA |
| MT461644.1 | 04/05/2020 | USA:WA |
| MT461649.1 | 04/05/2020 | USA |
| MT461653.1 | 04/05/2020 | USA:WA |
| MT461654.1 | 04/05/2020 | USA:WA |
| MT461655.1 | 04/05/2020 | USA:WA |
| MT461660.1 | 04/05/2020 | USA:WA |
| MT461661.1 | 04/05/2020 | USA:WA |
| MT474132.1 | 04/05/2020 | USA:CA |
| MT474133.1 | 04/05/2020 | USA:CA |
| MT483565.1 | 04/05/2020 | USA:CA |
| MT506974.1 | 04/05/2020 | USA:FL |
| MT506975.1 | 04/05/2020 | USA:FL |
| MT506976.1 | 04/05/2020 | USA:FL |
| MT506978.1 | 04/05/2020 | USA:FL |
| MT520384.1 | 04/05/2020 | USA:Massachusetts |
| MT520393.1 | 04/05/2020 | USA:Massachusetts |
| MT520416.1 | 04/05/2020 | USA:Massachusetts |
| MT520493.1 | 04/05/2020 | USA:Massachusetts |
| MT520506.1 | 04/05/2020 | USA:Massachusetts |
| MT535507.1 | 04/05/2020 | USA |
| MT628225.1 | 04/05/2020 | USA:CA |
| MT628226.1 | 04/05/2020 | USA:CA |
| MT628258.1 | 04/05/2020 | USA:CA |
| MT632825.1 | 04/05/2020 | USA:Washington-Yakima-County |
| MT632843.1 | 04/05/2020 | USA:Washington-Yakima-County |
| MT632886.1 | 04/05/2020 | USA:Washington-Yakima-County |
| MT632893.1 | 04/05/2020 | USA:Washington-Yakima-County |
| MT632905.1 | 04/05/2020 | USA:Washington-King-County |
| MT632937.1 | 04/05/2020 | USA:Washington-King-County |
| MT632940.1 | 04/05/2020 | USA:Washington-King-County |
| MT632962.1 | 04/05/2020 | USA:Washington-King-County |
| MT632971.1 | 04/05/2020 | USA:Washington-King-County |
| MT632984.1 | 04/05/2020 | USA:Washington-King-County |
| MT632987.1 | 04/05/2020 | USA:Washington-King-County |
| MT358401.1 | 04/06/2020 | USA:New-Orleans-LA |
| MT358402.1 | 04/06/2020 | USA:New-Orleans-LA |
| MT375457.1 | 04/06/2020 | USA |
| MT375458.1 | 04/06/2020 | USA:WA |
| MT375460.1 | 04/06/2020 | USA:WA |

| | | |
|---|---|---|
| MT375461.1 | 04/06/2020 | USA:WA |
| MT375462.1 | 04/06/2020 | USA:WA |
| MT375470.1 | 04/06/2020 | USA:CT |
| MT375471.1 | 04/06/2020 | USA:CT |
| MT375474.1 | 04/06/2020 | USA:CT |
| MT396243.1 | 04/06/2020 | India |
| MT396247.1 | 04/06/2020 | India |
| MT412270.1 | 04/06/2020 | USA:WA |
| MT412271.1 | 04/06/2020 | USA:WA |
| MT412272.1 | 04/06/2020 | USA:WA |
| MT412275.1 | 04/06/2020 | USA:WA |
| MT412278.1 | 04/06/2020 | USA:CT |
| MT429190.1 | 04/06/2020 | USA:Wisconsin |
| MT429191.1 | 04/06/2020 | USA:Wisconsin |
| MT444591.1 | 04/06/2020 | USA |
| MT444592.1 | 04/06/2020 | USA |
| MT444593.1 | 04/06/2020 | USA |
| MT444595.1 | 04/06/2020 | USA |
| MT444597.1 | 04/06/2020 | USA |
| MT444604.1 | 04/06/2020 | USA |
| MT446339.1 | 04/06/2020 | USA:MARINGOUIN-LA |
| MT446347.1 | 04/06/2020 | USA:RACELAND-LA |
| MT446348.1 | 04/06/2020 | USA:New-Orleans-LA |
| MT446350.1 | 04/06/2020 | USA:New-Orleans-LA |
| MT446352.1 | 04/06/2020 | USA:GHEENS-LA |
| MT446353.1 | 04/06/2020 | USA:THIBODAUX-LA |
| MT446357.1 | 04/06/2020 | USA:HOUMA-LA |
| MT446358.1 | 04/06/2020 | USA:RACELAND-LA |
| MT449644.1 | 04/06/2020 | USA:CA |
| MT449645.1 | 04/06/2020 | USA:CA |
| MT449649.1 | 04/06/2020 | USA:CA |
| MT451616.1 | 04/06/2020 | Australia:Victoria |
| MT451617.1 | 04/06/2020 | Australia:Victoria |
| MT451618.1 | 04/06/2020 | Australia:Victoria |
| MT451620.1 | 04/06/2020 | Australia:Victoria |
| MT451621.1 | 04/06/2020 | Australia:Victoria |
| MT451622.1 | 04/06/2020 | Australia:Victoria |
| MT451623.1 | 04/06/2020 | Australia:Victoria |
| MT451624.1 | 04/06/2020 | Australia:Victoria |
| MT451625.1 | 04/06/2020 | Australia:Victoria |
| MT451626.1 | 04/06/2020 | Australia:Victoria |
| MT451627.1 | 04/06/2020 | Australia:Victoria |
| MT451731.1 | 04/06/2020 | Australia:Victoria |
| MT451732.1 | 04/06/2020 | Australia:Victoria |
| MT451733.1 | 04/06/2020 | Australia:Victoria |
| MT451734.1 | 04/06/2020 | Australia:Victoria |

| | | |
|---|---|---|
| MT451735.1 | 04/06/2020 | Australia:Victoria |
| MT451736.1 | 04/06/2020 | Australia:Victoria |
| MT451737.1 | 04/06/2020 | Australia:Victoria |
| MT451738.1 | 04/06/2020 | Australia:Victoria |
| MT481934.1 | 04/06/2020 | USA:CA |
| MT483563.1 | 04/06/2020 | USA:CA |
| MT483564.1 | 04/06/2020 | USA:CA |
| MT506578.1 | 04/06/2020 | USA:Michigan |
| MT506579.1 | 04/06/2020 | USA:Michigan |
| MT506580.1 | 04/06/2020 | USA:Michigan |
| MT506581.1 | 04/06/2020 | USA:Michigan |
| MT506983.1 | 04/06/2020 | USA:FL |
| MT520196.1 | 04/06/2020 | USA:Massachusetts |
| MT520202.1 | 04/06/2020 | USA:Massachusetts |
| MT520207.1 | 04/06/2020 | USA:Massachusetts |
| MT520222.1 | 04/06/2020 | USA:Massachusetts |
| MT520228.1 | 04/06/2020 | USA:Massachusetts |
| MT520237.1 | 04/06/2020 | USA:Massachusetts |
| MT520248.1 | 04/06/2020 | USA:Massachusetts |
| MT520262.1 | 04/06/2020 | USA:Massachusetts |
| MT520263.1 | 04/06/2020 | USA:Massachusetts |
| MT520266.1 | 04/06/2020 | USA:Massachusetts |
| MT520288.1 | 04/06/2020 | USA:Massachusetts |
| MT520290.1 | 04/06/2020 | USA:Massachusetts |
| MT520308.1 | 04/06/2020 | USA:Massachusetts |
| MT520312.1 | 04/06/2020 | USA:Massachusetts |
| MT520332.1 | 04/06/2020 | USA:Massachusetts |
| MT520334.1 | 04/06/2020 | USA:Massachusetts |
| MT520343.1 | 04/06/2020 | USA:Massachusetts |
| MT520368.1 | 04/06/2020 | USA:Massachusetts |
| MT520374.1 | 04/06/2020 | USA:Massachusetts |
| MT520380.1 | 04/06/2020 | USA:Massachusetts |
| MT520400.1 | 04/06/2020 | USA:Massachusetts |
| MT520407.1 | 04/06/2020 | USA:Massachusetts |
| MT520411.1 | 04/06/2020 | USA:Massachusetts |
| MT520420.1 | 04/06/2020 | USA:Massachusetts |
| MT520421.1 | 04/06/2020 | USA:Massachusetts |
| MT520432.1 | 04/06/2020 | USA:Massachusetts |
| MT520433.1 | 04/06/2020 | USA:Massachusetts |
| MT520435.1 | 04/06/2020 | USA:Massachusetts |
| MT520440.1 | 04/06/2020 | USA:Massachusetts |
| MT520445.1 | 04/06/2020 | USA:Massachusetts |
| MT520448.1 | 04/06/2020 | USA:Massachusetts |
| MT520453.1 | 04/06/2020 | USA:Massachusetts |
| MT520464.1 | 04/06/2020 | USA:Massachusetts |
| MT520478.1 | 04/06/2020 | USA:Massachusetts |

| | | |
|---|---|---|
| MT520479.1 | 04/06/2020 | USA:Massachusetts |
| MT520482.1 | 04/06/2020 | USA:Massachusetts |
| MT520489.1 | 04/06/2020 | USA:Massachusetts |
| MT520497.1 | 04/06/2020 | USA:Massachusetts |
| MT520504.1 | 04/06/2020 | USA:Massachusetts |
| MT520510.1 | 04/06/2020 | USA:Massachusetts |
| MT520514.1 | 04/06/2020 | USA:Massachusetts |
| MT520525.1 | 04/06/2020 | USA:Massachusetts |
| MT520530.1 | 04/06/2020 | USA:Massachusetts |
| MT535504.1 | 04/06/2020 | USA |
| MT535505.1 | 04/06/2020 | USA |
| MT535506.1 | 04/06/2020 | USA |
| MT535508.1 | 04/06/2020 | USA |
| MT535509.1 | 04/06/2020 | USA |
| MT536176.1 | 04/06/2020 | USA:NAPOLEONVILLE-LA |
| MT536178.1 | 04/06/2020 | USA:LACOMBE-LA |
| MT536182.1 | 04/06/2020 | USA:CHALMETTE-LA |
| MT536186.1 | 04/06/2020 | USA:New-Orleans-LA |
| MT536187.1 | 04/06/2020 | USA:HAMMOND-LA |
| MT536190.1 | 04/06/2020 | USA:METAIRIE-LA |
| MT585082.1 | 04/06/2020 | USA |
| MT585083.1 | 04/06/2020 | USA |
| MT585085.1 | 04/06/2020 | USA |
| MT585086.1 | 04/06/2020 | USA |
| MT585096.1 | 04/06/2020 | USA |
| MT585099.1 | 04/06/2020 | USA |
| MT585101.1 | 04/06/2020 | USA |
| MT585104.1 | 04/06/2020 | USA |
| MT628187.1 | 04/06/2020 | USA:CA |
| MT628259.1 | 04/06/2020 | USA:CA |
| MT628260.1 | 04/06/2020 | USA:CA |
| MT632769.1 | 04/06/2020 | USA:Washington-Snohomish-County |
| MT632770.1 | 04/06/2020 | USA:Washington-Snohomish-County |
| MT632777.1 | 04/06/2020 | USA:Washington-Snohomish-County |
| MT632783.1 | 04/06/2020 | USA:Washington-Snohomish-County |
| MT632792.1 | 04/06/2020 | USA:Washington-Snohomish-County |
| MT641710.1 | 04/06/2020 | Australia:Victoria |
| MT375463.1 | 04/07/2020 | USA:WA |
| MT375465.1 | 04/07/2020 | USA:WA |
| MT375466.1 | 04/07/2020 | USA:WA |
| MT375467.1 | 04/07/2020 | USA:WA |
| MT375468.1 | 04/07/2020 | USA:WA |
| MT375469.1 | 04/07/2020 | USA:CT |
| MT412264.1 | 04/07/2020 | USA:CT |
| MT412265.1 | 04/07/2020 | USA:CT |
| MT412273.1 | 04/07/2020 | USA |

| | | |
|---|---|---|
| MT412277.1 | 04/07/2020 | USA:WA |
| MT412281.1 | 04/07/2020 | USA |
| MT412285.1 | 04/07/2020 | USA:WA |
| MT412286.1 | 04/07/2020 | USA:WA |
| MT412287.1 | 04/07/2020 | USA:WA |
| MT412288.1 | 04/07/2020 | USA:WA |
| MT412289.1 | 04/07/2020 | USA:WA |
| MT429188.1 | 04/07/2020 | USA:Wisconsin |
| MT435083.1 | 04/07/2020 | India:Ahmedabad |
| MT444599.1 | 04/07/2020 | USA |
| MT444600.1 | 04/07/2020 | USA |
| MT444601.1 | 04/07/2020 | USA |
| MT444603.1 | 04/07/2020 | USA |
| MT444605.1 | 04/07/2020 | USA |
| MT446346.1 | 04/07/2020 | USA:LULING-LA |
| MT449647.1 | 04/07/2020 | USA:CA |
| MT449648.1 | 04/07/2020 | USA:CA |
| MT449667.1 | 04/07/2020 | USA:CA |
| MT449668.1 | 04/07/2020 | USA:CA |
| MT449669.1 | 04/07/2020 | USA:CA |
| MT449675.1 | 04/07/2020 | USA:CA |
| MT451739.1 | 04/07/2020 | Australia:Victoria |
| MT451740.1 | 04/07/2020 | Australia:Victoria |
| MT451742.1 | 04/07/2020 | Australia:Victoria |
| MT451743.1 | 04/07/2020 | Australia:Victoria |
| MT451744.1 | 04/07/2020 | Australia:Victoria |
| MT451745.1 | 04/07/2020 | Australia:Victoria |
| MT451746.1 | 04/07/2020 | Australia:Victoria |
| MT451747.1 | 04/07/2020 | Australia:Victoria |
| MT451748.1 | 04/07/2020 | Australia:Victoria |
| MT451749.1 | 04/07/2020 | Australia:Victoria |
| MT451750.1 | 04/07/2020 | Australia:Victoria |
| MT451751.1 | 04/07/2020 | Australia:Victoria |
| MT451752.1 | 04/07/2020 | Australia:Victoria |
| MT451753.1 | 04/07/2020 | Australia:Victoria |
| MT451754.1 | 04/07/2020 | Australia:Victoria |
| MT451822.1 | 04/07/2020 | Australia:Victoria |
| MT477846.1 | 04/07/2020 | USA:AK |
| MT477855.1 | 04/07/2020 | USA:AK |
| MT477856.1 | 04/07/2020 | USA:AK |
| MT477857.1 | 04/07/2020 | USA:AK |
| MT477903.1 | 04/07/2020 | USA:FL |
| MT481936.1 | 04/07/2020 | USA:CA |
| MT499190.1 | 04/07/2020 | USA:CA |
| MT511071.1 | 04/07/2020 | Poland |
| MT511076.1 | 04/07/2020 | Poland |

| | | |
|---|---|---|
| MT511081.1 | 04/07/2020 | Poland |
| MT511673.1 | 04/07/2020 | USA:FL |
| MT511674.1 | 04/07/2020 | USA:FL |
| MT511675.1 | 04/07/2020 | USA:FL |
| MT520181.1 | 04/07/2020 | USA:Massachusetts |
| MT520187.1 | 04/07/2020 | USA:Massachusetts |
| MT520205.1 | 04/07/2020 | USA:Massachusetts |
| MT532437.1 | 04/07/2020 | USA |
| MT535486.1 | 04/07/2020 | USA |
| MT535487.1 | 04/07/2020 | USA |
| MT535488.1 | 04/07/2020 | USA |
| MT535489.1 | 04/07/2020 | USA |
| MT535490.1 | 04/07/2020 | USA |
| MT535491.1 | 04/07/2020 | USA |
| MT535492.1 | 04/07/2020 | USA |
| MT535493.1 | 04/07/2020 | USA |
| MT535494.1 | 04/07/2020 | USA |
| MT535495.1 | 04/07/2020 | USA |
| MT535496.1 | 04/07/2020 | USA |
| MT535497.1 | 04/07/2020 | USA |
| MT535498.1 | 04/07/2020 | USA |
| MT535499.1 | 04/07/2020 | USA |
| MT535500.1 | 04/07/2020 | USA |
| MT535501.1 | 04/07/2020 | USA |
| MT535502.1 | 04/07/2020 | USA |
| MT536177.1 | 04/07/2020 | USA:New-Orleans-LA |
| MT536179.1 | 04/07/2020 | USA:MARRERO-LA |
| MT536180.1 | 04/07/2020 | USA:New-Orleans-LA |
| MT536183.1 | 04/07/2020 | USA:VIOLET-LA |
| MT536189.1 | 04/07/2020 | USA:AVONDALE-LA |
| MT585095.1 | 04/07/2020 | USA |
| MT585097.1 | 04/07/2020 | USA |
| MT612123.1 | 04/07/2020 | Australia:Victoria |
| MT628150.1 | 04/07/2020 | USA:CA |
| MT628261.1 | 04/07/2020 | USA:CA |
| MT631841.1 | 04/07/2020 | USA:CA |
| MT631842.1 | 04/07/2020 | USA:CA |
| MT641494.1 | 04/07/2020 | USA:Washington-Yakima-County |
| MT641537.1 | 04/07/2020 | USA:Washington-King-County |
| MT641564.1 | 04/07/2020 | USA:Washington-Yakima-County |
| MT641578.1 | 04/07/2020 | USA:Washington-Adams-County |
| MT641608.1 | 04/07/2020 | USA:Washington-King-County |
| MT641624.1 | 04/07/2020 | USA:Washington-King-County |
| MT641627.1 | 04/07/2020 | USA:Washington-King-County |
| MT641629.1 | 04/07/2020 | USA:Washington-King-County |
| MT641632.1 | 04/07/2020 | USA:Washington-King-County |

| | | |
|---|---|---|
| MT641730.1 | 04/07/2020 | Australia:Victoria |
| MT375472.1 | 04/08/2020 | USA |
| MT375473.1 | 04/08/2020 | USA:WA |
| MT385416.1 | 04/08/2020 | USA:CA |
| MT385417.1 | 04/08/2020 | USA:CA |
| MT385419.1 | 04/08/2020 | USA:CA |
| MT385427.1 | 04/08/2020 | USA:CA |
| MT385433.1 | 04/08/2020 | USA:CA |
| MT385435.1 | 04/08/2020 | USA:CA |
| MT385438.1 | 04/08/2020 | USA:CA |
| MT385451.1 | 04/08/2020 | USA:CA |
| MT385455.1 | 04/08/2020 | USA:CA |
| MT385467.1 | 04/08/2020 | USA:CA |
| MT385468.1 | 04/08/2020 | USA:CA |
| MT385470.1 | 04/08/2020 | USA:CA |
| MT385471.1 | 04/08/2020 | USA:CA |
| MT385477.1 | 04/08/2020 | USA:CA |
| MT385478.1 | 04/08/2020 | USA:CA |
| MT385485.1 | 04/08/2020 | USA:CA |
| MT385486.1 | 04/08/2020 | USA:CA |
| MT396242.1 | 04/08/2020 | India |
| MT412266.1 | 04/08/2020 | USA |
| MT412267.1 | 04/08/2020 | USA |
| MT412268.1 | 04/08/2020 | USA:WA |
| MT412269.1 | 04/08/2020 | USA:WA |
| MT429185.1 | 04/08/2020 | USA:Wisconsin |
| MT429187.1 | 04/08/2020 | USA:Wisconsin |
| MT429189.1 | 04/08/2020 | USA:Wisconsin |
| MT444602.1 | 04/08/2020 | USA |
| MT444606.1 | 04/08/2020 | USA |
| MT444607.1 | 04/08/2020 | USA |
| MT444608.1 | 04/08/2020 | USA |
| MT444609.1 | 04/08/2020 | USA |
| MT444610.1 | 04/08/2020 | USA |
| MT444611.1 | 04/08/2020 | USA |
| MT446340.1 | 04/08/2020 | USA:KENNER-LA |
| MT446341.1 | 04/08/2020 | USA:KILLONA-LA |
| MT446342.1 | 04/08/2020 | USA:SAINT-ROSE-LA |
| MT446354.1 | 04/08/2020 | USA:MARINGOUIN-LA |
| MT446355.1 | 04/08/2020 | USA:Slidell-LA |
| MT449650.1 | 04/08/2020 | USA:CA |
| MT449651.1 | 04/08/2020 | USA:CA |
| MT449676.1 | 04/08/2020 | USA:CA |
| MT451629.1 | 04/08/2020 | Australia:Victoria |
| MT451755.1 | 04/08/2020 | Australia:Victoria |
| MT451756.1 | 04/08/2020 | Australia:Victoria |

| | | |
|---|---|---|
| MT451757.1 | 04/08/2020 | Australia:Victoria |
| MT451758.1 | 04/08/2020 | Australia:Victoria |
| MT451759.1 | 04/08/2020 | Australia:Victoria |
| MT451760.1 | 04/08/2020 | Australia:Victoria |
| MT451761.1 | 04/08/2020 | Australia:Victoria |
| MT451762.1 | 04/08/2020 | Australia:Victoria |
| MT451763.1 | 04/08/2020 | Australia:Victoria |
| MT451764.1 | 04/08/2020 | Australia:Victoria |
| MT451765.1 | 04/08/2020 | Australia:Victoria |
| MT451766.1 | 04/08/2020 | Australia:Victoria |
| MT451767.1 | 04/08/2020 | Australia:Victoria |
| MT451768.1 | 04/08/2020 | Australia:Victoria |
| MT451770.1 | 04/08/2020 | Australia:Victoria |
| MT451771.1 | 04/08/2020 | Australia:Victoria |
| MT451772.1 | 04/08/2020 | Australia:Victoria |
| MT474137.1 | 04/08/2020 | USA:CA |
| MT477858.1 | 04/08/2020 | USA:AK |
| MT477859.1 | 04/08/2020 | USA:AK |
| MT481935.1 | 04/08/2020 | USA:CA |
| MT506695.1 | 04/08/2020 | USA:Michigan |
| MT506986.1 | 04/08/2020 | USA:FL |
| MT506987.1 | 04/08/2020 | USA:FL |
| MT506990.1 | 04/08/2020 | USA:FL |
| MT507278.1 | 04/08/2020 | USA |
| MT520198.1 | 04/08/2020 | USA:Massachusetts |
| MT520321.1 | 04/08/2020 | USA:Massachusetts |
| MT520352.1 | 04/08/2020 | USA:Massachusetts |
| MT520353.1 | 04/08/2020 | USA:Massachusetts |
| MT520390.1 | 04/08/2020 | USA:Massachusetts |
| MT520398.1 | 04/08/2020 | USA:Massachusetts |
| MT536181.1 | 04/08/2020 | USA:METAIRIE-LA |
| MT536185.1 | 04/08/2020 | USA:New-Orleans-LA |
| MT585098.1 | 04/08/2020 | USA |
| MT628097.1 | 04/08/2020 | USA:CA |
| MT628098.1 | 04/08/2020 | USA:CA |
| MT628262.1 | 04/08/2020 | USA:CA |
| MT641490.1 | 04/08/2020 | USA:Washington-Pierce-County |
| MT641502.1 | 04/08/2020 | USA:Washington-Pierce-County |
| MT641504.1 | 04/08/2020 | USA:Washington-Pierce-County |
| MT641506.1 | 04/08/2020 | USA:Washington-Yakima-County |
| MT641509.1 | 04/08/2020 | USA:Washington-Pierce-County |
| MT641513.1 | 04/08/2020 | USA:Washington-Pierce-County |
| MT641514.1 | 04/08/2020 | USA:Washington-Yakima-County |
| MT641519.1 | 04/08/2020 | USA:Washington-Yakima-County |
| MT641520.1 | 04/08/2020 | USA:Washington-Pierce-County |
| MT641522.1 | 04/08/2020 | USA:Washington-Pierce-County |

| | | |
|---|---|---|
| MT641530.1 | 04/08/2020 | USA:Washington-Yakima-County |
| MT641535.1 | 04/08/2020 | USA:Washington-Pierce-County |
| MT641539.1 | 04/08/2020 | USA:Washington-Pierce-County |
| MT641546.1 | 04/08/2020 | USA:Washington-Yakima-County |
| MT641551.1 | 04/08/2020 | USA:Washington-Yakima-County |
| MT641552.1 | 04/08/2020 | USA:Washington-Yakima-County |
| MT641569.1 | 04/08/2020 | USA:Washington-Jefferson-County |
| MT641572.1 | 04/08/2020 | USA:Washington-King-County |
| MT641581.1 | 04/08/2020 | USA:Washington-Yakima-County |
| MT641590.1 | 04/08/2020 | USA:Washington-Yakima-County |
| MT641591.1 | 04/08/2020 | USA:Washington-Yakima-County |
| MT641593.1 | 04/08/2020 | USA:Washington-Yakima-County |
| MT641620.1 | 04/08/2020 | USA:Washington-Yakima-County |
| MT641642.1 | 04/08/2020 | USA:Washington-Yakima-County |
| MT641731.1 | 04/08/2020 | Australia:Victoria |
| MT432195.1 | 04/09/2020 | USA:East-Feliciana-Parish-Louisiana |
| MT444612.1 | 04/09/2020 | USA |
| MT449652.1 | 04/09/2020 | USA:CA |
| MT449674.1 | 04/09/2020 | USA:CA |
| MT451773.1 | 04/09/2020 | Australia:Victoria |
| MT451774.1 | 04/09/2020 | Australia:Victoria |
| MT451775.1 | 04/09/2020 | Australia:Victoria |
| MT451776.1 | 04/09/2020 | Australia:Victoria |
| MT451777.1 | 04/09/2020 | Australia:Victoria |
| MT451778.1 | 04/09/2020 | Australia:Victoria |
| MT451779.1 | 04/09/2020 | Australia:Victoria |
| MT451780.1 | 04/09/2020 | Australia:Victoria |
| MT451782.1 | 04/09/2020 | Australia:Victoria |
| MT451823.1 | 04/09/2020 | Australia:Victoria |
| MT451824.1 | 04/09/2020 | Australia:Victoria |
| MT451825.1 | 04/09/2020 | Australia:Victoria |
| MT477860.1 | 04/09/2020 | USA:AK |
| MT499179.1 | 04/09/2020 | USA:CA |
| MT506988.1 | 04/09/2020 | USA:FL |
| MT506989.1 | 04/09/2020 | USA:FL |
| MT506991.1 | 04/09/2020 | USA:FL |
| MT511676.1 | 04/09/2020 | USA:FL |
| MT511677.1 | 04/09/2020 | USA:FL |
| MT511678.1 | 04/09/2020 | USA:FL |
| MT511679.1 | 04/09/2020 | USA:FL |
| MT511680.1 | 04/09/2020 | USA:FL |
| MT511681.1 | 04/09/2020 | USA:FL |
| MT511683.1 | 04/09/2020 | USA:FL |
| MT620771.1 | 04/09/2020 | USA |
| MT628100.1 | 04/09/2020 | USA:CA |
| MT628263.1 | 04/09/2020 | USA:CA |

| | | |
|---|---|---|
| MT628264.1 | 04/09/2020 | USA:CA |
| MT641485.1 | 04/09/2020 | USA:Washington-Yakima-County |
| MT641571.1 | 04/09/2020 | USA:Washington-King-County |
| MT641609.1 | 04/09/2020 | USA:Washington-Snohomish-County |
| MT641617.1 | 04/09/2020 | USA:Washington-King-County |
| MT641623.1 | 04/09/2020 | USA:Washington-Yakima-County |
| MT641625.1 | 04/09/2020 | USA:Washington-King-County |
| MT641631.1 | 04/09/2020 | USA:Washington-Yakima-County |
| MT641733.1 | 04/09/2020 | Australia:Victoria |
| MT641735.1 | 04/09/2020 | Australia:Victoria |
| MT641764.1 | 04/09/2020 | Australia:Victoria |
| MT642241.1 | 04/09/2020 | USA:Washington |
| MT642380.1 | 04/09/2020 | USA:Washington |
| MT642417.1 | 04/09/2020 | USA:Washington |
| MT396245.1 | 04/10/2020 | India |
| MT396246.1 | 04/10/2020 | India |
| MT429184.1 | 04/10/2020 | USA:Wisconsin |
| MT444613.1 | 04/10/2020 | USA |
| MT444614.1 | 04/10/2020 | USA |
| MT444615.1 | 04/10/2020 | USA |
| MT449653.1 | 04/10/2020 | USA:CA |
| MT451769.1 | 04/10/2020 | Australia:Victoria |
| MT451783.1 | 04/10/2020 | Australia:Victoria |
| MT451784.1 | 04/10/2020 | Australia:Victoria |
| MT451785.1 | 04/10/2020 | Australia:Victoria |
| MT451786.1 | 04/10/2020 | Australia:Victoria |
| MT451826.1 | 04/10/2020 | Australia:Victoria |
| MT451827.1 | 04/10/2020 | Australia:Victoria |
| MT451828.1 | 04/10/2020 | Australia:Victoria |
| MT511682.1 | 04/10/2020 | USA:FL |
| MT511684.1 | 04/10/2020 | USA:FL |
| MT511685.1 | 04/10/2020 | USA:FL |
| MT511686.1 | 04/10/2020 | USA:FL |
| MT511687.1 | 04/10/2020 | USA:FL |
| MT511688.1 | 04/10/2020 | USA:FL |
| MT511690.1 | 04/10/2020 | USA:FL |
| MT511691.1 | 04/10/2020 | USA:FL |
| MT534303.1 | 04/10/2020 | USA:CA |
| MT534304.1 | 04/10/2020 | USA:CA |
| MT628101.1 | 04/10/2020 | USA:CA |
| MT641497.1 | 04/10/2020 | USA:Washington-Yakima-County |
| MT641526.1 | 04/10/2020 | USA:Washington-Yakima-County |
| MT641532.1 | 04/10/2020 | USA:Washington-Pierce-County |
| MT641599.1 | 04/10/2020 | USA:Washington-Pierce-County |
| MT641604.1 | 04/10/2020 | USA:Washington-Pierce-County |
| MT641607.1 | 04/10/2020 | USA:Washington-King-County |

| | | |
|---|---|---|
| MT641634.1 | 04/10/2020 | USA:Washington-Benton-County |
| MT641736.1 | 04/10/2020 | Australia:Victoria |
| MT641737.1 | 04/10/2020 | Australia:Victoria |
| MT641738.1 | 04/10/2020 | Australia:Victoria |
| MT641739.1 | 04/10/2020 | Australia:Victoria |
| MT641740.1 | 04/10/2020 | Australia:Victoria |
| MT641741.1 | 04/10/2020 | Australia:Victoria |
| MT642395.1 | 04/10/2020 | USA:Washington |
| MT642419.1 | 04/10/2020 | USA:Washington |
| MT449654.1 | 04/11/2020 | USA:CA |
| MT451787.1 | 04/11/2020 | Australia:Victoria |
| MT451788.1 | 04/11/2020 | Australia:Victoria |
| MT451789.1 | 04/11/2020 | Australia:Victoria |
| MT451790.1 | 04/11/2020 | Australia:Victoria |
| MT451791.1 | 04/11/2020 | Australia:Victoria |
| MT451829.1 | 04/11/2020 | Australia:Victoria |
| MT451830.1 | 04/11/2020 | Australia:Victoria |
| MT451831.1 | 04/11/2020 | Australia:Victoria |
| MT451832.1 | 04/11/2020 | Australia:Victoria |
| MT451833.1 | 04/11/2020 | Australia:Victoria |
| MT451834.1 | 04/11/2020 | Australia:Victoria |
| MT499208.1 | 04/11/2020 | Poland |
| MT499209.1 | 04/11/2020 | Poland |
| MT499210.1 | 04/11/2020 | Poland |
| MT509505.1 | 04/11/2020 | India:Ahmedabad |
| MT509509.1 | 04/11/2020 | India:Ahmedabad |
| MT511692.1 | 04/11/2020 | USA:FL |
| MT511693.1 | 04/11/2020 | USA:FL |
| MT511694.1 | 04/11/2020 | USA:FL |
| MT511695.1 | 04/11/2020 | USA:FL |
| MT511696.1 | 04/11/2020 | USA:FL |
| MT511697.1 | 04/11/2020 | USA:FL |
| MT511698.1 | 04/11/2020 | USA:FL |
| MT511699.1 | 04/11/2020 | USA:FL |
| MT511700.1 | 04/11/2020 | USA:FL |
| MT534305.1 | 04/11/2020 | USA:CA |
| MT534308.1 | 04/11/2020 | USA:CA |
| MT590600.1 | 04/11/2020 | Taiwan |
| MT612281.1 | 04/11/2020 | Timor-Leste |
| MT612289.1 | 04/11/2020 | Timor-Leste |
| MT628104.1 | 04/11/2020 | USA:CA |
| MT628152.1 | 04/11/2020 | USA:CA |
| MT628153.1 | 04/11/2020 | USA:CA |
| MT628265.1 | 04/11/2020 | USA:CA |
| MT641501.1 | 04/11/2020 | USA:Washington |
| MT641508.1 | 04/11/2020 | USA:Washington-Asotin-County |

| | | |
|---|---|---|
| MT641536.1 | 04/11/2020 | USA:Washington-Pierce-County |
| MT641543.1 | 04/11/2020 | USA:Washington-Pierce-County |
| MT641548.1 | 04/11/2020 | USA:Washington-Yakima-County |
| MT641554.1 | 04/11/2020 | USA:Washington-Yakima-County |
| MT641556.1 | 04/11/2020 | USA:Washington |
| MT641568.1 | 04/11/2020 | USA:Washington-Asotin-County |
| MT641580.1 | 04/11/2020 | USA:Washington-King-County |
| MT641588.1 | 04/11/2020 | USA:Washington-Whatcom-County |
| MT641592.1 | 04/11/2020 | USA:Washington-Yakima-County |
| MT641600.1 | 04/11/2020 | USA:Washington-Pierce-County |
| MT641628.1 | 04/11/2020 | USA:Washington-Yakima-County |
| MT641636.1 | 04/11/2020 | USA:Washington-Yakima-County |
| MT641742.1 | 04/11/2020 | Australia:Victoria |
| MT641743.1 | 04/11/2020 | Australia:Victoria |
| MT641744.1 | 04/11/2020 | Australia:Victoria |
| MT641745.1 | 04/11/2020 | Australia:Victoria |
| MT641746.1 | 04/11/2020 | Australia:Victoria |
| MT641747.1 | 04/11/2020 | Australia:Victoria |
| MT641748.1 | 04/11/2020 | Australia:Victoria |
| MT641766.1 | 04/11/2020 | Timor-Leste |
| MT641767.1 | 04/11/2020 | Timor-Leste |
| MT641768.1 | 04/11/2020 | Timor-Leste |
| MT429186.1 | 04/12/2020 | USA:Wisconsin |
| MT449655.1 | 04/12/2020 | USA:CA |
| MT451835.1 | 04/12/2020 | Australia:Victoria |
| MT451836.1 | 04/12/2020 | Australia:Victoria |
| MT451837.1 | 04/12/2020 | Australia:Victoria |
| MT474138.1 | 04/12/2020 | USA:CA |
| MT477861.1 | 04/12/2020 | USA:AK |
| MT511701.1 | 04/12/2020 | USA:FL |
| MT534317.1 | 04/12/2020 | USA:CA |
| MT534318.1 | 04/12/2020 | USA:CA |
| MT534320.1 | 04/12/2020 | USA:CA |
| MT534321.1 | 04/12/2020 | USA:CA |
| MT534323.1 | 04/12/2020 | USA:CA |
| MT534324.1 | 04/12/2020 | USA:CA |
| MT534329.1 | 04/12/2020 | USA:CA |
| MT534334.1 | 04/12/2020 | USA:CA |
| MT534336.1 | 04/12/2020 | USA:CA |
| MT622321.1 | 04/12/2020 | Italy |
| MT627752.1 | 04/12/2020 | USA:Washington-King-County |
| MT627758.1 | 04/12/2020 | USA:Washington-King-County |
| MT628105.1 | 04/12/2020 | USA:CA |
| MT628108.1 | 04/12/2020 | USA:CA |
| MT628227.1 | 04/12/2020 | USA:CA |
| MT628228.1 | 04/12/2020 | USA:CA |

| | | |
|---|---|---|
| MT641541.1 | 04/12/2020 | USA:Washington-Whatcom-County |
| MT641542.1 | 04/12/2020 | USA:Washington-Yakima-County |
| MT641577.1 | 04/12/2020 | USA:Washington-Yakima-County |
| MT641586.1 | 04/12/2020 | USA:Washington-Yakima-County |
| MT641603.1 | 04/12/2020 | USA:Washington-Yakima-County |
| MT641750.1 | 04/12/2020 | Australia:Victoria |
| MT641751.1 | 04/12/2020 | Australia:Victoria |
| MT641762.1 | 04/12/2020 | Australia:Victoria |
| MT641763.1 | 04/12/2020 | Australia:Victoria |
| MT641769.1 | 04/12/2020 | Timor-Leste |
| MT641771.1 | 04/12/2020 | Timor-Leste |
| MT641772.1 | 04/12/2020 | Timor-Leste |
| MT641773.1 | 04/12/2020 | Timor-Leste |
| MT641774.1 | 04/12/2020 | Timor-Leste |
| MT435079.1 | 04/13/2020 | India:Ahmedabad |
| MT435080.1 | 04/13/2020 | India:Ahmedabad |
| MT435081.1 | 04/13/2020 | India:Ahmedabad |
| MT435082.1 | 04/13/2020 | India:Ahmedabad |
| MT450872.1 | 04/13/2020 | Serbia |
| MT451838.1 | 04/13/2020 | Australia:Victoria |
| MT451839.1 | 04/13/2020 | Australia:Victoria |
| MT451840.1 | 04/13/2020 | Australia:Victoria |
| MT536954.1 | 04/13/2020 | USA |
| MT536955.1 | 04/13/2020 | USA |
| MT536956.1 | 04/13/2020 | USA |
| MT536957.1 | 04/13/2020 | USA |
| MT536958.1 | 04/13/2020 | USA |
| MT536959.1 | 04/13/2020 | USA |
| MT536960.1 | 04/13/2020 | USA |
| MT536961.1 | 04/13/2020 | USA |
| MT536962.1 | 04/13/2020 | USA |
| MT536963.1 | 04/13/2020 | USA |
| MT536964.1 | 04/13/2020 | USA |
| MT536965.1 | 04/13/2020 | USA |
| MT536966.1 | 04/13/2020 | USA |
| MT536967.1 | 04/13/2020 | USA |
| MT536968.1 | 04/13/2020 | USA |
| MT536969.1 | 04/13/2020 | USA |
| MT627757.1 | 04/13/2020 | USA:Washington-King-County |
| MT627926.1 | 04/13/2020 | USA:Washington |
| MT627928.1 | 04/13/2020 | USA:Washington |
| MT628110.1 | 04/13/2020 | USA:CA |
| MT641488.1 | 04/13/2020 | USA:Washington-Snohomish-County |
| MT641496.1 | 04/13/2020 | USA:Washington-King-County |
| MT641499.1 | 04/13/2020 | USA:Washington-Asotin-County |
| MT641500.1 | 04/13/2020 | USA:Washington-Snohomish-County |

| | | |
|---|---|---|
| MT641515.1 | 04/13/2020 | USA:Washington-Yakima-County |
| MT641516.1 | 04/13/2020 | USA:Washington-Asotin-County |
| MT641518.1 | 04/13/2020 | USA:Washington-Yakima-County |
| MT641525.1 | 04/13/2020 | USA:Washington-Snohomish-County |
| MT641527.1 | 04/13/2020 | USA:Washington-Yakima-County |
| MT641533.1 | 04/13/2020 | USA:Washington-Yakima-County |
| MT641534.1 | 04/13/2020 | USA:Washington-Yakima-County |
| MT641544.1 | 04/13/2020 | USA:Washington-Yakima-County |
| MT641545.1 | 04/13/2020 | USA:Washington-Yakima-County |
| MT641565.1 | 04/13/2020 | USA:Washington-Snohomish-County |
| MT641566.1 | 04/13/2020 | USA:Washington-Yakima-County |
| MT641574.1 | 04/13/2020 | USA:Washington-Yakima-County |
| MT641579.1 | 04/13/2020 | USA:Washington-Yakima-County |
| MT641583.1 | 04/13/2020 | USA:Washington-Yakima-County |
| MT641643.1 | 04/13/2020 | USA:Washington-Yakima-County |
| MT641752.1 | 04/13/2020 | Australia:Victoria |
| MT641754.1 | 04/13/2020 | Australia:Victoria |
| MT641757.1 | 04/13/2020 | Australia:Victoria |
| MT396244.1 | 04/14/2020 | India |
| MT396248.1 | 04/14/2020 | India |
| MT435084.1 | 04/14/2020 | India:Ahmedabad |
| MT444619.1 | 04/14/2020 | USA |
| MT444623.1 | 04/14/2020 | USA |
| MT444627.1 | 04/14/2020 | USA |
| MT444629.1 | 04/14/2020 | USA |
| MT444631.1 | 04/14/2020 | USA |
| MT444633.1 | 04/14/2020 | USA |
| MT451841.1 | 04/14/2020 | Australia:Victoria |
| MT451842.1 | 04/14/2020 | Australia:Victoria |
| MT506890.1 | 04/14/2020 | USA:Madison-Wisconsin |
| MT506892.1 | 04/14/2020 | USA:Illinois |
| MT506895.1 | 04/14/2020 | USA:Mount-Horeb-Wisconsin |
| MT506896.1 | 04/14/2020 | USA:Illinois |
| MT506905.1 | 04/14/2020 | USA:Illinois |
| MT506907.1 | 04/14/2020 | USA:Illinois |
| MT533207.1 | 04/14/2020 | USA:CA |
| MT534311.1 | 04/14/2020 | USA:CA |
| MT534312.1 | 04/14/2020 | USA:CA |
| MT534313.1 | 04/14/2020 | USA:CA |
| MT534314.1 | 04/14/2020 | USA:CA |
| MT534315.1 | 04/14/2020 | USA:CA |
| MT534316.1 | 04/14/2020 | USA:CA |
| MT612111.1 | 04/14/2020 | Australia:Victoria |
| MT612114.1 | 04/14/2020 | Australia:Victoria |
| MT612115.1 | 04/14/2020 | Australia:Victoria |
| MT628189.1 | 04/14/2020 | USA:CA |

| | | |
|---|---|---|
| MT628266.1 | 04/14/2020 | USA:CA |
| MT628267.1 | 04/14/2020 | USA:CA |
| MT641489.1 | 04/14/2020 | USA:Washington-Yakima-County |
| MT641498.1 | 04/14/2020 | USA:Washington-Snohomish-County |
| MT641517.1 | 04/14/2020 | USA:Washington-Yakima-County |
| MT641529.1 | 04/14/2020 | USA:Washington-Yakima-County |
| MT641531.1 | 04/14/2020 | USA:Washington-King-County |
| MT641547.1 | 04/14/2020 | USA:Washington-Yakima-County |
| MT641558.1 | 04/14/2020 | USA:Washington-Pierce-County |
| MT641559.1 | 04/14/2020 | USA:Washington-Pierce-County |
| MT641561.1 | 04/14/2020 | USA:Washington-Yakima-County |
| MT641563.1 | 04/14/2020 | USA:Washington-Pierce-County |
| MT641573.1 | 04/14/2020 | USA:Washington-King-County |
| MT641575.1 | 04/14/2020 | USA:Washington-King-County |
| MT641576.1 | 04/14/2020 | USA:Washington-Pierce-County |
| MT641589.1 | 04/14/2020 | USA:Washington-Yakima-County |
| MT641594.1 | 04/14/2020 | USA:Washington-Yakima-County |
| MT641601.1 | 04/14/2020 | USA:Washington-Yakima-County |
| MT641602.1 | 04/14/2020 | USA:Washington-King-County |
| MT641611.1 | 04/14/2020 | USA:Washington-King-County |
| MT641612.1 | 04/14/2020 | USA:Washington-Snohomish-County |
| MT641614.1 | 04/14/2020 | USA:Washington-King-County |
| MT641615.1 | 04/14/2020 | USA:Washington-King-County |
| MT641618.1 | 04/14/2020 | USA:Washington-King-County |
| MT641621.1 | 04/14/2020 | USA:Washington-King-County |
| MT641622.1 | 04/14/2020 | USA:Washington-King-County |
| MT641626.1 | 04/14/2020 | USA:Washington-King-County |
| MT641630.1 | 04/14/2020 | USA:Washington-Yakima-County |
| MT641635.1 | 04/14/2020 | USA:Washington-King-County |
| MT641641.1 | 04/14/2020 | USA:Washington-Yakima-County |
| MT641758.1 | 04/14/2020 | Australia:Victoria |
| MT641759.1 | 04/14/2020 | Australia:Victoria |
| MT641760.1 | 04/14/2020 | Australia:Victoria |
| MT641761.1 | 04/14/2020 | Australia:Victoria |
| MT641776.1 | 04/14/2020 | Timor-Leste |
| MT642302.1 | 04/14/2020 | USA:Washington |
| MT444616.1 | 04/15/2020 | USA |
| MT444617.1 | 04/15/2020 | USA |
| MT444618.1 | 04/15/2020 | USA |
| MT444620.1 | 04/15/2020 | USA |
| MT444621.1 | 04/15/2020 | USA |
| MT444622.1 | 04/15/2020 | USA |
| MT444624.1 | 04/15/2020 | USA |
| MT444625.1 | 04/15/2020 | USA |
| MT444626.1 | 04/15/2020 | USA |
| MT444628.1 | 04/15/2020 | USA |

| | | |
|---|---|---|
| MT444630.1 | 04/15/2020 | USA |
| MT444632.1 | 04/15/2020 | USA |
| MT444634.1 | 04/15/2020 | USA |
| MT444635.1 | 04/15/2020 | USA |
| MT506887.1 | 04/15/2020 | USA:Dane-County-Wisconsin |
| MT506888.1 | 04/15/2020 | USA:Illinois |
| MT506897.1 | 04/15/2020 | USA:Illinois |
| MT506904.1 | 04/15/2020 | USA:Illinois |
| MT506992.1 | 04/15/2020 | USA:FL |
| MT506993.1 | 04/15/2020 | USA:FL |
| MT506994.1 | 04/15/2020 | USA:FL |
| MT506995.1 | 04/15/2020 | USA:FL |
| MT506996.1 | 04/15/2020 | USA:FL |
| MT536971.1 | 04/15/2020 | USA |
| MT560530.1 | 04/15/2020 | Turkey |
| MT577596.1 | 04/15/2020 | USA:FL |
| MT577597.1 | 04/15/2020 | USA:FL |
| MT577598.1 | 04/15/2020 | USA:FL |
| MT584981.1 | 04/15/2020 | USA:Michigan |
| MT584982.1 | 04/15/2020 | USA:Michigan |
| MT612129.1 | 04/15/2020 | Australia:Victoria |
| MT627745.1 | 04/15/2020 | USA:Washington-King-County |
| MT627923.1 | 04/15/2020 | USA:Washington |
| MT628070.1 | 04/15/2020 | USA:Washington |
| MT641486.1 | 04/15/2020 | USA:Washington-Snohomish-County |
| MT641495.1 | 04/15/2020 | USA:Washington-Yakima-County |
| MT641510.1 | 04/15/2020 | USA:Washington-Yakima-County |
| MT641521.1 | 04/15/2020 | USA:Washington-Asotin-County |
| MT641528.1 | 04/15/2020 | USA:Washington-Snohomish-County |
| MT641538.1 | 04/15/2020 | USA:Washington-Yakima-County |
| MT641549.1 | 04/15/2020 | USA:Washington-Yakima-County |
| MT641560.1 | 04/15/2020 | USA:Washington-Yakima-County |
| MT641562.1 | 04/15/2020 | USA:Washington-Yakima-County |
| MT641582.1 | 04/15/2020 | USA:Washington-Yakima-County |
| MT641605.1 | 04/15/2020 | USA:Washington-Yakima-County |
| MT641613.1 | 04/15/2020 | USA:Washington-Yakima-County |
| MT641619.1 | 04/15/2020 | USA:Washington-Yakima-County |
| MT641633.1 | 04/15/2020 | USA:Washington-Yakima-County |
| MT642428.1 | 04/15/2020 | USA:Washington-Grant-County |
| MT449657.1 | 04/16/2020 | USA:CA |
| MT449658.1 | 04/16/2020 | USA:CA |
| MT506903.1 | 04/16/2020 | USA:Illinois |
| MT506997.1 | 04/16/2020 | USA:FL |
| MT506998.1 | 04/16/2020 | USA:FL |
| MT506999.1 | 04/16/2020 | USA:FL |
| MT507000.1 | 04/16/2020 | USA:FL |

| | | |
|---|---|---|
| MT507001.1 | 04/16/2020 | USA:FL |
| MT507002.1 | 04/16/2020 | USA:FL |
| MT507003.1 | 04/16/2020 | USA:FL |
| MT507004.1 | 04/16/2020 | USA:FL |
| MT507005.1 | 04/16/2020 | USA:FL |
| MT507006.1 | 04/16/2020 | USA:FL |
| MT507007.1 | 04/16/2020 | USA:FL |
| MT507008.1 | 04/16/2020 | USA:FL |
| MT507009.1 | 04/16/2020 | USA:FL |
| MT507010.1 | 04/16/2020 | USA:FL |
| MT533208.1 | 04/16/2020 | USA:CA |
| MT533209.1 | 04/16/2020 | USA:CA |
| MT533210.1 | 04/16/2020 | USA:CA |
| MT536970.1 | 04/16/2020 | USA |
| MT536973.1 | 04/16/2020 | USA |
| MT536975.1 | 04/16/2020 | USA |
| MT560531.1 | 04/16/2020 | Turkey |
| MT577599.1 | 04/16/2020 | USA:FL |
| MT577600.1 | 04/16/2020 | USA:FL |
| MT577601.1 | 04/16/2020 | USA:FL |
| MT577602.1 | 04/16/2020 | USA:FL |
| MT584980.1 | 04/16/2020 | USA:Michigan |
| MT584987.1 | 04/16/2020 | USA:Michigan |
| MT612122.1 | 04/16/2020 | Australia:Victoria |
| MT612125.1 | 04/16/2020 | Australia:Victoria |
| MT628071.1 | 04/16/2020 | USA:Washington-King-County |
| MT628268.1 | 04/16/2020 | USA:CA |
| MT641492.1 | 04/16/2020 | USA:Washington-Pierce-County |
| MT641493.1 | 04/16/2020 | USA:Washington-Snohomish-County |
| MT641512.1 | 04/16/2020 | USA:Washington-Yakima-County |
| MT641523.1 | 04/16/2020 | USA:Washington-Whatcom-County |
| MT641524.1 | 04/16/2020 | USA:Washington-Pierce-County |
| MT641553.1 | 04/16/2020 | USA:Washington-Yakima-County |
| MT641555.1 | 04/16/2020 | USA:Washington |
| MT641557.1 | 04/16/2020 | USA:Washington-Pierce-County |
| MT641567.1 | 04/16/2020 | USA:Washington-Snohomish-County |
| MT641570.1 | 04/16/2020 | USA:Washington-King-County |
| MT641584.1 | 04/16/2020 | USA:Washington-Yakima-County |
| MT641585.1 | 04/16/2020 | USA:Washington-Snohomish-County |
| MT641587.1 | 04/16/2020 | USA:Washington-Yakima-County |
| MT641595.1 | 04/16/2020 | USA:Washington-Yakima-County |
| MT641596.1 | 04/16/2020 | USA:Washington-Yakima-County |
| MT641598.1 | 04/16/2020 | USA:Washington-Yakima-County |
| MT641606.1 | 04/16/2020 | USA:Washington-Pierce-County |
| MT641610.1 | 04/16/2020 | USA:Washington-Yakima-County |
| MT642068.1 | 04/16/2020 | USA:Washington-Snohomish-County |

| | | |
|---|---|---|
| MT642359.1 | 04/16/2020 | USA:Washington-Pierce-County |
| MT642383.1 | 04/16/2020 | USA:Washington-Pierce-County |
| MT642405.1 | 04/16/2020 | USA:Washington-Pierce-County |
| MT434759.1 | 04/17/2020 | India |
| MT434760.1 | 04/17/2020 | India |
| MT506894.1 | 04/17/2020 | USA:Illinois |
| MT506901.1 | 04/17/2020 | USA:Mount-Horeb-Wisconsin |
| MT506902.1 | 04/17/2020 | USA:Illinois |
| MT507012.1 | 04/17/2020 | USA:FL |
| MT507013.1 | 04/17/2020 | USA:FL |
| MT507014.1 | 04/17/2020 | USA:FL |
| MT507015.1 | 04/17/2020 | USA:FL |
| MT507016.1 | 04/17/2020 | USA:FL |
| MT507017.1 | 04/17/2020 | USA:FL |
| MT507018.1 | 04/17/2020 | USA:FL |
| MT507019.1 | 04/17/2020 | USA:FL |
| MT507020.1 | 04/17/2020 | USA:FL |
| MT507021.1 | 04/17/2020 | USA:FL |
| MT507022.1 | 04/17/2020 | USA:FL |
| MT507023.1 | 04/17/2020 | USA:FL |
| MT507024.1 | 04/17/2020 | USA:FL |
| MT532438.1 | 04/17/2020 | USA |
| MT536972.1 | 04/17/2020 | USA |
| MT536974.1 | 04/17/2020 | USA |
| MT536976.1 | 04/17/2020 | USA |
| MT536977.1 | 04/17/2020 | USA |
| MT577603.1 | 04/17/2020 | USA:FL |
| MT577604.1 | 04/17/2020 | USA:FL |
| MT577605.1 | 04/17/2020 | USA:FL |
| MT577606.1 | 04/17/2020 | USA:FL |
| MT584983.1 | 04/17/2020 | USA:Michigan |
| MT584984.1 | 04/17/2020 | USA:Michigan |
| MT584985.1 | 04/17/2020 | USA:Michigan |
| MT584986.1 | 04/17/2020 | USA:Michigan |
| MT584988.1 | 04/17/2020 | USA:Michigan |
| MT584989.1 | 04/17/2020 | USA:Michigan |
| MT584990.1 | 04/17/2020 | USA:Michigan |
| MT584992.1 | 04/17/2020 | USA:Michigan |
| MT584993.1 | 04/17/2020 | USA:Michigan |
| MT584994.1 | 04/17/2020 | USA:Michigan |
| MT612131.1 | 04/17/2020 | Australia:Victoria |
| MT612133.1 | 04/17/2020 | Australia:Victoria |
| MT620770.1 | 04/17/2020 | USA |
| MT628037.1 | 04/17/2020 | USA:Washington-King-County |
| MT628038.1 | 04/17/2020 | USA:Washington-Pierce-County |
| MT628041.1 | 04/17/2020 | USA:Washington-Yakima-County |

| | | |
|---|---|---|
| MT628045.1 | 04/17/2020 | USA:Washington-Yakima-County |
| MT628049.1 | 04/17/2020 | USA:Washington-Snohomish-County |
| MT628050.1 | 04/17/2020 | USA:Washington-Pierce-County |
| MT628055.1 | 04/17/2020 | USA:Washington-Snohomish-County |
| MT628059.1 | 04/17/2020 | USA:Washington-Yakima-County |
| MT628060.1 | 04/17/2020 | USA:Washington-Snohomish-County |
| MT628073.1 | 04/17/2020 | USA:Washington-Yakima-County |
| MT628081.1 | 04/17/2020 | USA:Washington-Yakima-County |
| MT628269.1 | 04/17/2020 | USA:CA |
| MT641505.1 | 04/17/2020 | USA:Washington-Snohomish-County |
| MT641511.1 | 04/17/2020 | USA:Washington-Yakima-County |
| MT641550.1 | 04/17/2020 | USA:Washington-Snohomish-County |
| MT641639.1 | 04/17/2020 | USA:Washington-Snohomish-County |
| MT641640.1 | 04/17/2020 | USA:Washington-Yakima-County |
| MT641644.1 | 04/17/2020 | USA:Washington-Snohomish-County |
| MT642229.1 | 04/17/2020 | USA:Washington |
| MT642254.1 | 04/17/2020 | USA:Washington |
| MT642287.1 | 04/17/2020 | USA:Washington |
| MT642327.1 | 04/17/2020 | USA:Washington |
| MT642355.1 | 04/17/2020 | USA:Washington |
| MT476385.1 | 04/18/2020 | Bangladesh |
| MT506583.1 | 04/18/2020 | USA:Michigan |
| MT506886.1 | 04/18/2020 | USA:Oregon-Wisconsin |
| MT506889.1 | 04/18/2020 | USA:Illinois |
| MT506891.1 | 04/18/2020 | USA:Illinois |
| MT506898.1 | 04/18/2020 | USA:Verona-Wisconsin |
| MT506899.1 | 04/18/2020 | USA:Illinois |
| MT506900.1 | 04/18/2020 | USA:Illinois |
| MT584991.1 | 04/18/2020 | USA:Michigan |
| MT584995.1 | 04/18/2020 | USA:Michigan |
| MT612135.1 | 04/18/2020 | Australia:Victoria |
| MT612293.1 | 04/18/2020 | Timor-Leste |
| MT627760.1 | 04/18/2020 | USA:Washington-King-County |
| MT628043.1 | 04/18/2020 | USA:Washington-Pierce-County |
| MT628046.1 | 04/18/2020 | USA:Washington-Pierce-County |
| MT628048.1 | 04/18/2020 | USA:Washington-Pierce-County |
| MT628052.1 | 04/18/2020 | USA:Washington-Pierce-County |
| MT628058.1 | 04/18/2020 | USA:Washington-Yakima-County |
| MT628062.1 | 04/18/2020 | USA:Washington-Yakima-County |
| MT628064.1 | 04/18/2020 | USA:Washington-Yakima-County |
| MT628065.1 | 04/18/2020 | USA:Washington-Yakima-County |
| MT628077.1 | 04/18/2020 | USA:Washington-Pierce-County |
| MT628113.1 | 04/18/2020 | USA:CA |
| MT628270.1 | 04/18/2020 | USA:CA |
| MT631784.1 | 04/18/2020 | USA:CA |
| MT631786.1 | 04/18/2020 | USA:CA |

| | | |
|---|---|---|
| MT631787.1 | 04/18/2020 | USA:CA |
| MT631788.1 | 04/18/2020 | USA:CA |
| MT631789.1 | 04/18/2020 | USA:CA |
| MT631790.1 | 04/18/2020 | USA:CA |
| MT641777.1 | 04/18/2020 | Timor-Leste |
| MT642073.1 | 04/18/2020 | USA:Washington-Snohomish-County |
| MT642416.1 | 04/18/2020 | USA:Washington |
| MT506696.1 | 04/19/2020 | USA:Michigan |
| MT517436.1 | 04/19/2020 | Taiwan |
| MT517437.1 | 04/19/2020 | Taiwan |
| MT612136.1 | 04/19/2020 | Australia:Victoria |
| MT627763.1 | 04/19/2020 | USA:Washington-King-County |
| MT627766.1 | 04/19/2020 | USA:Washington-King-County |
| MT628039.1 | 04/19/2020 | USA:Washington-Pierce-County |
| MT628040.1 | 04/19/2020 | USA:Washington-Yakima-County |
| MT628063.1 | 04/19/2020 | USA:Washington-Yakima-County |
| MT628079.1 | 04/19/2020 | USA:Washington-Yakima-County |
| MT628082.1 | 04/19/2020 | USA:Washington-Yakima-County |
| MT628116.1 | 04/19/2020 | USA:CA |
| MT628154.1 | 04/19/2020 | USA:CA |
| MT628155.1 | 04/19/2020 | USA:CA |
| MT449660.1 | 04/20/2020 | USA:CA |
| MT506582.1 | 04/20/2020 | USA:Michigan |
| MT506584.1 | 04/20/2020 | USA:Michigan |
| MT507025.1 | 04/20/2020 | USA:FL |
| MT507027.1 | 04/20/2020 | USA:FL |
| MT577607.1 | 04/20/2020 | USA:FL |
| MT612145.1 | 04/20/2020 | Australia:Victoria |
| MT620767.1 | 04/20/2020 | USA |
| MT627750.1 | 04/20/2020 | USA:Washington-King-County |
| MT627755.1 | 04/20/2020 | USA:Washington-King-County |
| MT628042.1 | 04/20/2020 | USA:Washington-Asotin-County |
| MT628053.1 | 04/20/2020 | USA:Washington-Yakima-County |
| MT628075.1 | 04/20/2020 | USA:Washington-Pierce-County |
| MT628078.1 | 04/20/2020 | USA:Washington-Yakima-County |
| MT628119.1 | 04/20/2020 | USA:CA |
| MT628121.1 | 04/20/2020 | USA:CA |
| MT628123.1 | 04/20/2020 | USA:CA |
| MT628208.1 | 04/20/2020 | USA:CA |
| MT628271.1 | 04/20/2020 | USA:CA |
| MT642378.1 | 04/20/2020 | USA:Washington-Franklin-County |
| MT434757.1 | 04/21/2020 | India |
| MT435086.1 | 04/21/2020 | India:Mansa |
| MT449659.1 | 04/21/2020 | USA:CA |
| MT449663.1 | 04/21/2020 | USA:CA |
| MT449664.1 | 04/21/2020 | USA:CA |

| | | |
|---|---|---|
| MT449665.1 | 04/21/2020 | USA:CA |
| MT507029.1 | 04/21/2020 | USA:FL |
| MT507030.1 | 04/21/2020 | USA:FL |
| MT507031.1 | 04/21/2020 | USA:FL |
| MT532436.1 | 04/21/2020 | USA |
| MT577608.1 | 04/21/2020 | USA:FL |
| MT577609.1 | 04/21/2020 | USA:FL |
| MT577610.1 | 04/21/2020 | USA:FL |
| MT577611.1 | 04/21/2020 | USA:FL |
| MT612138.1 | 04/21/2020 | Australia:Victoria |
| MT612139.1 | 04/21/2020 | Australia:Victoria |
| MT612141.1 | 04/21/2020 | Australia:Victoria |
| MT628047.1 | 04/21/2020 | USA:Washington-Yakima-County |
| MT628051.1 | 04/21/2020 | USA:Washington-Yakima-County |
| MT628054.1 | 04/21/2020 | USA:Washington-Yakima-County |
| MT628056.1 | 04/21/2020 | USA:Washington-Snohomish-County |
| MT628057.1 | 04/21/2020 | USA:Washington-King-County |
| MT628067.1 | 04/21/2020 | USA:Washington-Pierce-County |
| MT628068.1 | 04/21/2020 | USA:Washington-Yakima-County |
| MT628074.1 | 04/21/2020 | USA:Washington-Snohomish-County |
| MT628080.1 | 04/21/2020 | USA:Washington-Yakima-County |
| MT628209.1 | 04/21/2020 | USA:CA |
| MT628272.1 | 04/21/2020 | USA:CA |
| MT642161.1 | 04/21/2020 | USA:Washington-Yakima-County |
| MT642268.1 | 04/21/2020 | USA:Washington-Snohomish-County |
| MT642343.1 | 04/21/2020 | USA:Washington-Franklin-County |
| MT642433.1 | 04/21/2020 | USA:Washington-Franklin-County |
| MT435085.1 | 04/22/2020 | India:Gandhinagar |
| MT438718.1 | 04/22/2020 | USA:CA |
| MT438719.1 | 04/22/2020 | USA:CA |
| MT438720.1 | 04/22/2020 | USA:CA |
| MT438721.1 | 04/22/2020 | USA:CA |
| MT438722.1 | 04/22/2020 | USA:CA |
| MT438723.1 | 04/22/2020 | USA:CA |
| MT438724.1 | 04/22/2020 | USA:CA |
| MT438725.1 | 04/22/2020 | USA:CA |
| MT438727.1 | 04/22/2020 | USA:CA |
| MT438728.1 | 04/22/2020 | USA:CA |
| MT438729.1 | 04/22/2020 | USA:CA |
| MT438730.1 | 04/22/2020 | USA:CA |
| MT438731.1 | 04/22/2020 | USA:CA |
| MT438732.1 | 04/22/2020 | USA:CA |
| MT438733.1 | 04/22/2020 | USA:CA |
| MT438734.1 | 04/22/2020 | USA:CA |
| MT438735.1 | 04/22/2020 | USA:CA |
| MT438736.1 | 04/22/2020 | USA:CA |

| | | |
|---|---|---|
| MT438737.1 | 04/22/2020 | USA:CA |
| MT438738.1 | 04/22/2020 | USA:CA |
| MT438739.1 | 04/22/2020 | USA:CA |
| MT438740.1 | 04/22/2020 | USA:CA |
| MT438741.1 | 04/22/2020 | USA:CA |
| MT438742.1 | 04/22/2020 | USA:CA |
| MT438743.1 | 04/22/2020 | USA:CA |
| MT438744.1 | 04/22/2020 | USA:CA |
| MT438745.1 | 04/22/2020 | USA:CA |
| MT438746.1 | 04/22/2020 | USA:CA |
| MT438747.1 | 04/22/2020 | USA:CA |
| MT438748.1 | 04/22/2020 | USA:CA |
| MT438749.1 | 04/22/2020 | USA:CA |
| MT438750.1 | 04/22/2020 | USA:CA |
| MT438751.1 | 04/22/2020 | USA:CA |
| MT438752.1 | 04/22/2020 | USA:CA |
| MT438753.1 | 04/22/2020 | USA:CA |
| MT438754.1 | 04/22/2020 | USA:CA |
| MT438755.1 | 04/22/2020 | USA:CA |
| MT438756.1 | 04/22/2020 | USA:CA |
| MT438757.1 | 04/22/2020 | USA:CA |
| MT438758.1 | 04/22/2020 | USA:CA |
| MT438759.1 | 04/22/2020 | USA:CA |
| MT438760.1 | 04/22/2020 | USA:CA |
| MT449661.1 | 04/22/2020 | USA:CA |
| MT449666.1 | 04/22/2020 | USA:CA |
| MT506697.1 | 04/22/2020 | USA:Michigan |
| MT506698.1 | 04/22/2020 | USA:Michigan |
| MT506699.1 | 04/22/2020 | USA:Michigan |
| MT506700.1 | 04/22/2020 | USA:Michigan |
| MT506701.1 | 04/22/2020 | USA:Michigan |
| MT569474.1 | 04/22/2020 | Serbia:Kraljevo |
| MT612144.1 | 04/22/2020 | Australia:Victoria |
| MT620769.1 | 04/22/2020 | USA |
| MT620772.1 | 04/22/2020 | USA |
| MT628061.1 | 04/22/2020 | USA:Washington-Yakima-County |
| MT628066.1 | 04/22/2020 | USA:Washington-Yakima-County |
| MT628069.1 | 04/22/2020 | USA:Washington-Yakima-County |
| MT628076.1 | 04/22/2020 | USA:Washington-Yakima-County |
| MT628210.1 | 04/22/2020 | USA:CA |
| MT631793.1 | 04/22/2020 | USA:CA |
| MT631794.1 | 04/22/2020 | USA:CA |
| MT631795.1 | 04/22/2020 | USA:CA |
| MT631797.1 | 04/22/2020 | USA:CA |
| MT642216.1 | 04/22/2020 | USA:Washington-Yakima-County |
| MT642235.1 | 04/22/2020 | USA:Washington-Pierce-County |

| | | |
|---|---|---|
| MT642244.1 | 04/22/2020 | USA:Washington-Yakima-County |
| MT642293.1 | 04/22/2020 | USA:Washington-Yakima-County |
| MT642305.1 | 04/22/2020 | USA:Washington-Cowlitz-County |
| MT642341.1 | 04/22/2020 | USA:Washington-Grant-County |
| MT642353.1 | 04/22/2020 | USA:Washington-Yakima-County |
| MT642364.1 | 04/22/2020 | USA:Washington-Mason-County |
| MT642423.1 | 04/22/2020 | USA:Washington-Yakima-County |
| MT434758.1 | 04/23/2020 | India |
| MT439597.1 | 04/23/2020 | India |
| MT449662.1 | 04/23/2020 | USA:CA |
| MT449673.1 | 04/23/2020 | USA:CA |
| MT506587.1 | 04/23/2020 | USA:Michigan |
| MT513758.1 | 04/23/2020 | Morocco |
| MT539729.1 | 04/23/2020 | USA |
| MT569473.1 | 04/23/2020 | Serbia:Kraljevo |
| MT585001.1 | 04/23/2020 | USA:Michigan |
| MT585008.1 | 04/23/2020 | USA:Michigan |
| MT585009.1 | 04/23/2020 | USA:Michigan |
| MT585010.1 | 04/23/2020 | USA:Michigan |
| MT585011.1 | 04/23/2020 | USA:Michigan |
| MT627754.1 | 04/23/2020 | USA:Washington-King-County |
| MT627762.1 | 04/23/2020 | USA:Washington-King-County |
| MT627920.1 | 04/23/2020 | USA:Washington |
| MT627921.1 | 04/23/2020 | USA:Washington |
| MT627922.1 | 04/23/2020 | USA:Washington |
| MT627925.1 | 04/23/2020 | USA:Washington |
| MT627927.1 | 04/23/2020 | USA:Washington |
| MT627929.1 | 04/23/2020 | USA:Washington |
| MT628156.1 | 04/23/2020 | USA:CA |
| MT628157.1 | 04/23/2020 | USA:CA |
| MT628211.1 | 04/23/2020 | USA:CA |
| MT642081.1 | 04/23/2020 | USA:Washington-Snohomish-County |
| MT642086.1 | 04/23/2020 | USA:Washington-Snohomish-County |
| MT642102.1 | 04/23/2020 | USA:Washington-Pierce-County |
| MT642104.1 | 04/23/2020 | USA:Washington-Snohomish-County |
| MT642143.1 | 04/23/2020 | USA:Washington-Yakima-County |
| MT642152.1 | 04/23/2020 | USA:Washington-Yakima-County |
| MT642172.1 | 04/23/2020 | USA:Washington-Snohomish-County |
| MT642194.1 | 04/23/2020 | USA:Washington-Snohomish-County |
| MT642201.1 | 04/23/2020 | USA:Washington-Snohomish-County |
| MT642203.1 | 04/23/2020 | USA:Washington-Yakima-County |
| MT642212.1 | 04/23/2020 | USA:Washington-Yakima-County |
| MT642220.1 | 04/23/2020 | USA:Washington-Yakima-County |
| MT642240.1 | 04/23/2020 | USA:Washington-Yakima-County |
| MT642249.1 | 04/23/2020 | USA:Washington-Yakima-County |
| MT642252.1 | 04/23/2020 | USA:Washington-Snohomish-County |

| | | |
|---|---|---|
| MT642257.1 | 04/23/2020 | USA:Washington-Yakima-County |
| MT642263.1 | 04/23/2020 | USA:Washington-Snohomish-County |
| MT642265.1 | 04/23/2020 | USA:Washington-Snohomish-County |
| MT642277.1 | 04/23/2020 | USA:Washington-Snohomish-County |
| MT642288.1 | 04/23/2020 | USA:Washington-Yakima-County |
| MT642290.1 | 04/23/2020 | USA:Washington-Yakima-County |
| MT642313.1 | 04/23/2020 | USA:Washington-Snohomish-County |
| MT642315.1 | 04/23/2020 | USA:Washington-Yakima-County |
| MT642319.1 | 04/23/2020 | USA:Washington-Yakima-County |
| MT642391.1 | 04/23/2020 | USA:Washington-Yakima-County |
| MT396266.1 | 04/24/2020 | Netherlands:Milheeze |
| MT439595.1 | 04/24/2020 | India |
| MT439596.1 | 04/24/2020 | India |
| MT451874.1 | 04/24/2020 | India:Surat |
| MT506702.1 | 04/24/2020 | USA:Michigan |
| MT584999.1 | 04/24/2020 | USA:Michigan |
| MT585000.1 | 04/24/2020 | USA:Michigan |
| MT585002.1 | 04/24/2020 | USA:Michigan |
| MT585003.1 | 04/24/2020 | USA:Michigan |
| MT585004.1 | 04/24/2020 | USA:Michigan |
| MT585005.1 | 04/24/2020 | USA:Michigan |
| MT585006.1 | 04/24/2020 | USA:Michigan |
| MT585007.1 | 04/24/2020 | USA:Michigan |
| MT585012.1 | 04/24/2020 | USA:Michigan |
| MT642074.1 | 04/24/2020 | USA:Washington-King-County |
| MT642079.1 | 04/24/2020 | USA:Washington-King-County |
| MT642089.1 | 04/24/2020 | USA:Washington-King-County |
| MT642111.1 | 04/24/2020 | USA:Washington-Yakima-County |
| MT642120.1 | 04/24/2020 | USA:Washington-Snohomish-County |
| MT642148.1 | 04/24/2020 | USA:Washington-Pierce-County |
| MT642163.1 | 04/24/2020 | USA:Washington-King-County |
| MT642230.1 | 04/24/2020 | USA:Washington-Asotin-County |
| MT642243.1 | 04/24/2020 | USA:Washington-Snohomish-County |
| MT642247.1 | 04/24/2020 | USA:Washington-Snohomish-County |
| MT642261.1 | 04/24/2020 | USA:Washington-Snohomish-County |
| MT642308.1 | 04/24/2020 | USA:Washington-Yakima-County |
| MT642346.1 | 04/24/2020 | USA:Washington-Yakima-County |
| MT449672.1 | 04/25/2020 | USA:CA |
| MT481904.1 | 04/25/2020 | India:Gandhinagar |
| MT533199.1 | 04/25/2020 | USA |
| MT577359.1 | 04/25/2020 | Bangladesh |
| MT612146.1 | 04/25/2020 | Australia:Victoria |
| MT627749.1 | 04/25/2020 | USA:Washington-King-County |
| MT628085.1 | 04/25/2020 | USA:CA |
| MT628158.1 | 04/25/2020 | USA:CA |
| MT642185.1 | 04/25/2020 | USA:Washington-Yakima-County |

| | | |
|---|---|---|
| MT642217.1 | 04/25/2020 | USA:Washington-Yakima-County |
| MT642223.1 | 04/25/2020 | USA:Washington-King-County |
| MT642298.1 | 04/25/2020 | USA:Washington-Yakima-County |
| MT642316.1 | 04/25/2020 | USA:Washington |
| MT449670.1 | 04/26/2020 | USA:CA |
| MT451875.1 | 04/26/2020 | India:Surat |
| MT451876.1 | 04/26/2020 | India:Surat |
| MT451877.1 | 04/26/2020 | India:Surat |
| MT451880.1 | 04/26/2020 | India:Surat |
| MT451881.1 | 04/26/2020 | India:Ahmedabad |
| MT451882.1 | 04/26/2020 | India:Ahmedabad |
| MT451883.1 | 04/26/2020 | India:Ahmedabad |
| MT451884.1 | 04/26/2020 | India:Ahmedabad |
| MT451885.1 | 04/26/2020 | India:Ahmedabad |
| MT451886.1 | 04/26/2020 | India:Ahmedabad |
| MT451887.1 | 04/26/2020 | India:Ahmedabad |
| MT451888.1 | 04/26/2020 | India:Ahmedabad |
| MT451889.1 | 04/26/2020 | India:Ahmedabad |
| MT451890.1 | 04/26/2020 | India:Ahmedabad |
| MT460117.1 | 04/26/2020 | USA:CA |
| MT460118.1 | 04/26/2020 | USA:CA |
| MT467237.1 | 04/26/2020 | India:Ahmedabad |
| MT467239.1 | 04/26/2020 | India:Ahmedabad |
| MT467241.1 | 04/26/2020 | India:Ahmedabad |
| MT477862.1 | 04/26/2020 | USA:AK |
| MT481907.1 | 04/26/2020 | India:Gandhinagar |
| MT533198.1 | 04/26/2020 | USA |
| MT533212.1 | 04/26/2020 | USA:CA |
| MT533213.1 | 04/26/2020 | USA:CA |
| MT533214.1 | 04/26/2020 | USA:CA |
| MT533215.1 | 04/26/2020 | USA:CA |
| MT533216.1 | 04/26/2020 | USA:CA |
| MT584996.1 | 04/26/2020 | USA:Michigan |
| MT584997.1 | 04/26/2020 | USA:Michigan |
| MT584998.1 | 04/26/2020 | USA:Michigan |
| MT627764.1 | 04/26/2020 | USA:Washington-King-County |
| MT627765.1 | 04/26/2020 | USA:Washington-King-County |
| MT627767.1 | 04/26/2020 | USA:Washington-King-County |
| MT628084.1 | 04/26/2020 | USA:CA |
| MT631799.1 | 04/26/2020 | USA:CA |
| MT642078.1 | 04/26/2020 | USA:Washington-Snohomish-County |
| MT642142.1 | 04/26/2020 | USA:Washington-Yakima-County |
| MT642242.1 | 04/26/2020 | USA:Washington-Yakima-County |
| MT642280.1 | 04/26/2020 | USA:Washington-Snohomish-County |
| MT642356.1 | 04/26/2020 | USA:Washington |
| MT642382.1 | 04/26/2020 | USA:Washington-Yakima-County |

| | | |
|---|---|---|
| MT451878.1 | 04/27/2020 | India:Surat |
| MT451879.1 | 04/27/2020 | India:Surat |
| MT460113.1 | 04/27/2020 | USA:CA |
| MT460114.1 | 04/27/2020 | USA:CA |
| MT460115.1 | 04/27/2020 | USA:CA |
| MT460121.1 | 04/27/2020 | USA:CA |
| MT460122.1 | 04/27/2020 | USA:CA |
| MT460123.1 | 04/27/2020 | USA:CA |
| MT481906.1 | 04/27/2020 | India:Gandhinagar |
| MT496991.1 | 04/27/2020 | India:Gandhinagar |
| MT506585.1 | 04/27/2020 | USA:Michigan |
| MT506586.1 | 04/27/2020 | USA:Michigan |
| MT509506.1 | 04/27/2020 | India:Rajkot |
| MT517426.1 | 04/27/2020 | Czech-Republic |
| MT532435.1 | 04/27/2020 | USA |
| MT533197.1 | 04/27/2020 | USA |
| MT533200.1 | 04/27/2020 | USA |
| MT533201.1 | 04/27/2020 | USA |
| MT533202.1 | 04/27/2020 | USA |
| MT533203.1 | 04/27/2020 | USA |
| MT533217.1 | 04/27/2020 | USA:CA |
| MT533218.1 | 04/27/2020 | USA:CA |
| MT533219.1 | 04/27/2020 | USA:CA |
| MT534338.1 | 04/27/2020 | USA:CA |
| MT534339.1 | 04/27/2020 | USA:CA |
| MT539174.1 | 04/27/2020 | India:Rajkot |
| MT576034.1 | 04/27/2020 | India:Rajkot |
| MT576038.1 | 04/27/2020 | India:Rajkot |
| MT577612.1 | 04/27/2020 | USA:FL |
| MT577613.1 | 04/27/2020 | USA:FL |
| MT585013.1 | 04/27/2020 | USA:Michigan |
| MT585014.1 | 04/27/2020 | USA:Michigan |
| MT585015.1 | 04/27/2020 | USA:Michigan |
| MT612147.1 | 04/27/2020 | Australia:Victoria |
| MT612154.1 | 04/27/2020 | Australia:Victoria |
| MT627746.1 | 04/27/2020 | USA:Washington-King-County |
| MT628159.1 | 04/27/2020 | USA:CA |
| MT642116.1 | 04/27/2020 | USA:Washington-King-County |
| MT642138.1 | 04/27/2020 | USA:Washington-Snohomish-County |
| MT642144.1 | 04/27/2020 | USA:Washington-Snohomish-County |
| MT642158.1 | 04/27/2020 | USA:Washington-Yakima-County |
| MT642165.1 | 04/27/2020 | USA:Washington-Yakima-County |
| MT642169.1 | 04/27/2020 | USA:Washington-Yakima-County |
| MT642184.1 | 04/27/2020 | USA:Washington-Snohomish-County |
| MT642206.1 | 04/27/2020 | USA:Washington-Yakima-County |
| MT642255.1 | 04/27/2020 | USA:Washington-Yakima-County |

| | | |
|---|---|---|
| MT642258.1 | 04/27/2020 | USA:Washington-Snohomish-County |
| MT642266.1 | 04/27/2020 | USA:Washington-Yakima-County |
| MT642283.1 | 04/27/2020 | USA:Washington-Adams-County |
| MT642299.1 | 04/27/2020 | USA:Washington-Cowlitz-County |
| MT642311.1 | 04/27/2020 | USA:Washington-Yakima-County |
| MT642334.1 | 04/27/2020 | USA:Washington-Yakima-County |
| MT642352.1 | 04/27/2020 | USA:Washington-Snohomish-County |
| MT642389.1 | 04/27/2020 | USA:Washington-Yakima-County |
| MT642401.1 | 04/27/2020 | USA:Washington-Snohomish-County |
| MT642410.1 | 04/27/2020 | USA:Washington-Yakima-County |
| MT642411.1 | 04/27/2020 | USA:Washington-Yakima-County |
| MT642412.1 | 04/27/2020 | USA:Washington-Cowlitz-County |
| MT642420.1 | 04/27/2020 | USA:Washington-Yakima-County |
| MT642422.1 | 04/27/2020 | USA:Washington-Snohomish-County |
| MT642427.1 | 04/27/2020 | USA:Washington-Snohomish-County |
| MT460119.1 | 04/28/2020 | USA:CA |
| MT460120.1 | 04/28/2020 | USA:CA |
| MT481900.1 | 04/28/2020 | India:Gandhinagar |
| MT481901.1 | 04/28/2020 | India:Dahegam |
| MT481908.1 | 04/28/2020 | India:Mansa |
| MT509511.1 | 04/28/2020 | India:Rajkot |
| MT512644.1 | 04/28/2020 | USA |
| MT539170.1 | 04/28/2020 | India:Rajkot |
| MT566434.1 | 04/28/2020 | Bangladesh |
| MT566435.1 | 04/28/2020 | Bangladesh |
| MT566436.1 | 04/28/2020 | Bangladesh |
| MT566437.1 | 04/28/2020 | Bangladesh |
| MT566438.1 | 04/28/2020 | Bangladesh |
| MT569472.1 | 04/28/2020 | Serbia:Uzice |
| MT577615.1 | 04/28/2020 | USA:FL |
| MT577616.1 | 04/28/2020 | USA:FL |
| MT577617.1 | 04/28/2020 | USA:FL |
| MT577618.1 | 04/28/2020 | USA:FL |
| MT577619.1 | 04/28/2020 | USA:FL |
| MT577620.1 | 04/28/2020 | USA:FL |
| MT577621.1 | 04/28/2020 | USA:FL |
| MT577622.1 | 04/28/2020 | USA:FL |
| MT577623.1 | 04/28/2020 | USA:FL |
| MT577624.1 | 04/28/2020 | USA:FL |
| MT577625.1 | 04/28/2020 | USA:FL |
| MT577627.1 | 04/28/2020 | USA:FL |
| MT577628.1 | 04/28/2020 | USA:FL |
| MT577629.1 | 04/28/2020 | USA:FL |
| MT577630.1 | 04/28/2020 | USA:FL |
| MT598175.1 | 04/28/2020 | USA:San-Diego-California |
| MT612162.1 | 04/28/2020 | Australia:Victoria |

| | | |
|---|---|---|
| MT628160.1 | 04/28/2020 | USA:CA |
| MT628207.1 | 04/28/2020 | USA:CA |
| MT628212.1 | 04/28/2020 | USA:CA |
| MT628213.1 | 04/28/2020 | USA:CA |
| MT642105.1 | 04/28/2020 | USA:Washington-Snohomish-County |
| MT642109.1 | 04/28/2020 | USA:Washington-Snohomish-County |
| MT642179.1 | 04/28/2020 | USA:Washington-Yakima-County |
| MT642187.1 | 04/28/2020 | USA:Washington-Yakima-County |
| MT642195.1 | 04/28/2020 | USA:Washington-Yakima-County |
| MT642270.1 | 04/28/2020 | USA:Washington-Yakima-County |
| MT642272.1 | 04/28/2020 | USA:Washington-Yakima-County |
| MT642284.1 | 04/28/2020 | USA:Washington-Yakima-County |
| MT642285.1 | 04/28/2020 | USA:Washington-Spokane-County |
| MT642286.1 | 04/28/2020 | USA:Washington-Yakima-County |
| MT642300.1 | 04/28/2020 | USA:Washington-Skagit-County |
| MT642307.1 | 04/28/2020 | USA:Washington-Benton-County |
| MT642312.1 | 04/28/2020 | USA:Washington-Spokane-County |
| MT642330.1 | 04/28/2020 | USA:Washington-Yakima-County |
| MT642360.1 | 04/28/2020 | USA:Washington-Skagit-County |
| MT642366.1 | 04/28/2020 | USA:Washington-Benton-County |
| MT642370.1 | 04/28/2020 | USA:Washington-Yakima-County |
| MT642375.1 | 04/28/2020 | USA:Washington-Yakima-County |
| MT642398.1 | 04/28/2020 | USA:Washington-Yakima-County |
| MT642415.1 | 04/28/2020 | USA:Washington-Skagit-County |
| MT457391.1 | 04/29/2020 | Netherlands |
| MT457392.1 | 04/29/2020 | Netherlands |
| MT457393.1 | 04/29/2020 | Netherlands |
| MT460116.1 | 04/29/2020 | USA:CA |
| MT460140.1 | 04/29/2020 | USA:CA |
| MT467238.1 | 04/29/2020 | India:Ahmedabad |
| MT467242.1 | 04/29/2020 | India:Ahmedabad |
| MT467245.1 | 04/29/2020 | India:Ahmedabad |
| MT467246.1 | 04/29/2020 | India:Ahmedabad |
| MT467247.1 | 04/29/2020 | India:Ahmedabad |
| MT467248.1 | 04/29/2020 | India:Ahmedabad |
| MT467249.1 | 04/29/2020 | India:Ahmedabad |
| MT467250.1 | 04/29/2020 | India:Ahmedabad |
| MT467251.1 | 04/29/2020 | India:Ahmedabad |
| MT467252.1 | 04/29/2020 | India:Ahmedabad |
| MT467253.1 | 04/29/2020 | India:Ahmedabad |
| MT467254.1 | 04/29/2020 | India:Ahmedabad |
| MT467256.1 | 04/29/2020 | India:Ahmedabad |
| MT467258.1 | 04/29/2020 | India:Ahmedabad |
| MT477904.1 | 04/29/2020 | USA:FL |
| MT496972.1 | 04/29/2020 | India:Ahmedabad |
| MT496973.1 | 04/29/2020 | India:Ahmedabad |

| | | |
|---|---|---|
| MT496974.1 | 04/29/2020 | India:Ahmedabad |
| MT496975.1 | 04/29/2020 | India:Ahmedabad |
| MT496990.1 | 04/29/2020 | India:Gandhinagar |
| MT496993.1 | 04/29/2020 | India:Gandhinagar |
| MT496994.1 | 04/29/2020 | India:Gandhinagar |
| MT496995.1 | 04/29/2020 | India:Gandhinagar |
| MT496997.1 | 04/29/2020 | India:Gandhinagar |
| MT533211.1 | 04/29/2020 | USA:CA |
| MT533220.1 | 04/29/2020 | USA:CA |
| MT533228.1 | 04/29/2020 | USA:CA |
| MT533229.1 | 04/29/2020 | USA:CA |
| MT533230.1 | 04/29/2020 | USA:CA |
| MT533231.1 | 04/29/2020 | USA:CA |
| MT577631.1 | 04/29/2020 | USA:FL |
| MT577632.1 | 04/29/2020 | USA:FL |
| MT577634.1 | 04/29/2020 | USA:FL |
| MT585016.1 | 04/29/2020 | USA:Michigan |
| MT585017.1 | 04/29/2020 | USA:Michigan |
| MT585018.1 | 04/29/2020 | USA:Michigan |
| MT585019.1 | 04/29/2020 | USA:Michigan |
| MT585020.1 | 04/29/2020 | USA:Michigan |
| MT585021.1 | 04/29/2020 | USA:Michigan |
| MT585022.1 | 04/29/2020 | USA:Michigan |
| MT585023.1 | 04/29/2020 | USA:Michigan |
| MT585024.1 | 04/29/2020 | USA:Michigan |
| MT585025.1 | 04/29/2020 | USA:Michigan |
| MT585026.1 | 04/29/2020 | USA:Michigan |
| MT585027.1 | 04/29/2020 | USA:Michigan |
| MT585028.1 | 04/29/2020 | USA:Michigan |
| MT585029.1 | 04/29/2020 | USA:Michigan |
| MT585030.1 | 04/29/2020 | USA:Michigan |
| MT585031.1 | 04/29/2020 | USA:Michigan |
| MT585032.1 | 04/29/2020 | USA:Michigan |
| MT585033.1 | 04/29/2020 | USA:Michigan |
| MT585034.1 | 04/29/2020 | USA:Michigan |
| MT585035.1 | 04/29/2020 | USA:Michigan |
| MT585036.1 | 04/29/2020 | USA:Michigan |
| MT585037.1 | 04/29/2020 | USA:Michigan |
| MT585038.1 | 04/29/2020 | USA:Michigan |
| MT585039.1 | 04/29/2020 | USA:Michigan |
| MT585040.1 | 04/29/2020 | USA:Michigan |
| MT585041.1 | 04/29/2020 | USA:Michigan |
| MT585042.1 | 04/29/2020 | USA:Michigan |
| MT585043.1 | 04/29/2020 | USA:Michigan |
| MT585044.1 | 04/29/2020 | USA:Michigan |
| MT585045.1 | 04/29/2020 | USA:Michigan |

| | | |
|---|---|---|
| MT585046.1 | 04/29/2020 | USA:Michigan |
| MT585047.1 | 04/29/2020 | USA:Michigan |
| MT585048.1 | 04/29/2020 | USA:Michigan |
| MT585049.1 | 04/29/2020 | USA:Michigan |
| MT598176.1 | 04/29/2020 | USA:San-Diego-California |
| MT612150.1 | 04/29/2020 | Australia:Victoria |
| MT612151.1 | 04/29/2020 | Australia:Victoria |
| MT612153.1 | 04/29/2020 | Australia:Victoria |
| MT612155.1 | 04/29/2020 | Australia:Victoria |
| MT612163.1 | 04/29/2020 | Australia:Victoria |
| MT612311.1 | 04/29/2020 | Australia:Victoria |
| MT627747.1 | 04/29/2020 | USA:Washington-King-County |
| MT628206.1 | 04/29/2020 | USA:CA |
| MT628218.1 | 04/29/2020 | USA:CA |
| MT642085.1 | 04/29/2020 | USA:Washington-Pierce-County |
| MT642091.1 | 04/29/2020 | USA:Washington-Yakima-County |
| MT642092.1 | 04/29/2020 | USA:Washington-Pierce-County |
| MT642093.1 | 04/29/2020 | USA:Washington-Yakima-County |
| MT642122.1 | 04/29/2020 | USA:Washington-Yakima-County |
| MT642129.1 | 04/29/2020 | USA:Washington-Yakima-County |
| MT642136.1 | 04/29/2020 | USA:Washington-Pierce-County |
| MT642137.1 | 04/29/2020 | USA:Washington-Snohomish-County |
| MT642147.1 | 04/29/2020 | USA:Washington-Pierce-County |
| MT642150.1 | 04/29/2020 | USA:Washington-Yakima-County |
| MT642151.1 | 04/29/2020 | USA:Washington-Pierce-County |
| MT642181.1 | 04/29/2020 | USA:Washington-Yakima-County |
| MT642188.1 | 04/29/2020 | USA:Washington-Yakima-County |
| MT642193.1 | 04/29/2020 | USA:Washington-Pierce-County |
| MT642197.1 | 04/29/2020 | USA:Washington-Pierce-County |
| MT642199.1 | 04/29/2020 | USA:Washington-Yakima-County |
| MT642210.1 | 04/29/2020 | USA:Washington-Pierce-County |
| MT642215.1 | 04/29/2020 | USA:Washington-Yakima-County |
| MT642239.1 | 04/29/2020 | USA:Washington-Yakima-County |
| MT642256.1 | 04/29/2020 | USA:Washington-Yakima-County |
| MT642262.1 | 04/29/2020 | USA:Washington-Yakima-County |
| MT642275.1 | 04/29/2020 | USA:Washington-King-County |
| MT642303.1 | 04/29/2020 | USA:Washington-Yakima-County |
| MT642317.1 | 04/29/2020 | USA:Washington-Yakima-County |
| MT642318.1 | 04/29/2020 | USA:Washington-Snohomish-County |
| MT642322.1 | 04/29/2020 | USA:Washington |
| MT642323.1 | 04/29/2020 | USA:Washington-Yakima-County |
| MT642342.1 | 04/29/2020 | USA:Washington-Yakima-County |
| MT642344.1 | 04/29/2020 | USA:Washington-Yakima-County |
| MT642347.1 | 04/29/2020 | USA:Washington-Yakima-County |
| MT642385.1 | 04/29/2020 | USA:Washington-Skagit-County |
| MT642425.1 | 04/29/2020 | USA:Washington-Yakima-County |

| | | |
|---|---|---|
| MT642431.1 | 04/29/2020 | USA:Washington-Yakima-County |
| MT460125.1 | 04/30/2020 | USA:CA |
| MT460126.1 | 04/30/2020 | USA:CA |
| MT460127.1 | 04/30/2020 | USA:CA |
| MT460128.1 | 04/30/2020 | USA:CA |
| MT460129.1 | 04/30/2020 | USA:CA |
| MT460130.1 | 04/30/2020 | USA:CA |
| MT460131.1 | 04/30/2020 | USA:CA |
| MT460132.1 | 04/30/2020 | USA:CA |
| MT460133.1 | 04/30/2020 | USA:CA |
| MT460134.1 | 04/30/2020 | USA:CA |
| MT460135.1 | 04/30/2020 | USA:CA |
| MT460136.1 | 04/30/2020 | USA:CA |
| MT460137.1 | 04/30/2020 | USA:CA |
| MT460138.1 | 04/30/2020 | USA:CA |
| MT460139.1 | 04/30/2020 | USA:CA |
| MT467240.1 | 04/30/2020 | India:Ahmedabad |
| MT467243.1 | 04/30/2020 | India:Ahmedabad |
| MT467244.1 | 04/30/2020 | India:Ahmedabad |
| MT467255.1 | 04/30/2020 | India:Ahmedabad |
| MT467257.1 | 04/30/2020 | India:Ahmedabad |
| MT506229.1 | 04/30/2020 | USA:Michigan |
| MT533221.1 | 04/30/2020 | USA:CA |
| MT533222.1 | 04/30/2020 | USA:CA |
| MT533223.1 | 04/30/2020 | USA:CA |
| MT533224.1 | 04/30/2020 | USA:CA |
| MT533225.1 | 04/30/2020 | USA:CA |
| MT533226.1 | 04/30/2020 | USA:CA |
| MT533227.1 | 04/30/2020 | USA:CA |
| MT577635.1 | 04/30/2020 | USA:FL |
| MT577636.1 | 04/30/2020 | USA:FL |
| MT577637.1 | 04/30/2020 | USA:FL |
| MT577638.1 | 04/30/2020 | USA:FL |
| MT577639.1 | 04/30/2020 | USA:FL |
| MT577640.1 | 04/30/2020 | USA:FL |
| MT577641.1 | 04/30/2020 | USA:FL |
| MT577642.1 | 04/30/2020 | USA:FL |
| MT577643.1 | 04/30/2020 | USA:FL |
| MT577644.1 | 04/30/2020 | USA:FL |
| MT585050.1 | 04/30/2020 | USA:Michigan |
| MT585051.1 | 04/30/2020 | USA:Michigan |
| MT585052.1 | 04/30/2020 | USA:Michigan |
| MT585053.1 | 04/30/2020 | USA:Michigan |
| MT585054.1 | 04/30/2020 | USA:Michigan |
| MT585055.1 | 04/30/2020 | USA:Michigan |
| MT585056.1 | 04/30/2020 | USA:Michigan |

| | | |
|---|---|---|
| MT585057.1 | 04/30/2020 | USA:Michigan |
| MT585058.1 | 04/30/2020 | USA:Michigan |
| MT585059.1 | 04/30/2020 | USA:Michigan |
| MT585060.1 | 04/30/2020 | USA:Michigan |
| MT585061.1 | 04/30/2020 | USA:Michigan |
| MT585062.1 | 04/30/2020 | USA:Michigan |
| MT585063.1 | 04/30/2020 | USA:Michigan |
| MT585064.1 | 04/30/2020 | USA:Michigan |
| MT585065.1 | 04/30/2020 | USA:Michigan |
| MT585066.1 | 04/30/2020 | USA:Michigan |
| MT585067.1 | 04/30/2020 | USA:Michigan |
| MT585068.1 | 04/30/2020 | USA:Michigan |
| MT585069.1 | 04/30/2020 | USA:Michigan |
| MT585070.1 | 04/30/2020 | USA:Michigan |
| MT585071.1 | 04/30/2020 | USA:Michigan |
| MT585072.1 | 04/30/2020 | USA:Michigan |
| MT585073.1 | 04/30/2020 | USA:Michigan |
| MT585074.1 | 04/30/2020 | USA:Michigan |
| MT585075.1 | 04/30/2020 | USA:Michigan |
| MT585076.1 | 04/30/2020 | USA:Michigan |
| MT585077.1 | 04/30/2020 | USA:Michigan |
| MT585078.1 | 04/30/2020 | USA:Michigan |
| MT612159.1 | 04/30/2020 | Australia:Victoria |
| MT628204.1 | 04/30/2020 | USA:CA |
| MT628205.1 | 04/30/2020 | USA:CA |
| MT642087.1 | 04/30/2020 | USA:Washington-Snohomish-County |
| MT642107.1 | 04/30/2020 | USA:Washington-Yakima-County |
| MT642115.1 | 04/30/2020 | USA:Washington-Snohomish-County |
| MT642121.1 | 04/30/2020 | USA:Washington-Yakima-County |
| MT642124.1 | 04/30/2020 | USA:Washington-Snohomish-County |
| MT642125.1 | 04/30/2020 | USA:Washington-Yakima-County |
| MT642133.1 | 04/30/2020 | USA:Washington-Snohomish-County |
| MT642154.1 | 04/30/2020 | USA:Washington-Yakima-County |
| MT642160.1 | 04/30/2020 | USA:Washington-Yakima-County |
| MT642190.1 | 04/30/2020 | USA:Washington-Yakima-County |
| MT642202.1 | 04/30/2020 | USA:Washington-Yakima-County |
| MT642204.1 | 04/30/2020 | USA:Washington-Yakima-County |
| MT642207.1 | 04/30/2020 | USA:Washington-Yakima-County |
| MT642218.1 | 04/30/2020 | USA:Washington-Yakima-County |
| MT642226.1 | 04/30/2020 | USA:Washington-Skagit-County |
| MT642233.1 | 04/30/2020 | USA:Washington-Yakima-County |
| MT642253.1 | 04/30/2020 | USA:Washington-Yakima-County |
| MT642269.1 | 04/30/2020 | USA:Washington-Yakima-County |
| MT642271.1 | 04/30/2020 | USA:Washington-Yakima-County |
| MT642278.1 | 04/30/2020 | USA:Washington-Yakima-County |
| MT642295.1 | 04/30/2020 | USA:Washington-Yakima-County |

| | | |
|---|---|---|
| MT642320.1 | 04/30/2020 | USA:Washington-Yakima-County |
| MT642321.1 | 04/30/2020 | USA:Washington-Pierce-County |
| MT642331.1 | 04/30/2020 | USA:Washington-Yakima-County |
| MT642340.1 | 04/30/2020 | USA:Washington-Yakima-County |
| MT642349.1 | 04/30/2020 | USA:Washington-Yakima-County |
| MT642357.1 | 04/30/2020 | USA:Washington-Yakima-County |
| MT642369.1 | 04/30/2020 | USA:Washington-Yakima-County |
| MT642372.1 | 04/30/2020 | USA:Washington-Skagit-County |
| MT460089.1 | 05/01/2020 | USA:CA |
| MT460090.1 | 05/01/2020 | USA:CA |
| MT460091.1 | 05/01/2020 | USA:CA |
| MT460092.1 | 05/01/2020 | USA:CA |
| MT509512.1 | 05/01/2020 | India:Dahod |
| MT585079.1 | 05/01/2020 | USA:Michigan |
| MT585080.1 | 05/01/2020 | USA:Michigan |
| MT628163.1 | 05/01/2020 | USA:CA |
| MT628164.1 | 05/01/2020 | USA:CA |
| MT628165.1 | 05/01/2020 | USA:CA |
| MT628166.1 | 05/01/2020 | USA:CA |
| MT628169.1 | 05/01/2020 | USA:CA |
| MT628203.1 | 05/01/2020 | USA:CA |
| MT642082.1 | 05/01/2020 | USA:Washington-Yakima-County |
| MT642083.1 | 05/01/2020 | USA:Washington-Yakima-County |
| MT642113.1 | 05/01/2020 | USA:Washington-Yakima-County |
| MT642126.1 | 05/01/2020 | USA:Washington-Yakima-County |
| MT642130.1 | 05/01/2020 | USA:Washington-Yakima-County |
| MT642139.1 | 05/01/2020 | USA:Washington-Yakima-County |
| MT642149.1 | 05/01/2020 | USA:Washington-Yakima-County |
| MT642162.1 | 05/01/2020 | USA:Washington-Yakima-County |
| MT642171.1 | 05/01/2020 | USA:Washington-Yakima-County |
| MT642175.1 | 05/01/2020 | USA:Washington-Yakima-County |
| MT642177.1 | 05/01/2020 | USA:Washington-Yakima-County |
| MT642178.1 | 05/01/2020 | USA:Washington-Yakima-County |
| MT642182.1 | 05/01/2020 | USA:Washington-Yakima-County |
| MT642191.1 | 05/01/2020 | USA:Washington-Yakima-County |
| MT642200.1 | 05/01/2020 | USA:Washington-Yakima-County |
| MT642222.1 | 05/01/2020 | USA:Washington-Yakima-County |
| MT642246.1 | 05/01/2020 | USA:Washington-Yakima-County |
| MT642248.1 | 05/01/2020 | USA:Washington-Yakima-County |
| MT642259.1 | 05/01/2020 | USA:Washington-Yakima-County |
| MT642276.1 | 05/01/2020 | USA:Washington-Yakima-County |
| MT642301.1 | 05/01/2020 | USA:Washington-Skagit-County |
| MT642309.1 | 05/01/2020 | USA:Washington-Yakima-County |
| MT642329.1 | 05/01/2020 | USA:Washington-Yakima-County |
| MT642336.1 | 05/01/2020 | USA:Washington-Yakima-County |
| MT642337.1 | 05/01/2020 | USA:Washington-Yakima-County |

| | | |
|---|---|---|
| MT642338.1 | 05/01/2020 | USA:Washington-Yakima-County |
| MT642339.1 | 05/01/2020 | USA:Washington-Yakima-County |
| MT642351.1 | 05/01/2020 | USA:Washington-Yakima-County |
| MT642361.1 | 05/01/2020 | USA:Washington-Yakima-County |
| MT642374.1 | 05/01/2020 | USA:Washington-Yakima-County |
| MT642384.1 | 05/01/2020 | USA:Washington-Snohomish-County |
| MT642394.1 | 05/01/2020 | USA:Washington-Yakima-County |
| MT642413.1 | 05/01/2020 | USA:Washington-Skagit-County |
| MT642421.1 | 05/01/2020 | USA:Washington-Yakima-County |
| MT642426.1 | 05/01/2020 | USA:Washington-Yakima-County |
| MT467259.1 | 05/02/2020 | India:Prantij |
| MT467260.1 | 05/02/2020 | India:Prantij |
| MT481905.1 | 05/02/2020 | India:Gandhinagar |
| MT496996.1 | 05/02/2020 | India:Gandhinagar |
| MT509649.1 | 05/02/2020 | India:Vadodara |
| MT509650.1 | 05/02/2020 | India:Vadodara |
| MT510690.1 | 05/02/2020 | Egypt |
| MT510691.1 | 05/02/2020 | Egypt |
| MT510692.1 | 05/02/2020 | Egypt |
| MT510693.1 | 05/02/2020 | Egypt |
| MT510694.1 | 05/02/2020 | Egypt |
| MT510695.1 | 05/02/2020 | Egypt |
| MT510696.1 | 05/02/2020 | Egypt |
| MT510697.1 | 05/02/2020 | Egypt |
| MT510698.1 | 05/02/2020 | Egypt |
| MT510699.1 | 05/02/2020 | Egypt |
| MT510700.1 | 05/02/2020 | Egypt |
| MT510701.1 | 05/02/2020 | Egypt |
| MT510702.1 | 05/02/2020 | Egypt |
| MT510703.1 | 05/02/2020 | Egypt |
| MT511065.1 | 05/02/2020 | Egypt |
| MT511083.1 | 05/02/2020 | Egypt |
| MT539164.1 | 05/02/2020 | India:Vadodara |
| MT539165.1 | 05/02/2020 | India:Vadodara |
| MT539166.1 | 05/02/2020 | India:Vadodara |
| MT539167.1 | 05/02/2020 | India:Vadodara |
| MT539168.1 | 05/02/2020 | India:Vadodara |
| MT539169.1 | 05/02/2020 | India:Vadodara |
| MT539171.1 | 05/02/2020 | India:Vadodara |
| MT539172.1 | 05/02/2020 | India:Vadodara |
| MT539173.1 | 05/02/2020 | India:Vadodara |
| MT539175.1 | 05/02/2020 | India:Vadodara |
| MT539176.1 | 05/02/2020 | India:Vadodara |
| MT576035.1 | 05/02/2020 | India:Vadodara |
| MT576037.1 | 05/02/2020 | India:Vadodara |
| MT576039.1 | 05/02/2020 | India:Vadodara |

| | | |
|---|---|---|
| MT612168.1 | 05/02/2020 | Australia:Victoria |
| MT612170.1 | 05/02/2020 | Australia:Victoria |
| MT612229.1 | 05/02/2020 | Australia:Victoria |
| MT612230.1 | 05/02/2020 | Australia:Victoria |
| MT612231.1 | 05/02/2020 | Australia:Victoria |
| MT612245.1 | 05/02/2020 | Australia:Victoria |
| MT628202.1 | 05/02/2020 | USA:CA |
| MT642231.1 | 05/02/2020 | USA:Washington-Yakima-County |
| MT642294.1 | 05/02/2020 | USA:Washington-Yakima-County |
| MT481895.1 | 05/03/2020 | India:Ahmedabad |
| MT481896.1 | 05/03/2020 | India:Ahmedabad |
| MT481902.1 | 05/03/2020 | India:Dahegam |
| MT481903.1 | 05/03/2020 | India:Dahegam |
| MT496976.1 | 05/03/2020 | India:Ahmedabad |
| MT496977.1 | 05/03/2020 | India:Ahmedabad |
| MT496978.1 | 05/03/2020 | India:Ahmedabad |
| MT496980.1 | 05/03/2020 | India:Ahmedabad |
| MT496981.1 | 05/03/2020 | India:Ahmedabad |
| MT496982.1 | 05/03/2020 | India:Ahmedabad |
| MT496988.1 | 05/03/2020 | India:Ahmedabad |
| MT496989.1 | 05/03/2020 | India:Ahmedabad |
| MT496992.1 | 05/03/2020 | India:Ahmedabad |
| MT509494.1 | 05/03/2020 | India:Vadodara |
| MT509500.1 | 05/03/2020 | India:Dahod |
| MT509651.1 | 05/03/2020 | India:Vadodara |
| MT509656.1 | 05/03/2020 | India:Vadodara |
| MT509657.1 | 05/03/2020 | India:Dahod |
| MT509658.1 | 05/03/2020 | India:Dahod |
| MT509659.1 | 05/03/2020 | India:Vadodara |
| MT612173.1 | 05/03/2020 | Australia:Victoria |
| MT612174.1 | 05/03/2020 | Australia:Victoria |
| MT612228.1 | 05/03/2020 | Australia:Victoria |
| MT628201.1 | 05/03/2020 | USA:CA |
| MT642080.1 | 05/03/2020 | USA:Washington-Yakima-County |
| MT642097.1 | 05/03/2020 | USA:Washington-Yakima-County |
| MT642098.1 | 05/03/2020 | USA:Washington-Yakima-County |
| MT642123.1 | 05/03/2020 | USA:Washington-Yakima-County |
| MT642127.1 | 05/03/2020 | USA:Washington-Yakima-County |
| MT642174.1 | 05/03/2020 | USA:Washington-Yakima-County |
| MT642176.1 | 05/03/2020 | USA:Washington-Yakima-County |
| MT642186.1 | 05/03/2020 | USA:Washington-Yakima-County |
| MT642238.1 | 05/03/2020 | USA:Washington-Pierce-County |
| MT642260.1 | 05/03/2020 | USA:Washington-Snohomish-County |
| MT642264.1 | 05/03/2020 | USA:Washington-Yakima-County |
| MT642273.1 | 05/03/2020 | USA:Washington-Yakima-County |
| MT642314.1 | 05/03/2020 | USA:Washington-Yakima-County |

| | | |
|---|---|---|
| MT642326.1 | 05/03/2020 | USA:Washington-Pierce-County |
| MT642365.1 | 05/03/2020 | USA:Washington-Pierce-County |
| MT642403.1 | 05/03/2020 | USA:Washington-Pierce-County |
| MT642432.1 | 05/03/2020 | USA:Washington-Yakima-County |
| MT467261.1 | 05/04/2020 | India:Modasa |
| MT467262.1 | 05/04/2020 | India:Modasa |
| MT467263.1 | 05/04/2020 | India:Dhansura |
| MT496987.1 | 05/04/2020 | India:Ahmedabad |
| MT598177.1 | 05/04/2020 | USA:San-Diego-California |
| MT612171.1 | 05/04/2020 | Australia:Victoria |
| MT612183.1 | 05/04/2020 | Australia:Victoria |
| MT612227.1 | 05/04/2020 | Australia:Victoria |
| MT642095.1 | 05/04/2020 | USA:Washington-Yakima-County |
| MT642096.1 | 05/04/2020 | USA:Washington-Yakima-County |
| MT642100.1 | 05/04/2020 | USA:Washington-Yakima-County |
| MT642106.1 | 05/04/2020 | USA:Washington-Yakima-County |
| MT642119.1 | 05/04/2020 | USA:Washington-Yakima-County |
| MT642128.1 | 05/04/2020 | USA:Washington-Yakima-County |
| MT642132.1 | 05/04/2020 | USA:Washington-Pierce-County |
| MT642141.1 | 05/04/2020 | USA:Washington-Yakima-County |
| MT642145.1 | 05/04/2020 | USA:Washington-Yakima-County |
| MT642146.1 | 05/04/2020 | USA:Washington-Yakima-County |
| MT642153.1 | 05/04/2020 | USA:Washington-Yakima-County |
| MT642155.1 | 05/04/2020 | USA:Washington-Pierce-County |
| MT642156.1 | 05/04/2020 | USA:Washington-Yakima-County |
| MT642167.1 | 05/04/2020 | USA:Washington-Skagit-County |
| MT642170.1 | 05/04/2020 | USA:Washington-Yakima-County |
| MT642205.1 | 05/04/2020 | USA:Washington-Skagit-County |
| MT642219.1 | 05/04/2020 | USA:Washington-Pierce-County |
| MT642250.1 | 05/04/2020 | USA:Washington-Pierce-County |
| MT642267.1 | 05/04/2020 | USA:Washington-Pierce-County |
| MT642279.1 | 05/04/2020 | USA:Washington-Yakima-County |
| MT642281.1 | 05/04/2020 | USA:Washington-Pierce-County |
| MT642291.1 | 05/04/2020 | USA:Washington-Pierce-County |
| MT642292.1 | 05/04/2020 | USA:Washington-Snohomish-County |
| MT642296.1 | 05/04/2020 | USA:Washington-Pierce-County |
| MT642297.1 | 05/04/2020 | USA:Washington-Pierce-County |
| MT642304.1 | 05/04/2020 | USA:Washington-Pierce-County |
| MT642310.1 | 05/04/2020 | USA:Washington-Pierce-County |
| MT642324.1 | 05/04/2020 | USA:Washington-Pierce-County |
| MT642328.1 | 05/04/2020 | USA:Washington-Yakima-County |
| MT642362.1 | 05/04/2020 | USA:Washington-Pierce-County |
| MT642363.1 | 05/04/2020 | USA:Washington-Pierce-County |
| MT642367.1 | 05/04/2020 | USA:Washington-Pierce-County |
| MT642371.1 | 05/04/2020 | USA:Washington-Pierce-County |
| MT642373.1 | 05/04/2020 | USA:Washington-Pierce-County |

| | | |
|---|---|---|
| MT642377.1 | 05/04/2020 | USA:Washington-Pierce-County |
| MT642379.1 | 05/04/2020 | USA:Washington-Pierce-County |
| MT642381.1 | 05/04/2020 | USA:Washington-Pierce-County |
| MT642388.1 | 05/04/2020 | USA:Washington-Mason-County |
| MT642392.1 | 05/04/2020 | USA:Washington-Pierce-County |
| MT642393.1 | 05/04/2020 | USA:Washington-Pierce-County |
| MT642396.1 | 05/04/2020 | USA:Washington-Pierce-County |
| MT642397.1 | 05/04/2020 | USA:Washington-Pierce-County |
| MT642399.1 | 05/04/2020 | USA:Washington-Pierce-County |
| MT642400.1 | 05/04/2020 | USA:Washington-Pierce-County |
| MT642402.1 | 05/04/2020 | USA:Washington-Pierce-County |
| MT642404.1 | 05/04/2020 | USA:Washington-Pierce-County |
| MT642406.1 | 05/04/2020 | USA:Washington-Pierce-County |
| MT642408.1 | 05/04/2020 | USA:Washington-Yakima-County |
| MT642424.1 | 05/04/2020 | USA:Washington-Yakima-County |
| MT642430.1 | 05/04/2020 | USA:Washington-Skagit-County |
| MT642434.1 | 05/04/2020 | USA:Washington-Pierce-County |
| MT481897.1 | 05/05/2020 | India:Modasa |
| MT481898.1 | 05/05/2020 | India:Himatnagar |
| MT481899.1 | 05/05/2020 | India:Modasa |
| MT481909.1 | 05/05/2020 | India:Modasa |
| MT483553.1 | 05/05/2020 | India:Modasa |
| MT483554.1 | 05/05/2020 | India:Modasa |
| MT483555.1 | 05/05/2020 | India:Modasa |
| MT483556.1 | 05/05/2020 | India:Modasa |
| MT483557.1 | 05/05/2020 | India:Modasa |
| MT483558.1 | 05/05/2020 | India:Modasa |
| MT483559.1 | 05/05/2020 | India:Prantij |
| MT483560.1 | 05/05/2020 | India:Modasa |
| MT483702.1 | 05/05/2020 | India:Modasa |
| MT496979.1 | 05/05/2020 | India:Ahmedabad |
| MT496983.1 | 05/05/2020 | India:Ahmedabad |
| MT496985.1 | 05/05/2020 | India:Ahmedabad |
| MT496986.1 | 05/05/2020 | India:Ahmedabad |
| MT509508.1 | 05/05/2020 | India:Jamnagar |
| MT512645.1 | 05/05/2020 | USA |
| MT560689.1 | 05/05/2020 | India:Bayad |
| MT612190.1 | 05/05/2020 | Australia:Victoria |
| MT612198.1 | 05/05/2020 | Australia:Victoria |
| MT612216.1 | 05/05/2020 | Australia:Victoria |
| MT628087.1 | 05/05/2020 | USA:CA |
| MT628200.1 | 05/05/2020 | USA:CA |
| MT642094.1 | 05/05/2020 | USA:Washington-Yakima-County |
| MT642099.1 | 05/05/2020 | USA:Washington-Yakima-County |
| MT642101.1 | 05/05/2020 | USA:Washington-Yakima-County |
| MT642112.1 | 05/05/2020 | USA:Washington-Snohomish-County |

| | | |
|---|---|---|
| MT642114.1 | 05/05/2020 | USA:Washington-Snohomish-County |
| MT642118.1 | 05/05/2020 | USA:Washington-King-County |
| MT642131.1 | 05/05/2020 | USA:Washington-Yakima-County |
| MT642166.1 | 05/05/2020 | USA:Washington-Yakima-County |
| MT642173.1 | 05/05/2020 | USA:Washington-Yakima-County |
| MT642192.1 | 05/05/2020 | USA:Washington-Yakima-County |
| MT642196.1 | 05/05/2020 | USA:Washington-Yakima-County |
| MT642211.1 | 05/05/2020 | USA:Washington-Yakima-County |
| MT642214.1 | 05/05/2020 | USA:Washington-King-County |
| MT642225.1 | 05/05/2020 | USA:Washington-Yakima-County |
| MT642232.1 | 05/05/2020 | USA:Washington-Yakima-County |
| MT642289.1 | 05/05/2020 | USA:Washington-Yakima-County |
| MT642306.1 | 05/05/2020 | USA:Washington-Yakima-County |
| MT642333.1 | 05/05/2020 | USA:Washington-Yakima-County |
| MT642358.1 | 05/05/2020 | USA:Washington-Yakima-County |
| MT642407.1 | 05/05/2020 | USA:Washington-Yakima-County |
| MT642429.1 | 05/05/2020 | USA:Washington-Yakima-County |
| MT457400.1 | 05/06/2020 | Netherlands |
| MT457401.1 | 05/06/2020 | Netherlands |
| MT509501.1 | 05/06/2020 | India:Jamnagar |
| MT509504.1 | 05/06/2020 | India:Rajkot |
| MT560673.1 | 05/06/2020 | India:Prantij |
| MT560827.1 | 05/06/2020 | India:Prantij |
| MT612169.1 | 05/06/2020 | Australia:Victoria |
| MT612175.1 | 05/06/2020 | Australia:Victoria |
| MT612192.1 | 05/06/2020 | Australia:Victoria |
| MT612193.1 | 05/06/2020 | Australia:Victoria |
| MT612194.1 | 05/06/2020 | Australia:Victoria |
| MT612219.1 | 05/06/2020 | Australia:Victoria |
| MT620766.1 | 05/06/2020 | USA |
| MT628088.1 | 05/06/2020 | USA:CA |
| MT628089.1 | 05/06/2020 | USA:CA |
| MT628170.1 | 05/06/2020 | USA:CA |
| MT628198.1 | 05/06/2020 | USA:CA |
| MT628199.1 | 05/06/2020 | USA:CA |
| MT642103.1 | 05/06/2020 | USA:Washington-Yakima-County |
| MT642157.1 | 05/06/2020 | USA:Washington-Yakima-County |
| MT642164.1 | 05/06/2020 | USA:Washington-Yakima-County |
| MT642168.1 | 05/06/2020 | USA:Washington-Yakima-County |
| MT642180.1 | 05/06/2020 | USA:Washington-Yakima-County |
| MT642209.1 | 05/06/2020 | USA:Washington-Yakima-County |
| MT642224.1 | 05/06/2020 | USA:Washington-Yakima-County |
| MT642234.1 | 05/06/2020 | USA:Washington-Yakima-County |
| MT642245.1 | 05/06/2020 | USA:Washington-Yakima-County |
| MT642251.1 | 05/06/2020 | USA:Washington-Yakima-County |
| MT642348.1 | 05/06/2020 | USA:Washington-Yakima-County |

| | | |
|---|---|---|
| MT642350.1 | 05/06/2020 | USA:Washington-Yakima-County |
| MT642387.1 | 05/06/2020 | USA:Washington-Yakima-County |
| MT642414.1 | 05/06/2020 | USA:Washington-Yakima-County |
| MT642435.1 | 05/06/2020 | USA:California-Monterey-County |
| MT509499.1 | 05/07/2020 | India:Jamnagar |
| MT601275.1 | 05/07/2020 | Bangladesh:Narayanganj |
| MT607254.1 | 05/07/2020 | Bangladesh:Narayanganj |
| MT612199.1 | 05/07/2020 | Australia:Victoria |
| MT612200.1 | 05/07/2020 | Australia:Victoria |
| MT628171.1 | 05/07/2020 | USA:CA |
| MT628197.1 | 05/07/2020 | USA:CA |
| MT628274.1 | 05/07/2020 | USA:CA |
| MT628275.1 | 05/07/2020 | USA:CA |
| MT628276.1 | 05/07/2020 | USA:CA |
| MT628277.1 | 05/07/2020 | USA:CA |
| MT642117.1 | 05/07/2020 | USA:Washington-Yakima-County |
| MT642134.1 | 05/07/2020 | USA:Washington-Yakima-County |
| MT642140.1 | 05/07/2020 | USA:Washington-Yakima-County |
| MT642183.1 | 05/07/2020 | USA:Washington-Yakima-County |
| MT642213.1 | 05/07/2020 | USA:Washington-Yakima-County |
| MT642227.1 | 05/07/2020 | USA:Washington-Yakima-County |
| MT642228.1 | 05/07/2020 | USA:Washington-Yakima-County |
| MT642282.1 | 05/07/2020 | USA:Washington-Yakima-County |
| MT642325.1 | 05/07/2020 | USA:Washington-Yakima-County |
| MT642332.1 | 05/07/2020 | USA:Washington-Yakima-County |
| MT642345.1 | 05/07/2020 | USA:Washington-Yakima-County |
| MT598178.1 | 05/08/2020 | USA:San-Diego-California |
| MT628196.1 | 05/08/2020 | USA:CA |
| MT628279.1 | 05/08/2020 | USA:CA |
| MT509496.1 | 05/09/2020 | India:Botad |
| MT509498.1 | 05/09/2020 | India:Jamnagar |
| MT509503.1 | 05/09/2020 | India:Junagadh |
| MT601295.1 | 05/09/2020 | Bangladesh:Narayanganj |
| MT607246.1 | 05/09/2020 | Bangladesh:Narayanganj |
| MT612211.1 | 05/09/2020 | Australia:Victoria |
| MT612232.1 | 05/09/2020 | Australia:Victoria |
| MT612236.1 | 05/09/2020 | Australia:Victoria |
| MT612238.1 | 05/09/2020 | Australia:Victoria |
| MT612254.1 | 05/09/2020 | Australia:Victoria |
| MT612260.1 | 05/09/2020 | Australia:Victoria |
| MT628176.1 | 05/09/2020 | USA:CA |
| MT628178.1 | 05/09/2020 | USA:CA |
| MT628280.1 | 05/09/2020 | USA:CA |
| MT509495.1 | 05/10/2020 | India:Kodinar |
| MT509502.1 | 05/10/2020 | India:Jamnagar |
| MT509507.1 | 05/10/2020 | India:Una |

| | | |
|---|---|---|
| MT509510.1 | 05/10/2020 | India:Una |
| MT569471.1 | 05/10/2020 | Serbia:Kraljevo |
| MT601278.1 | 05/10/2020 | Bangladesh:Narayanganj |
| MT601279.1 | 05/10/2020 | Bangladesh:Narayanganj |
| MT601280.1 | 05/10/2020 | Bangladesh:Narayanganj |
| MT607252.1 | 05/10/2020 | Bangladesh:Narayanganj |
| MT607255.1 | 05/10/2020 | Bangladesh:Narayanganj |
| MT612189.1 | 05/10/2020 | Australia:Victoria |
| MT612206.1 | 05/10/2020 | Australia:Victoria |
| MT612221.1 | 05/10/2020 | Australia:Victoria |
| MT612253.1 | 05/10/2020 | Australia:Victoria |
| MT612255.1 | 05/10/2020 | Australia:Victoria |
| MT509497.1 | 05/11/2020 | India:Una |
| MT509958.1 | 05/11/2020 | Bangladesh |
| MT568643.1 | 05/11/2020 | Bangladesh |
| MT578015.1 | 05/11/2020 | USA |
| MT612201.1 | 05/11/2020 | Australia:Victoria |
| MT612202.1 | 05/11/2020 | Australia:Victoria |
| MT612203.1 | 05/11/2020 | Australia:Victoria |
| MT628193.1 | 05/11/2020 | USA:CA |
| MT628194.1 | 05/11/2020 | USA:CA |
| MT628195.1 | 05/11/2020 | USA:CA |
| MT628214.1 | 05/11/2020 | USA:CA |
| MT628215.1 | 05/11/2020 | USA:CA |
| MT628216.1 | 05/11/2020 | USA:CA |
| MT628217.1 | 05/11/2020 | USA:CA |
| MT534306.1 | 05/12/2020 | USA:CA |
| MT534307.1 | 05/12/2020 | USA:CA |
| MT534309.1 | 05/12/2020 | USA:CA |
| MT534319.1 | 05/12/2020 | USA:CA |
| MT534322.1 | 05/12/2020 | USA:CA |
| MT534325.1 | 05/12/2020 | USA:CA |
| MT534326.1 | 05/12/2020 | USA:CA |
| MT534327.1 | 05/12/2020 | USA:CA |
| MT534328.1 | 05/12/2020 | USA:CA |
| MT534330.1 | 05/12/2020 | USA:CA |
| MT534331.1 | 05/12/2020 | USA:CA |
| MT534332.1 | 05/12/2020 | USA:CA |
| MT534333.1 | 05/12/2020 | USA:CA |
| MT534335.1 | 05/12/2020 | USA:CA |
| MT534337.1 | 05/12/2020 | USA:CA |
| MT612209.1 | 05/12/2020 | Australia:Victoria |
| MT612210.1 | 05/12/2020 | Australia:Victoria |
| MT612246.1 | 05/12/2020 | Australia:Victoria |
| MT612247.1 | 05/12/2020 | Australia:Victoria |
| MT612248.1 | 05/12/2020 | Australia:Victoria |

| | | |
|---|---|---|
| MT612250.1 | 05/12/2020 | Australia:Victoria |
| MT612251.1 | 05/12/2020 | Australia:Victoria |
| MT612259.1 | 05/12/2020 | Australia:Victoria |
| MT612261.1 | 05/12/2020 | Australia:Victoria |
| MT628192.1 | 05/12/2020 | USA:CA |
| MT502774.1 | 05/13/2020 | Bangladesh |
| MT576645.1 | 05/13/2020 | Poland |
| MT612220.1 | 05/13/2020 | Australia:Victoria |
| MT628191.1 | 05/13/2020 | USA:CA |
| MT612225.1 | 05/14/2020 | Australia:Victoria |
| MT612241.1 | 05/14/2020 | Australia:Victoria |
| MT612243.1 | 05/14/2020 | Australia:Victoria |
| MT631800.1 | 05/14/2020 | USA:CA |
| MT631802.1 | 05/14/2020 | USA:CA |
| MT631803.1 | 05/14/2020 | USA:CA |
| MT631805.1 | 05/14/2020 | USA:CA |
| MT568645.1 | 05/15/2020 | Morocco:Casablanca |
| MT628190.1 | 05/15/2020 | USA:CA |
| MT560669.1 | 05/16/2020 | India:Himatnagar |
| MT560675.1 | 05/16/2020 | India:Himatnagar |
| MT560677.1 | 05/16/2020 | India:Khedbrahma |
| MT612266.1 | 05/16/2020 | Australia:Victoria |
| MT612267.1 | 05/16/2020 | Australia:Victoria |
| MT612279.1 | 05/16/2020 | Australia:Victoria |
| MT628090.1 | 05/16/2020 | USA:CA |
| MT560694.1 | 05/17/2020 | India:Khedbrahma |
| MT560705.1 | 05/17/2020 | India:Himatnagar |
| MT560706.1 | 05/17/2020 | India:Khedbrahma |
| MT576529.1 | 05/18/2020 | India:Himatnagar |
| MT576532.1 | 05/18/2020 | India:Himatnagar |
| MT612298.1 | 05/19/2020 | Australia:Victoria |
| MT612299.1 | 05/19/2020 | Australia:Victoria |
| MT612301.1 | 05/19/2020 | Australia:Victoria |
| MT612302.1 | 05/19/2020 | Australia:Victoria |
| MT612322.1 | 05/20/2020 | Australia:Victoria |
| MT628091.1 | 05/20/2020 | USA:CA |
| MT628092.1 | 05/20/2020 | USA:CA |
| MT539158.1 | 05/21/2020 | Bangladesh |
| MT539159.1 | 05/21/2020 | Bangladesh |
| MT539160.1 | 05/21/2020 | Bangladesh |
| MT576639.1 | 05/21/2020 | Bangladesh:Dhaka |
| MT628093.1 | 05/21/2020 | USA:CA |
| MT549887.1 | 05/22/2020 | Kenya |
| MT576640.1 | 05/23/2020 | Bangladesh:Dhaka |
| MT576641.1 | 05/23/2020 | Bangladesh:Dhaka |
| MT576642.1 | 05/23/2020 | Bangladesh:Dhaka |

| | | |
|---|---|---|
| MT576643.1 | 05/23/2020 | Bangladesh:Dhaka |
| MT576644.1 | 05/23/2020 | Bangladesh:Dhaka |
| MT576689.1 | 05/23/2020 | Bangladesh |
| MT578016.1 | 05/23/2020 | Bangladesh |
| MT578017.1 | 05/23/2020 | Bangladesh |
| MT560656.1 | 05/24/2020 | India:Ahmedabad |
| MT560657.1 | 05/24/2020 | India:Ahmedabad |
| MT560667.1 | 05/24/2020 | India:Ahmedabad |
| MT560668.1 | 05/24/2020 | India:Ahmedabad |
| MT560670.1 | 05/24/2020 | India:Ahmedabad |
| MT560671.1 | 05/24/2020 | India:Ahmedabad |
| MT560672.1 | 05/24/2020 | India:Ahmedabad |
| MT560674.1 | 05/24/2020 | India:Ahmedabad |
| MT560679.1 | 05/24/2020 | India:Ahmedabad |
| MT560680.1 | 05/24/2020 | India:Ahmedabad |
| MT560682.1 | 05/24/2020 | India:Ahmedabad |
| MT560690.1 | 05/24/2020 | India:Ahmedabad |
| MT560692.1 | 05/24/2020 | India:Ahmedabad |
| MT560693.1 | 05/24/2020 | India:Ahmedabad |
| MT560704.1 | 05/24/2020 | India:Ahmedabad |
| MT576530.1 | 05/24/2020 | India:Ahmedabad |
| MT576531.1 | 05/24/2020 | India:Ahmedabad |
| MT601281.1 | 05/26/2020 | Bangladesh:Chattogram |
| MT601282.1 | 05/26/2020 | Bangladesh:Chattogram |
| MT601283.1 | 05/26/2020 | Bangladesh:Chattogram |
| MT601284.1 | 05/26/2020 | Bangladesh:Chattogram |
| MT612325.1 | 05/26/2020 | Australia:Victoria |
| MT576031.1 | 05/27/2020 | India:Ahmedabad |
| MT576032.1 | 05/27/2020 | India:Ahmedabad |
| MT576033.1 | 05/27/2020 | India:Ahmedabad |
| MT576040.1 | 05/27/2020 | India:Ahmedabad |
| MT576041.1 | 05/27/2020 | India:Ahmedabad |
| MT576042.1 | 05/27/2020 | India:Ahmedabad |
| MT576043.1 | 05/27/2020 | India:Ahmedabad |
| MT576044.1 | 05/27/2020 | India:Ahmedabad |
| MT576045.1 | 05/27/2020 | India:Ahmedabad |
| MT576046.1 | 05/27/2020 | India:Ahmedabad |
| MT576047.1 | 05/27/2020 | India:Ahmedabad |
| MT576048.1 | 05/27/2020 | India:Ahmedabad |
| MT576049.1 | 05/27/2020 | India:Ahmedabad |
| MT576052.1 | 05/27/2020 | India:Ahmedabad |
| MT576053.1 | 05/27/2020 | India:Ahmedabad |
| MT576054.1 | 05/27/2020 | India:Ahmedabad |
| MT576055.1 | 05/27/2020 | India:Ahmedabad |
| MT576056.1 | 05/27/2020 | India:Ahmedabad |
| MT576057.1 | 05/27/2020 | India:Ahmedabad |

| | | |
|---|---|---|
| MT576058.1 | 05/27/2020 | India:Ahmedabad |
| MT576059.1 | 05/27/2020 | India:Ahmedabad |
| MT576060.1 | 05/27/2020 | India:Ahmedabad |
| MT576061.1 | 05/27/2020 | India:Ahmedabad |
| MT594112.1 | 05/27/2020 | India:Ahmedabad |
| MT612327.1 | 05/27/2020 | Australia:Victoria |
| MT635270.1 | 05/27/2020 | India:Gujarat-Ahmedabad |
| MT576025.1 | 05/30/2020 | Spain:ASTURIAS |
| MT576026.1 | 05/30/2020 | Spain:ASTURIAS |
| MT576027.1 | 05/30/2020 | Spain:ASTURIAS |
| MT576028.1 | 05/30/2020 | Spain:ASTURIAS |
| MT576029.1 | 05/30/2020 | Spain:ASTURIAS |
| MT576030.1 | 05/30/2020 | Spain:ASTURIAS |
| MT607253.1 | 05/31/2020 | Bangladesh:Narayanganj |
| MT601276.1 | 06/01/2020 | Bangladesh:Narayanganj |
| MT601277.1 | 06/01/2020 | Bangladesh:Narayanganj |
| MT601285.1 | 06/01/2020 | Bangladesh:Dhaka |
| MT601286.1 | 06/01/2020 | Bangladesh:Dhaka |
| MT601287.1 | 06/01/2020 | Bangladesh:Dhaka |
| MT601288.1 | 06/01/2020 | Bangladesh:Dhaka |
| MT601289.1 | 06/01/2020 | Bangladesh:Dhaka |
| MT601290.1 | 06/01/2020 | Bangladesh:Dhaka |
| MT601291.1 | 06/01/2020 | Bangladesh:Dhaka |
| MT601292.1 | 06/01/2020 | Bangladesh:Dhaka |
| MT601293.1 | 06/01/2020 | Bangladesh:Dhaka |
| MT601294.1 | 06/01/2020 | Bangladesh:Dhaka |
| MT607245.1 | 06/01/2020 | Bangladesh:Dhaka |
| MT607247.1 | 06/01/2020 | Bangladesh:Dhaka |
| MT607248.1 | 06/01/2020 | Bangladesh:Dhaka |
| MT648676.1 | 06/02/2020 | Bangladesh |
| MT607603.1 | 06/03/2020 | India:Ahmedabad |
| MT607604.1 | 06/03/2020 | India:Ahmedabad |
| MT607605.1 | 06/03/2020 | India:Ahmedabad |
| MT607606.1 | 06/03/2020 | India:Ahmedabad |
| MT607607.1 | 06/03/2020 | India:Ahmedabad |
| MT607608.1 | 06/03/2020 | India:Ahmedabad |
| MT607609.1 | 06/03/2020 | India:Ahmedabad |
| MT607611.1 | 06/03/2020 | India:Ahmedabad |
| MT607614.1 | 06/03/2020 | India:Ahmedabad |
| MT607617.1 | 06/03/2020 | India:Ahmedabad |
| MT607619.1 | 06/03/2020 | India:Ahmedabad |
| MT607620.1 | 06/03/2020 | India:Ahmedabad |
| MT607621.1 | 06/03/2020 | India:Ahmedabad |
| MT608648.1 | 06/03/2020 | India:Ahmedabad |
| MT635269.1 | 06/03/2020 | India:Gujarat-Ahmedabad |
| MT635271.1 | 06/03/2020 | India:Gujarat-Ahmedabad |

| | | |
|---|---|---|
| MT635272.1 | 06/03/2020 | India:Gujarat-Ahmedabad |
| MT635328.1 | 06/03/2020 | India:Gujarat-Ahmedabad |
| MT635392.1 | 06/03/2020 | India:Gujarat-Vadodara |
| MT635393.1 | 06/03/2020 | India:Gujarat-Vadodara |
| MT607244.1 | 06/05/2020 | India:Gandhinagar |
| MT607249.1 | 06/05/2020 | India:Kadi |
| MT607250.1 | 06/05/2020 | India:Kadi |
| MT607251.1 | 06/05/2020 | India:Gandhinagar |
| MT607601.1 | 06/05/2020 | India:Gandhinagar |
| MT607602.1 | 06/05/2020 | India:Kalol |
| MT607610.1 | 06/05/2020 | India:Gandhinagar |
| MT607612.1 | 06/05/2020 | India:Gandhinagar |
| MT607613.1 | 06/05/2020 | India:Kalol |
| MT607615.1 | 06/05/2020 | India:Vadodara |
| MT607616.1 | 06/05/2020 | India:Kalol |
| MT607618.1 | 06/05/2020 | India:Vadodara |
| MT635339.1 | 06/05/2020 | India:Gujarat-Vadodara |
| MT635391.1 | 06/05/2020 | India:Gujarat-Vadodara |
| MT635397.1 | 06/05/2020 | India:Gujarat-Vadodara |
| MT635403.1 | 06/05/2020 | India:Gujarat-Vadodara |
| MT635406.1 | 06/05/2020 | India:Gujarat-Vadodara |
| MT635408.1 | 06/05/2020 | India:Gujarat-Vadodara |
| MT635856.1 | 06/05/2020 | India:Gujarat-Dhanera |
| MT635672.1 | 06/07/2020 | Bangladesh |
| MT635404.1 | 06/08/2020 | India:Gujarat-Vadodara |
| MT635405.1 | 06/08/2020 | India:Gujarat-Vadodara |
| MT635407.1 | 06/08/2020 | India:Gujarat-Vadodara |
| MT635409.1 | 06/08/2020 | India:Gujarat-Nadiad |
| MT635410.1 | 06/08/2020 | India:Gujarat-Vadodara |
| MT635855.1 | 06/08/2020 | India:Gujarat-Mahemdavad |
| MT635857.1 | 06/08/2020 | India:Gujarat-Mahemdavad |
| MT635858.1 | 06/08/2020 | India:Gujarat-Nadiad |
